# Cell-state–dependent responses to PLK1 inhibition reveal a non-canonical microtubule-endolysosomal vulnerability in quiescent leukemia stem cells

**DOI:** 10.64898/2026.08.25.746864

**Authors:** Qiang Liu, Milena Gojsevic, Angelica Varesi, Amit Subedi, Changjiang Xu, Faith Au Yeung, Bailey Dinel, Nathan Mbong, Liqing Jin, Amanda Mitchell, Cassie Lim, Pepijn J. Janssen, Helena Boutzen, Andrea Arruda, Mark D. Minden, Eric R. Lechman, Steven M. Chan, Gary D. Bader, Brian Raught, Kerstin B. Kaufmann, Jean C.Y. Wang

## Abstract

Relapse in acute myeloid leukemia (AML) is frequently driven by therapy-resistant quiescent leukemia stem cells (LSCs). We show that inhibition of the mitotic master regulator Polo-like kinase 1 (PLK1), a kinase extensively pursued for antiproliferative therapy, unexpectedly eradicates quiescent LSCs through a mechanism distinct from its canonical mitotic function. In proliferating AML cells, PLK1 inhibition (PLK1i) induced G2/M arrest and mitotic catastrophe. In contrast, quiescence-enriched LSCs underwent apoptosis independent of mitotic arrest, revealing a cell-state-dependent mode of drug action. Mechanistically, PLK1i perturbed vesicle trafficking and endolysosomal homeostasis in LSC by disrupting a previously unrecognized PLK1-MAP1A interaction, leading to apoptotic cell death. Combinatorial pharmacologic perturbation studies established microtubule regulation as a critical determinant of quiescent LSC survival, while ex vivo and in vivo assays demonstrated depletion of functionally-defined LSC following PLK1i. These findings identify non-canonical effects of PLK1i in LSC and uncover a targetable vulnerability linked to microtubule and endolysosomal homeostasis.

**Statement of significance:** Conventional chemotherapy has been designed to target proliferating cells and is generally presumed to be ineffective against non-cycling cells. This study demonstrates that microtubule-targeting agents and PLK1 inhibitors can eradicate both cycling leukemic blasts and quiescent LSCs through distinct, cell state–dependent mechanisms, challenging proliferation-centric models of chemotherapy action.

## Introduction

Cancer is fundamentally a disease of abnormal and uncontrolled cell growth, and for decades anti-proliferative chemotherapy has proven effective in reducing tumor burden and providing a cure for some patients. However, these treatments were developed prior to the recognition that many cancers are functionally heterogeneous, where growth is sustained by a typically rare subset of cells with stem-like properties at the apex of a cellular hierarchy.(1) These cancer stem cells (CSC), like normal stem cells, are often quiescent and slow cycling, and hence, not effectively eliminated by conventional drugs. (2–6) Thus, while some cancers exhibit high cure rates with traditional anti-proliferative therapies, post-treatment persistence of CSCs driving metastasis and disease relapse has been a significant barrier to cure in patients whose cancers are hierarchically-organized. (7–13) The reason for this dichotomy likely lies in intrinsic biological differences among various cancer types, but it may also reflect an incomplete understanding of how anti-proliferative therapies act across distinct cellular states, particularly within quiescent stem cell populations.

Acute myeloid leukemia (AML) is a hierarchically-organized cancer in which a small population of leukemia stem cells (LSC) drives the disease and is responsible for relapse after treatment.(8,14,15) While patient-derived xenograft (PDX) models are the most stringent way to examine LSC properties in human AML,(8) this labor-intensive assay is not amenable to high throughput approaches to identify druggable vulnerabilities. We established a scalable, multi-parameter, high-throughput screen of 1200 bioactive compounds (including metabolic inhibitors and classic anti-proliferative drugs) that integrates LSC biology in surrogate readouts, thereby enabling identification of molecules that directly antagonize human LSC properties. As proof of principle, this approach led to identification of NAMPT inhibitors as LSC-targeting drugs.(16) Using the same approach, we identified PLK1 as a regulator of LSC survival, prompting investigation of its function in quiescent stem cells.

The serine/threonine-protein kinase Polo-like kinase 1 (PLK1) is well known for its role as a master regulator of mitosis, and is involved in control of centrosome maturation, spindle assembly, and microtubule attachment to kinetochores. The regulation of its catalytic activity throughout cell-cycle progression is a complex spatial and temporal interplay of autoinhibition, conformational changes, oligomerization, localization and stabilization involving a broad interactome.(17,18) PLK1 overexpression is observed in a number of solid tumors, and hypersensitivity to PLK1i has been reported in cytogenetically complex AML.(19,20) Hence, various ATP-competitive and allosteric inhibitors have been tested in preclinical and clinical trials for their anti-proliferative effects against a variety of cancers including metastatic colorectal cancer, small-cell lung cancer, pancreatic cancer and AML, with variable results.(21) For example, the ATP-competitive PLK1 inhibitor volasertib (also known as BI-6727) in combination with low-dose cytarabine achieved 30% complete remission across several AML phase I to III trials; however, potential benefits were offset by infection-related mortality in non-responders, shifting the focus of recent research to identifying predictive biomarkers of response and adverse effects.(22–26)

Of note, drugs are often moved to human trials without a clear understanding of the mechanism of action, especially in the relevant disease-driving stem cell populations, which can cause effective drugs to be prematurely discarded.(1) Recently, deeper structural insights into PLK1 and the conformational perturbations induced by certain ATP-competitive inhibitors including volasertib suggest these drugs may also promote or mitigate PLK1 non-catalytic functions by altering protein-protein interactions and thus might thereby also exert effects on non-mitotic, quiescent cells.(18,27,28) Indeed, PLK1i was shown to increase PLK1 binding to and acetylation of microtubules in interphase MCF-7 cells (29) and other cell cycle-independent and non-catalytic functions of PLK1 have been reported in specific contexts.(21,30–33) However, whether PLK1 performs cell cycle–independent functions in quiescent stem cells remains unknown.

Multiple cellular processes rely on microtubule dynamics and their stabilization has been shown to be crucial for coordination of endolysosomal trafficking and autophagosome-lysosome fusion.(34,35) Not surprisingly, common anti-cancer drugs that induce mitotic cell death not only interfere with microtubule dynamics during mitosis but are also effective blockers of vesicle fusion (vinblastine, nocodazole).(36) Paclitaxel, a microtubule stabilizing agent, and endogenous microtubule stabilizing proteins like microtubule associated protein 1A (MAP1A) further showcase the reliance of endolysosomal trafficking on stabilized microtubules e.g. for coordinating receptor internalization and degradation or their re-routing back to the cell surface.(37–39) We previously showed that endolysosomal activity and autophagy control normal human hematopoietic stem cell (HSC) quiescence and self-renewal(40,41), although their role in leukemia is complex and appears to be highly context-specific.(42) Enhanced autophagy has been shown either to promote leukemia cell death or to induce pro-survival mechanisms endowing chemoresistance.(42) Similarly, PLK1i has been shown either to mitigate or promote autophagy, depending on the cell line or model studied.(43–47) None of these findings have thus far been explored in the context of quiescent CSC. Hence, while PLK1 inhibitors were originally developed as antimitotic agents, whether their biological effects are uniform across distinct AML cell states remains unclear.

Here, we applied a stemness-based approach integrating primary AML samples, PDX models, and mechanistic studies in hierarchical human AML cell models to investigate PLK1 function in quiescent LSCs. We identify a previously unrecognized PLK1–MAP1A axis that contributes to microtubule-dependent vesicle trafficking and endolysosomal homeostasis, thereby linking cytoskeletal organization to cell-state–dependent LSC survival.

## Results

### Drug screen identifies PLK1 inhibitors as candidate LSC-depleting compounds

To identify small molecule antagonists of LSC activity, we employed a high-throughput screen of a library of 1219 curated bioactive small molecules targeting a wide variety of cellular processes and pathways. Hits were identified based on their ability to deplete primitive CD34^+^CD38^−^ cells in a hierarchical AML cell culture model (OCI-AML8227) (Fig 1A; Suppl. Table 1).(16,48–50) In contrast to traditional cell lines, functional LSCs in OCI-AML8227, as assayed by xenotransplantation, are restricted to the CD34^+^CD38^−^ fraction.(48) Interestingly, all 4 selective ATP-competitive PLK1 inhibitors included in the compound library (BI-2536, volasertib, onvansertib, GSK-461364) were top hits in the LSC-directed screen (Fig. 1B, Suppl. Table 1), an unexpected finding given their well-known role in disrupting the cell cycle in proliferating cells. Of note, a dual PLK1/PLK3 inhibitor (GW-843682X) and a non-ATP competitive, highly promiscuous PLK1 inhibitor (rigosertib)(51) did not selectively target CD34⁺CD38⁻ cells consistently at nM concentrations (Suppl. Table 1). PLK1 is expressed in all phases of the cell cycle and can be activated by Aurora kinases, key cell cycle regulators, to initiate G2/M transition.(52) However, 14 of 16 selective Aurora kinase A (AURKA) inhibitors in the drug library showed no consistent LSC targeting potential (Fig. 1B, Suppl. Table 1), suggesting that LSC depletion by ATP-competitive PLK1 inhibitors was via a cell cycle-independent mechanism.

**Figure 1:**
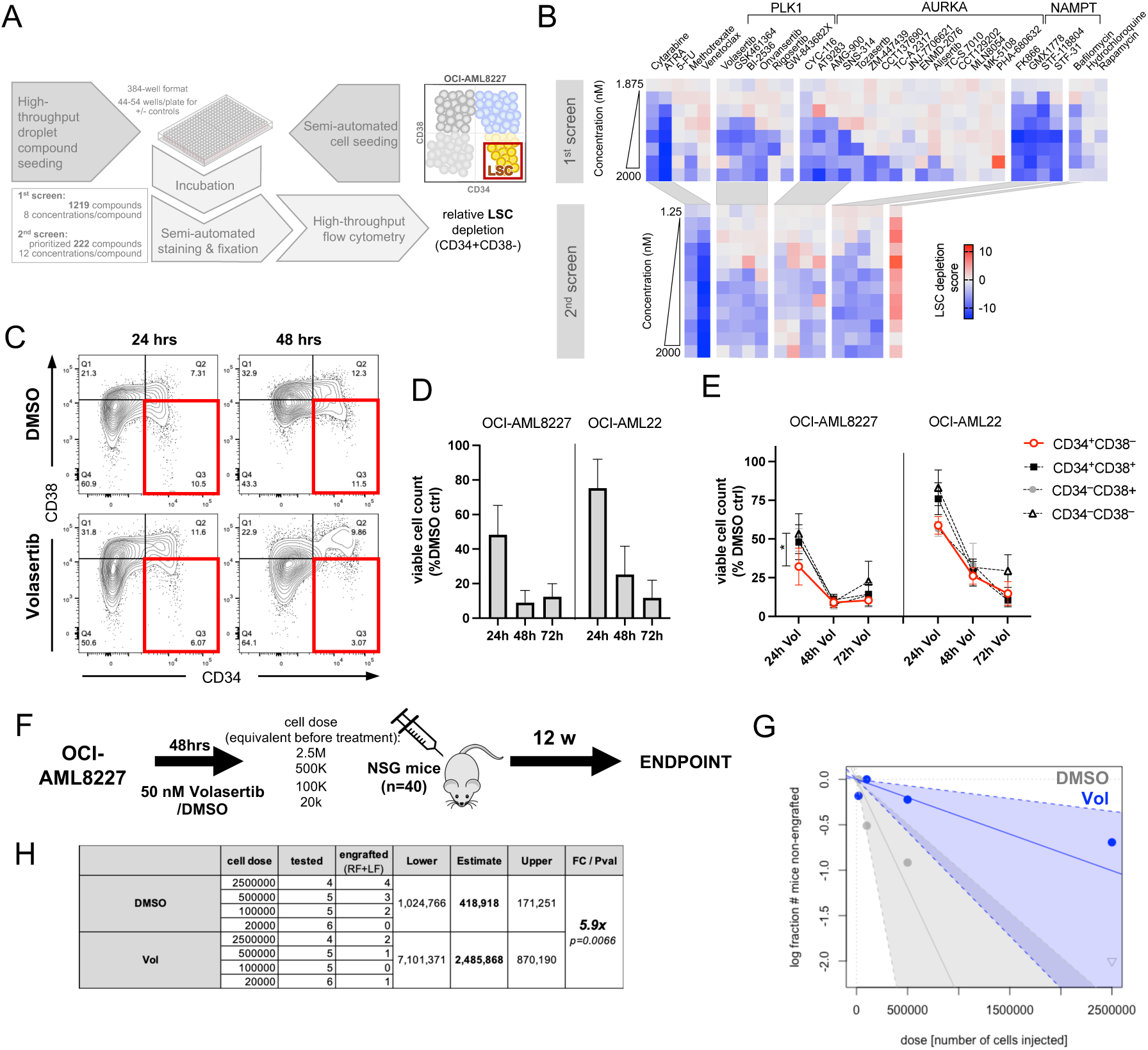
Drug screen identifies PLK1 inhibitors as candidate LSC depleting compounds. A) Schematic outline of high throughput leukemia stemness drug screen B) Heatmap of top scoring drug candidates across 12 concentrations according to flow cytometric analysis and relative depletion of LSC-containing CD34+CD38-fraction compared to DMSO controls is shown. C) Representative flow cytometric analysis of viable, AnnexinV-OCI-AML8227 cells to showcase gating and depletion of CD34+CD38^-^ cell population upon volasertib (50 nM) treatment. D) Absolute total viable cell count as % of DMSO control for indicated time points of treatment (50nM; n=2-6 per time point and model). E) Absolute viable cell counts of sub-gated (CD34/CD38) fractions as % of corresponding fraction of DMSO control for indicated time points of treatment (50nM; n=2-6 per time point, population and model). F) Experimental outline of limiting dilution xenotransplantation assay (LDA) into NSG mice of *in vitro* volasertib treated OCI-AML8227. G) ELDA analysis of LDA results.(90) H) Engraftment scoring (>0.1% hCD45^++^ in right/injected(RF) and left femora(LF)) and statistics according to ELDA analysis.

In validation studies, treatment of OCI-AML8227 cells for 72 hrs *in vitro* with 3 different PLK1 inhibitors at 10-100 nM reduced bulk viability as well as the proportion of CD34⁺CD38⁻ cells within the total viable cell compartment (Fig. 1C-E, Supplementary Fig. 1A), demonstrating activity against both rapidly- and slowly-proliferating cells. Detailed analysis of volasertib drug response curves and time course experiments revealed that the LSC-enriched CD34^+^CD38^−^fraction was depleted at a similar or slightly faster rate than other OCI-AML8227 subpopulations (Fig. 1E, Supplementary Fig. 1B). These results were confirmed with an independent patient-derived AML cell model (OCI-AML22) that similarly maintains a phenotypic and functional cellular hierarchy during culture (Fig. 1D, E, Supplementary Fig. 1C, D).(53,54)

To assess the functional impact and underlying mechanism of action of pharmacological PLK1i specifically on LSC, we selected volasertib as the principal compound for detailed studies, as it represents the most extensively characterized and clinically evaluated PLK1 inhibitor in AML. To quantify effects on LSC, bulk OCI-AML8227 cells were treated with volasertib (50 nM) or DMSO for 48 hrs in culture followed by transplantation into NOD.Cg-*Prkdc^scid^Il2rg^tm1Wjl^*/SzJ (NSG) mice at limiting dilution according to the initial cell dose treated (n=40; 4 cell doses; Fig. 1F). Analysis of engraftment after 12 weeks revealed a 5.9-fold reduction in LSC frequency following *ex vivo* volasertib vs DMSO treatment (\*\**p=0.007,* Fig. 1G, H). Together, these *in vitro* and *ex vivo* findings strongly suggest that PLK1i reduces stemness by depleting functional LSC.

### PLK1i depletes functional LSC *in vivo*

We next assessed whether PLK1i targets LSCs *in vivo* using a cohort of AML patient samples (n=9) comprising multiple subtypes that had been previously assayed in PDX models (Suppl. Table 2).(55,56) NSG mice were treated with vehicle or volasertib (10 mg/kg twice weekly) starting 2-4 weeks after intrafemoral transplantation of AML cells (n=3-8 mice/treatment group for each patient sample, total of 148 mice; Fig. 2A). Human CD45^+^CD33^+^ leukemic cell engraftment in the injected femur (IF) and dissemination to non-injected bone marrow (BM) were analyzed by flow cytometry after 4 weeks of treatment. Volasertib treatment significantly reduced leukemic engraftment compared to vehicle in 7 of 9 PDX models in either the IF or the non-injected BM (Fig. 2B, C), and for 4 samples in both compartments (Fig. 2D).

**Figure 2:**
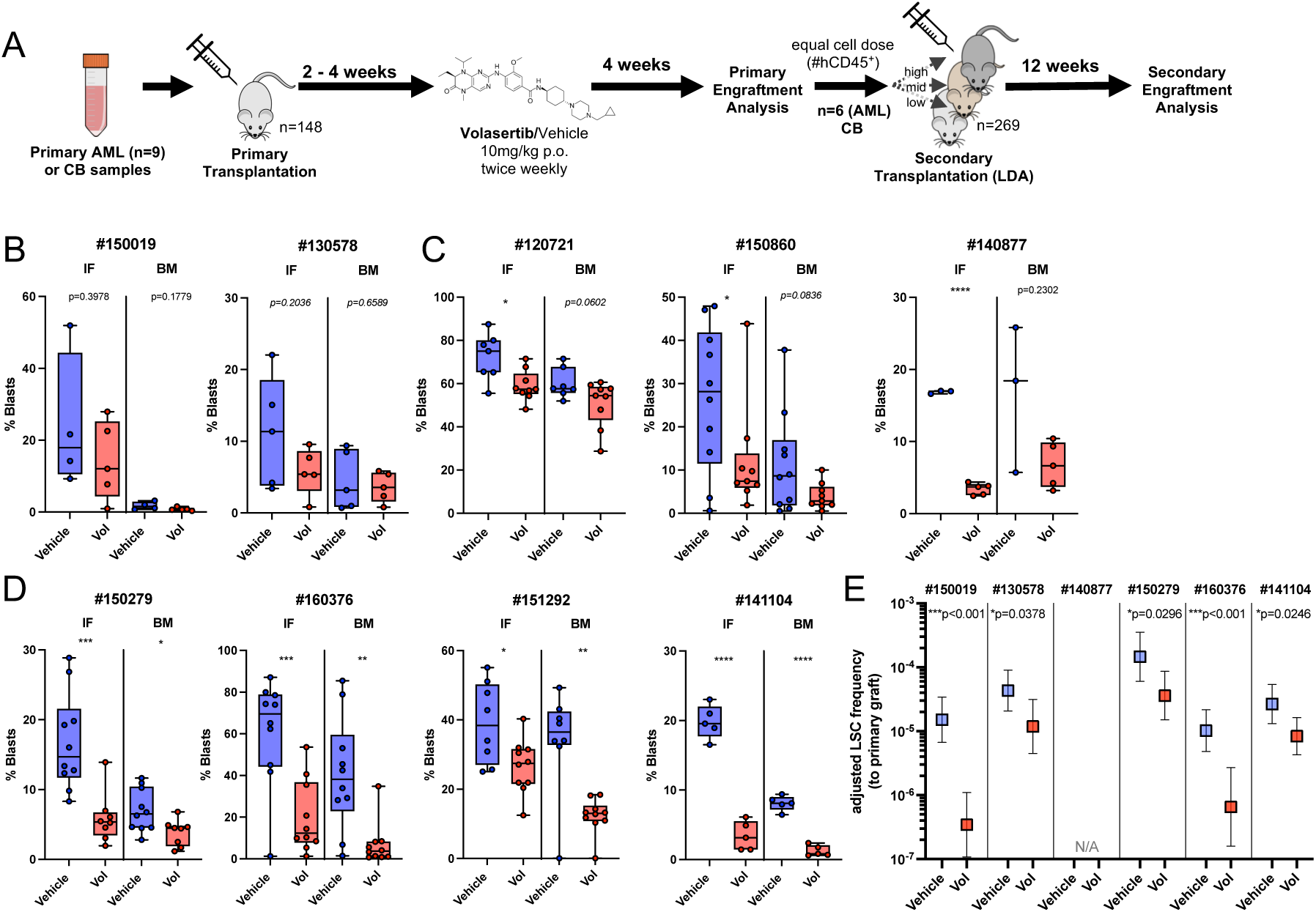
PLK1i depletes functional LSC *in vivo*. A) Experimental outline of in vivo volasertib treatment with serial xenotransplantation. B-D) Flow cytometric analysis (%hCD45+CD33+/Blasts) of PDX after DMSO/Volasertib in vivo treatment with AML samples showing no significant response in primary xenografts (B), with a significant response in the injected femur(IF), with significant responses in IF and non-injected bone marrow (BM; D). E) Summary of secondary LDA according to ELDA analysis. No graft was detected for DMSO or volasertib for sample #140877.

To quantify depletion of LSC in primary treated mice, we performed secondary limiting dilution assays (LDA; additional 12 weeks) with remaining viable human CD45^+^CD33^+^ cells sorted from the primary xenografts of the 2 non-responder samples and 4 of the responders, transplanted as pools at matching cell doses (n= 200 NSG mice total; Fig. 2A, E). After correcting transplanted cell doses for the mean engraftment levels in primary recipients, there was a 3.2-to 43.8-fold reduction in LSC frequency after volasertib treatment in all 5 evaluable samples (Fig. 2E, Suppl. Table 3), including AML#1500019 that was classified as a non-responding sample in the primary PDX assay (Fig. 2B). Collectively, our PDX studies across a diverse cohort of AML samples highlight the potential of PLK1i for targeting LSC.

### Cell state determines cell death pathways induced by PLK1i

As PLK1 inhibitors are generally categorized as a classic anti-proliferative drugs, we wanted to understand if they are acting through different mechanisms in proliferating vs quiescent cells. After 48 hrs of volasertib treatment, 30-35% of remaining viable (PI^−^) cells across all CD34/CD38-defined populations of OCI-AML8227 showed mitotic catastrophe (Fig. 3A, B), with mono-polar spindles characteristic of PLK1i in proliferating cells(57) and increased DAPI intensity consistent with 4n DNA content (Fig. 3A, C).

**Fig. 3:**
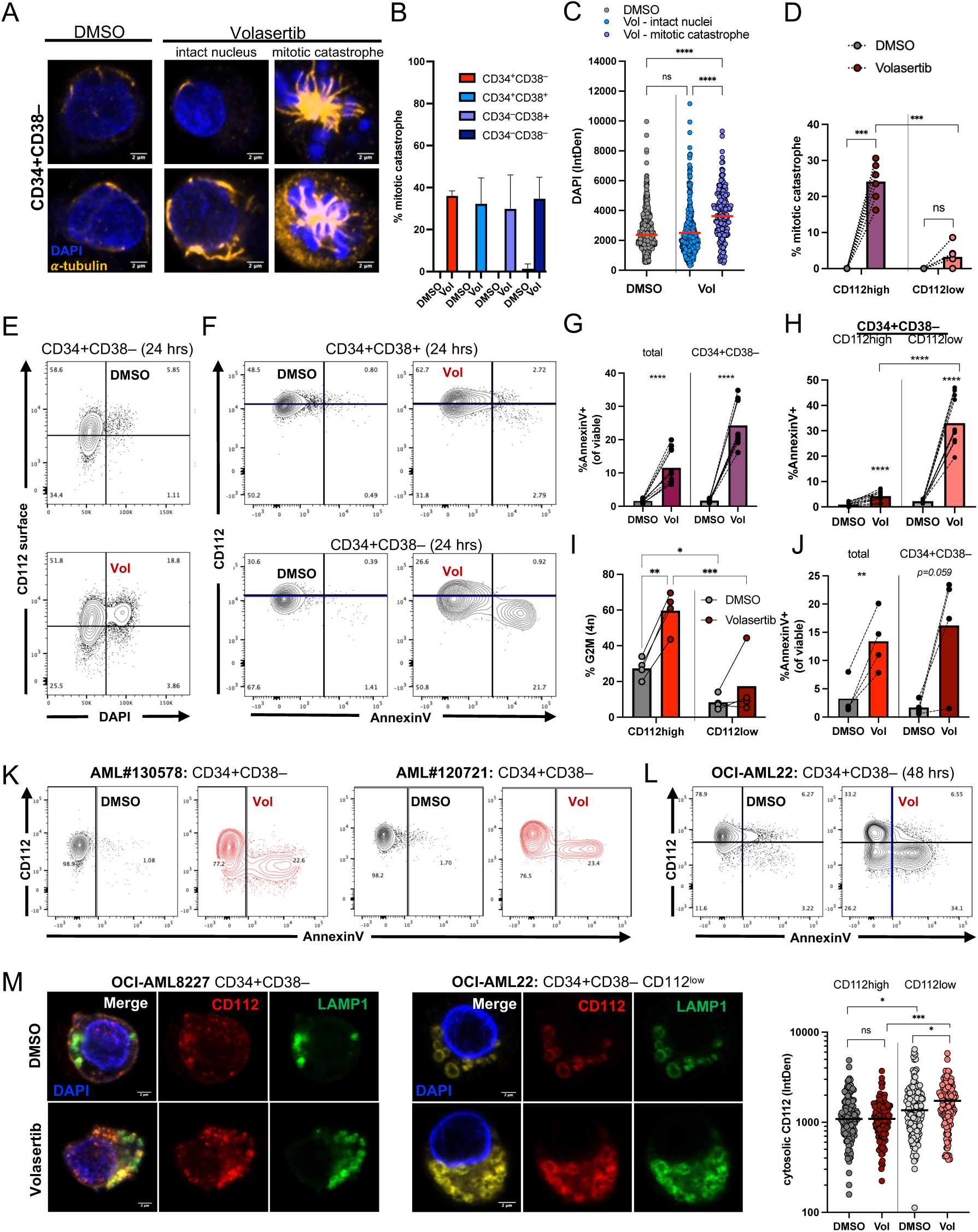
Cell state determines cell death pathways induced by PLK1i. A. Confocal analysis of CD34_+_CD38^-^ OCI-AML8227 with DAPI/DNA and alpha-tubulin staining after DMSO/volasertib treatment (48 hrs). B. Quantification of mitotic catastrophe in 4 OCI-AML8227 fractions sorted according to CD34/CD38 surface expression (confocal imaging; n=3; ∼ 50 cells per fraction and cultured batch). C. DAPI intensity for DMSO and volasertib treated OCI-AML8227 cells (all 4 fractions combined, n=3) with the latter stratified according to mitotic catastrophe scoring vs. intact nucleus scoring. D. Mitotic catastrophe scoring in prospectively isolated and 24 hrs treated CD34_+_CD38^-^ CD112^high^ and CD112^low^ OCI-AML22 cells (n=6; 0 50 cells per fraction and cultured batch). E. Flow cytometric analysis of 24 hrs DMSO/50 nM volasertib treated, surface antibody stained, fixed, permeabilized and DAPI stained OCI-AML22. Only AnnexinV^-^CD34^+^CD38^-^ subpopulations are shown. F. Representative flow plots of 24 hrs DMSO/50 nM volasertib treated OCI-AML22 cells subgated for PI^-^ CD34^+^CD38^+^ and PI^-^CD34^+^CD38^-^ cell populations. G. Flow cytometric analysis of %AnnexinV^+^ among total viable and CD34^+^CD38^-^ subgated OCI-AML22 cells after 24 hrs of DMSO/Volasertib treatment (50 nM, n=12) H. %AnnexinV^+^ cells among subgated CD112^high^ and CD112^low^ CD34^+^CD38^-^ OCI-AML22 cells after 24 hrs of DMSO/volasertib treatment (n=12). I. DNA content analysis by flow cytometry of cultured and 48 hrs DMSO/volasertib treated AML patient samples (n=4) that were surface antibody stained, fixed, permeabilized and DAPI stained. J. Flow cytometric analysis of %AnnexinV^+^ among total viable and CD34^+^CD38^-^ subgated cells of 4 primary AML patient samples after 24 hrs of DMSO/volasertib treatment (50 nM, n=4) following one day of culture after thawing. K. Flow cytometric analysis of indicated AML patient samples (n=2) after 48 hrs DMSO/50 nM volasertib treatment that were subgated for PI^-^CD34^+^CD38^-^ populations. L. Representative flow plot of DMSO/volasertib treated PI^-^CD34^+^CD38^-^ OCI-AML22 cells after 48 hrs of treatment. M. Confocal analysis of prospectively isolated and 24 hrs DMSO/volasertib treated LSC-enriched populations of OCI-AML8227 (CD34^+^CD38^-^) and OCI-AML22 (CD34^+^CD38^-^CD112^high^ and CD112^low^; quantification on right from 0 50 cells per batch from 3 individually cultured batches) as indicated.

In the OCI-AML8227 model, less than 20% of CD34^+^CD38^−^ cells are in G_0_ (Ki-67^−^).(48,49) We previously showed that two distinct quiescent populations can be isolated within the human long-term HSC compartment based on the top and bottom 20% of CD112 surface expression levels, with the most deeply quiescent (CDK6^−^) cells enriched in the CD112^low^ fraction.(58) However, G_0_ cells were only modestly enriched in the CD34^+^CD38^−^CD112^low^ fraction of OCI-AML8227 (Supplementary Fig. 2A,B); hence this model was suboptimal for dissecting mechanisms specific to quiescent leukemic cells. Even so, short (24 hrs) volasertib treatment of OCI-AML8227 induced an AnnexinV^+^ population that was exclusively CD112^low^ and an increase in the proportion of viable, AnnexinV^−^ cells with high CD112 expression in the CD34^+^CD38^−^ compartment, pointing to rapid, selective apoptosis and depletion of the most quiescent cells (Supplementary Fig. 2C-F).

For more precise evaluation of cell-state dependent effects in quiescence-enriched LSC populations, we turned to the OCI-AML22 model, in which we previously showed that CD112^low^ cells in the CD34^+^CD38^−^ fraction exhibit slower engraftment kinetics, greater cytarabine resistance, and a markedly higher proportion of G_0_ (75% vs. 55%) and deeply quiescent CDK6^−^ cells (60% vs. 10%) compared to CD112^high^ cells.(53) The actively-cycling G2/M fraction makes up 21% of total OCI-AML22 cells. Within the CD34^+^CD38^−^ population, 8% of CD112^high^ are in G2/M, compared to 2% of CD112^low^ cells (Supplementary Fig. 2G). Consistent with these observations, volasertib induced mitotic catastrophe almost exclusively in prospectively-isolated CD112^high^ but not in CD112^low^ CD34^+^CD38^−^ OCI-AML22 cells (Fig. 3D). Similarly, CD112^high^ CD34^+^CD38^−^ cells accumulated in G2/M with 4n DNA content more than CD112^low^ cells (Fig. 3E; Supplementary Fig. 2G). In contrast, short (24 hrs) volasertib treatment induced an AnnexinV^+^ population marked by low CD112 expression exclusively within the LSC-enriched CD34^+^CD38^−^ fraction and not among CD34^+^CD38^+^ progenitors (Fig. 3F-H). Importantly, these findings were recapitulated using both the highly PLK1-selective ATP-competitive inhibitor onvansertib and the structurally-distinct allosteric inhibitor rigosertib, supporting PLK1 as the relevant target and suggesting that the observed phenotype is not restricted to a single inhibitor class or mode of PLK1 perturbation (Supplementary Fig. 2H,I).

We next extended this analysis to 4 AML patient samples with functional LSC content established by xenografting.(4,54) Volasertib-induced mitotic arrest based on DNA content was significantly more pronounced in the CD112^high^ vs CD112^low^ CD34^+^CD38^−^ subfractions (Fig. 3I, Supplementary Fig. 2J, Suppl. Table 2). In contrast, volasertib treatment increased the proportion of AnnexinV^+^ cells within the CD34^+^CD38^−^ compartment in 3 of 4 samples (Fig. 3J) to a greater extent than in the bulk population. Indeed, AnnexinV^+^ cells were exclusively CD112^low^ or CD112^−^ in 2 of these samples (Fig. 3K, Supplementary Fig. 2K), one of which (AML#130578) has been shown to be enriched for deeply quiescent (CDK6^−^Ki67^−^) cells in the CD34^+^CD38^−^CD112^low^ fraction.(53)

Interestingly, volasertib treatment induced a CD112^−^ (below CD112^low^), AnnexinV-negative population in the CD34^+^CD38^−^ fraction in samples AML#130578 and AML#120721 (Fig. 3K). This was also observed after longer volasertib treatment (48 hrs; Fig. 3L) of OCI-AML22 cells in the CD34^+^CD38^−^ but not CD34^+^CD38^+^ progenitor fractions (Supplementary Fig. 2L). Similar to AnnexinV, Caspase 3/7 activity was largely confined to PI^−^CD34^+^CD38^−^CD112^−/low^ cells (Supplementary Fig. 2M), suggesting that PLK1i may promote internalization of CD112 prior to induction of apoptosis. We have previously shown that CD112 surface expression on quiescent vs activated HSC is regulated by endocytosis.(58) Consistent with this, we observed an increase in intracellular CD112 protein abundance in vesicles visualized by the lysosome and endolysosomal marker LAMP1 in sorted CD34^+^CD38^−^CD112^low^ OCI-AML22 and CD34^+^CD38^−^ OCI-AML8227 cells following volasertib treatment but not in the more differentiated cell fractions (Fig. 3M; Supplementary Fig. 2N).

Taken together, our findings suggest that PLK1 inhibitors have dual mechanisms of action, inducing mitotic arrest predominantly in proliferation-primed CD112^high^ LSC and cycling non-LSC, while inducing apoptosis in the quiescent LSC-enriched CD112^low^ fraction.

### A starved PLK1 interactome points to PLK1 regulating endolysosomal trafficking in quiescent cells

To elucidate how PLK1i might induce apoptosis specifically in quiescent cells, we examined the PLK1 interactome via proximity-dependent biotinylation (BioID) in non-mitotic cells (in contrast to prior studies exclusively focusing on its roles in mitotic cells).(59,60) To enrich for non-mitotic PLK1 interactors, we performed BioID under serum starvation of Flp-In T-Rex 293 cells and employed the fast miniTurbo BirA mutant followed by mass spectrometry and protein identification.(60,61) We generated miniTurbo constructs for wild-type PLK1 (wt-PLK1), a constitutively active mutant (ca-T210D) and a kinase dead mutant (kd-K82R)(62,63) and retrieved differential as well as common interactors among the 3 PLK1 constructs (Fig. 4A,B; Suppl. Table 4).

**Figure 4:**
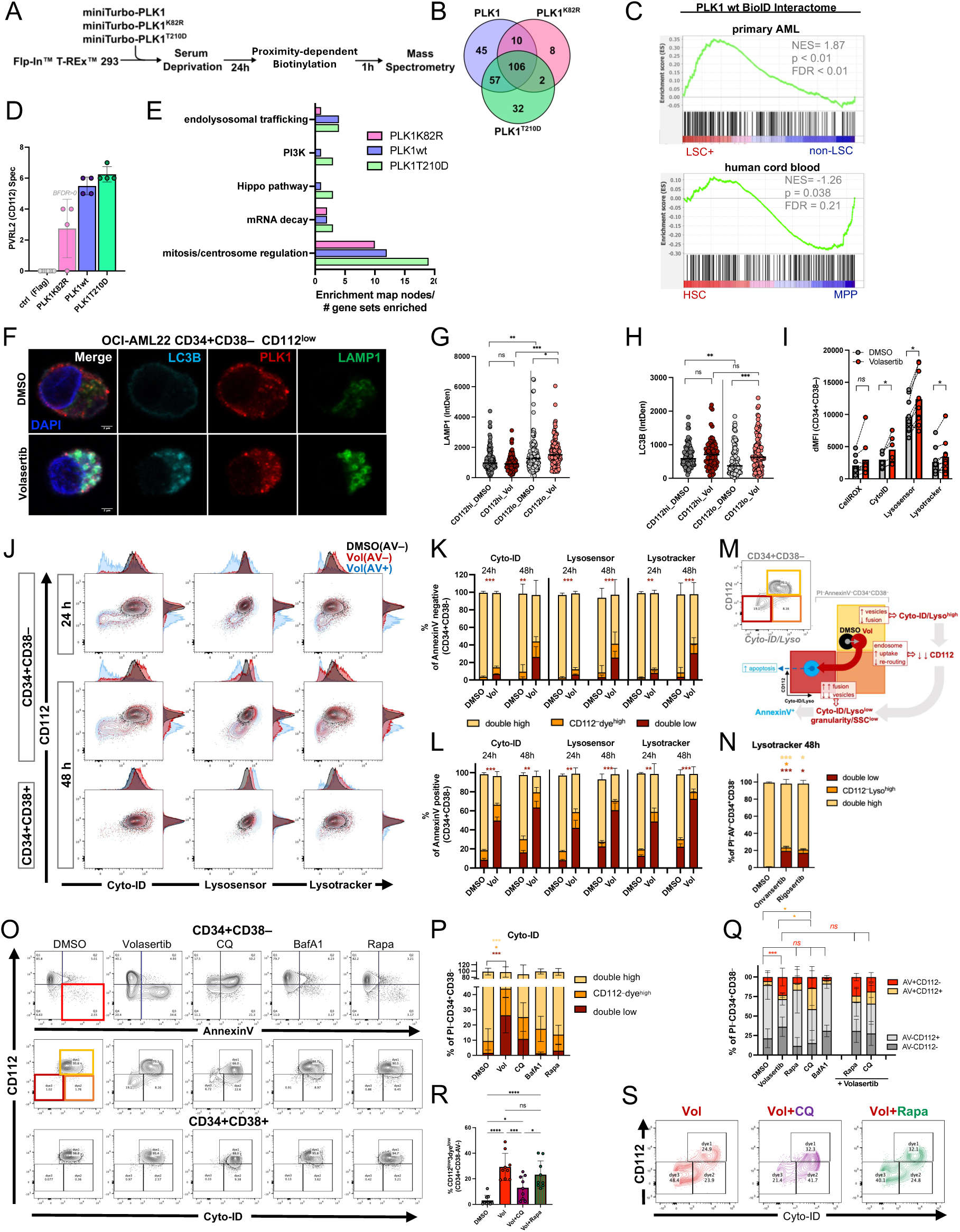
A starved PLK1 interactome points to PLK1 navigating endolysosomal trafficking in quiescent cells. A. Experimental outline of the performed Bio-ID assay. B. Venn-Diagram of unique and shared interactors for PLK1wt, kd-K82R and ca-T210D according to Bio-ID. C. Gene set enrichment analysis (GSEA) using the retrieved PLK1 wt interactome under serum deprivation as gene set on RNAseq expression data of functionally validated primary AML LSC+ vs. non-LSC fractions and human umbilical cord blood HSC and multipotent progenitors (MPP).^4,40^ D. Mass spectrometry results for PVRL2/CD112 (4 measurements from 2 biological and 2 technical replicates each); *BFDR: Bayesian false discovery rate.* E. Pathway enrichment analysis of interactomes from BioID assay according to PLK1 constructs reflective of enrichment map nodes in Suppl. Fig. 3A. F. Confocal imaging of prospectively isolated OCI-AML CD34+CD38-CD112^low^ cells, treated with DMSO/volasertib for 24hrs. G. LAMP1 quantification by confocal imaging of sorted and DMSO/Volasertib 24 hrs treated OCI-AML22 CD34^+^CD38^-^ CD112^high^ and CD112^low^ cells (0 50 cells per batch from 3 individually cultured batches). H. Quantification of LC3B staining in sorted and DMSO/Volasertib 24 hrs treated OCI-AML22 CD34^+^CD38^-^ CD112^low^ and CD112^high^ cells (0 50 cells/batch from 3 individually cultured batches) by confocal imaging. I. Combined flow cytometric analysis of 48 hrs DMSO/Volasertib (50 nM) treated OCI-AML22 and OCI-AML8227 (4 independent assays with 6-7 and 2-3 measurements from independent cultures, respectively). J. Flow cytometric analysis for Cyto-ID, Lysosensor and Lysotracker assays of 24 and 48 hrs DMSO/Volasertib treated PI^-^AnnexinV^-^CD34^+^CD3^8-^ subgated OCI-AML22 cells overlaid with volasertib treated AnnexinV^+^CD34^+^CD38^-^ cells (blue). K. Flow cytometric analysis for Cyto-ID, Lysosensor and Lysotracker assays of 24 and 48 hrs DMSO/Volasertib treated PI^-^AnnexinV^—^CD34^+^CD38^—^OCI-AML22 cells further subgated into double high (CD112^high^3dye^high^), CD112^low^3dye^high^ and double low (n=4) - see gating in scheme on the right. L. Gating scheme from K) applied to AnnexinV+ cells within subgated CD34^+^CD38^-^ population for 24h and 48 hrs treatment for individual dyes as indicated (n=4). M. Schematic drawing of underlying processes induced by volasertib treatment and traced by AnnexinV and CD112 surface expression and vesicle dye signals by flow cytometry. N. Quantification of flow cytometric analysis of OCI-AML22 cells treated for 48 hrs with either DMSO equivalent or 50 nM of the indicated drugs (n=3). O. Flow cytometric analysis of AnnexinV vs CD112 staining on OCI-AML22 PI^-^CD34^+^CD38^-^ subgated cells treated with the indicated drugs for 48 hrs (50 nM volasertib, 60 nM chloroquine=CQ, 20 nM bafilomycin A1=BafA1, 500 nM rapamycin=Rapa) in top row and respective CD112 vs Cyto-ID staining on PI-CD34+CD38- and PI^-^CD34^+^CD38^+^ cells in bottom rows. P. Summary data of n=4 for the middle row of M). PI^-^AnnexinV^+^CD34^+^CD38^-^ cells were subgated for analysis. Q. Summary of flow cytometric subpopulation analysis of 48 hrs single or double drug treated OCI-AML22 PI^-^ CD34^+^CD38^-^ cells as indicated. R. Flow cytometric analysis of %induction of a CD112^low^3dye^low^ population (=dye3; see gate in N) within the PI^-^AnnexinV^-^ CD34^+^CD38^-^ fraction upon indicated single or double drug treatment (50 nM volasertib, 60 nM chloroquine=CQ, 500 nM rapamycin=Rapa) for 48 hrs for all 3 dyes combined from n=3. S. Representative flow plots of 48 hrs treated,subgated OCI-AML22 PI^-^CD34^+^CD38^-^ cells for analysis in R.

In total we identified 260 significant PLK1 interactors, of which only 22 have been previously reported in interactome studies.(64,65) Gene set enrichment analysis (GSEA) showed that our wt-PLK1 interactome is enriched in functionally-validated LSC⁺ vs non-LSC fractions from primary AML samples (n=62, 112 fractions) but not in normal HSC vs progenitor cells (MPP; Fig. 4C), supporting the relevance of our starved PLK1 interactome to LSCs.(4,41) Interestingly, PVRL2 (=CD112=Nectin2) was also among the differentially-retrieved PLK1 interactors (only interacting with wt-PLK1 and ca-T210D; Fig. 4D), suggesting that PLK1 may be directly involved in stabilizing CD112 surface presentation during cell cycle progression (see Fig. 3E). Not surprisingly, pathway enrichment analysis demonstrated that the majority of interactors of all 3 PLK1 constructs are involved in cell cycle-associated processes (Fig. 4E; Supplementary Fig. 3A). Pathways retrieved using wt-PLK1 and ca-T210D but not kd-K82R as bait were related to hippo signaling, endocytosis, and vesicle trafficking, suggesting only catalytically-competent PLK1 interacts with components of these processes.

As we have previously shown that endolysosomal activity and autophagy are important in normal human HSC biology,(40,41) we next assessed these potentially PLK1 inhibitor-sensitive intracellular vesicle trafficking pathways in our two hierarchical AML cell models. Immunostaining of prospectively-isolated subpopulations showed an accumulation of the endolysosomal marker LAMP1 and/or the autophagosome marker LC3B following volasertib treatment specifically in the quiescent LSC-enriched fraction (OCI-AML22 CD34^+^CD38^−^CD112^low^ Fig. 4F-H, OCI-AML8227 CD34^+^CD38^−^ Supplementary Fig. 3B; see also Fig. 3M). Similarly, volasertib treatment of both AML models induced increased levels of Cyto-ID (formation of autophagosomes), Lysosensor (lysosomal acidity) and LysoTracker (abundance of acidic organelles) staining in viable (PI^−^AnnexinV^−^) cells within the CD34^+^CD38^−^compartment without an increase in oxidative stress (CellROX; Fig. 4I). These findings suggest that PLK1i in this stem cell context results in accumulation of autophagosomes and lysosomes, either due to enhanced vesicle formation, reduced vesicle fusion, or both.

In addition to the overall shift to higher fluorescence for Cyto-ID, Lysosensor and LysoTracker, we also noted at the lower end of the fluorescence spectrum of all 3 dyes (3dye^low^) a small population of cells that completely lost CD112 expression (CD112^−^) and was only found in the CD34^+^CD38^−^, but not CD34^+^CD38^+^ fraction (Fig. 4J, Supplementary Fig. 3C). This small AnnexinV^−^3dye^low^CD112^−^ population increased over time (24 vs 48 hrs of treatment; Fig. 4J, K), showed reduced cellular granularity (SSC^low^; Supplementary Fig. 3D) consistent with lower overall intracellular vesicle abundance and had a similar dye/CD112 staining profile as the majority of AnnexinV^+^ volasertib-treated cells (Fig. 4L). In contrast, AnnexinV^+^ CD34^+^CD38^−^control-treated cells and both AnnexinV^+^ and AnnexinV^−^ volasertib-treated CD34^+^CD38^+^ progenitors retained 3dye^high^ staining (Fig. 4L, Supplementary Fig. 3E). Thus, CD34^+^CD38^−^cells appeared to transition through a 3dye^high^ and then a 3dye^low^ state while losing CD112 expression before undergoing apoptosis (AnnexinV^+^). These findings suggest that the initial accumulation of autophagosomes and endolysosomes induced by PLK1i is transient and resolves at later timepoints in still non-apoptotic (AnnexinV^−^) cells (Fig. 4M). Importantly, this sequence of events with induction of a CD34^+^CD38^−^dye^low^CD112^−^ population was also recapitulated in AML patient samples following volasertib treatment (Fig. 3K; Supplementary Fig. 3F) and in OCI-AML22 cells treated with onvansertib or rigosertib (Fig. 3N).

Next, we investigated whether these phenotypes could be recapitulated by pharmacological modulators of the autophagy-endolysosomal systems, namely the classic lysosomal disruptors bafilomycin A1 (BafA1) and chloroquine and the autophagy-promoting drug rapamycin. BafA1 and rapamycin did not induce marked apoptosis in the CD34^+^CD38^−^compartment or impact dye intensities (Fig 4O, Supplementary Fig. 3G). Chloroquine shifted dye intensities in the CD112^high^ compartment to similar levels as volasertib, consistent with stalled vesicle fusion (66), but induced apoptosis in the CD34^+^CD38^−^compartment irrespective of CD112 expression (Fig. 4O-Q). A small Cyto-ID^low^CD112^−^ population appeared after chloroquine treatment in the CD34^+^CD38^−^ compartment but was not significantly different from controls (*p=0.097,* Fig. 4 O, P; Supplementary Fig. 3H). Of note, these drugs and their derivatives were not identified as consistent hits in our initial drug screen (Suppl. Table 1). Together, these findings suggest that the observed initial accumulation of intracellular vesicles following PLK1i cannot be explained by impaired lysosomal acidification (BafA1) or enhanced autophagy induction (rapamycin) alone as seen in other cellular contexts,(44,47) and point to a more complex cascade of events that is not entirely recapitulated by chloroquine-mediated disruption of vesicle fusion and lysosomal function.(66)

To assess whether additional perturbation of vesicle homeostasis can influence the effects of PLK1i, we carried out combination treatment experiments. Addition of chloroquine or rapamycin to volasertib did not alter overall cell counts, %AnnexinV^+^CD112^−^ cells or Cyto-ID staining within the original CD34^+^CD38^−^ population compared to volasertib alone (Fig. 4Q, Supplementary Fig. 3I-L), suggesting that vesicle accumulation induced by volasertib is not further enhanced by either drug in the remaining CD112^high^ population. Importantly, addition of chloroquine but not rapamycin to volasertib-treated cells reduced the emergence of the 3dye^low^CD112^−^ population without altering the concomitant overall loss of CD112 surface expression, consistent with chloroquine stalling vesicle fusion without restoring endosomal receptor (CD112) re-routing (Fig. 4R; S; Supplementary Fig. 3K, L). These findings support the hypothesis that the observed changes in lysosomal dye and CD112 staining induced by PLK1i represent vesicle accumulation due to a transient reduction in autophagosomal-lysosomal fusion (shift of CD112^high^ cells to 3dye^high^ intensities). The vesicle buildup affects endolysosomal homeostasis with continued endosomal uptake (loss of CD112 surface expression in AnnexinV^−^ 3dye^high^ cells) and is then followed by excess vesicle clearance (loss of dye staining and granularity in AnnexinV^−^ CD112^−^ cells; reduced by chloroquine) leading to apoptosis (appearance of AnnexinV^+^ CD112^−^ cells; see schematic Fig. 4M and Figs. 4J-O; Supplementary Fig. 3D, E). Collectively, these studies of the PLK1 interactome and the candidate pathways retrieved point to a distinct, LSC-specific role for PLK1 in regulating vesicular homeostasis and CD112 surface presentation in this quiescent stem cell-enriched context.

### Interference with PLK1-MAP1A interaction disrupts microtubule dynamics in quiescent LSC

To understand how PLK1i triggers the observed changes in the cellular vesicle machinery specifically in LSC, we identified MAP1A as a potential candidate in our BioID data. MAP1A has predominantly been studied in murine neurons where it is known to form a complex with the autophagosome marker and fusion mediator LC3 and its reported function is to stabilize microtubule structures that are essential to vesicle trafficking.(35,67–70) Interestingly, in addition to PLK1’s established role in regulating cytoskeletal changes during mitosis, PLK1i has been found to stabilize microtubules in interphase cells as measured by alpha-tubulin(K40) acetylation, a posttranslational modification known to facilitate vesicle trafficking and ultimately fusion.(29,34)

MAP1A was retrieved as a high confidence interactor only for wt-PLK1 and not ca-T210D (Supplementary Fig. 4A; Suppl. Table 4). As wt-PLK1 only becomes catalytically active upon cell cycle progression, this finding suggests that MAP1A interaction with PLK1 is restricted to interphase or quiescent cells. *MAP1A* expression across multiple functionally-validated LSC datasets derived from AML patient cohorts and our AML cell models showed a positive correlation with LSC frequency and specificity for LSC over normal HSC, and was transcriptionally upregulated at relapse in samples where relapse originated from primitive (ROp) versus committed (ROc) LSC clones (Fig. 5 A-C; Supplementary Fig. 4B-E). *MAP1A* levels were higher in CD112^low^ vs. CD112^high^ OCI-AML22 CD34^+^CD38^−^ cells (Supplementary Fig. 4F).

**Figure 5:**
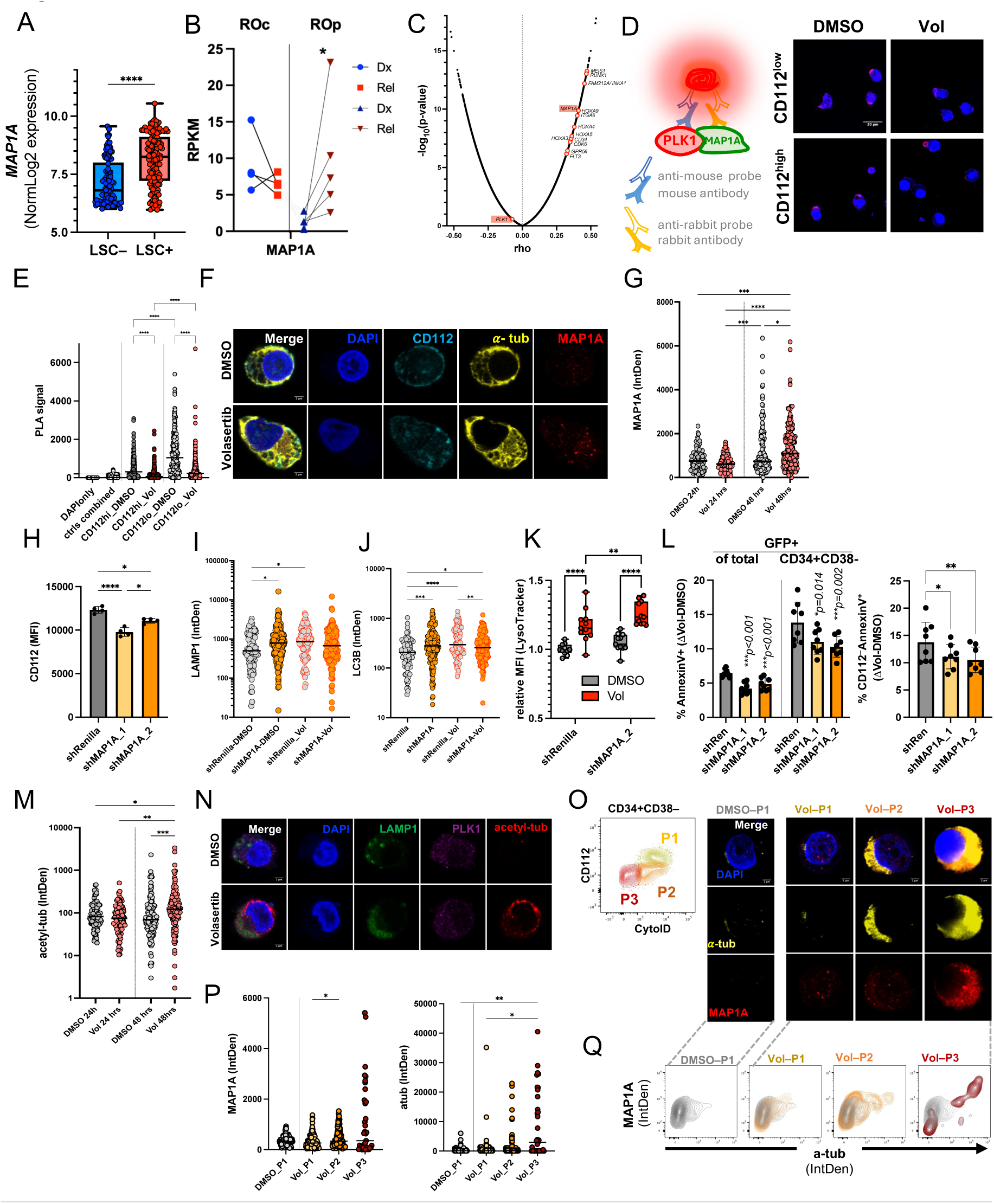
Interference with PLK1-MAP1A interaction disrupts microtubule dynamics in quiescent LSC. A. *MAP1A* expression in sorted and functionally validated LSC^-^ (non-engrafting) and LSC^+^ (engrafting) fractions of primary AML patient samples.(4) B. *MAPI A* expression in paired diagnosis-relapse AML RNAseq samples with inferred Relapse Origin (RO) committed (ROc) or primitive (ROp), either maintaining an already at diagnosis established or acquiring a stem-like bulk transcriptome upon relapse, respectively.(15) C. Spearman Correlation (rho) of gene expression with LSC frequency vs p-value according to functionally (LDA in xenografts) validated LSC^+^ fractions and their transcriptomic profile.(4,48) D. Proximity ligation assay (PLA) to assess interaction between PLK1 and MAP1A with and without volasertib treatment in sorted OCIAML22 CD34^+^CD38^-^ CD112^high^ and CD112^low^ cells. E. Quantification of 5D (161-346 cells in total per condition and population from n=4, total of 375 cells for negative controls analyzed). F. Prospectively isolated OCI-AML22 CD34^+^CD38^-^CD112^low^ cells were treated with DMSO/volasertib for 48 hrs, intracellularly stained for CD112, alpha-tubulin and MAP1A and analyzed by confocal microscopy. G. Quantification of MAP1A levels as shown in F and analogue analysis at 24 hrs of treatment (0 50 cells per batch from 4 individually cultured batches). H. Median fluorescence intensity (MFI) of surface CD112 upon lentivirally delivered, shRNA-mediated MAP1A knockdown vs control (shRenilla) in CD34^+^CD38^-^ OCI-AML22 (subgated on vector co-expressed GFP; 2 independent shRNAs targeting MAP1A shown; n=4; 2 weeks post transduction). I. Confocal analysis for LAMP1 abundance in prospectively sorted and DMSO/volasertib treated GFP^+^CD34^+^CD38^-^ cells in the context of shRenilla or shMAP1A. (shMAP1A_1+_2 combined;0 100 cells from 3 individually transduced and sorted cultures). J. Same as I) for LC3B staining. K. Relative MFI (delta MFI to unstained normalized to delta MFI of DMSO shRenilla) of Lysotracker signal for control and shMAP1A-2 transduced OCI-AML22 CD34^+^CD38^-^ cells 24 hrs after DMSO/volasertib treatment was initiated (4 individually transduced cultures; 3 repetitions; 2 weeks after transduction). L. Flow cytometric analysis of additional % AnnexinV^+^ cells compared to DMSO controls among total P^I-^GFP^+^ and within subgated PI^-^CD34^+^CD38^-^ OCI-AML22 cells upon 50 nM volasertib treatment in the context of shRenilla or shMAP1A_1 or shMAP1A_2 (2 independent shRNAs, 4 individually cultured and sorted batches; 2 repetitions; 2-3 weeks post transduction). Second panel shows additional % CD112^low^AnnexinV+ cells within PI^-^GFP^+^CD34^+^CD38^-^ OCI-AML22 upon 50 nM volasertib treatment vs. DMSO controls. M. Prospectively isolated OCI-AML22 CD34^+^CD38^-^CD112^low^ cells were treated with DMSO/volasertib for 48 hrs. Quantification of acetyl-K40 tubulin levels as shown and analogous analysis at 24hrs of treatment (0 50 cells per batch from 3 individually cultured batches). N. Prospectively isolated OCI-AML22 CD34^+^CD38^-^CD112^low^ cells were treated with DMSO/volasertib for 48 hrs and intracellularly stained for LAMP1, PLK1 and acetyl K40 tubulin and analyzed by confocal microscopy. Quantification in M. O. Confocal images of 48 hrs DMSO/volasertib treated and then sorted OCI-AML22 PI^-^AnnexinV^-^ CD34^+^CD38^-^ cells subgated as indicated in scheme for volasertib into P1, P2, P3 and DMSO only into P1. P. Quantification of MAP1A and alpha-tubulin levels within sorted fractions from O (37-156 cells/images per population combined from 3 individually cultured and sorted batches). Q. Density plots of confocal data shown in O and P to visualize levels of MAP1A and alpha-tubulin within single cells. DMSO treated cells are overlaid in grey in all plots (data of a total of 498 cells combined from 3 individually cultured and sorted batches are shown).

To understand the relevance of MAP1A in the context of LSC and their susceptibility to PLK1i, we investigated PLK1 interaction with MAP1A in sorted OCI-AML22 cells by proximity ligation assay (PLA). The highest PLA signal was observed in CD112^low^ compared to CD112^high^ CD34^+^CD38^−^ cells and this signal was reduced upon volasertib treatment, indicating loss of PLK1-MAP1A interaction (Fig. 5 D, E; Supplementary Fig. 4G). Following volasertib treatment, MAP1A protein levels were increased in CD34^+^CD38^−^CD112^low^ OCI-AML22 at 48 hrs but not 24 hrs (Fig. 5F, G).

To investigate the dependency of the observed volasertib-induced effects in LSC on MAP1A, we performed shRNA-mediated knock-down of *MAP1A* in CD34^+^CD38^−^ OCI-AML22 cells. OCI-AML22 cells transduced with four independent shRNAs targeting MAP1A and co-expressing GFP did not reveal any competitive growth advantage or disadvantage in untreated cultures of bulk or CD34^+^CD38^−^ cells over 24 days compared to control shRNA (shRen)-transduced cells (Supplementary Fig. 4H, I). However, *MAP1A*-targeted shRNAs led to a reduction in CD112 surface expression as a marker of endosomal receptor internalization on GFP^+^CD34^+^CD38^−^ cells compared to controls (Fig. 5H). LC3B and LAMP1 levels in untreated shMAP1A-transduced CD34^+^CD38^−^ cells were elevated to similar levels as control cells treated with volasertib (Fig. 5I, J), indicative of increased vesicle endocytosis and/or reduced exocytosis. Blocking lysosomal acidification and degradation with bafilomycin A1 treatment levelled these differences (Supplementary Fig. 4J). Together, these observations indicate that shMAP1A impacts endolysosomal turnover and suggest that MAP1A regulates CD112 surface expression and recycling through its interaction with PLK1; disruption of this interaction either by volasertib or by knocking down MAP1A levels leads to internalization of CD112.

Volasertib treatment increased lysosomal activity in both control- and shMAP1A-transduced cells, but more so in the latter (Fig. 5K). Importantly, shMAP1A ameliorated apoptosis induction in the LSC-enriched subpopulation following volasertib treatment compared to control-transduced cells (Fig. 5L). The reduction in apoptotic rate by shMAP1A after volasertib treatment was mitigated after extended culture (Supplementary Fig. 4K), likely related to upregulation of *MAP1A* transcript levels during culture in GFP-sorted, shMAP1A-transduced cells (Supplementary Fig. 4L), pointing to tight cellular control of MAP1A levels.

Given the known role of MAP1A in microtubule stabilization, we next examined whether the accumulation of MAP1A following volasertib treatment alters microtubule dynamics by assessing alpha-tubulin K40 acetylation through confocal microscopy. In sorted CD34^+^CD38^−^CD112^low^ OCI-AML22 cells, tubulin acetylation levels increased at 48 hrs but not at 24 hrs after volasertib treatment (Fig. 5M, N; see also Fig. 5F, G). Confocal microscopy of volasertib-treated AnnexinV^−^ CD34^+^CD38^−^ OCI-AML22 cells sorted based on Cyto-ID and CD112 levels demonstrated an increase in both MAP1A and alpha-tubulin levels as Cyto-ID and CD112 surface levels decreased (Fig. 5O-Q). Together, these results suggest that the increase in MAP1A levels upon PLK1i delays clearance of the vesicle build-up via microtubule stabilization, ultimately leading to apoptosis.

Overall, the volasertib-sensitive interaction between PLK1 and MAP1A, occurring predominantly in quiescence-enriched LSC, together with the LSC-specific expression pattern of *MAP1A*, strongly implicate MAP1A and microtubule dynamics as key mechanisms underlying the selective effects of PLK1i in quiescence-enriched LSCs.

### Classic microtubule-targeting agents validate the importance of microtubule stabilization in eliminating LSC

In light of these findings, we reevaluated our initial drug screen data to look at drugs that interfere with microtubule dynamics. Accordingly, microtubule stabilizers (paclitaxel, docetaxel) and tubulin polymerization blockers (vinca alkaloids), drugs commonly used for their anti-proliferative effects, did show some LSC targeting potential. However, this was only observed at high concentration (100 - 2000 nM) while lower drug concentrations (2.5 – 40 nM) enriched the CD34^+^CD38^−^ LSC fraction, consistent with depletion of the proliferative cell fraction (Supplementary Fig. 5A; Suppl. Table 1).

Microtubule targeting agents have been shown to alter vesicle trafficking and are also used as compounds preventing vesicle fusion.(36)(71) Here, paclitaxel treatment of OCI-AML22 induced a 3dye^low^, CD112^−^ cell population and an AnnexinV^+^CD112^−^ population within only the LSC-enriched CD34^+^CD38^−^ and not the CD34^+^CD38^+^ fraction (Fig. 6A-D; Supplementary Fig. 5B); these findings faithfully phenocopied the effects of PLK1i. No additive effects were observed when cells were treated simultaneously with the 2 drugs, pointing to saturating activity on the same pathways. Paclitaxel treatment resulted in loss of PLA signal (Fig. 6E), consistent with disruption of the interaction between PLK1 and MAP1A and providing an explanation for the recapitulation of volasertib-induced effects including downregulation of CD112 surface expression and transient vesicle accumulation.

**Figure 6:**
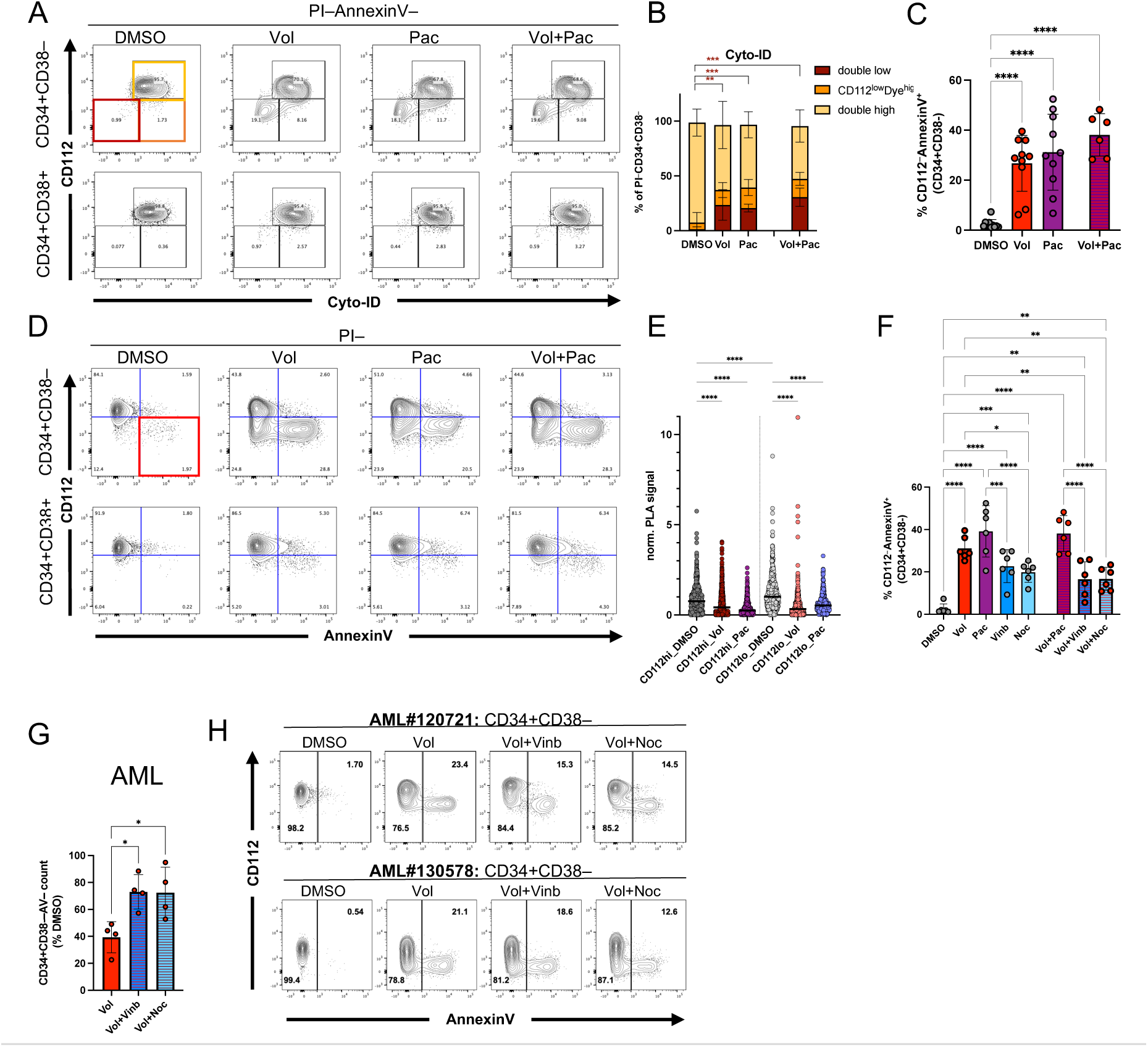
Classic microtubule-targeting agents validate the importance of microtubule stabilization in eliminating LSC. A-D) OCI-AML22 treated with indicated drugs for 48 hrs (50 nM volasertib, 500 nM paclitaxel). E) Proximity ligation assay of 24 hrs treated OCI-AML22 CD112^high^ and CD112^low^ cells ( 0 575 cells per condition from 3 individual cultures assessed in 2 assays). F) %CD112^-^ AnnexinV^+^ OCI-AML22 cells (CD34^+^CD38^-^) with single and double treatment (50 nM volasertib, 1 pM paclitaxel, 100 nM vinblastine, 500 ng/mL nocodazole). G) Absolute cell counts of primary AML PI^-^CD34^+^CD38^-^ cells after 48 hrs single or double treatment (n=4; 50 nM volasertib, 100 nM vinblastine, 500 ng/mL nocodazole). H) Flow cytometric analysis of 2 primary AML samples after 48 hrs single or double treatment as indicated and subgated for P^i-^Cd34^+^cD38^-^.

To assess whether volasertib-induced effects could be mitigated by microtubule destabilization, we treated OCI-AML22 cells and 4 primary AML patient samples with the microtubule assembly inhibitors vinblastine and nocodazole alone or in combination with volasertib. Consistent with their anti-proliferative activity, all drug combinations induced mitotic arrest (%G2/M) at different rates depending on the AML patient sample (Supplementary Fig. 5D). Although vinblastine and nocodazole induced a small AnnexinV^+^CD112^−^ population in CD34^+^CD38^−^ OCI-AML22, there was no substantial increase in apoptosis rate from 24 to 48 hrs after treatment, in contrast to what was seen with volasertib or paclitaxel treatment (Supplementary Fig. 5B-D). Importantly, both vinblastine and nocodazole alleviated the volasertib-induced effects on apoptosis rate and viability of CD34^+^CD38^−^ cells upon combination treatment in OCI-AML22 and AML patient samples (Fig. 6F-H; Supplementary Fig. 5E-G), regardless of whether they were added simultaneously or sequentially 4 hrs before or after volasertib (Fig. 6 F; Supplementary Fig. 5H). Together, these findings provide strong evidence for the critical role of microtubule stabilization in the selective killing of LSC by both volasertib and paclitaxel.

## Discussion

Here, we identify a previously unrecognized non-mitotic and cell-state-dependent function of PLK1 in quiescence-enriched LSC populations. Specifically, we show that perturbation of PLK1 function has fundamentally different consequences across cellular states: PLK1i induces mitotic catastrophe in cycling cells but triggers an apoptotic program in quiescent LSC through disruption of the PLK1–MAP1A interaction, leading to a complex cascade of changes in endolysosomal trafficking and turnover that ultimately compromises stem cell survival. The ability of microtubule-stabilizing and microtubule-destabilizing agents to respectively recapitulate and rescue these phenotypes establishes microtubule–endolysosomal dynamics as a key determinant of quiescent stem cell survival and challenges the prevailing view that quiescence confers resistance to conventional antimitotic therapies.

Accumulating evidence over recent years has indicated that the pharmacodynamics of PLK1 inhibitors and microtubule interfering agents cannot be explained solely by their canonical modes of action that target mitotic cells.(71–77) The reported divergence in molecular and cellular effects between ATP-competitive, allosteric and PLK1-degrading inhibitors, combined with the complex spatial and conformational regulation of this kinase, point to a PLK1 interactome that governs cellular processes that extend beyond mitosis and do not strictly rely on its kinase activity(18,78). Recently it was shown that ATP-competitive PLK1 inhibitors (including volasertib) induce conformational changes in PLK1 that relieve the kinase from its autoinhibited state while simultaneously blocking kinase-dependent functions.(27,28) This observation predicts that the interactome of volasertib-bound PLK1 would resemble that of a ca-T210D rather than a kd-K82R mutant under starvation conditions. In agreement with our findings, an open PLK1 conformation induced by volasertib, T210D mutation or by activation would allow interactions with the endolysosomal trafficking machinery and CD112 in LSC, but not with MAP1A. Of note, despite differences in structure and selectivity profiles, volasertib, onvansertib, and rigosertib all produced similar cell-state-dependent effects, confirming that the LSC-specific endolysosomal vulnerability identified here is PLK1-related and not specific to any one inhibitor.

Our findings provide additional context for past and future clinical development of PLK1 inhibitors, as conventional assessments of drug response have primarily focused on elimination of proliferating leukemic blasts and reduction of bulk disease burden. In view of the identification of cell state as a previously unrecognized determinant of PLK1 inhibitor response, PLK1-targeted therapies may warrant re-evaluation in biologically-defined patient subsets. Future studies should determine whether biomarkers such as stemness- or cell state-associated signatures, for example using clinically available RNA-based assays (79), can help identify patient populations most likely to benefit from PLK1-directed therapies.

MAP1A emerged as a candidate mediator of the cell-state-dependent effects observed following PLK1 perturbation. Disruption of PLK1-MAP1A interaction may occur through volasertib-induced conformational changes in PLK1, PLK1 activation or paclitaxel-mediated increase in microtubule stability. Of note, MAP1A expression in untreated LSC did not result in marked baseline microtubule acetylation/stabilization. This observation indicates that its well-known function as a stabilizer is restrained by its interaction with PLK1; disruption of this bond uncouples both proteins, which appear to reciprocally regulate their localization, interactome and function under these conditions. PLK1 has been shown to stall endocytic vesicle fusion during mitosis (80). Additionally, PLK1i has previously been shown to induce autophagy in the acute promyelocytic leukemia cell line NB4 as well as in other settings,(43,44,47) and we and others have shown that autophagy and endolysosomal activity regulate stemness in both normal and leukemic haematopoiesis,(40,41,81–84), lending further support for a critical role of PLK1 in regulating these processes in LSC.

After volasertib or paclitaxel treatment, an initial increase in endolysosomal trafficking and intracellular vesicle accumulation is followed by microtubule stabilization as delineated by the sequence of events captured by flow cytometry (CD112 vs vesicle 3dye staining in AnnexinV-cells). Microtubule stabilization is essential for re-initiation of stalled fusion and vesicle clearance, which cannot occur in the presence of the fusion inhibitor chloroquine or microtubule destabilizing agents. The fact that chloroquine does not prevent the accompanying loss of CD112 surface expression indicates that endosomal receptor re-routing cannot be restored when vesicle fusion is inhibited, evidence of already irreversible effects on cellular integrity. In the absence of chloroquine, the subsequent delayed, but excess vesicle fusion marked by a complete loss of acidic organelles, cannot re-establish the status quo of the LSC state, leading to initiation of apoptosis. This cascade reflects the convergence of multiple cellular mechanisms commonly targeted by anti-cancer therapies and highlights the close coupling between microtubule dynamics, vesicle trafficking, and endolysosomal homeostasis in the maintenance of quiescent stem cell integrity. Although our studies were performed in AML, cytoskeletal regulation of vesicle trafficking and endolysosomal homeostasis is unlikely to be restricted to this context, suggesting that similar mechanisms may operate in other stem-like or quiescent cell populations. The PLK1–MAP1A interaction therefore represents a previously unrecognized vulnerability of quiescent stem cells that may be therapeutically exploited by compounds that are already in the clinic.

Our findings additionally highlight both the challenges and opportunities associated with combinatorial drug treatments, as certain compounds may potentiate, saturate or interfere with one another’s mechanisms of action and efficacy. For example, induction of endocytosis by severing the PLK1-MAP1A bond may enhance uptake of surface receptor targeted therapies and may underlie the reversal of resistance to an EGFR tyrosine kinase inhibitor previously reported for volasertib.(85) Numerous combination therapies that include paclitaxel have already been tested preclinically in combination with PLK1 inhibitors, aiming for synergism or overcoming of drug resistance across a range of cancer models.(21) Our findings could inform the design of monotherapies capable of targeting both the tumor bulk and quiescent CSC, and raise the possibility that some of the historic success achieved with conventional antimitotic chemotherapy may also involve previously unrecognized effects on intracellular trafficking pathways in non-proliferating cells.

Discovery of the cell-state dependent effects of PLK1i required resolution of LSC populations with distinct quiescence properties based on differential surface expression of the adhesion molecule and immune checkpoint ligand CD112, a PLK1 interactor that also enabled visualization of endolysosomal receptor internalization. Small changes in CD112 surface abundance allow prospective enrichment for deeply dormant versus quiescent but activation-primed long-term HSC and LSC.(53,58) The governance of CD112 surface expression by interaction with proteins that appear central to HSC and LSC function indicates that its biological role may be important in both contexts. As a ligand for the immune checkpoint receptors TIGIT and CD226, CD112 likely modulates immune cell interactions with HSC and LSC, transducing either activating or inhibitory signals depending on the context.(86) Future studies are needed to assess how the excessive downregulation of CD112 on LSC following disruption of PLK1-MAP1A may impact immune responses within the cancer microenvironment in immunocompetent settings.

Our data introduce a quiescent stem cell-specific gatekeeper function of the interaction between PLK1 and MAP1A that appears to contribute to cell surface integrity by coordinating endolysosomal receptor recycling, a process that is closely coupled to the microtubule network. MAP1A has been predominantly studied in neurogenesis and brain tissue due to its high expression in these contexts.(87) It has also been implicated in retrograde intracellular transport of the HIV nucleocapsid in human macrophages by tethering the viral complexes to microtubules, and in contributing to cell rigidity in ATRA-resistant acute promyelocytic leukemia.(39,88) In glioblastoma, MAP1A facilitates EGFR trafficking and signaling, processes that are also sensitive to paclitaxel and volasertib treatment in non-small cell lung cancer (NSCLC), where *MAP1A* expression is being explored as a potential prognostic marker.(37,38,89–91) Overall, these observations underscore the tight interconnection of these molecules and their functions, and suggest that this axis and the vulnerabilities uncovered here may extend to other tissues and tumor hierarchies.

## Methods

### Human specimen collection and processing

All biological samples were collected with written informed consent according to procedures approved by the Research Ethics Board of the University Health Network (UHN; REB# 01-0573-C), and AML patient samples were viably frozen in the Leukemia Tissue Bank at Princess Margaret Cancer Centre (Toronto, Ontario, Canada). The UHN Research Ethics Board operates in compliance with the Tri-Council Policy Statement; International Council for Harmonization Guideline for Good Clinical Practice E6(R1); Ontario Personal Health Information Protection Act (2004); Part C Division 5 of the Food and Drug Regulations; and Part 4 of the Natural Health Products Regulations and the Medical Devices Regulations of Health Canada. The inclusion criteria for AML samples were based on their prior established ability to engraft NSG mice (RRID:IMSR_JAX:005557) and their availability. Samples were thawed by dropwise addition of X-VIVO + 50% FBS supplemented with DNase (100 μg/mL final concentration, Roche, Cat#11284932001). For transplantation, AML cells were CD3-depleted by LS column purification with MACS magnet technology (Miltenyi). For short-term culture, cells were either CD34^+^ enriched (MS columns, Miltenyi) or after overnight culture AnnexinV^+^ depleted (Easy Sep, Stem cell technologies) depending on the known CD34/CD38 profile and LSC content within thereby defined fractions, and their overall viability.

### AML cultures and treatment

OCI-AML22 and OCI-AML8227 cells were derived from the long-term expansion of individual primary AML patient samples (48,49,54) and tested for *Mycoplasma* contamination using the MycoAlert Mycoplasma Detection Kit (Lonza, Cat # LT07-318) after conclusion of experiments.

OCI-AML8227 were cultured in X-VIVO 10 medium (Lonza, Cat # 04-380Q) supplemented with 20% (v/v) BIT 9500 serum substitute (Stem Cell Technologies, Cat # 09500), 2 mM L-Glutamine (GIBCO, Cat #25030081), and the following cytokines: 10 ng/mL IL-6 (Peprotech, Cat # 200-06), 5 ng/mL IL-3 (Peprotech, Cat # 200-03),100 ng/mL SCF (Peprotech, Cat # 300-07), 50 ng/mL FLT3L (Peprotech, Cat # 300-19), 5 ng/mL G-CSF (Peprotech, Cat # 300-23) and 25 ng/mL TPO (Peprotech, Cat # 300-18). Antimicrobial agents were not added to the culture medium. Cells were maintained at a density of 0.4 × 10^6^ cells/mL and passaged every 3 to 5 days in 24 or 96-well, flat-bottom plates.

OCI-AML22 and primary patient AML cells were cultured in X-VIVO 10 (Lonza, BE04-380Q) supplemented with 20% BIT 9500 Serum Substitute (STEMCELL Technologies, 09500), 1× Glutamax Supplement (Thermo Fisher Scientific, 35050061), Primocin 0.1 mg/mL (Thermo Fisher Scientific), SCF (200 ng/mL; Miltenyi Biotec, 130-096-696), IL3 (20 ng/mL; Miltenyi Biotec, 130-095-069), TPO (20 ng/mL; PeproTech, 300-18), FLT3L (40 ng/mL; PeproTech, 300-19 B), IL6 (10 ng/mL; Miltenyi Biotec, 130-093-934), and G-CSF (10 ng/mL; Miltenyi Biotec, 130-093-861). OCI-AML22 cells were maintained at a density of 0.8 × 10^6^ cells/mL and passaged every 3 to 5 days in 96-well, flat-bottom plates. To serially expand the cells, the CD34^+^ fraction was regularly sorted, or dead cell depletion was performed using the EasySep Dead Cell Removal (Annexin V) Kit (Cat# NC1408982; Stem Cell Technologies) according to the manufacturer’s protocol. All cell cultures were maintained in a humidified incubator at 37°C in 5% CO_2_. Drug treatments of primary AML samples were initiated after overnight culture and of OCI-AML8227 or OCI-AML22 freshly or +1 day passaged bulk cells or directly after sorting (aiming for passaging cell concentrations) adding to pre-plated cell suspension same volume of media containing 2x of final single or double drug concentrations. Final drug concentration if not otherwise indicated in figure or figure legends were 50 nM volasertib (Selleck Chemicals Cat #S2235), 50 nM onvansertib (Selleck Chemicals Cat # S7255), 50 nM rigosertib (Selleck Chemicals # S1362), 400 nM GW-843682X (“GW”, Selleck Chemicals #S2880), 0.5 or 1uM paclitaxel (Sigma, Cat #T7191), 500 ng/mL nocodazole (Sigma, Cat #487929), 100 nM vinblastine (Sigma, Cat #V1377), 60 nM chloroquine (Enzo, ENZ-KIT175), 500 nM rapamycin (Enzo, ENZ-KIT175), 20 nM bafilomycin A1(Cedarlane/Cayman #11038). For inhibiting lysosomal degradation bafilomycin A1 (final concentration 20 nM) was added 2 hrs before cells were harvested for immunostainings or stained for flow cytometry.

### Primary and secondary LSC drug screen

The 1219 compounds used in the primary screen were dissolved in dimethyl sulfoxide (DMSO) and deposited into 96-well microplates. The master plates were sealed with aluminum foil and stored in −80°C until use. The compounds from the master plates were transferred to wells in 384-well cell culture plates (Sigma, Cat # CLS3571) using an acoustic liquid handler (Echo, Labcyte). The volume of compound transferred in each well was pre-determined to achieve the desired drug concentration in the final total volume. Eight serial concentrations were tested for each compound in the primary screen. Each well in the 384-well plates were back-filled with DMSO to maintain the same final DMSO concentration in each well. Wells containing DMSO only were included in each plate as controls.

OCI-AML8227 cells (20,000) in culture medium were then added to each well in a volume of 80 μl using a robotic dispenser (Multidrop Combi Reagent Dispenser, Thermo Fisher Scientific). The cells were incubated at 37°C in 5% CO2 for 3 days. After the 3-day incubation, the plates were centrifuged at 500xg for 5 minutes to collect cells at the bottom of the plates and the supernatant removed. The cells were resuspended in 10 μL per well of FACS buffer (2% [v/v] FBS in HBSS with calcium) containing 10% (v/v) FcR blocking reagent (Miltenyi Biotech, Cat # 130-059-901; RRID:AB_2892112), anti-CD34 APC at 1:200 dilution (BD Biosciences, Cat # 340441; RRID:AB_2228982), anti-CD38 PE at 1:50 dilution (BD Biosciences, Cat # 12-0388-42; RRID:AB_1518748), Annexin V FITC at 1:20 dilution (BD Biosciences, Cat # 556419; RRID: AB_2665412), and 7-AAD at 1:20 dilution (BD Biosciences, Cat # 559925; RRID:AB_2869266). The cells were stained at room temperature for 15 minutes and washed once in FACS buffer. The cells were then fixed in 20 μL per well of fixation buffer (Biolegend, Cat # 420801) for 20 minutes at room temperature and washed once in FACS buffer. The fixed cells were resuspended in 50 μL of freezing medium (10% [v/v] DMSO in FBS) per well. The cell culture plates were then sealed with aluminum foil and stored in −80°C for later analysis. At the time of analysis, the cells were thawed at 37°C for 30 minutes, centrifuged, and resuspended in 60 μL of FACS buffer per well. The cells were analyzed using a BD FACS Canto II flow cytometer with a High Throughout Sampler attachment.

The proportion of CD34^+^CD38^−^ cells in the viable population (Annexin V^-^ and 7-AAD^-^) was determined for each drug at the different concentrations using FlowJo Software (RRID:SCR_008520, version 10). To estimate drug effect, a *t*-statistic was calculated for each drug as described previously.(16) Next, 222 compounds were selected for confirmation in a secondary screen. The design of the secondary screen was identical to the primary screen except for a greater number of DMSO control wells per plate and an increase in the number of concentrations tested per drug to 12 (Supplementary Table 1).

### Flow cytometric analysis and sorting

Cells were incubated (15-20 min, RT) in their medium with the following antibodies: PE-Cy7-anti-CD38 (BD #335790, RRID:AB_399969, 1/100 dilution) and APC-Cy7-anti-CD34 (Biolegend #343514, RRID:AB_1877168, 1/200 dilution), APC-anti-CD112 (Biolegend # 337412, RRID:AB_2565729, 1/100 dilution) and PE-Annexin V (BD # 556422, RRID:AB_2869071, 1/25 dilution, BD) for sorting and/or flow cytometric analysis. After washing, cells were resuspended at 0.5–10 × 10^6^ cells per ml in AnnexinV-buffer with 0.1 μg l^−1^ propidium iodide (PI). Cell preparation for intracellular DAPI (1 mg/mL; Thermo Fisher) staining or detailed cell cycle analysis was performed with BD Cytofix/Cytoperm (BD, RRID:AB_2869008) after surface antibody staining as above and as described before(58) and if applicable incubated with PE-anti-Ki67 (BD #556027, RRID:AB_2266296, 1/30 dilution) in PermWash solution (BD) overnight. Other flow cytometry-based assays were performed according to the manufacturers’ protocols: CellROX (Thermo Fisher #C10444, 1/500), LysoTracker Green or Blue (Thermo Fisher #L7526//#L7525, 1/13,000), LysoSensor Green or Blue (Thermo Fisher #L7535/#L7533, 1/500), Cyto-ID (Enzo #ENZ-51031-200, 1/500) and CellEvent Caspase-3/7 Green Flow Cytometry Assay Kit (Thermo Fisher #C10427, 1/1000) staining in X-vivo 10 (Lonza), 20% BIT 9500 (Stem Cell Technologies) for 30 min at 37 °C while co-staining with antibody panel as above. After washing, cells were resuspended in AnnexinV-buffer with 0.1 μg l^−1^ propidium iodide (PI). Cells were sorted on BD FACSAria III and BD Symphony S6 Instruments or analyzed on BD Celesta or BD Symphony A1 instruments using a high-throughput sampler allowing for flow cytometry-based cell counting. Positive and negative gates were set according to unstained or isotype controls or according to set percentages (CD112^high^ and CD112^low^ gates as the 25% maximum or minimum, respectively, of the CD112 spectrum within CD34^+^CD38^−^ parental gate). Data were analyzed by FlowJo (version 10.10, RRID:SCR_008520).

### Patient-derived xenografts and *in vivo* treatment

Animal experiments were done in accordance with institutional guidelines approved by the UHN Animal care, and we complied with all relevant ethical regulations for animal testing and research. NOD.Cg-*Prkdc^scid^Il2rg^tm1Wjl^*/SzJ (NSG; RRID:IMSR_JAX:005557) mice were housed in a controlled environment with a 12-h:12-h light:dark cycle including a 30-min transition, a room temperature (RT) of 21–23 °C and a humidity of 30–60%, and had ad libitum access to dry laboratory food and water at the animal facility (ARC) at Princess Margaret Cancer Centre. They were kept in a room designated only for immunocompromised mice with individually ventilated racks equipped with complete sterile micro-isolator caging (IVC), on corn-cob bedding and supplied with environmental enrichment in the form of a red house/tube and a cotton nestlet. Cages were changed every <7 d under a biological safety cabinet. Health status was monitored using a combination of soiled bedding sentinels and environmental monitoring.

Twelve-to 16-week-old male and female NSG mice (RRID:IMSR_JAX:005557) were sublethally irradiated (225 cGy) 24 hours before intrafemoral injection of OCI-AML8227, primary AML cells or hCD45^+^CD33^+^ sorted (BD Fusion or Beckman Coulter MoFlo Astrios) cells from primary PDX (hCD45+ from CB derived xenografts) and randomly assigned to treatment groups. Mice were euthanized 6 to 8 weeks (primary PDX) or 12 weeks (OCI-AML8227, secondary PDX) after transplant and AML engraftment (hCD45^+^CD33^+^) in the injected right femur, the non-injected left femur (+tibiae/”BM”) was assessed by flow cytometry (BD Celesta and BD Symphony A1) after long bones were flushed separately in Iscove’s modified Dulbecco’s medium (IMDM). The remaining, unstained cell suspensions (90%) were frozen viably at −150 °C for secondary transplantation. Primary PDX were treated after xenografts were established (2-4 weeks). For this, volasertib (BI-6727; MedChem Express #HY-12137) was reconstituted in prewarmed DMSO to 50 mg/mL, diluted to a final concentration of 1.25 mg/mL with water and administered by oral gavage at the indicated dosing schedule (Fig. 2A). Control mice were given an equivalent volume of vehicle by oral gavage.

The flow cytometry panels were for OCI-AML8227 LDA hCD45-V500 (clone HL30; BD #560777; RRID:AB_1937324; 1/100), hCD45-APC (clone 2D1;BD #340943; RRID:AB_400555; 1/100); CD34-APC-Cy7 (Biolegend #343514, RRID:AB_1877168, 1/200), CD38-PE (BD 347687, RRID:AB_1518748, 1/100), Propidium Iodide/PI (Thermo Fisher # P3566, 1/10,000); for PDX CD15-FITC (BD # 347423, *RRID*:AB_400297, 1/100), CD117-PE (BD # 340529, RRID:AB_400044, 1/100), hCD45-V500 (clone HL30; BD #560777; RRID:AB_1937324, 1/100), CD33-BV786 (BD # 740974, RRID:AB_2740599, 1/200), CD14-PC5 (Beckman Coulter #IM2640U, 1/100), CD34-APC-Cy7 (Biolegend #343514, RRID:AB_1877168, 1/200), CD38-APC (BD 340439, RRID:AB_400512, 1/100), SytoxBlue (Thermo Fisher # S34857, 1/2,000); for sorts for secondary transplantation: hCD45-FITC (BD #347463, RRID:AB_400306, 1/100), CD33-APC (BD # 340474, RRID:AB_400518, 1/100), PI (Thermo Fisher # P3566, 1/10,000). For Limiting dilution analysis (LDA) a cut off of 0.1% hCD45^+^ engraftment in RF (right/injected femur) and LF (left/non-injected femur) was applied. Sample acquisition was performed with a BD Celesta Instrument and BD FACSDiva Software (RRID:SCR_001456, Version 8.0.1.1), analysis with FlowJo_v10.10 software (RRID:SCR_008520) and LSC frequency and statistical significance were calculated by ELDA software (RRID:SCR_018933, https://bioinf.wehi.edu.au/software/elda/).(92)

### Immunofluorescence assays

Cells were spun onto Poly-L-Lysine (Sigma # P8920)-coated glass slides (Ibidi # 81817, 200 xg, 10 min), fixed with 4% paraformaldehyde (VWR, 15714-S) and permeabilized with 0.5% Triton (Sigma) before blocking (PBS, 10% FBS, 5% BSA, 2% donkey serum/Sigma #D9663). Slides were incubated with primary antibodies in blocking solution O/N at 4°C as follows: a-tubulin (1:500; abcam #ab6160, RRID:AB_305328), LAMP1-AF488 (1:50 – 1:100; R&D Systems # IC7985G, RRID:AB_2928967); CD112/Nectin2 (1:100; Novus Biological #AF2229, RRID:AB_2269089), LC3B (1:100; Nanotools #5F10, RRID:AB_2722733), LC3B-AF647 (1:50; Novus Biologicals #NBP2-59800AF647, variant of RRID: AB_3094850), PLK1 (1:50; Santa Cruz sc-17783, RRID:AB_628157), MAP1A (1:500; abcam #ab184350, RRID:AB_2732015), Acetyl-a-tubulin (1:500; abcam #ab179484). Secondary antibodies (all Thermo Fisher Scientific: donkey anti-mouse AF555, goat anti-rabbit AF647, goat anti-mouse 568, donkey anti-rat AF555, goat anti-mouse AF647, donkey anti-mouse AF488, donkey anti-goat AF647, donkey anti-goat AF488, donkey anti-rat AF488) were added (PBS, 0.025% Tween, Sigma, 1.5 h, RT) at 1:400. After washing, nuclei were stained with 1 mg/mL DAPI (Thermo Fisher Scientific) and slides were mounted (Fluoromount G, Thermo Fisher Scientific). An average of 50 single cell images per specimen and condition were captured by a Zeiss LSM700 Confocal (oil, 63x/1.4NA, Zen 2012) or a Leica Stellaris 5 (HC PL APO 63x/1.40 NA Oil immersion CS2/for PLA: HC PL APO 20x/0.75 NA CS2, Leica LAS-X Software, RRID:SCR_013673). In an estimated one third of repetitions experimenter were blinded for specimen, treatment and parameters. Downstream analysis was performed with ImageJ/Fiji (RRID:SCR_002285) and FlowJo10 (RRID:SCR_008520). Mitotic catastrophes were omitted or excluded from further analysis in case of unbiased acquisition of mitotic catastrophes and intact nuclei. In case of high inter-experimental variability between independent stainings, data were normalized relative to the median of controls.

For proximity ligation assay/PLA slides were prepared for Sigma Duolink In Situ PLA assay (Cat #DUO92007; Cat #DUO82047; Cat #DUO92004; Cat #DUO92002) as follows: Slides were after permeabilization blocked with in-house blocking solution (PBS, 10% FBS, 5% BSA, 2% donkey serum/Sigma) for 5 hrs. Primary antibodies including controls (see below) were incubated in in-house blocking solution O/N at 4°C, then 3x rinsed and 2x incubated (15 min, RT) with Sigma Wash Buffer A, blocked with Sigma blocking solution (3 hr, 37 °C) and washed with Sigma Wash A for 5 min, RT before probe solution with antibody diluent and plus/minus probes were added (1 hr) and protocol was followed according to manufactures instructions with 100 min of polymerase reaction. After final wash in 0.01x Wash buffer B nuclei were stained with 1 mg/mL DAPI (Thermo Fisher Scientific #D21490) and slides were mounted (Fluoromount G, Thermo Fisher Scientific #00-4958-02). Background signal was determined according to several controls: Bulk or viable sorted CD34^+^CD38^−^ OCI-AML22 cells incubated with no primary antibody (AB), no primary nor secondary AB (DAPI only), anti-PLK1 (RRID:AB_628157) and anti-MAP1A (RRID:AB_2732015) with no secondary AB, anti-PLK1 in combination with rabbit IgG (anti-Flag; 1/800 dilution, RRID:AB_2572291), anti-MAP1A in combination with mouse IgG (anti-Flag; 1/250, RRID:AB_262044), rabbit IgG in combination with mouse IgG. Images were captured by a Leica Stellaris 5 (HC PL APO 20x/0.75 NA CS2, Leica LAS-X Software/RRID:SCR_013673).

### BioID (miniTurbo) assay

Flp-In T-Rex 293 cells (Thermo Fisher, R78007, RRID:CVCL_U427) were cultured in Dulbecco’s Modified Eagle Medium (DMEM, Wisent #319-016-CL) containing 100 μg/ml hygromycin B and supplemented with 10% fetal bovine serum (FBS). Tetracycline-inducible, miniTurbo-tagged PLK1 variants were stably expressed in Flp-In T-Rex 293 cells (RRID:CVCL_U427) using the Flp-In site-specific recombination system. Cells were then co-transfected with pOG44 (Flp-recombinase expression vector, RRID:Addgene_209087) and the respective PLK1 variant miniTurbo fusions encoding pcDNA5-FRT-TO plasmid (see below) and stable cell lines were generated by selection with 200 μg/ml hygromycin B. The expression of fusion proteins in the resulting isogenic cell pools was validated by intracellular flow cytometry detecting PLK1(Santa Cruz, sc-17783, RRID:AB_628157). Two independent pools (biological replicates) were created for each condition. The expression of the fusion protein was induced by the addition of 1 μg/ml tetracycline to the culture media for 24 hrs while cells were simultaneously starved by FBS withdrawal. Cells were then treated with iso-or hyper-osmotic solutions with 50 μM biotin for 1 hr and were subsequently collected by scraping and centrifugation. Cell pellets were washed twice with PBS and stored at −80 °C until lysis.

Biotin-streptavidin affinity purification and mass spectrometric analysis and data processing for the BioID assay were performed as described previously. (93,94). Briefly, high-confidence interactors were identified that fulfilled the following criteria: iProphet > 0.9, at least two unique peptides and a FDR < 1% comparing the two highest peptide counts. Twenty control runs (from cells treated with isosmotic condition) were condensed to the four highest spectral counts for each prey and compared to the experimental data, consisting of two biological replicates (each analyzed with two technical replicates). High-confidence interactors were defined as those with a Bayesian false discovery rate (BFDR) ≤0.01. Data were loaded into Cytoscape (RRID:SCR_003032; v3.9.1 (95)) for network visualization and in Cytoscape STRING enrichment (Gene Ontology biological processes from http://baderlab.org/GeneSets/; version February 2020) was used for pathway analysis to subsequently generate the enrichment map (EnrichmentMap, RRID:SCR_016052, version 3.1.0 in Cytoscape; RRID:SCR_016052) to visualize enriched pathway gene sets with cut-off values for P-value= 0.05 and FDR Q-value = 0.05. For GSEA analysis (GSEA software, RRID:SCR_003199, v4.0.3;) the list of high-confidence interactors for PLK1wt (PLK1wt BioID interactome) was used as gene set and applied to rank files from the comparisons as indicated using 2,000 permutations and default parameters.

### Quantitative gene expression assay

Total RNA extraction was performed with RNeasy Plus Micro Kit (QIAGEN) according to the manufacturer’s instructions. SuperScript VILO IV cDNA Synthesis Kit (Thermo Fisher) with EzDNAse treatment was used for cDNA synthesis. Digital droplet PCR (ddPCR) was performed with 5-50ng cDNA using the QX200 Droplet Digital PCR System (Bio-Rad) following manufacturer’s instructions. Commercial primers and probes were purchased from Thermo Fisher (HPRT1 Hs02800695_m1, MAP1A Hs00357973_m1). MAP1A gene expression for each sample was determined over HPRT1 normalizer using the following formula: number of MAP1A+ droplets/number of HPRT1+ droplets.

### Cloning shRNA lentiviral vectors and miniTurbo PLK1 fusion plasmids

PLK1 wt and mutants were PCR amplified from pCer-C3 plasmids (RRID:Addgene_68132, RRID:Addgene_68133, RRID:Addgene_68134) and inserted into the N- and C-terminal multiple-cloning-sites (MCS) of miniTurbo expressing plasmids (pcDNA5_FRT-TO_3xFLAG-MiniTurboID-(MCS) and pcDNA5_FRT-TO_(MCS)-MiniTurboID-3xFLAG) using NotI and XbaI restriction enzymes (NEB). PLK1 overexpression was confirmed in HEK293T (RRID:CVCL_0063) by intracellular flow cytometry detecting PLK1(Santa Cruz, sc-17783, RRID:AB_628157).

Oligonucleotides (5’-3’):

MCS_PLK1_F_NotI: AAAAGCGGCCGCCATGAGTGCTGCAGTGACTGCAGGGAAG

MCS_PLK1_R_XbaI: AAAACTCNGAGTTAGGAGGCCTTGAGACGGTTGCTG

mut_PLK1_K82R_F: TTCGCGGGCATGATTGTGCCT

mut_PLK1_K82R_R: CACCTCCTTGGTGTCCGCGTCC

mut_PLK1_T210D_F: AGAGGAAGAAGGCCCTGTGTGGGAC

mut_PLK1_T210D_R: CCCCGTCATATTCGACTTTGGTTG.

ShRNAs sequences were predicted based on the Sherwood algorithm described in Knott et al. 2014(96). shRNAs oligos were designed to include BsmBI-v2 flanking recognition sites to generate 4bp overhangs. The inserts were generated by PCR using Q5 high fidelity DNA polymerase (NEB#M0491S). The PCR products were purified prior to Golden gate assembly. The amplified PCR products was subcloned into the pLV[miR30]-SFFV>EGFP:5’ miR-30a:BsmBI lentiviral vector using T4 DNA ligation buffer(NEB#M0202), and BsmBI-v2 enzyme mix (NEB#E1602S).

ShRNA knock-down sequences:

shMAP1A-1: TGCTGTTGACAGTGAGCGCTGCAGGAACCCTTGAAGGTAATAGTGAAGCCACAGATGT ATTACCTTCAAGGGTTCCTGCATTGCCTACTGCCTCGGA

shMAP1A-2: TGCTGTTGACAGTGAGCGCAAGCAATAGTCTTTGAGATTATAGTGAAGCCACAGATGTA TAATCTCAAAGACTATTGCTTTTGCCTACTGCCTCGGA

shMAP1A-3: TGCTGTTGACAGTGAGCGAAGACCTAGAACAGACAGACAATAGTGAAGCCACAGATGT ATTGTCTGTCTGTTCTAGGTCTCTGCCTACTGCCTCGGA

shMAP1A-4: TGCTGTTGACAGTGAGCGACCCAGTGGAAGAAAAGTCTGATAGTGAAGCCACAGATGT ATCAGACTTTTCTTCCACTGGGGTGCCTACTGCCTCGGA

BsmBI-v2 Oligonucleotides:

MAP1A-1 Fw: GGCTACCGTCTCGAGCGTGCTGTTGACAGTGAGCG

MAP1A-1 Rev: GGCTACCGTCTCCGGCATCCGAGGCAGTAGGCAAT

MAP1A-2 Fw: GGCTACCGTCTCAAGCGTGCTGTTGACAGTGAGCG

MAP1A-2 Rev: GGCTACCGTCTCGGGCATCCGAGGCAGTAGGCAAAAG

MAP1A-3 Fw: GGCCTACCGTCTCCAGCGTGCTGTTGACAGTGAGCG

MAP1A-3 Rev: GGCTACCGTCTCTGGCATCCGAGGCAGTAGGCAGAG

MAP1A-4 Fw: GGCTACCGTCTCTAGCGTGCTGTTGACAGTGAGCG

MAP1A-4 Rev: GGCTACCGTCTCAGGCATCCGAGGCAGTAGGCACC

### Lentiviral production and transduction

VSV-G pseudotyped lentiviral vector particles were produced by polyethyleneimine (PEI)-based co-transfection of 10.5 μg of pMD2.G, 20.5 μg of pCMVR8.74 (both from Addgene, RRID:Addgene_12259 and RRID:Addgene_22036) and 38 μg of transfer vector into HEK293T (RRID:CVCL_0063) cells. Viral particles harvested after 44 and 70 hrs were sterile filtered (0.45-µm filter), concentrated 100x by LentiX Concentrator (Takara), resuspended in X-VIVO 10 (Lonza) and stored at –80 °C until use. Lentiviral transductions of OCI-AML22 were carried out at a cell density of 0.5 × 10^6^ to 0.8 × 10^6^ cells/mL in 96-well, round-bottom plates after 2 days of recovery from FACS sorting for CD34^+^ cells by adding <1/5 of the volume of 100× concentrated viral supernatant. After 24 hrs, a half-medium exchange was performed, and cells were subsequently passaged every 3 to 5 days maintaining a density of 0.8 × 10^6^ cells/mL. GFP expression, CD34 (APCCy7), CD38 (PECy7), and CD112(APC) surface expression was monitored every 5-7 days over 24 days of culture by flow cytometry using a BD Symphony A1.

### Quantification and statistical analysis

GraphPad Prism 10 (RRID:SCR_002798) was used for all statistical analyses except for GSEA (RRID:SCR_003199) and ELDA analysis (RRID:SCR_018933), which was analyzed with the respective software. Unless otherwise indicated in the figure legends, mean ± s.d. values are reported in bar graphs, median values in scatter dot and violin plots, and minimal to maximal values are represented in box-and-whisker plots. Unless otherwise noted, statistical significance (*P<0.05, **P<0.01 and ***P<0.001) was determined using two-tailed Student’s t-test paired and unpaired and with or without Welch’s correction as applicable using GraphPad Prism 10 (RRID:SCR_002798). If normal-distribution assumptions were not valid, statistical significance was evaluated using the Mann-Whitney U test (two-tailed) for single comparisons or the Kruskal-Wallis test for multiple comparisons using GraphPad Prism 10 (RRID:SCR_002798). A normal distribution of the data was tested using the Kolmogorov-Smirnov test if the sample size allowed. One-way ANOVA was used to compare means among three or more independent groups. Applicable post hoc tests were used to compare all pairs of treatment groups when the overall P value was <0.05.

## Data and Material Availability

Previously published data sets used in this study are under accession numbers GSE125345 (RNA-seq, human CB hierarchy), GSE76008 (RNA array, LSC+ vs LSC-), GSE199452 (RNA-seq, LSC+ vs LSC-), GSE199451 (RNA-seq, diagnosis-relapse AML pairs), GSE211595 (RNA-seq, OCI-AML22 fractions). *MAP1A* expression across the bone marrow reference map was generated with https://cellxgene.cziscience.com/e/cd2f23c1-aef1-48ae-8eb4-0bcf124e567d.cxg/. Other data and materials relevant to this study are available from the corresponding authors upon reasonable request.

## Acknowledgments

We acknowledge support from the Princess Margaret Leukemia Tissue Bank and the patients who donated samples used in this study. This project was supported by the Leukemia and Lymphoma Society of Canada. We also wish to thank the SPARC BioCentre at the Hospital for Sick Children and OICR Drug Discovery Program for their support with drug screening and the SickKids-UHN Flow Cytometry Facility and the Advanced Optical Microscopy Facility (AOMF) for their services. AI (MS Copilot) has only been used for proofreading and to improve clarity of human author written text.

## Author contributions

K.B.K and J.C.Y.W. wrote the manuscript; Q.L., K.B.K. and J.C.Y.W. designed the study and analyzed results. Q.L., M.G., A.V., A.S., F.Y., B.D., N.M., L.J., C.L., P.J.J, E.R.L. and K.B.K. performed the experiments and analyzed data. C.X. and G.D.B. performed the bioinformatic analysis of the drug screen. A.M., H.B. and E.R.L. analyzed data and provided conceptual input. A.M., A.A. and M.M provided AML samples and assisted with sample selection. S.M.C. and B.R. supervised research. J.C.Y.W. and K.B.K. provided funding and study supervision. All authors reviewed the manuscript.

## Supplementary Tables

**Supplementary Table 1:** LSC drug screens: drugs, drug targets, t-scores per concentration and LSC depletion scores.

**Supplementary Table 2:** Experimental and clinical parameters of AML patient samples used for *in vivo* and *in vitro* studies.

**Supplementary Table 3:** Secondary limiting dilution assays: cell doses, engraftment scoring and calculations.

**Supplementary Table 4:** BioID interactome results.

