## Supplementary Figures for "Cell-state–dependent responses to PLK1 inhibition reveal a non-canonical microtubule-endolysosomal vulnerability in quiescent leukemia stem cells"

### Supplementary Figure 1

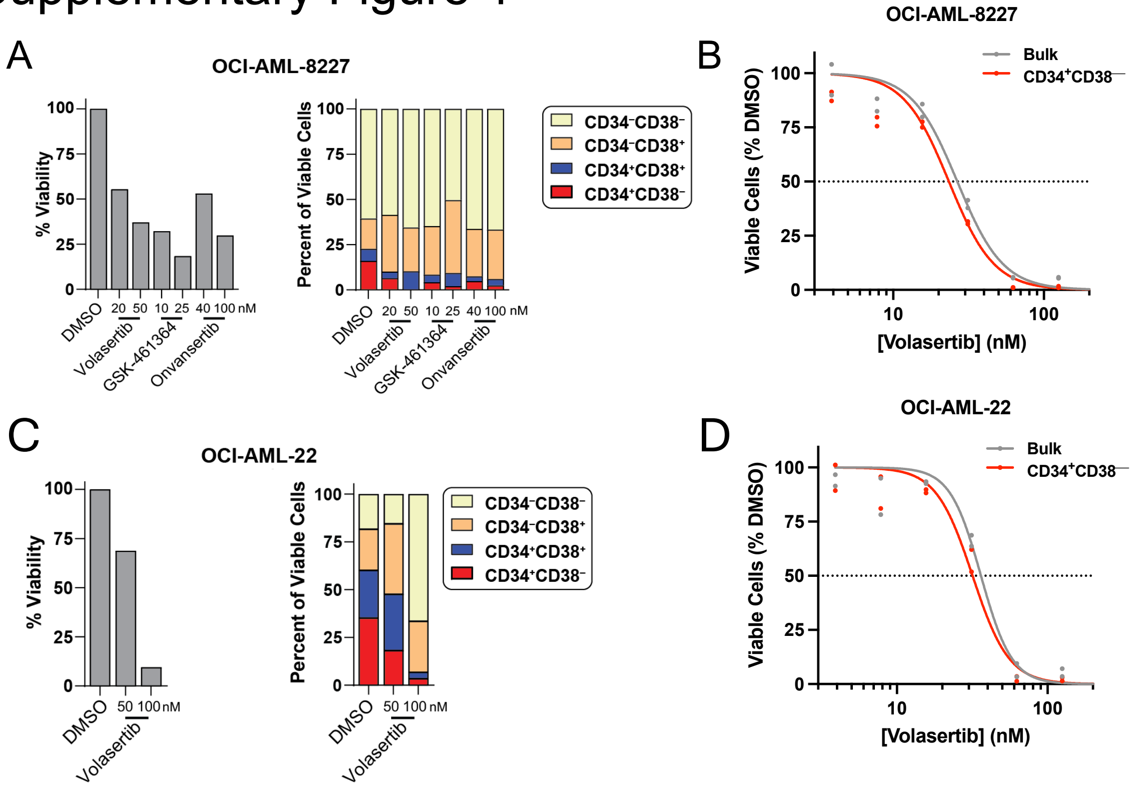

**Suppl. Fig.1**

A) OCI-AML8227 cells were treated with the indicated PLK1 inhibitors at indicated doses for 72 h and viability relative to DMSO (%7AAD$^{-}$AnnexinV$^{-}$) and relative subpopulation abundance (defined by CD34 and CD38) within viable cell fraction was assessed by flow cytometry.

B) Dose-escalation curve for volasertib on bulk and subgated LSC-containing CD34$^{+}$CD38$^{-}$ OCI-AML8227 (n=2, 72 h).

C) OCI-AML22 cells were treated with the indicated PLK1 inhibitors at indicated doses for 72 h and viability relative to DMSO (%7AAD$^{-}$AnnexinV$^{-}$) and relative subpopulation abundance (defined by CD34 and CD38) within viable cell fraction was assessed by flow cytometry.

D) Dose-escalation curve for volasertib on bulk and LSC-containing CD34+CD38–OCI-AML22 (n=2; 72 h).

### Supplementary Figure 2

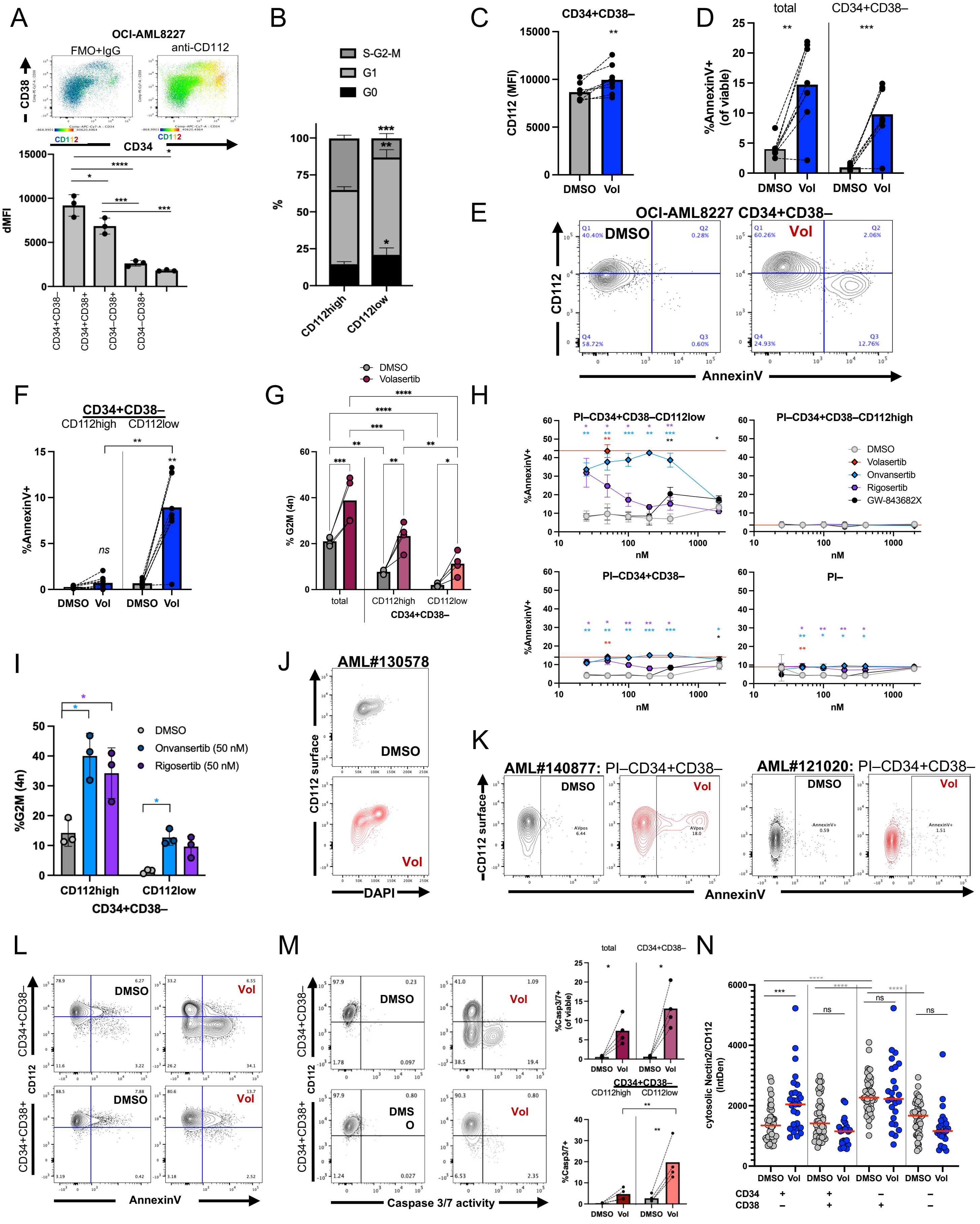

**Suppl. Fig.2**

A) CD112 surface staining overlayed onto CD34/CD38 profile of OCI-AML8227 (flow cytometry) and delta median fluorescence intensity (dMFI) of CD112 of subgated 4 CD34/CD38 fractions (n=3).

B) Prospectively isolated OCI-AML8227 PI–CD34+CD38– CD112$^{high}$ (top 25%) and CD112low (bottom 25%) cells were subjected to cell cycle analysis (Ki67, Hoechst; n=4).

C) CD112 Median fluorescence intensity (MFI) on OCI-AML8227 PI$^{-}$AnnexinV$^{-}$CD34+CD38– cells upon DMSO/volasertib treatment (24 hrs, 50 mM, n=8).

D) Flow cytometric analysis of %AnnexinV+ among total viable (PI–) and CD34+CD38$^{-}$ subgated OCI-AML8227 cells after 24 hrs of DMSO/Volasertib treatment (50 nM; n=8).

E) Representative flow cytometric analysis plot showing gating strategy and effects of DMSO/Volasertib treatment on the CD34+CD38– cell fraction of OCI-AML8227.

F) Flow cytometric analysis of %AnnexinV+ among CD112$^{high}$ and CD112$^{low}$ subgated PI–CD34+CD38–OCI-AML8227 cells after 24 hrs of DMSO/Volasertib treatment (50 nM; n=8).

G) DNA content analysis by flow cytometry of 24 hr DMSO/volasertib treated OCI-AML22 that were surface antibody stained, fixed, permeabilized and DAPI stained (n=4). Analysis of total cells and CD34+CD38– cells sub-fractionated into CD112$^{high}$ (top 25%) and CD112$^{low}$ (bottom 25%) populations analysed for cells in G2/M-phase according to 4n DNA content.

H) Flow cytometric analysis of %AnnexinV+ among CD112$^{low}$ and CD112$^{high}$ subgated PI–CD34+CD38–, PI–CD34+CD38– or PI– OCI-AML22 cells after 24 hrs of DMSO/volasertib/onvansertib/rigosertib/GW-843682X treatment (at indicated concentrations; n=3; redlines highlight mean of 50 nM volasertib).

I) DNA content analysis by flow cytometry of 24 hr DMSO/volasertib/onvansertib/rigosertib treated OCI-AML22 that were surface antibody stained, fixed, permeabilized and DAPI stained (n=3). Analysis of CD34+CD38– cells sub-fractionated into CD112$^{high}$ and CD112$^{low}$ populations analysed for cells in G2/M-phase according to 4n DNA content.

J) Flow cytometric analysis of cultured and 48 hr DMSO/volasertib treated primary AML patient sample AML#130578. Cells were surface antibody stained, fixed, permeabilized and stained with DAPI.

K) Flow cytometric analysis of indicated AML patent samples (n=2) after 48 hr DMSO/ 50 nM volasertib treatment that were subgated for PI–CD34+CD38– populations.

L) Representative flow cytometric analysis plot of 48hr treated OCI-AML22 subgated into CD34+CD38– and CD34+CD38+ cells and analysed for CD112 surface levels and AnnexinV.

M) Flow cytometric analysis of %CellEvent Caspase 3/7 Green+ cells vs CD112 surface staining in subgated PI–CD34+CD38– and PI–CD34+CD38+ OCI-AML22 cells and quantification of %Caspase 3/7 activity+ cells in PI–CD34+CD38–CD112$^{low}$ and CD112$^{high}$, PI–CD34+CD38–, or total (PI–)CD34+CD38 OCI-AML22 cells after 48 hrs of DMSO/volasertib treatment (n=4).

N) Quantification of Nectin-2/CD112 as determined by confocal analysis of DMSO/volasertib treated and sorted OCI-AML8227 (23-50 cells per population and condition) after exclusion of cells in mitotic catastrophe.

### Supplementary Figure 3

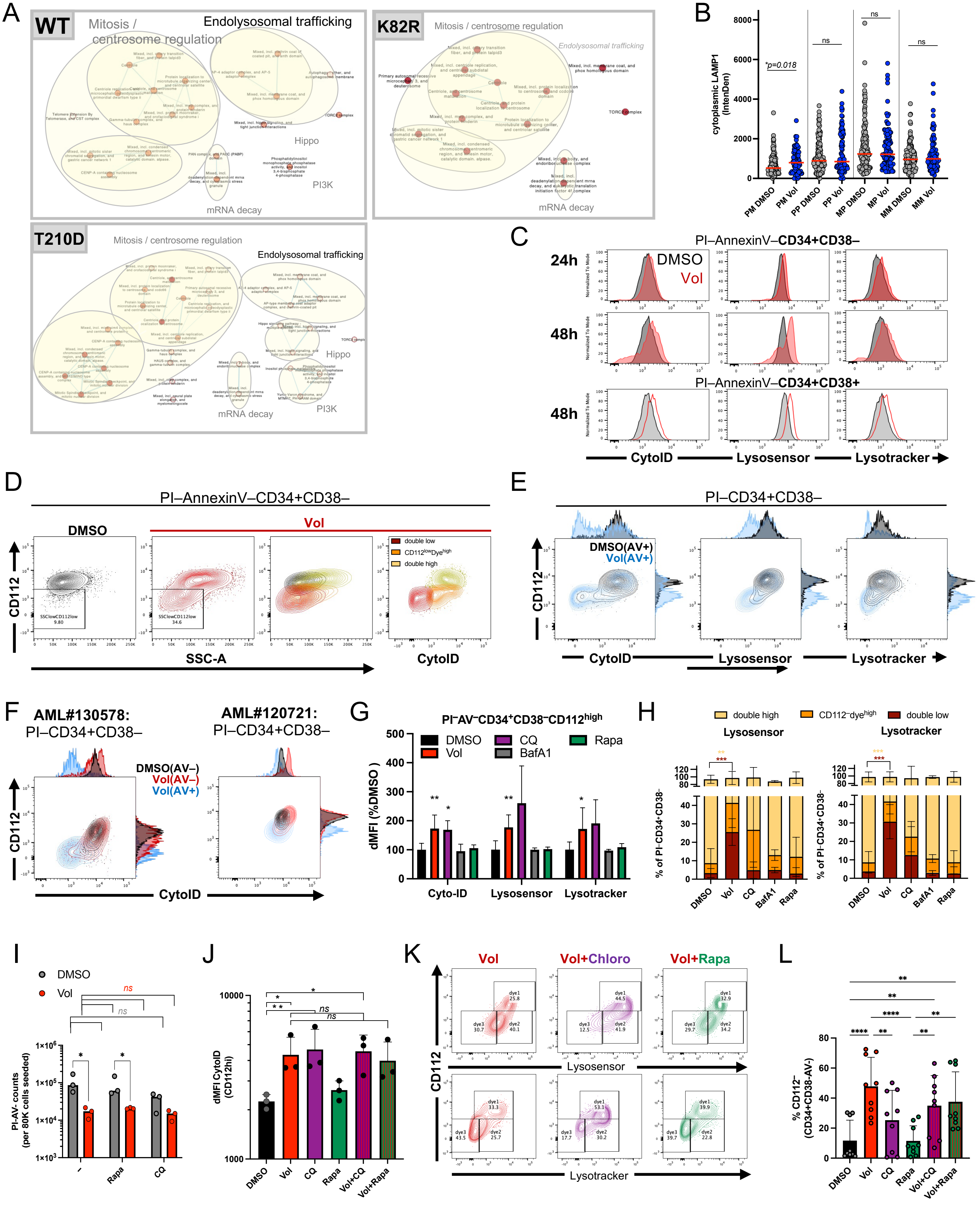

**Suppl. Fig. 3**

A) Enrichment map of GOBP gene set annotations enriched in the BioID data according to PLK1 construct using Cytoscape.
B) Confocal analysis of LAMP1 in 48hrs treated and CD34/CD38 sorted OCI-AML8227 populations.
C) CytoID, Lysosensor and Lysotracker signal in OCI-AML22 PI–AnnexinV–CD34+CD38– cells after 24 and 48 hrs of DMSO/ volasertib treatment.
D) Granularity assessment by flow cytometry using side scatter (SSC) and superimposing subgated populations as delineated in the right panel.
E) Superimposed pre-apoptotic (AV+) $PI^{-}CD34^{+}CD38^{-}$ OCI-AML22 cells after 48 hrs DMSO/Volasertib treatment.
F) Flow cytometric representation Cyto-ID assay of primary AML patient samples cultured and treated with DMSO/ volasertib for 48 hrs and subgated for $PI^{-}CD34^{+}CD38^{-}$ and either $AnnexinV^{+}$ or $AnnexinV^{-}$ as indicated (n=3-7).
G) Summary of flow cytometric analysis of Cyto-ID, Lysosensor and Lysotracker dMFI relative to DMSO controls within subgated OCI-AML22 $PI^{-}AnnexinV^{-}$ $CD34^{+}CD38^{-}CD112^{high}$ cells after treatment with the indicated drugs for 48 hrs.
H) Summary data of flow cytometric analysis of Lysosensor and Lysotracker signals vs CD112 staining on subgated OCI-AML22 $PI^{-}AnnexinV^{-}$ $CD34^{+}CD38^{-}$cells after treatment with the indicated drugs for 48 hrs.
I) Viable cell counts of single drug or double drug treated OCI-AML22 after 48 hrs of *in vitro* treatment with the indicated drugs.
J) Delta Median fluorescence intensity (dMFI) for Cyto-ID signal in 48 hrs single and double drug treated $PI^{-}AnnexinV^{-}CD34^{+}CD38^{-}$ OCI-AML22 (n=3).
K) Representative flow cytometric analysis for volasertib only or in combination with chloroquine/CQ or rapamycin/Rapa treated OCI-AML22 cells subjected to Lysotracker or Lysosensor analysis and subgated for $PI^{-}AnnexinV^{-}CD34^{+}CD38^{-}$ cells.
L) Flow cytometric analysis of %induction of a $CD112^{low}$ population within the $P^{I-}AnnexinV^{-}$ $CD34^{+}CD38^{-}$ fraction upon indicated single or double drug treatment for 48 hrs for all 3 dyes combined from n=3.

### Supplementary Figure 4

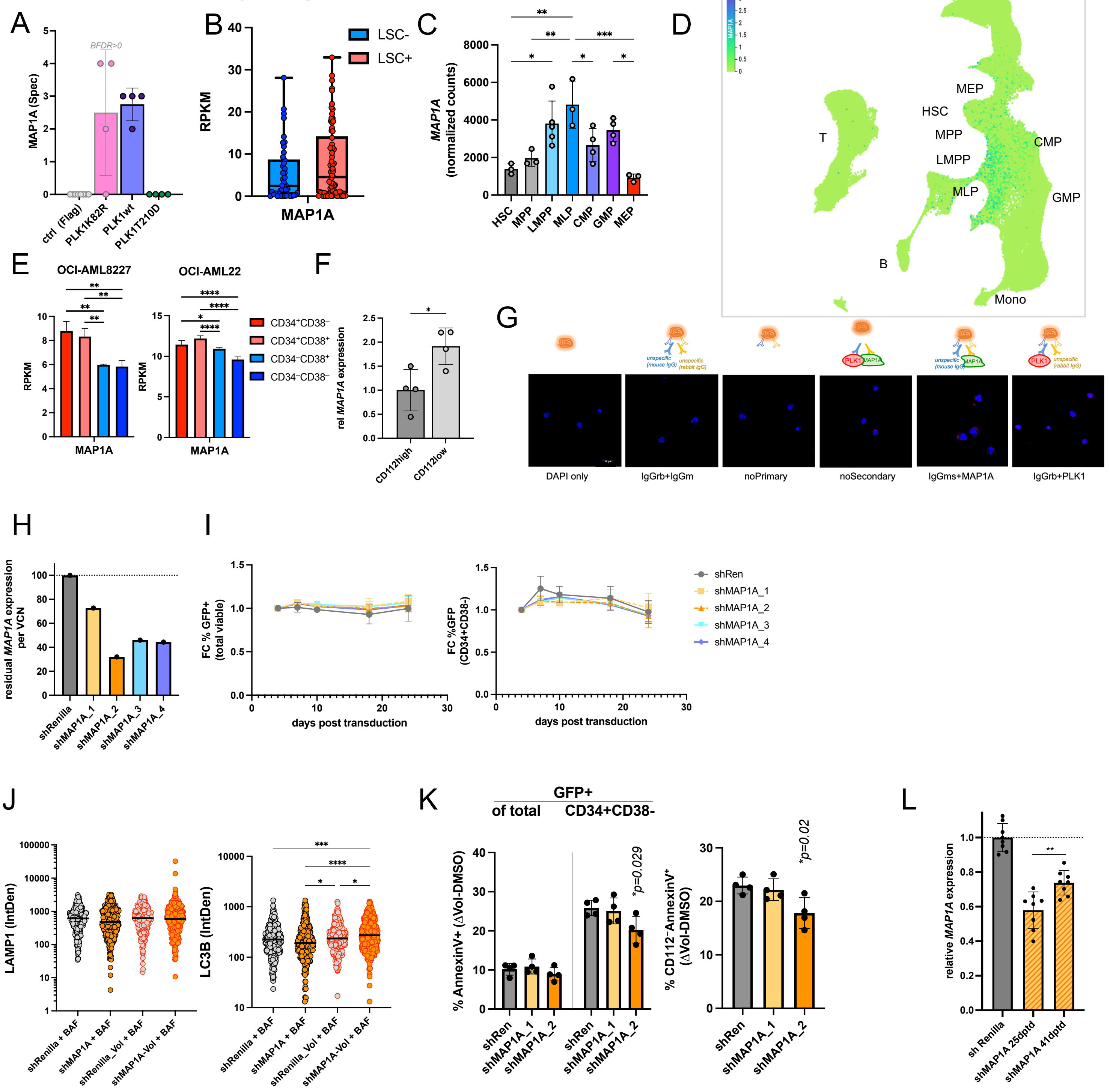

**Suppl. Fig. 4:**

A) Mass spectrometry results for MAP1A in BioID assay (4 measurements from 2 biological and 2 technical replicates each).
B) *MAP1A* expression in bulk RNAseq data generated from sorted and functionally validated LSC– (non-engrafting) and LSC+ (engrafting) fractions of primary AML patient samples (as in[4]).
C) *MAP1A* transcript expression across normal haematopoietic progenitors (RNAseq[40])
D) *MAP1A* expression in human haematopoietic reference map[6].
E) *MAP1A* expression in bulk RNAseq of CD34/CD38 sorted OCI-AML8227 and OCI-AML22 subpopulations[53].
F) *MAP1A* expression on sorted $PI^{-}CD34^{+}CD38^{-}$ OCI-AML22 cells subfractionated into top 25% $CD112^{high}$ and bottom 25% $CD112^{low}$ subpopulations according to digital droplet PCR.
G) Representative images of negative controls with various primary antibody combinations to assess PLA background signal (total of 375 control cells acquired).
H) Knockdown efficiency measured by residual *MAP1A* expression on day 18 post transduction according to digital droplet PCR performed on shRNA- transduced OCI-AML22. Vector copy number (VCN) was estimated by transduction efficiency (%$GFP^{+}$) and assuming Poisson distribution.
I) Flow cytometric monitoring of %$GFP^{+}$ cell marking in total OCI-AML22 cell population and within CD34+CD38– fraction of 4 independent shRNAs targeting *MAP1A and a* control shRNA targeting Renilla luciferase *(shRen)* (n=4 individual transductions per construct).
J) Confocal analysis for LAMP1 and LC3B abundance in prospectively sorted and DMSO/volasertib treated $GFP^{+}CD34^{+}CD38^{-}$ cells in the context of shRenilla or shMAP1A that were additionally treated with 20 nM bafilomycin A1/BafA1 2 hours before fixation. (shMAP1A_1+_2 combined; Ø 100 cells from 3 individually transduced and sorted cultures; 2 weeks post transduction).
K) % AnnexinV+ cells in indicated subsets of shRNA transduced OCI-AML22 treated with volasertib vs DMSO (24h) after additional 3 weeks of propagation culture (n=4; 5-6 weeks post transduction).
L) Digital droplet PCR for *MAP1A* transcript expression in shRNA transduced and $GFP^{+}$ sorted OCI-AML22 harvested at 25 d and 41 d post transduction (n=8 individual transductions).

### Supplementary Figure 5

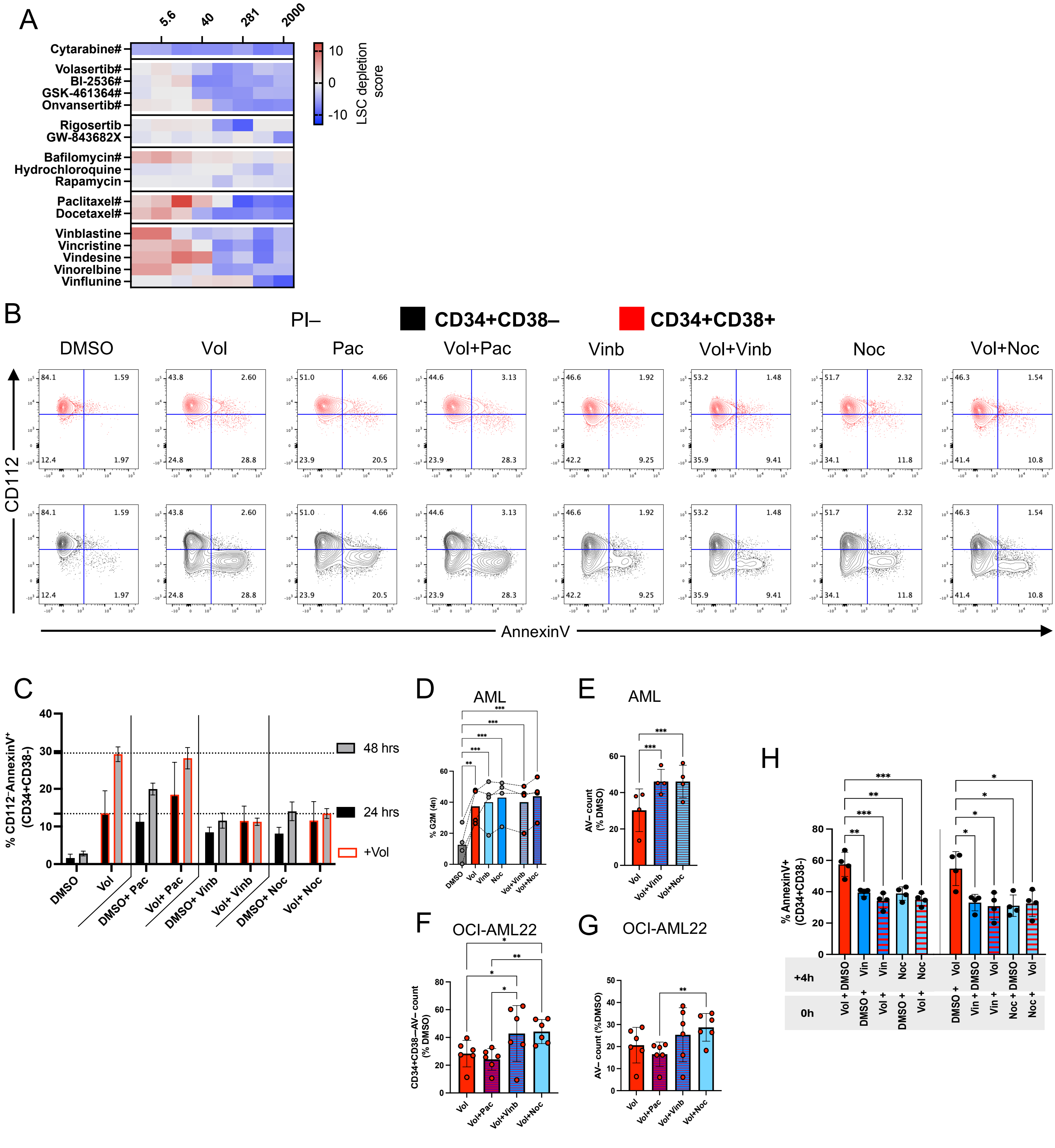

**Suppl. Fig.5:**

A) Excerpt and summary of selected drugs from the two iterative LSC stem cell drug screens as described in Fig. 1. #: drugs in first and second screen.
B) Representative flow cytometric plots for treated OCI-AML22 with the indicated drugs/ single / double treatment and their effect on apoptosis (in $CD34^{+}CD38–$ / black vs $CD34^{+}CD38^{+}$/ red overlay).
C) Apoptosis induction in $CD34^{+}CD38^{-}$ OCI-AML22 cells after single/double drug treatment at 24 and 48 hrs (n=4).
D) DNA content analysis by flow cytometry of cultured and 24 hr as indicated treated primary AML patient samples (n=4).
E) Viable ($PI^{-}AnnexinV^{-}$) cell counts relative to DMSO controls after 48 hrs of indicated treatment of 4 cultured AML patient samples.
F) Viable ($PI^{-}AnnexinV^{-}$) $CD34^{+}CD38^{-}$ cell counts relative to DMSO controls after 48 hrs of indicated treatment of OCI-AML22 (n=6).
G) Viable ($PI^{-}AnnexinV^{-}$) total cell counts relative to DMSO controls after 48 hrs of indicated treatment of OCI-AML22 (n=6).
H) OCI-AML22 were with 4 hrs delay sequentially treated with the indicated drug combinations and after a total of 48 hrs analysed for apoptosis induction (%$AnnexinV^{+}$) in the $CD34^{+}CD38^{-}$ fraction.
