## Supplementary Table 3 for "Cell-state–dependent responses to PLK1 inhibition reveal a non-canonical microtubule-endolysosomal vulnerability in quiescent leukemia stem cells"

| | A | B | C | D | E | F | G | H | I | J | K | L | M | N | O | P |
|---|---|---|---|---|---|---|---|---|---|---|---|---|---|---|---|---|
| 1 | | | cell dose | cell dose | # mice | # mice | 1/(stem cell frequency) - actual cell dose | | | | | 1/(stem cell frequency)- adjusted cell dose | | | | |
| 2 | AML ID | group | transplanted | adjusted | transplanted | engrafted | Lower | Estimate | Upper | Fold change | P.value | Lower | Estimate | Upper | Fold change | P.value |
| 3 | | | 12000 | 12000 | 8 | 1 | | | | | | | | | | |
| 4 | | vehicle | 60000 | 60000 | 8 | 5 | 149,752 | 66,247 | 29,306 | | | 149,752 | 66,247 | 29,306 | | |
| 5 | | | 3000000 | 3000000 | 4 | 4 | | | | | | | | | | |
| 6 | #150019 | | 12000 | 20753 | 8 | 0 | | | | -25.33 | 8.45E-06 | | | | -43.82 | 3.42E-07 |
| 7 | | volasertib | 60000 | 103767 | 7 | 1 | 5,333,513 | 1,677,800 | 527,797 | | | 9,227,627 | 2,902,818 | 913,166 | | |
| 8 | | | 3000000 | 5188327 | 4 | 3 | | | | | | | | | | |
| 9 | | | 1000 | 1000 | 2 | 0 | | | | | | | | | | |
| 10 | | vehicle | 5000 | 5000 | 8 | 0 | 48,377 | 23,149 | 11,077 | | | 48,377 | 23,149 | 11,077 | | |
| 11 | | | 20000 | 20000 | 10 | 7 | | | | | | | | | | |
| 12 | #130578 | | 1000 | 1687 | 2 | 0 | | | | -2.15 | 2.18E-01 | | | | -3.63 | 3.78E-02 |
| 13 | | volasertib | 5000 | 8435 | 8 | 0 | 131,741 | 49,833 | 18,850 | | | 222,258 | 84,072 | 31,801 | | |
| 14 | | | 20000 | 33742 | 10 | 4 | | | | | | | | | | |
| 15 | | | 5000 | 5000 | 5 | 2 | | | | | | | | | | |
| 16 | | vehicle | 20000 | 20000 | 5 | 5 | 16,588 | 6,849 | 2,828 | | | 16,588 | 6,849 | 2,828 | | |
| 17 | | | 100000 | 100000 | 5 | 5 | | | | | | | | | | |
| 18 | | | 500000 | 500000 | 5 | 5 | | | | | | | | | | |
| 19 | #150279 | | 5000 | 12185 | 5 | 2 | | | | -1.67 | 4.28E-01 | | | | -4.06 | 2.96E-02 |
| 20 | | volasertib | 20000 | 48741 | 5 | 4 | 27,336 | 11,404 | 4,758 | | | 66,620 | 27,792 | 11,594 | | |
| 21 | | | 100000 | 243704 | 5 | 5 | | | | | | | | | | |
| 22 | | | 500000 | 1218522 | 5 | 5 | | | | | | | | | | |
| 23 | | | 15000 | 15000 | 10 | 1 | | | | | | | | | | |
| 24 | | vehicle | 75000 | 75000 | 8 | 4 | 208,163 | 98,079 | 46,211 | | | 208,163 | 98,079 | 46,211 | | |
| 25 | | | 150000 | 150000 | 2 | 2 | | | | | | | | | | |
| 26 | #160376 | | 15000 | 45614 | 10 | 1 | | | | -5.12 | 2.49E-02 | | | | -15.57 | 1.28E-04 |
| 27 | | volasertib | 75000 | 228071 | 8 | 1 | 2,069,106 | 502,015 | 121,801 | | | 6,292,036 | 1,526,600 | 370,390 | | |
| 28 | | | 150000 | 456142 | 2 | 0 | | | | | | | | | | |
| 29 | | | 1000 | 1000 | 2 | 1 | | | | | | | | | | |
| 30 | | vehicle | 10000 | 10000 | 8 | 1 | 76,221 | 37,681 | 18,628 | | | 76,221 | 37,681 | 18,628 | | |
| 31 | | | 40000 | 40000 | 8 | 5 | | | | | | | | | | |
| 32 | | | 100000 | 100000 | 2 | 2 | | | | | | | | | | |
| 33 | #141104 | | 1000 | 5962 | 2 | 0 | | | | 0.53 | 2.05E-01 | | | | -3.18 | 2.46E-02 |
| 34 | | volasertib | 10000 | 59621 | 8 | 3 | 39,263 | 20,069 | 10,258 | | | 234,087 | 119,651 | 61,158 | | |
| 35 | | | 40000 | 238483 | 8 | 7 | | | | | | | | | | |
| 36 | | | 100000 | 596208 | 2 | 2 | | | | | | | | | | |
