## Supplementary Table 2 for "Cell-state–dependent responses to PLK1 inhibition reveal a non-canonical microtubule-endolysosomal vulnerability in quiescent leukemia stem cells"

| | A | B | C | D | E | F | G | H | I | J | K | L | M | N | O | P | Q | R |
|---|---|---|---|---|---|---|---|---|---|---|---|---|---|---|---|---|---|---|
| 1 | | | previously assessed functional parameters | | | | | | | | | | | | | | | |
| 2 | **SampleID** | **Exp** | **LSC frequency in CD34+CD38-** | **Engraftment (CD34/CD38 fractions)** | **% CD34+ invitro assays** | **in vitro enrichment/depletion** | **SEX** | **AGE.AT.AML.DX** | **DE.NOVO.vs.SECONDARY** | **TYPE.OF.SAMPLE** | **SAMPLE.MATERIAL** | **FAB** | **WBC.count at.AML.dx** | **BM.blast. at.AML.dx** | **Cytogenetics.at.AML.dx** | **NPM1.at.diag** | **FLT3.ITD.at.diag** | **FLT3.TKD.at.diag** |
| 3 | 130578 | *in vivo / in vitro* | <1/7,000 | 34+ only | 60% | CD34+ | M | 62.3 | de novo | diagnosis | PB | M4 | 155.3 | 63 | 46,XY[20] | negative | negative | negative |
| 4 | 141104 | *in vivo* | 1/12,717 | 3 Fractions | | N/A | F | 67.2 | de novo | diagnosis | PB | nd | 99.8 | 94 | 46,XX[10] | positive | intermediate | negative |
| 5 | 150860 | *in vivo* | N/A | 34+ only | | N/A | M | 54.6 | (MPN, | relapse1 | PB | nd | 114 | nd | 46 XY [8] | negative | positive | nd |
| 6 | 151292 | *in vivo* | <1/10,000 | All Fractions | | N/A | F | 54.3 | de novo | diagnosis | PB | nd | 74.2 | 80 | 46,XX[20] | negative | intermediate | negative |
| 7 | 121020 | *in vivo / in vitro* | <1/10,000 | All Fractions | 33% | AnnexinV depletion | M | 33.4 | de novo | diagnosis | PB | M4 | 150.8 | 90 | 46,XY[20] | positive | high | negative |
| 8 | 150019 | *in vivo* | 1/10,735 | 34+ only | | N/A | M | 37 | de novo | diagnosis | PB | nd | 20.8 | 90 | 46,XY,del(1)(p13p31),del(5)(q13),del(11)(q21q23),del(13)(q12q22)[9]/46,XY[1] | Negative | Negative | Negative |
| 9 | 140877 | *in vivo / in vitro* | <1/3,000 | nd | 35% | CD34+/ Annexin V depl | F | 28 | de novo | diagnosis | PB | nd | 187.4 | 80 | 46,XX[20] | Positive | Positive | Negative |
| 10 | 150279 | *in vivo* | N/A | nd | | N/A | F | 63 | de novo | diagnosis | PB | nd | 227.1 | nd | Not assessable | Positive | Positive | Negative |
| 11 | 160376 | *in vivo* | N/A | nd | | N/A | M | 45 | de novo | diagnosis | PB | nd | 97 | 48 | 46,XY,t(3;3)(q21;q26.2),ins(7;12)(p15;q1?1q13),der(12)del(12)(p11.2p12)ins(7;12)[13] | nd | nd | nd |
| 12 | 120721 | *in vivo / in vitro* | N/A | nd | 90% | Annexin V depletion | M | 36 | secondary (MDS) | refractory | PB | nd | 4.6 | 51 | 46,XY,t(1;3)(q32;q26~27),del(20)(q13.1)[11] | nd | nd | nd |
