## Supplementary Table 1 for "Cell-state–dependent responses to PLK1 inhibition reveal a non-canonical microtubule-endolysosomal vulnerability in quiescent leukemia stem cells"

| | A | B | C | D | E | F | G | H | I | J | K | L |
|---|---|---|---|---|---|---|---|---|---|---|---|---|
| 1 | | | viable t_score at different concentration (nM) | | | | | | | | | |
| 2 | drug.ID | Drug name | 1.875 | 5.625 | 15.000 | 40.000 | 93.750 | 281.250 | 750.000 | 2000.000 | LSC depletion score | Target(s) |
| 3 | 1 | 2-Thio(3-iodobenzyl)-5-(1-pyridyl)-[1,3,4]-oxadiazole, GSK-3 inhibitor II | -0.47159 | -0.54383 | -0.56373 | -0.14003 | -1.74853 | -0.47648 | -2.12222 | -0.80147 | -0.43562 | GSK3B |
| 4 | 2 | AS-601245 | 0.50166 | 1.03593 | -0.48292 | -2.89262 | -1.69061 | -2.41444 | -1.99276 | -6.11521 | -0.89123 | JNK1 (JNK) |
| 5 | 3 | NU-6102 | -0.79947 | 0.55269 | -1.10167 | 0.81353 | -0.09633 | -0.26679 | 0.80448 | -0.22472 | -0.02019 | CDC2 (CDK1, p34cdc2, CDC28p)/Cyclin B, CDK2/Cyclin A |
| 6 | 4 | BX-795 | -1.65947 | -0.24918 | 0.46686 | 0.01444 | 1.67983 | 0.31539 | -0.05970 | -0.41064 | 0.00619 | PDK1 (PDPK1), TBK1, IKKe |
| 7 | 5 | CHIR-98014 isomer, CT-98014 | 0.43647 | -0.76116 | -0.64713 | 0.59311 | -0.14896 | -1.71256 | -1.34886 | -4.83216 | -0.53415 | GSK3 |
| 8 | 6 | CT-99021, CHIR-99021 | -2.17873 | 0.07338 | 0.41248 | 1.75187 | 0.59789 | 0.16175 | -0.36978 | -2.04183 | -0.10104 | GSK3 |
| 9 | 7 | GDC-0941 bismesylate, RG-7321, Pictilisib | 0.95397 | 0.20923 | 1.10422 | 1.79043 | 1.46398 | 5.05904 | 7.12557 | 6.73330 | 1.55018 | PI3K |
| 10 | 8 | GSK-269962A | -0.78040 | -1.79276 | -1.29347 | -1.59638 | -2.85375 | 1.79456 | 1.04910 | 1.96293 | -0.22265 | ROCK1, ROCK2 (ROCKa) |
| 11 | 9 | GW-843682X | -0.99319 | -0.26318 | -1.10634 | -0.40537 | -1.56281 | 0.47376 | -1.38876 | -6.22658 | -0.72768 | PLK1 (PLK, STPK13), PLK3 (PRK, FNK, CNK) |
| 12 | 10 | MLN-518, CT53518, Tandutinib | 0.20125 | -1.09090 | 0.63425 | 0.35435 | -0.64952 | 0.10865 | -2.35350 | -0.01061 | -0.17798 | FLT3 (STK1, FLK2), KIT (c-KIT), PDGFRb (PDGFR, PDGFR1) |
| 13 | 11 | NSC 625987 | -0.95742 | 1.39553 | -0.24514 | 0.37376 | -1.26982 | -0.33384 | -1.33206 | -3.32219 | -0.36098 | CDK4/Cyclin D1 |
| 14 | 12 | NVP-BEZ235, BEZ235, BEZ-235, Dactolisib | 1.83050 | 0.44844 | -0.22192 | 0.47984 | -1.67528 | 2.13894 | 0.23307 | 3.29175 | 0.41389 | PI3K |
| 15 | 13 | PD-169316 | -2.69166 | -0.16120 | -1.60087 | -0.98590 | -2.65314 | -3.63036 | -3.92996 | -4.32240 | -1.26702 | p38 MAPK |
| 16 | 14 | PD-180970 | 0.34613 | 0.07721 | -1.28884 | -1.80168 | -2.79690 | -4.74749 | -6.12800 | -5.45602 | -1.38247 | ABL1 (ABL), SRC (c-SRC) |
| 17 | 15 | PD-184352, CI-1040 | -0.19569 | 0.47248 | 0.50995 | 1.41115 | -0.24251 | 4.17869 | 3.14488 | 2.25269 | 0.73144 | MAP2K1 (MEK1), MAP2K2 (MEK2), Erk2 (ERK, p38), RAF1 (c-Raf) |
| 18 | 16 | PIK-90 | -0.95100 | -0.42639 | 1.51888 | 1.89770 | 0.10561 | 2.95601 | 3.47960 | 3.84878 | 0.78837 | PI3Ka |
| 19 | 17 | PIK-75 hydrochloride | -0.21417 | 0.96120 | 0.47075 | -0.67217 | -5.62470 | -4.71807 | -7.22527 | -13.28113 | -1.92212 | PI3K |
| 20 | 18 | Tyrphostin AG-1296 | -2.26238 | -1.17126 | 0.44030 | 0.77649 | 0.40740 | 1.60347 | 1.42619 | -1.15826 | 0.00393 | PDGFRb (PDGFR, PDGFR1) |
| 21 | 19 | AS604850, AS-604850, AS 604850 | -0.09865 | -1.03077 | -0.94623 | -0.43896 | -1.18653 | -0.15326 | -0.16463 | -1.10683 | -0.32513 | PI3K |
| 22 | 20 | JNJ-10198409, RWJ-540973 | -0.55148 | -1.08797 | -0.20920 | -2.19722 | -7.15113 | -4.25594 | -4.51426 | -5.82686 | -1.63608 | PDGFRb (PDGFR, PDGFR1) |
| 23 | 21 | NH125 | -0.42749 | 0.22184 | -0.06694 | -0.10014 | 0.75371 | 3.24096 | -9.56186 | -8.78027 | -0.93368 | eEF2K activator |
| 24 | 22 | PI-103 | -0.47885 | -0.79581 | 0.37970 | 1.25545 | -1.33691 | 2.49906 | 6.17760 | 2.45989 | 0.64444 | PI3K |
| 25 | 23 | Axitinib, AG-013736, AG-13736, Inlyta | -1.83945 | -0.63826 | -0.58887 | 0.12812 | -1.09909 | 0.85071 | -1.10813 | 0.70582 | -0.22766 | PDGFRb (PDGFR, PDGFR1), FLT1 (VEGFR1), KDR (VEGFR2, VEGFR, FLK1), FLT4 (VEGFR3), FMS (c-FMS, CSF-1R, CSF1R) |
| 26 | 24 | Gefitinib, Iressa, ZD1839, ZD-1839 | -0.12207 | -0.77061 | -0.59643 | 0.85677 | -1.01959 | -0.18199 | -0.03362 | 0.96161 | -0.05746 | EGFR (ERBB1, HER1), ErbB2 (TKR1, HER2, NEU) |
| 27 | 25 | Lapatinib ditosylate, Tykerb, GW572016 | -1.02000 | 0.30828 | 0.16577 | 0.18815 | -3.39962 | -0.56666 | 0.36272 | -0.53365 | -0.28511 | EGFR (ERBB1, HER1), ErbB2 (TKR1, HER2, NEU) |
| 28 | 26 | Vatalanib dihydrochloride, PTK-787, ZK-222584, CGP79787D | 0.03794 | 1.62704 | 0.65729 | 0.22848 | -1.02254 | -2.53960 | -3.86027 | -3.81788 | -0.55117 | KDR (VEGFR2, VEGFR, FLK1), PDGFRb (PDGFR, PDGFR1) |
| 29 | 27 | Rapamycin, Sirolimus, Rapamune | -1.02417 | -0.26842 | -0.71148 | -0.33776 | -3.08784 | -0.63990 | -1.18743 | -1.55769 | -0.55910 | FRAP (MTOR, FRAP1), CCR5 expression inhibitor |
| 30 | 28 | VX-680, MK-0457, Tozasertib, VX6 | -0.15551 | -0.04697 | 0.28368 | 3.23956 | -0.35574 | -9.06967 | -4.88689 | -5.00397 | -1.01457 | AurA (AuroraA), AurB (AuroraB), AurC (AuroraC), ABL1 (ABL), FLT3 (STK1, FLK2), JAK2 |
| 31 | 29 | Vandetanib, Zactima, ZD6474, AZD-6474 | -1.27462 | 0.11297 | -1.20443 | -1.58860 | -4.52559 | -0.93279 | -1.00882 | 2.02547 | -0.53257 | EGFR (ERBB1, HER1), KDR (VEGFR2, VEGFR, FLK1), RET, ABL1 (ABL), KIT (c-KIT), FLT1 (VEGFR1), FLT4 (VEGFR3), TRKA (TRK) |
| 32 | 30 | Everolimus, RAD-001, Certican | -2.63751 | -1.21711 | -1.15089 | -0.83792 | -0.79777 | 0.11205 | -1.52073 | 0.92908 | -0.45166 | FRAP (MTOR, FRAP1), FKBP12 (Rotamase), Angiogenesis inhibitor |
| 33 | 31 | Fasudil HCl, AT-877, HA-1077 (diHCl), Eril | -0.13838 | 0.24057 | -1.68566 | 0.06136 | -1.45579 | -0.60485 | -2.01685 | -1.33543 | -0.43988 | ROCK1, ROCK2 (ROCKa), Calcium sensitizer |
| 34 | 32 | Imatinib Mesylate, Gleevec, Glivec, CGP-57148B, STI-571 | -0.83966 | -0.92272 | 0.87714 | 0.87972 | -1.01217 | 0.11271 | 0.62962 | -0.76149 | -0.06577 | BCR, ABL1 (ABL), PDGFRb (PDGFR, PDGFR1), KIT (c-KIT) |
| 35 | 33 | K-252a, K-2151 | 0.31039 | 0.44777 | -1.14413 | 0.70435 | -5.69789 | -6.31117 | -3.81902 | -11.54201 | -1.71586 | TRKA (TRK), Na+/H+ Exchanger (NHE) Inhibitor |
| 36 | 34 | Lestaurtinib, CEP-701, KT-5555, SPM-924 | -0.46004 | 0.79415 | -0.27099 | -1.23022 | -3.75333 | -7.90925 | -11.50696 | -12.48879 | -2.33579 | FLT3 (STK1, FLK2), TRKA (TRK), JAK2 |
| 37 | 35 | LY-294002, LY-924002, SF-1101 | 0.34712 | 0.24734 | 1.27465 | 0.17907 | -0.71549 | -1.67106 | 1.25292 | -0.64478 | 0.01711 | PI3K |
| 38 | 36 | Go 6976, PD-406976, Go-6976, Goe-6976 | -0.35847 | 0.50146 | -0.10267 | 1.93625 | -1.78392 | 1.22429 | -4.50045 | 3.60503 | 0.03308 | PKC, JAK2 |
| 39 | 37 | PD-98059 | -1.40557 | -0.00545 | -1.57455 | 0.86582 | -2.29867 | -1.81871 | -2.86318 | -2.22285 | -0.71821 | MAP2K1 (MEK1), MAP2K2 (MEK2), Erk1 |
| 40 | 38 | PKC-412, CGP41251, Midostaurin | 0.28828 | -0.30949 | 0.31686 | 1.63168 | -0.53735 | -4.30055 | -8.90763 | -7.09735 | -1.19979 | FLT3 (STK1, FLK2) |
| 41 | 39 | Roscovitine, CYC202, Seliciclib | 0.19746 | 0.74593 | -0.65083 | -0.13611 | 1.01774 | -0.26196 | -1.18211 | -2.53684 | -0.17803 | CDC2 (CDK1, p34cdc2, CDC28p)/Cyclin B, CDK2/Cyclin A, CDK2/Cyclin E, CDK5/p35 |
| 42 | 40 | FK-506, Tacrolimus, Fujimycin, Prograf | 0.66170 | 0.14309 | 0.18717 | 0.98310 | 0.68621 | -0.73990 | -0.09321 | -0.88438 | 0.05986 | Calcineurin (CN, PP-2B), INS expression inhibitor, FKBP12 (Rotamase) |
| 43 | 41 | Erlotinib HCl, CP-358774, OSI-774, Tarceva, NSC-718781, RG-1415, Ro-50-8231 | -0.18364 | 1.17467 | -1.94057 | 1.01977 | -0.43260 | -0.34705 | -1.00657 | -1.37951 | -0.19634 | EGFR (ERBB1, HER1), ErbB2 (TKR1, HER2, NEU) |
| 44 | 42 | Staurosporine, AM-2282 | 1.09199 | 1.15045 | -2.58249 | -6.55752 | -5.27457 | -4.67056 | -8.02838 | -8.81638 | -2.13675 | Pan kinase |
| 45 | 43 | Wortmannin, KY 12420 | -0.75752 | 0.54905 | 2.76350 | 2.09557 | 0.63837 | 2.33753 | 1.56649 | 4.21946 | 0.85073 | PI3K - Irreversible |
| 46 | 44 | 6-bromoindirubin-3'-oxime, BIO, 6-BIO, MLS-2407 | -0.96835 | -1.55933 | -1.51577 | -2.64311 | -6.29352 | -8.78372 | -9.49586 | -12.75332 | -2.79169 | GSK3 |
| 47 | 45 | AR-A014418 | -1.12905 | 0.49441 | -0.36825 | -0.84816 | -2.22616 | -0.20099 | 0.45074 | -0.84400 | -0.29631 | GSK3 |
| 48 | 46 | BAY 61-3606 hydrochloride, sc-202351, BAY-61-3606 | 0.18933 | 0.57207 | -1.80590 | -0.30394 | -2.50134 | -1.45296 | -3.21714 | -6.61534 | -0.96001 | SYK |
| 49 | 47 | BMS-345541 | -0.02128 | -1.37387 | 0.48788 | -1.27909 | -0.12095 | -1.84430 | -3.59239 | -2.69239 | -0.66197 | IKKb (IKK2) |
| 50 | 48 | CGP-74514A hydrochloride | -0.36012 | 0.50465 | -0.05993 | 0.75680 | -0.68476 | -0.89683 | -1.00786 | -3.28409 | -0.31918 | CDC2 (CDK1, p34cdc2, CDC28p)/Cyclin B |
| 51 | 49 | GW2974, GW-2974, GW-282974 | -0.43767 | -1.65720 | 0.87087 | 1.60693 | -1.32573 | -1.51848 | -1.29779 | -3.14403 | -0.43785 | EGFR (ERBB1, HER1), ErbB2 (TKR1, HER2, NEU) |
| 52 | 50 | GW441756 hydrochloride, GW-441756 | -1.30073 | -1.63652 | -3.45526 | -5.62739 | -8.22244 | -7.39153 | -5.17950 | -4.79485 | -2.38544 | TRKA (TRK) |
| 53 | 51 | GW583340 dihydrochloride, GW-583340 | -0.09612 | -1.61529 | -0.69910 | 0.97330 | -0.12268 | -0.40632 | 1.78640 | -1.20753 | -0.08800 | EGFR (ERBB1, HER1), ErbB2 (TKR1, HER2, NEU) |
| 54 | 52 | GW8510, GW-8510 | -0.22631 | 1.40519 | 0.89242 | 2.00553 | -0.32966 | -1.04210 | -1.33025 | -2.26153 | -0.05624 | CDK2/Cyclin A |
| 55 | 53 | IRAK-1/4 Inhibitor I | -0.57341 | -0.77639 | -0.16087 | -0.26034 | 0.32914 | 0.26193 | -3.04598 | -3.65280 | -0.49974 | IRAK1 (IRAK) |
| 56 | 54 | PD-173074 | 0.72519 | 0.87627 | 0.25187 | 0.80327 | -0.09330 | -0.15243 | -0.88012 | 2.36311 | 0.24698 | FGFR1 (FLT2), FGFR2, FGFR3 |
| 57 | 55 | PS1145 dihydrochloride, PS-1145, MLN-1145 | 0.34678 | 0.88718 | -0.43963 | 0.75705 | -1.29890 | -1.75313 | -2.71878 | -3.91116 | -0.51571 | IKKb (IKK2) |
| 58 | 56 | SB-505124 hydrochloride hydrate | -1.41068 | -0.34884 | 1.34931 | 2.13158 | 0.02716 | 2.39450 | -2.11825 | 2.00148 | 0.25538 | TGFbR1 (ALK5), ALK4 (ACVR1B, ACTRIB), ALK7 (ACVR1C) |
| 59 | 57 | SD 208, SD-208 | -0.61091 | -0.09035 | 0.00343 | 0.35591 | -1.45806 | 0.40623 | 0.41002 | -2.96126 | -0.25023 | TGFbR1 (ALK5) |
| 60 | 58 | SIS3, SIS-3 | -0.27363 | -0.08760 | -0.37505 | -0.18346 | -0.59769 | 0.92288 | -0.83844 | -0.60955 | -0.12956 | TGFbR1 (ALK5) |
| 61 | 59 | SKF-86002, SKF 86002, SK&F-86002 | -1.07810 | 0.77431 | -0.99959 | 0.03513 | -0.44461 | -0.46349 | 0.63521 | -1.08862 | -0.16680 | p38 MAPK |
| 62 | 60 | STO-609 acetic acid | -0.81470 | -0.34264 | 0.52887 | 1.82335 | 0.33980 | -0.64733 | -2.10969 | -2.21128 | -0.21779 | CAMKK2 (CAMKK) |
| 63 | 61 | SU 6656, SU-6656 | -2.04226 | -1.11785 | -2.57358 | -0.88133 | -3.90499 | -4.52689 | -2.54267 | -3.85519 | -1.36021 | SRC (c-SRC), YES, LYN, FYN (p59fyn) |
| 64 | 62 | ZM 39923 hydrochloride | -1.24848 | 0.12457 | -0.31090 | 1.12842 | -1.67697 | -0.24886 | -0.31972 | -5.37783 | -0.50298 | JAK3 |
| 65 | 63 | 6-[4-(2-Piperidin-1-ylethoxy)phenyl]-3-pyridin-4-ylpyrazolo[1,5-a]pyrimidine, Dorsomorphin, BML-275, AMPK Inhibitor, Compound C | -1.52948 | -1.14286 | -1.11448 | -1.21416 | 0.25828 | -1.75452 | -1.68819 | -3.63891 | -0.75000 | AMPK, ALK2, BMPR1A (ALK3), BMPR1B (ALK6) |
| 66 | 64 | Chk2 Inhibitor II, 339253, BML-277 | -0.32509 | 1.38888 | -0.39799 | -0.47230 | -1.88660 | -0.63829 | -0.38730 | -2.09252 | -0.30517 | CHK2 |
| 67 | 65 | Compound 52, NG-52, NG 52 | -2.87672 | -0.78070 | -0.83777 | -2.17961 | -5.94136 | -6.87621 | -5.22897 | -7.01740 | -2.01315 | CDC2 (CDK1, p34cdc2, CDC28p)/Cyclin B |
| 68 | 66 | AEG 3482 | -1.07923 | 0.87581 | 0.20879 | -0.32484 | 0.61144 | -0.88442 | 1.23313 | -0.13732 | 0.03193 | JNK1 (JNK) |
| 69 | 67 | BIBU 1361 dihydrochloride | -0.98686 | 0.55468 | 1.16681 | 1.16384 | 0.62710 | 2.34380 | 2.20839 | 4.45389 | 0.73144 | EGFR (ERBB1, HER1) |
| 70 | 68 | CGP-57380, CGP 57380 | -1.44617 | -2.00151 | -3.01981 | -2.02545 | -2.65482 | -5.30413 | -3.87119 | -3.56945 | -1.51547 | MNK1 |
| 71 | 69 | BIBX 1382 dihydrochloride, Falnidamol | -0.82530 | -0.46884 | -0.63059 | 0.28827 | -0.37566 | -0.79997 | -0.35532 | -2.33563 | -0.34905 | EGFR (ERBB1, HER1) |
| 72 | 70 | GTP 14564, GTP-14564 | -0.82364 | 0.68963 | -0.53582 | -1.72969 | -7.89276 | -8.13644 | -6.99548 | -6.09952 | -1.99951 | FLT3 (STK1, FLK2) |
| 73 | 71 | GW-5074 | 0.43480 | 0.09754 | -1.24794 | 1.60057 | -0.32397 | 1.11576 | 0.18998 | -1.41229 | 0.02882 | RAF1 (c-Raf) |
| 74 | 72 | LY-364947, HTS 466284 | -0.47742 | 1.19061 | -0.37242 | 0.47302 | -0.47717 | 1.62462 | 1.65691 | 0.84108 | 0.28284 | TGFbR1 (ALK5) |
| 75 | 73 | PI-828 | -1.28339 | -0.87055 | 0.58960 | 1.66150 | -1.41020 | 0.31174 | 2.06158 | 1.03306 | 0.13278 | PI3K |
| 76 | 74 | PQ401 | -0.09414 | -0.62253 | 0.30904 | -0.34171 | -0.99254 | -0.15115 | -2.82297 | -3.04035 | -0.49198 | IGF1R (JTK13, IGFIR) |
| 77 | 75 | SB 203580, SB-203580 | -1.13858 | 1.18656 | -0.03028 | 0.28313 | -1.83464 | -1.77748 | -1.07266 | -1.88329 | -0.39752 | p38 MAPK |
| 78 | 76 | SB 216763 | 0.76479 | -0.64112 | 0.46776 | -0.09478 | -1.86951 | 0.73979 | 0.91398 | -1.96953 | -0.10711 | GSK3 |
| 79 | 77 | SB 239063 | 0.99561 | -0.38561 | -1.65903 | 0.04210 | -1.43509 | -0.01124 | 3.29359 | -0.89420 | -0.00342 | p38 MAPK |
| 80 | 78 | SB-415286 | -0.20809 | 0.68182 | -0.63916 | 2.13908 | 1.06714 | -1.09686 | -1.17697 | -1.84296 | -0.06825 | GSK3 |
| 81 | 79 | SB 431542 | -0.15003 | -0.11019 | -0.78142 | 0.34337 | -0.59358 | 1.04102 | 3.12244 | 1.12674 | 0.25361 | TGFbR1 (ALK5) |
| 82 | 80 | SD 169, SD-169 | -0.50901 | -0.13768 | -1.32732 | 0.55052 | -0.35199 | -0.24550 | -1.30696 | -1.31473 | -0.29448 | p38 MAPK |
| 83 | 81 | N-(4-Pyridyl)-N'-(2,4,6-trichlorophenyl)urea, Rho Kinase Inhibitor II | -1.31340 | -0.86003 | 0.15559 | 0.30407 | -0.86831 | -0.28603 | -1.97166 | -2.03069 | -0.43578 | ROCK1, ROCK2 (ROCKa) |
| 84 | 82 | 2-(p-Hydroxyanilino)-4-(p-chlorophenyl) thiazole, HCl, SKI-II | 0.39937 | -1.09995 | 0.78915 | 0.78601 | 0.06242 | -3.02826 | -1.37259 | -0.05079 | -0.22293 | SPK1 (SPHK, SK1, Sphingosine Kinase), Transitional Endoplasmic Reticulum ATPase (VCP, p97) Inhibitor |
| 85 | 83 | 2-Dimethylamino-4,5,6,7-tetrabromo-1H-benzimidazole, DMAT, Casein Kinase II, K25 | -1.55461 | -0.51652 | -2.02323 | -2.28842 | -3.74946 | -3.56700 | -3.76016 | -4.60188 | -1.39932 | CK1a (Casein kinase 1, CK1) |
| 86 | 84 | 5-(3-Methoxy-4-((4-methoxybenzyl)oxy)benzyl)-pyrimidine-2,4-diamine, GW-2580 | 0.13404 | 0.31383 | 0.40345 | -0.08046 | 0.68297 | -0.45182 | 0.76097 | -1.06875 | 0.04403 | FMS (c-FMS, CSF-1R, CSF1R) |
| 87 | 85 | PP-2, PP2, AG-1879 | -0.86791 | -0.44216 | 1.46760 | 0.88633 | -0.27931 | -0.82701 | -0.73527 | -0.53962 | -0.08483 | LCK (p56lck), FYN (p59fyn), HCK (p56hck), SRC (c-SRC) |
| 88 | 86 | TPL2 | 0.10578 | -0.09541 | 0.82634 | 1.07099 | -1.73750 | -2.16451 | -4.31591 | -4.66187 | -0.69595 | COT (TPL2, EST, MAP3K8) |
| 89 | 87 | TWS-119 | -1.58333 | -0.28318 | 0.50280 | 0.94305 | -1.62129 | 1.80063 | 1.15665 | -1.33269 | -0.02647 | GSK3 |
| 90 | 88 | Sorafenib,p-Toluenesulfonate Salt, Bay 43-9006, Nexavar | 0.61571 | -0.85906 | -0.95301 | 0.08266 | -2.60874 | -1.61526 | -3.92043 | -3.35033 | -0.79974 | FLT3 (STK1, FLK2), KIT (c-KIT), PDGFRb (PDGFR, PDGFR1), RET, RAF1 (c-Raf), BRAF (B-raf), FLT1 (VEGFR1), KDR (VEGFR2, VEGFR, FLK1), FLT4 (VEGFR3) |
| 91 | 89 | IC-261, SU-5607 | 0.09306 | 0.76185 | -0.16390 | 0.14538 | -1.18125 | -1.07346 | 0.70064 | -6.95800 | -0.48686 | CK1a (Casein kinase 1, CK1) |
| 92 | 90 | JNJ-7706621 | -0.95656 | -0.53005 | -0.28837 | -0.69668 | -3.20112 | -1.18849 | -1.83668 | -0.50712 | -0.58387 | AurA (AuroraA), AurB (AuroraB), AurC (AuroraC), CDC2 (CDK1, p34cdc2, CDC28p)/Cyclin B |
| 93 | 91 | KN-62 | 0.27663 | 0.35762 | -0.69237 | 0.95197 | -0.23658 | -1.01895 | 0.64120 | -0.47499 | -0.01240 | CAMKK2 (CAMKK) |
| 94 | 92 | KU-55933, KU-0064 | 0.66342 | 0.44531 | -1.24848 | 0.72362 | -0.59593 | -0.43881 | 1.46602 | -0.54277 | 0.02996 | ATM, PI3K |
| 95 | 93 | Kenpaullone, NSC-664704 | 1.10573 | 0.11833 | -2.92778 | -1.35346 | -5.15207 | -3.65221 | -3.32216 | -7.18118 | -1.41857 | CDC2 (CDK1, p34cdc2, CDC28p)/Cyclin B, CDK2/Cyclin A, CDK2/Cyclin E, CDK5/p35, GSK3, LCK (p56lck) |
| 96 | 94 | NU-7026, LY-293646 | -1.56164 | 0.98779 | 0.14259 | -1.14213 | -2.10526 | -0.69532 | -1.18211 | -1.72410 | -0.46177 | DNAPK (DNA-PKcs) |
| 97 | 95 | PD-153035, AG-1517, Compound 32, SU-5271, ZM-252868, WHI-P79 | -1.11321 | -0.91951 | 0.88167 | -0.06773 | -0.52673 | -0.17633 | -0.99841 | -2.77551 | -0.36128 | EGFR (ERBB1, HER1) |
| 98 | 96 | PIM1/2 Kinase Inhibitor VI | -0.66553 | -1.08772 | 0.63463 | 0.01261 | -1.81065 | 0.42158 | 0.21590 | -2.25790 | -0.28778 | PIM1, PIM2 |
| 99 | 97 | Sal003 | -0.68335 | -1.32909 | 0.02084 | 1.13004 | -0.44271 | 0.44938 | -1.31528 | -0.75009 | -0.18523 | eIF2a |
| 100 | 98 | Rho Kinase Inhibitor V | -1.34373 | 0.02379 | -1.34140 | 0.37771 | -1.71073 | -1.39997 | -0.02187 | -4.13130 | -0.60559 | ROCK1, ROCK2 (ROCKa) |
| 101 | 99 | Nilotinib, Tasigna, AMN-107 | 0.87602 | 0.29857 | -0.59093 | 1.15109 | -1.05978 | -1.35921 | -1.68492 | -2.57274 | -0.31346 | BCR, ABL1 (ABL), PDGFRb (PDGFR, PDGFR1), KIT (c-KIT) |
| 102 | 100 | AZD6244, ARRY-142886, AZD-6244, Selumetinib, ARRY-886 | 0.02211 | 0.48149 | 1.26341 | 2.27080 | 1.68438 | 3.13342 | 2.46812 | 3.50589 | 0.94062 | MAP2K1 (MEK1), MAP2K2 (MEK2), Erk2 (ERK, p38) |
| 103 | 101 | BIBW-2992, Tovok, Afatinib, Gilotrif, Giotrif, Tomtovok | 0.66193 | -0.71643 | 0.47400 | -1.47748 | -1.08835 | -0.42368 | 0.71668 | -0.37157 | -0.14112 | EGFR (ERBB1, HER1) |
| 104 | 102 | Purvalanol A, NG 60, N-60 | 0.28745 | -2.12242 | -0.29205 | -0.37922 | -1.86931 | -0.14352 | 0.38591 | -3.95461 | -0.51300 | CDC2 (CDK1, p34cdc2, CDC28p)/Cyclin B, CDK2/Cyclin A, CDK2/Cyclin E, CDK4/Cyclin D1, CDK5/p35, DYRK1A (DYRK) |
| 105 | 103 | Dasatinib, BMS-354825, Sprycel, NSC-732517 | -1.46662 | -0.83403 | 0.23633 | -0.31428 | -3.54117 | -1.33258 | -2.27221 | -3.13330 | -0.80287 | BCR, ABL1 (ABL), SRC (c-SRC), EphA1 (EPH), FYN (p59fyn), LCK (p56lck), PDGFRb (PDGFR, PDGFR1), YES, Erk2 (ERK, p38) |
| 106 | 104 | CI-1033, Canertinib, PD-183805, SN-26606 | -1.05629 | -0.87657 | -0.68222 | -0.56100 | -0.95903 | 0.53254 | 1.36032 | 0.98196 | -0.07994 | EGFR (ERBB1, HER1), ErbB2 (TKR1, HER2, NEU) - Irreversible |
| 107 | 105 | SC-514 | -0.16870 | -2.04388 | -1.45928 | -0.66433 | -0.14577 | -1.68982 | -0.23261 | -1.21883 | -0.48353 | IKKb (IKK2) |
| 108 | 106 | CP-690550, CP-690,550, Tasocitinib, Tofacitinib, Jaquinus, Xeljanz | -0.97221 | -0.17494 | -1.11862 | -1.40392 | -1.59356 | 2.59826 | 0.54694 | 1.18159 | -0.05940 | JAK3 |
| 109 | 107 | Bosutinib, SKI-606, Bosulif | 0.60697 | -0.52897 | -0.02361 | 0.93055 | -0.88882 | 0.10174 | -3.14588 | -7.19257 | -0.64321 | BCR, ABL1 (ABL), SRC (c-SRC), FGR (SRC2), LYN |
| 110 | 108 | Pimecrolimus, Elidel, SDZ-ASM-981 | -0.86441 | -0.09352 | 1.69227 | 0.52388 | -1.11771 | -0.18686 | 0.56967 | -2.40454 | -0.11932 | Calcineurin (CN, PP-2B) |
| 111 | 109 | Rottlerin | 0.26035 | 0.97634 | 0.68715 | -0.62214 | -2.12145 | -1.80530 | -2.17270 | -3.15627 | -0.50451 | PKCd, TGM2 (Tissue Transglutaminase), signal transduction pathway inhibitor |
| 112 | 110 | Temsirolimus, CCI-779, Torisel | -1.65759 | -1.57061 | -2.55868 | -0.25027 | -3.02219 | -1.32368 | -2.28379 | -2.50102 | -0.96208 | FRAP (MTOR, FRAP1) |
| 113 | 111 | Ro-31-8220 mesylate, Bisindolylmaleimide IX, Ro-318220 | 0.99179 | 0.52035 | 0.66407 | 1.01935 | -1.08982 | 5.24404 | 11.69089 | 9.18229 | 1.79015 | PKC |
| 114 | 112 | 7-Cyclopentyl-5-(4-phenoxyphenyl)-7H-pyrrolo[2,3-d]pyrimidin-4-ylamine | -1.78344 | -0.90809 | 0.22302 | 1.19297 | -1.49333 | -0.52636 | 0.53236 | 1.22833 | -0.09733 | LCK (p56lck), SRC (c-SRC) |
| 115 | 113 | D4476, D-4476 | 0.77801 | -0.22058 | -1.19367 | 1.27680 | -1.43362 | 1.30205 | -0.62521 | 0.07478 | -0.00263 | CK1a (Casein kinase 1, CK1) |
| 116 | 114 | SU-11274, PKI-SU11274, SU-MI-2 | 0.16471 | -0.32063 | -0.76218 | 0.18627 | -0.62139 | -1.80658 | -2.05431 | -2.49685 | -0.48910 | MET (c-MET, HGFR) |
| 117 | 115 | Masitinib mesylate, AB1010, AB-1010, Masiviera | -1.80428 | -0.07120 | -0.05621 | -0.21475 | -0.02609 | -0.95983 | -0.72925 | -0.65920 | -0.28675 | KIT (c-KIT), PDGFRb (PDGFR, PDGFR1), FGFR3 |
| 118 | 116 | H 89 dihydrochloride, H-89 dihydrochloride | -1.30448 | 0.21439 | 0.73830 | 0.31013 | 1.79413 | -1.00519 | 0.36616 | -0.25723 | 0.05431 | PKACa (PKA) |
| 119 | 117 | ER 27319 maleate, ER-27319 | -0.20906 | 0.46538 | 0.01635 | -0.91390 | -4.76132 | -4.70724 | -9.40797 | -10.13961 | -1.88113 | SYK, Mediator Release Inhibitors |
| 120 | 118 | CID 755673 | -0.49857 | -0.68661 | -0.70981 | 0.19002 | -0.48954 | -0.31954 | 2.13275 | 1.12114 | 0.04693 | PKD1 (PKD) |
| 121 | 119 | PF-670462 | 0.17291 | 3.10910 | -0.43869 | 0.82768 | -1.68736 | -0.39133 | 0.06254 | -0.69342 | 0.06098 | CK1e, CK1d |
| 122 | 120 | SKF 86002 dihydrochloride, SKF-86002 dihydrochloride, SK&F-86002 | 0.03537 | 0.09345 | -0.50189 | -0.88563 | 0.06085 | -0.32628 | 2.02410 | -0.83506 | -0.02125 | p38 MAPK |
| 123 | 121 | SL327, SL-327 | -0.96649 | -0.71310 | -1.55905 | -1.12806 | -0.88777 | -0.82073 | 0.24476 | 0.14353 | -0.36071 | MAP2K1 (MEK1), MAP2K2 (MEK2) |
| 124 | 122 | SP-600125 | 1.46626 | 0.52167 | -0.54883 | -0.31250 | -2.63791 | -1.17016 | -0.56745 | -1.96421 | -0.33066 | JNK1 (JNK) |
| 125 | 123 | Sphingosine kinase Inhibitor 2, D-erythro-Sphingosine | 0.60420 | -0.54682 | 0.04683 | -0.04875 | -0.12500 | -1.62449 | -1.91481 | -0.51580 | -0.26162 | SPK1 (SPHK, SK1, Sphingosine Kinase) |
| 126 | 124 | SU 4312, DMBI, SU-4312 | -0.39984 | -1.08495 | -1.42095 | 0.43777 | -1.15288 | -2.00473 | -1.79205 | -1.57828 | -0.57060 | KDR (VEGFR2, VEGFR, FLK1) |
| 127 | 125 | SU 5416, SU-5416, TSU-16, NSC-696819, Semaxanib, Semaxnib, Semoxind | -2.51932 | -3.34659 | -5.37630 | -6.46704 | -8.57874 | -9.10871 | -8.79347 | -10.59191 | -3.47476 | FLT3 (STK1, FLK2), MET (c-MET, HGFR), KDR (VEGFR2, VEGFR, FLK1) |
| 128 | 126 | PF-573228 | 0.53454 | 1.60395 | 0.28795 | 1.04918 | 1.15147 | 0.37973 | -0.35206 | 0.92336 | 0.35381 | FAK (FAK1) |
| 129 | 127 | TPCA-1 | -0.87107 | 0.32796 | -0.53894 | -0.62160 | -0.78310 | -1.64841 | -0.79677 | 2.26052 | -0.16944 | IKKb (IKK2) |
| 130 | 128 | Quercetin, LIM-5662, LNS-5662, Quer | -0.28935 | -0.14497 | 0.51184 | 0.02305 | 0.81790 | -0.38750 | 0.90934 | -1.37566 | 0.00410 | PI3K |
| 131 | 129 | FAK Inhibitor 14, CFAK-Y15, Y-15 | -0.48902 | -0.75065 | -1.03144 | -0.37772 | -1.10260 | 1.06809 | -0.68478 | 3.58673 | 0.01387 | FAK (FAK1) |
| 132 | 130 | ZM 336372, ZM-336372 | -1.17942 | -1.30562 | -0.82959 | 1.90989 | -1.64618 | -1.77246 | -1.81565 | -1.23993 | -0.49975 | RAF1 (c-Raf) |
| 133 | 131 | XRP44X, XRP-44X, WCH-44 | -0.74253 | 1.99193 | -1.89070 | 0.68438 | -1.33301 | -1.07257 | -0.40021 | -9.69488 | -0.79017 | Ras-NET (Elk3), Tubulin polimerization |
| 134 | 132 | Necrostatin-1, MTH-DL-Tryptophan, MTH-trp, Nec-1 | -1.40758 | -0.00691 | -1.11616 | -0.83960 | -2.33488 | -3.95024 | -4.89366 | -5.83005 | -1.29262 | RIPK1 (RIP) |
| 135 | 133 | PD-407824 | -0.99345 | 0.05402 | 0.16575 | 0.21055 | -1.15050 | -3.18357 | -2.96481 | -0.68080 | -0.54186 | CHK1, WEE1 |
| 136 | 134 | PHA-767491 hydrochloride | 0.84806 | 0.03604 | -1.24255 | -0.81872 | -2.69207 | -5.22465 | -1.46794 | -8.40480 | -1.20303 | CDC7, CDK9/Cyclin T1, MAPKAPK2 (MK2) |
| 137 | 135 | TCS PIM-1 1 | -0.69185 | -0.46832 | -1.05168 | -0.02402 | -0.25366 | 1.16783 | 0.77991 | -0.10532 | -0.04105 | PIM1, PIM2 |
| 138 | 136 | 2-(p-Hydroxyanilino)-4-(p-chlorophenyl) thiazole, SKI-II, ABC-294640 | -0.20952 | -0.90415 | -0.70515 | 0.06801 | -2.30186 | -3.63199 | -2.59992 | -2.53941 | -0.81341 | SPK1 (SPHK, SK1, Sphingosine Kinase) |
| 139 | 137 | 10-DEBC hydrochloride, 10-NCP | -0.16339 | -0.66325 | -0.04979 | 0.32637 | -0.58039 | 0.70507 | 0.06404 | 3.04054 | 0.16994 | AKT1 (PKBa) |
| 140 | 138 | FPA 124, FPA-124 | -0.97248 | -0.75049 | -0.50244 | -0.33594 | -2.38164 | -0.14071 | -0.22861 | -1.74145 | -0.44741 | AKT1 (PKBa) |
| 141 | 139 | API-2, Triciribine, NSC154020, TCN, Tricibine, VQD-002 | -0.37694 | 0.75315 | -0.28369 | 0.28926 | -1.06983 | -2.71083 | -0.42969 | -5.15596 | -0.56988 | AKT1 (PKBa) |
| 142 | 140 | SGI-1776 | -0.10679 | 0.48627 | 0.37797 | -0.35956 | -0.19542 | -0.28997 | 1.56960 | 2.45909 | 0.24998 | PIM1, PIM2, FLT3 (STK1, FLK2) |
| 143 | 141 | A-443654, A-654 | -1.54599 | -1.20560 | 0.24927 | -0.91007 | -3.36734 | -5.65293 | -0.88275 | -6.97926 | -1.28726 | AKT1 (PKBa) |
| 144 | 142 | CGP 57380, CGP-57380 | -1.90638 | -0.85692 | -0.71640 | -3.52198 | -2.99162 | -2.09472 | -4.53162 | -3.74229 | -1.29153 | MNK1 |
| 145 | 143 | SU-6668, TSU-68, Orantinib | -1.09698 | -0.06505 | -0.18696 | 0.11182 | 0.79364 | -1.85619 | -0.57623 | -1.56130 | -0.28145 | PDGFRb (PDGFR, PDGFR1), KDR (VEGFR2, VEGFR, FLK1), FGFR1 (FLT2), AurA (AuroraA), AurB (AuroraB), TBK1 |
| 146 | 144 | TAE-684, NVP-TAE684 | -0.27612 | -1.60650 | 0.12603 | 0.42251 | 1.39781 | 5.32403 | 7.03266 | -3.98677 | 0.53494 | ALK |
| 147 | 145 | AZD0530, Saracatinib, NSC-735464, AZD-0530 | -2.23521 | -0.52548 | 0.43609 | -0.54344 | -1.27747 | -0.19184 | 0.79408 | -1.68199 | -0.33143 | ABL1 (ABL), SRC (c-SRC), KIT (c-KIT) |
| 148 | 146 | Sorafenib, Bay 43-9006, Nexavar | 0.02528 | -0.26545 | -1.11260 | -0.47299 | -2.74898 | -1.74372 | -2.68553 | -4.23313 | -0.83961 | FLT3 (STK1, FLK2), KIT (c-KIT), PDGFRb (PDGFR, PDGFR1), RET, RAF1 (c-Raf), BRAF (B-raf), FLT1 (VEGFR1), KDR (VEGFR2, VEGFR, FLK1), FLT4 (VEGFR3) |

| | A | B | C | D | E | F | G | H | I | J | K | L |
|---|---|---|---|---|---|---|---|---|---|---|---|---|
| 149 | 147 | L-779450, Raf Kinase Inhibitor IV | -1.11398 | 0.58317 | -1.38116 | -0.19161 | -1.79987 | -0.96476 | -3.86260 | -5.34843 | -0.89303 | RAF1 (c-Raf) |
| 150 | 148 | PLX-4720, Raf Kinase Inhibitor V | -0.88817 | -0.00759 | -0.63156 | 1.12391 | -1.37335 | 0.29190 | -0.84691 | -2.31892 | -0.29499 | BRAF (B-raf) |
| 151 | 149 | BI 2536, BI-2536 | -1.97684 | 0.69251 | 2.00678 | -6.74103 | -8.54315 | -5.50120 | -6.88131 | -6.73975 | -2.13653 | PLK1 (PLK, STPK13), Bromodomain-Containing Protein 4 (Brd4, HUNK1) Inhibitor |
| 152 | 150 | Mubritinib, TAK-165, D04025 | -2.11862 | 0.29094 | -0.04467 | -0.21088 | -1.55176 | -0.60633 | -1.39967 | -2.34153 | -0.50632 | ErbB2 (TKR1, HER2, NEU) |
| 153 | 151 | BMS-599626, AC-480 | -0.75870 | -0.12397 | -0.37711 | 0.77378 | 0.31541 | -1.16656 | -0.07474 | -1.95553 | -0.21359 | EGFR (ERBB1, HER1), ErbB2 (TKR1, HER2, NEU), ErbB3 (HER3), ErbB4 (HER4) |
| 154 | 152 | Motesanib diphosphate salt, AMG-706 | 0.07991 | -0.72779 | -0.86448 | 0.01721 | 0.44755 | -1.08084 | -2.21987 | -6.33538 | -0.67765 | KIT (c-KIT), PDGFRb (PDGFR, PDGFR1), KDR (VEGFR2, VEGFR, FLK1) |
| 155 | 153 | ZSTK474 | -0.59788 | -0.02571 | 1.70390 | 1.01383 | 2.19794 | 3.76510 | 4.18084 | 6.89504 | 1.21359 | PI3K |
| 156 | 154 | Bisindolylmaleimide I hydrochloride, GF 109203X HCl, Go 6850 HCl | -1.57708 | -1.42740 | 0.55120 | -0.89120 | -1.32842 | 0.54362 | -2.74796 | -6.70108 | -0.86126 | PKC |
| 157 | 155 | VX-702, VX-850, VX-954, KVK-702 | 0.49078 | -0.04966 | 0.23226 | 1.50155 | -1.09574 | -0.23067 | 0.66158 | -0.75656 | 0.04780 | p38 MAPK |
| 158 | 156 | ZM-447439 | 1.35667 | 1.15174 | -0.90957 | 0.90194 | 0.11473 | 0.31252 | -9.29908 | -9.19405 | -0.98727 | AurA (AuroraA), AurB (AuroraB), AurC (AuroraC) |
| 159 | 157 | Tyrphostin AG 1478 mesylate, AG-1478 | -1.09317 | -0.66600 | -1.46281 | 0.86625 | -1.58238 | -1.37287 | -2.00072 | -1.77225 | -0.57618 | EGFR (ERBB1, HER1) |
| 160 | 158 | Tyrphostin SU 1498, SU 1498, SU-1498 | -0.35709 | 0.39627 | -1.33761 | -0.27653 | -0.55235 | 0.29887 | -1.66516 | -1.86715 | -0.34002 | KDR (VEGFR2, VEGFR, FLK1) |
| 161 | 159 | U0126, U-0126, U 0126 | -2.09068 | -0.39591 | -0.08259 | 0.34058 | -1.93059 | -2.82072 | -2.81828 | -6.84107 | -1.05541 | MAP2K1 (MEK1), MAP2K2 (MEK2), JAK2, Erk2 (ERK, p38) |
| 162 | 160 | Chelerythrine chloride, MP-0922, NSC-646662, Toddalin | -1.71077 | -0.38902 | -0.69385 | -0.87149 | -1.26132 | -0.32920 | 2.25462 | -1.22246 | -0.26789 | PKC |
| 163 | 161 | Imatinib (free base), Gleevec, Glivec, CGP-57148B, STI-571, NSC-716051, QTI-571, Ruvise | -1.23923 | 1.30791 | 0.33190 | 0.63955 | -1.37187 | 0.56721 | -1.96801 | -1.29315 | -0.19192 | BCR, ABL1 (ABL), PDGFRb (PDGFR, PDGFR1), KIT (c-KIT) |
| 164 | 162 | RDEA-119, AR-119, RDEA119, BAY-869766, BAY-86-9766, Refametinib | 0.00694 | 0.49277 | -0.26006 | 2.52303 | 1.87292 | 3.05487 | 3.26352 | -3.58560 | 0.46737 | MAP2K1 (MEK1), MAP2K2 (MEK2) |
| 165 | 163 | SB-202190, FHPI | -0.09038 | -0.53350 | -0.63418 | 0.31597 | 0.64188 | -0.07255 | -0.13753 | -0.68574 | -0.07586 | p38 MAPK |
| 166 | 164 | Alsterpaullone, NSC-705701, 9-Nitropaullone | -1.13317 | 0.42687 | -0.98109 | 1.17003 | 0.06864 | -1.69685 | -1.74484 | -4.46229 | -0.52980 | CDC2 (CDK1, p34cdc2, CDC28p)/Cyclin B, CDK2/Cyclin A, CDK2/Cyclin E, CDK5/p35, GSK3 |
| 167 | 165 | NSC-663284, DA-3003-1 | 0.03409 | -1.22598 | -0.55345 | -0.10147 | -0.57808 | 0.33604 | 0.21935 | 0.66335 | -0.07650 | cdc25 |
| 168 | 166 | NU-2058 | -1.71784 | -0.68303 | 0.85629 | 0.48440 | -0.93855 | 0.96062 | -1.90349 | -1.74233 | -0.29710 | CDC2 (CDK1, p34cdc2, CDC28p)/Cyclin B, CDK2/Cyclin A, CDK2/Cyclin E, CDK5/p35, GSK3 |
| 169 | 167 | PD-0325901, PD 0325901 | -0.58463 | 0.94617 | 2.60387 | 2.36335 | 0.84314 | 2.25953 | 1.56718 | -2.05074 | 0.50412 | MAP2K1 (MEK1), MAP2K2 (MEK2) |
| 170 | 168 | PD-184161 | -0.29574 | 0.25325 | -0.38812 | 0.89437 | -0.11184 | 1.13785 | 1.19303 | 1.50381 | 0.26555 | MAP2K1 (MEK1), MAP2K2 (MEK2) |
| 171 | 169 | EKI-785, CL-387785 | 0.17998 | 0.09741 | -0.35997 | -0.40120 | -0.92892 | 0.66026 | 0.92180 | 2.00593 | 0.13798 | EGFR (ERBB1, HER1) - Irreversible |
| 172 | 170 | Olomoucine | -2.63203 | 0.46824 | 0.30165 | 0.16816 | -0.38429 | 0.84166 | -1.66781 | -13.60284 | -1.04703 | CDC2 (CDK1, p34cdc2, CDC28p)/Cyclin B, CDK2/Cyclin A, CDK2/Cyclin E, CDK5/p35 |
| 173 | 171 | BMS-536924 | -0.89730 | 0.28812 | -0.87565 | -0.17320 | -2.35992 | -2.90825 | -1.87802 | 2.56680 | -0.39563 | IGF1R (JTK13, IGFIR) |
| 174 | 172 | NVP-AEW-541, AEW-541, NVP-AEW541-NX, NVP-AEW541-NX7 | -0.51233 | -0.46463 | 1.31098 | -0.20728 | -0.14355 | 1.11185 | 4.90991 | 4.13182 | 0.64296 | IGF1R (JTK13, IGFIR) |
| 175 | 173 | Sunitinib Malate, Sutent, SU-11248, PHA-290940AD, PNU-290940AD | -0.08039 | 0.72984 | -1.30770 | 0.31692 | -0.75924 | -0.50427 | -1.01560 | -4.95858 | -0.48073 | KIT (c-KIT), FLT3 (STK1, FLK2), FMS (c-FMS, CSF-1R, CSF1R), KDR (VEGFR2, VEGFR, FLK1), PDGFRb (PDGFR, PDGFR1) |
| 176 | 174 | Y-27632 | 2.41380 | 1.13806 | -0.40658 | 0.68306 | -0.24682 | -0.21280 | 0.29103 | -0.84110 | 0.17878 | LRRK2, ROCK1, ROCK2 (ROCKa), Calcium sensitizer |
| 177 | 175 | AS-252424 | 0.81476 | 0.02241 | 0.52819 | -0.82261 | -0.32507 | -1.08184 | -0.25378 | -0.84022 | -0.12420 | PI3K |
| 178 | 176 | Ryuvidine | -0.00321 | 1.00095 | -1.01066 | 0.36371 | -0.45677 | -0.36649 | 4.54289 | 20.70703 | 1.57160 | CDK4/Cyclin D1, SETD8 inhibitor |
| 179 | 177 | TGX-221 | -0.48420 | -0.39125 | -1.19256 | 2.10747 | 2.64076 | 0.27887 | 0.52786 | 0.69392 | 0.26519 | PI3K |
| 180 | 178 | Dovitinib, CHIR-258, TKI-258, GFKI-258 | 0.15713 | 0.08329 | -0.32910 | 0.45801 | -2.24505 | 1.58605 | -2.65563 | -9.38032 | -0.78180 | KDR (VEGFR2, VEGFR, FLK1), FLT3 (STK1, FLK2), KIT (c-KIT), PDGFRb (PDGFR, PDGFR1) |
| 181 | 179 | TX-1918 | -1.32187 | 0.18804 | 0.75586 | 0.33932 | -0.91454 | 0.24270 | 0.33402 | 0.40679 | 0.00192 | eEF2K |
| 182 | 180 | OSU-03012 hydrochloride, AR-12, NSC-728209 | -0.74251 | 1.59327 | -0.50281 | 0.04326 | -0.73095 | -0.39929 | 0.43680 | -0.58907 | -0.05653 | PDK1 (PDPK1) |
| 183 | 181 | K-252c, Staurosporine Aglycone | -0.03238 | -0.85819 | -1.27547 | -1.95194 | -3.38709 | -3.25887 | -0.67580 | -2.63068 | -0.89247 | PKC |
| 184 | 182 | IC87114, D-030, IC-87114, GS-548202 | -0.80601 | 0.35046 | 0.78271 | 1.41988 | 0.55097 | 1.94978 | 0.21639 | -0.79499 | 0.23273 | PI3Kd |
| 185 | 183 | Akt 1/2 Kinase inhibitor, 352556 | -1.11006 | 0.39859 | 1.58528 | -0.44946 | 0.77638 | 0.37378 | -0.23232 | 0.26927 | 0.10221 | AKT1 (PKBa), AKT2 (PKBb) |
| 186 | 184 | Imidazolo-oxindole PKR inhibitor C16, PKRi | -0.80282 | -0.15406 | 1.23577 | 0.88449 | -0.43176 | -4.15879 | -5.61795 | -6.76469 | -1.00280 | PKR (EIF2AK1, EIF2AK2) |
| 187 | 185 | SU-9516 | -1.17350 | 0.72480 | -0.89980 | 1.87640 | -0.80061 | -0.33519 | -0.63807 | -2.26308 | -0.22257 | CDC2 (CDK1, p34cdc2, CDC28p)/Cyclin B, CDK2/Cyclin A, CDK4/Cyclin D1 |
| 188 | 186 | Compound 401 | -1.23028 | -0.50026 | -1.39452 | -0.61832 | -1.78508 | -0.56050 | -0.75682 | -2.14819 | -0.57048 | DNAPK (DNA-PKcs), FRAP (MTOR, FRAP1) |
| 189 | 187 | NSC 109555 ditosylate, DDUG, MES-10950 | -1.93671 | -0.47227 | 0.76682 | 0.80911 | -1.36883 | -2.19785 | -1.05109 | -0.21181 | -0.35917 | CHK2 |
| 190 | 188 | PHA-665752 | -0.06431 | -0.31623 | -0.00052 | 0.74050 | -0.75023 | 0.13933 | 0.16946 | 0.37617 | 0.01866 | MET (c-MET, HGFR), RON (MST1R), KDR (VEGFR2, VEGFR, FLK1) |
| 191 | 189 | Dioctanoylglycol | -0.28913 | -0.71115 | 0.80362 | -0.15654 | -0.74746 | -0.25192 | 1.03177 | -0.77549 | -0.06954 | DAGK (DGK, Diacylglycerol kinase) |
| 192 | 190 | R 59-022, R-59022 | -0.12189 | -0.43575 | 0.22456 | 1.27768 | 0.42991 | 0.19874 | 0.76800 | 0.65190 | 0.18985 | DAGK (DGK, Diacylglycerol kinase) |
| 193 | 191 | AG 490, Tyrphostin AG 490, Tyrphostin B42 | -0.55688 | -0.81012 | 0.94730 | 0.80913 | 0.29965 | 0.10124 | -1.86848 | 0.00853 | -0.06784 | EGFR (ERBB1, HER1), JAK2, JAK3/STAT, JAK3/AP-1, JAK3/MAPK |
| 194 | 192 | PD-158780 | 0.68703 | 1.16475 | -1.46857 | 0.93270 | -0.87714 | -0.77771 | -2.92687 | -0.80238 | -0.25804 | EGFR (ERBB1, HER1), ErbB2 (TKR1, HER2, NEU), ErbB4 (HER4) |
| 195 | 193 | AG 18, RG-50810, TyrphostinA23 | -2.78608 | -0.25491 | 0.94225 | 1.55159 | -1.44092 | -0.36332 | -0.84277 | -0.33987 | -0.22416 | EGFR (ERBB1, HER1), PDGFRb (PDGFR, PDGFR1) |
| 196 | 194 | AG 213, Tyrphostin AG 213 | -0.38209 | 0.09060 | -1.08314 | 1.36070 | 1.29492 | -0.05760 | -0.11550 | -1.89319 | -0.04981 | EGFR (ERBB1, HER1), PDGFRb (PDGFR, PDGFR1) |
| 197 | 195 | AG 494 | 1.26317 | 0.35553 | -0.43918 | 1.25377 | -0.00890 | -0.64592 | -0.09956 | -1.77711 | -0.00623 | EGFR (ERBB1, HER1) |
| 198 | 196 | AG 825, Tyrphostin AG 825 | -0.15715 | 0.52458 | 0.40067 | 0.00866 | -0.42443 | -0.48775 | 0.11323 | 0.51350 | 0.03116 | ErbB2 (TKR1, HER2, NEU) |
| 199 | 197 | Genistein, BIO-300, G-2535, PTI-G4660, SIPI-9764-I, Bonistein, 4',5,7-Trihydroxyisoflavone | -0.17312 | -0.64403 | 1.41987 | 1.16187 | -0.54016 | 0.37035 | 1.46112 | -0.93517 | 0.13452 | EGFR (ERBB1, HER1), CFTR Channel Activator, DNA Topoisomerase II Inhibitor, NADH Oxidase (NOX) Inhibitor, Tyrosine Kinase Inhibitor |
| 200 | 198 | HDS 029 | 0.06810 | 1.52026 | 0.46421 | 0.09785 | -1.58445 | -0.78019 | -1.60914 | -3.13537 | -0.31453 | EGFR (ERBB1, HER1), ErbB2 (TKR1, HER2, NEU), ErbB4 (HER4) |
| 201 | 199 | JNJ 28871063 hydrochloride | -1.54372 | 0.41063 | -0.29081 | 1.29306 | -0.83623 | -0.62675 | -0.50283 | -0.36348 | -0.15604 | EGFR (ERBB1, HER1), ErbB2 (TKR1, HER2, NEU), ErbB4 (HER4) |
| 202 | 200 | Lavendustin A, RG 14355 | 0.10413 | 1.92727 | 1.20405 | -0.47279 | -0.23695 | -1.25099 | -1.07250 | -1.02928 | -0.05246 | EGFR (ERBB1, HER1), SRC (c-SRC) |
| 203 | 201 | PP-3 | 0.58420 | 0.07552 | 0.02742 | -0.27156 | -0.68232 | -0.27808 | -0.57020 | -3.79469 | -0.31142 | EGFR (ERBB1, HER1) |
| 204 | 202 | Tyrphostin B44, (-) enantiomer | -0.00492 | 0.13020 | -0.62613 | 1.21633 | -0.94076 | -1.85777 | -0.78059 | -0.46572 | -0.21118 | EGFR (ERBB1, HER1), ErbB2 (TKR1, HER2, NEU) |
| 205 | 203 | SB-218078 | -0.92222 | 0.83797 | -0.13788 | 3.00148 | -2.97002 | -8.28851 | -5.82658 | -3.79753 | -1.14827 | CHK1, CDC2 (CDK1, p34cdc2, CDC28p)/Cyclin B, PKC |
| 206 | 204 | 1-Naphthyl PP1, 1-NA-PP 1, 1-NaPP1 | -0.83403 | 0.00917 | 1.42078 | -0.18496 | 1.18584 | 0.36067 | 1.62387 | 2.17453 | 0.36509 | SRC (c-SRC), FYN (p59fyn), ABL1 (ABL) |
| 207 | 205 | ABT-702 dihydrochloride | -0.16267 | 0.09303 | -1.37025 | 1.11305 | -0.11408 | 0.26559 | 0.62096 | -0.96206 | -0.03276 | AK (Adenosine Kinase) |
| 208 | 206 | 5-Iodotubercidin, IT | -0.09426 | 0.48033 | -0.48050 | -0.82763 | -1.40953 | -2.70874 | -1.03197 | -6.33265 | -0.78683 | AK (Adenosine Kinase), CK1a (Casein kinase 1, CK1), Decrease DNA methylation |
| 209 | 207 | A 83-01, A-83-01 | -0.07398 | 0.54801 | 1.78188 | 2.82471 | 2.05161 | 2.11762 | 3.57128 | 1.61274 | 0.91552 | TGFbR1 (ALK5) |
| 210 | 208 | Dorsomorphin dihydrochloride, BML-275, AMPK Inhibitor, Compound C | -0.37119 | 0.31524 | 0.74849 | -0.79398 | -1.42353 | -0.97167 | -1.19170 | -2.12551 | -0.36877 | AMPK, ALK2, BMPR1A (ALK3), BMPR1B (ALK6) |
| 211 | 209 | CGK 733, CGK-733 | -0.05337 | -1.04408 | -1.13919 | 0.06359 | -1.10876 | 0.03270 | -0.60430 | -0.22852 | -0.25891 | ATR, ATM |
| 212 | 210 | LFM-A13, DDE-28, LFM-A13-F (oral formulation) | -0.41807 | 0.22715 | 0.59244 | 0.52277 | 0.01658 | -0.77346 | -1.31181 | -0.96898 | -0.13405 | BTK (Bruton's tyrosine kinase) |
| 213 | 211 | TCS JNK 5a | -0.01012 | -0.58435 | -1.07645 | 0.64035 | -1.47547 | -1.90725 | -2.74373 | -9.37343 | -1.04851 | JNK1 (JNK), JNK2, JNK3, p38a |
| 214 | 212 | BI 78D3, BI-78D3 | -1.36130 | -0.43526 | 1.29098 | 0.11216 | -1.17664 | -0.08075 | -0.70888 | 0.12886 | -0.14150 | JNK1 (JNK) |
| 215 | 213 | Arcyriaflavin A | -0.68975 | 0.04077 | -0.72148 | -0.53774 | -5.11699 | -4.33063 | -6.81146 | -5.84815 | -1.52327 | CDK4/Cyclin D1, CaMK2b (CaMK2) |
| 216 | 214 | ML 9 hydrochloride, ML-9 | 0.63850 | -0.43459 | 1.86866 | 0.24814 | 1.48159 | 1.65375 | 2.57644 | 4.61783 | 0.80239 | smMLCK (MLCK, myosin light chain kinase, MYLK) |
| 217 | 215 | Ellagic acid, LTK-20 | 0.48689 | -0.65069 | -1.56112 | -0.58231 | -1.61163 | -0.67246 | -0.13513 | -1.07906 | -0.36824 | CK2a1 (CK2, CKII, Casein kinase2) |
| 218 | 216 | TBB, NSC 231634, Tetrabromobenzotriazole | -0.80588 | -1.16556 | 0.31688 | 1.96000 | -0.71530 | 0.79629 | -0.11617 | -0.56925 | -0.01896 | CK2a1 (CK2, CKII, Casein kinase2) |
| 219 | 217 | GSK-650394 | -0.66491 | 0.15809 | -0.40373 | 0.12823 | -1.01365 | -0.79445 | -1.05425 | -0.56075 | -0.26675 | SGK1 (SGK), SGK2 |
| 220 | 218 | DCA, Sodium Dichloroacetate, CPC-211, X-11S | -0.62483 | 2.03060 | 0.00097 | -0.34304 | -0.97016 | -0.46611 | 1.01225 | -1.54975 | -0.05772 | PDK1 (PDPK1) |
| 221 | 219 | IPA 3, IPA-3 | -1.52682 | -0.54176 | -0.27810 | -0.24269 | -1.90387 | -0.36890 | 0.38242 | -1.23936 | -0.36275 | PAK1 (PAKa) |
| 222 | 220 | (5Z)-7-Oxozeaenol, C-292, FR-148083, L-783278, LL-Z1640-2, f-152A1 | 0.25211 | 0.78081 | -0.06244 | -0.76406 | -1.15882 | 2.75387 | -1.11518 | -6.14657 | -0.34634 | TAK1 (TGF1a, MAP3K7) |
| 223 | 221 | Purvalanol B, NG 95, NG-95 | -0.95539 | 0.64142 | -0.14424 | 1.13971 | -0.37949 | -0.46539 | 0.27987 | -0.50420 | -0.02459 | CDC2 (CDK1, p34cdc2, CDC28p)/Cyclin B, CDK2/Cyclin A, CDK2/Cyclin E, CDK5/p35 |
| 224 | 222 | WHI-P 154, MASTPROM, WHI-P154 | -0.33389 | -0.54509 | -0.13296 | -0.17215 | -0.17376 | -1.40330 | 0.32389 | 0.59278 | -0.11699 | JAK3, EGFR (ERBB1, HER1), SRC (c-SRC), ABL1 (ABL), KDR (VEGFR2, VEGFR, FLK1), p38 MAPK, PI3K |
| 225 | 223 | SU-5402 | 0.72382 | 0.37045 | 0.59047 | 0.57589 | -1.61746 | 0.06664 | -2.30222 | -5.65413 | -0.45964 | KDR (VEGFR2, VEGFR, FLK1), FGFR1 (FLT2), PDGFRb (PDGFR, PDGFR1) |
| 226 | 224 | Ki-8751 | -2.04791 | -0.05494 | 0.66119 | -0.38585 | -1.22668 | -1.17027 | 1.22232 | -4.57029 | -0.48031 | KDR (VEGFR2, VEGFR, FLK1), KIT (c-KIT), PDGFRa (PDGFR2), FGFR2, FLT3 (STK1, FLK2) |
| 227 | 225 | ZM 306416 hydrochloride, ZM-306416 | 1.43439 | -0.52746 | 0.70980 | -0.34657 | -2.32742 | -0.06554 | -1.40884 | -2.44965 | -0.31596 | KDR (VEGFR2, VEGFR, FLK1) |
| 228 | 226 | ZM 323881 hydrochloride, ZM-323881 | -0.74841 | -1.39662 | -0.59164 | -1.36986 | -2.14440 | -1.27059 | -3.60924 | -3.12400 | -0.90416 | KDR (VEGFR2, VEGFR, FLK1) |
| 229 | 227 | NSC 693868, NSC-693868 | -1.32271 | 0.49861 | -1.04760 | 1.18276 | 0.58961 | 0.44513 | 1.53908 | -3.35825 | -0.09345 | CDC2 (CDK1, p34cdc2, CDC28p)/Cyclin B, CDK5/p25, GSK3 |
| 230 | 228 | Arctigenin, (-)-Arctigenin | 0.10474 | 0.72546 | 0.70582 | 1.00035 | 0.25816 | 0.42697 | -1.17930 | -3.00883 | -0.06131 | MAP2K1 (MEK1), DNA Topoisomerase II, HIV Integrase, NFKB, AP-1 |
| 231 | 229 | IKK 16, IKK-16 | 0.23824 | -0.94153 | -0.28194 | 2.11797 | -0.14523 | 0.46817 | 3.54788 | -0.45519 | 0.28850 | IKKa (IKK1), IKKb (IKK2) |
| 232 | 230 | IMD 0354, IMD-0354 | 0.55301 | 0.23570 | -0.03951 | -0.08225 | 0.58467 | -0.69855 | -0.24300 | -1.47529 | -0.07391 | IKKb (IKK2) |
| 233 | 231 | JX-401 | -1.34958 | 1.11549 | 0.14432 | -0.46172 | -0.56778 | -0.30093 | 1.98614 | -0.35762 | 0.01321 | p38a |
| 234 | 232 | EO-1428 | -0.68257 | -0.04293 | -1.02042 | 0.82361 | -0.99648 | 1.54299 | 0.36671 | -1.11528 | -0.07132 | p38a, p38b |
| 235 | 233 | RWJ-67657 | -1.11095 | 0.03879 | 0.89227 | -0.37624 | -1.30602 | 0.58795 | -1.12123 | -0.78033 | -0.20143 | p38a, p38b |
| 236 | 234 | PD-198306 | 0.71619 | 0.16229 | 0.00707 | -0.13786 | -0.27024 | 1.69176 | 0.21981 | 1.80461 | 0.26600 | MAP2K1 (MEK1), MAP2K2 (MEK2) |
| 237 | 235 | TCS 359, TCS-359 | -0.42287 | 0.01391 | -0.55984 | 0.70082 | -0.04888 | -0.35635 | -1.93827 | -0.98249 | -0.22796 | FLT3 (STK1, FLK2) |
| 238 | 236 | TCS 2312 dihydrochloride, TCS-2312 | -2.18113 | 0.74168 | -0.62696 | 0.39150 | -0.46444 | 4.30057 | 4.37234 | -6.54909 | -0.00099 | CHK1 |
| 239 | 237 | H 1152, Glycyl dihydrochloride, H-1152, Dimethylfasudil, AL-38565A, H-1152P, HMN-1152, diMF | -1.29131 | -0.04992 | -1.18739 | -0.31955 | -1.31865 | -1.07765 | 1.89784 | 6.05964 | 0.17208 | ROCK1, ROCK2 (ROCKa), Leucine-Rich Repeat Kinase 2 (LRRK2; Dardarin) Inhibitor |
| 240 | 238 | SU 16f | -1.20460 | 1.01877 | -1.84864 | -1.47007 | -3.21789 | -3.69699 | -3.97215 | -5.60636 | -1.26844 | PDGFRb (PDGFR, PDGFR1), KDR (VEGFR2, VEGFR, FLK1) |
| 241 | 239 | PIM 1 Inhibitor 2 | -1.64004 | -1.96256 | -2.22430 | -2.59516 | -4.84094 | -5.97249 | -6.61822 | -7.51969 | -2.11683 | PIM1, PIM2, Glycogen Synthase Kinase 3 (GSK-3) Inhibitor, Janus Kinase (Jak) Inhibitor |
| 242 | 240 | Src I1 | -0.47480 | -0.47647 | -0.71156 | 0.04363 | -1.27750 | -0.68742 | 0.50762 | -0.26274 | -0.21180 | SRC (c-SRC), LCK (p56lck), KDR (VEGFR2, VEGFR, FLK1) |
| 243 | 241 | PP-1, PP1, AGL-1872, Tyrphostin-PP1 | -1.98439 | -0.90913 | 1.30381 | -0.42018 | -0.61224 | -0.70950 | 0.14935 | -0.80147 | -0.25268 | SRC (c-SRC) |
| 244 | 242 | HA 1100 hydrochloride, Hydroxyfasudil | -0.90889 | -1.05931 | 0.03018 | 1.17883 | -0.07553 | 0.09138 | -2.54700 | -0.88188 | -0.26464 | ROCK1, ROCK2 (ROCKa) |
| 245 | 243 | Cyclapolin 9, Cyclapolin9 | 0.52188 | 1.10394 | -1.21710 | 0.42077 | 2.01164 | 0.30045 | 0.12529 | -0.47241 | 0.17725 | PLK1 (PLK, STPK13) |
| 246 | 244 | C-1 | -0.37322 | -1.47209 | -0.21050 | 0.09560 | -0.03311 | -1.16552 | 1.34213 | -0.81800 | -0.16712 | PKC |
| 247 | 245 | DMPQ dihydrochloride | -0.34735 | 1.19022 | -1.37814 | 1.09015 | -1.95064 | -2.18066 | -0.53183 | -2.59821 | -0.42538 | PDGFRb (PDGFR, PDGFR1) |
| 248 | 246 | PF-02341066, PF-2341066, Crizotinib | -0.32781 | 1.42131 | -0.62692 | -1.04257 | -0.73240 | -0.88960 | -0.51903 | 1.91914 | -0.05061 | ALK, MET (c-MET, HGFR) |
| 249 | 247 | Pazopanib hydrochloride, GW-786034, Armala, Votrient, Patorma, 786034, GSK-786034, GW-786034B | 0.07956 | 0.87980 | -0.27025 | 0.20274 | 0.61735 | 1.16561 | -1.83270 | -2.21579 | -0.08713 | FLT1 (VEGFR1), KDR (VEGFR2, VEGFR, FLK1), FLT4 (VEGFR3), KIT (c-KIT), PDGFRa (PDGFR2), PDGFRb (PDGFR, PDGFR1) |
| 250 | 248 | Cediranib, AZD-2171, Recentin, NSC-732208 | -1.32627 | 0.80523 | -0.40097 | 0.33702 | -1.55455 | 0.68026 | 2.63881 | -0.75659 | 0.02683 | FLT1 (VEGFR1), KDR (VEGFR2, VEGFR, FLK1), FLT4 (VEGFR3), KIT (c-KIT), PDGFRb (PDGFR, PDGFR1) |
| 251 | 249 | XL-880, GSK-1363089, EXEL-2880, GSK-089, foretinib, 1363089 | -1.35682 | -0.55814 | -0.28957 | -0.36737 | -1.07962 | 1.06744 | -7.08482 | -5.52493 | -0.96373 | MET (c-MET, HGFR), KDR (VEGFR2, VEGFR, FLK1), FLT3 (STK1, FLK2), TIE2 (TEK) |
| 252 | 250 | PD04217903, PF-04217903, PF-4217903 | 1.56075 | -2.28729 | 1.21673 | 1.30180 | -1.35829 | 0.57467 | 0.47454 | 0.73631 | 0.14076 | MET (c-MET, HGFR) |
| 253 | 251 | GDC-0879, AR-00341677 | -0.28300 | -1.34317 | -0.31178 | 0.08146 | -0.68488 | -0.45177 | -2.06087 | -3.05088 | -0.51408 | BRAF (B-raf) |
| 254 | 252 | ABT-869, Linifanib, AL-39324, RG-3635 | -0.94982 | 1.81339 | -1.02615 | -0.16808 | -0.87208 | 0.33900 | -0.67357 | -2.46496 | -0.25386 | KDR (VEGFR2, VEGFR, FLK1), PDGFRb (PDGFR, PDGFR1), FMS (c-FMS, CSF-1R, CSF1R), FLT3 (STK1, FLK2) |
| 255 | 253 | AZD-1152, Barasertib | 0.72536 | -0.08247 | 11.06316 | 5.14415 | -8.96141 | -2.11816 | -10.71649 | -7.44289 | -0.78580 | AurB (AuroraB) |
| 256 | 254 | Enzastaurin, LY-317615, DB-102, 317615 | -0.52875 | 1.14171 | -0.23732 | 0.50152 | -1.45234 | -0.35703 | 0.57426 | -0.06871 | -0.02706 | AKT1 (PKBa), PKCb, p70S6K (S6K, S6K1), GSK3B |
| 257 | 255 | AP-24534, Ponatinib, Iclusig | -0.55658 | -0.71688 | -1.36526 | -0.86898 | -4.75076 | -2.46403 | -2.57319 | -4.86371 | -1.15183 | BCR, ABL1 (ABL), SRC (c-SRC), FLT3 (STK1, FLK2), KIT (c-KIT), RET, PDGFRb (PDGFR, PDGFR1), FGFR1 (FLT2), FGFR2, FGFR3 |
| 258 | 256 | MLN-8237, Alisertib | -0.61264 | -0.76077 | 1.45515 | 1.04006 | 1.81268 | 2.08853 | -6.54587 | -4.56143 | -0.38592 | AurA (AuroraA) |
| 259 | 257 | BIBF-1120, Intedanib, Vargatef, Nintedanib, Ofev | -1.34204 | 0.33467 | -1.34146 | -0.02637 | -0.71192 | 0.20415 | 2.54630 | 2.22297 | 0.11965 | KDR (VEGFR2, VEGFR, FLK1), PDGFRb (PDGFR, PDGFR1), MET (c-MET, HGFR) |
| 260 | 258 | PHA-739358, Danusertib | -1.84501 | -0.28963 | -1.12116 | 3.63226 | 10.84291 | -4.58896 | -4.42593 | -7.52877 | -0.33771 | ABL1 (ABL), RET, TRKA (TRK), AurA (AuroraA), FGFR1 (FLT2) |
| 261 | 259 | AZD 7762, AZD-7762 hydrochloride | 0.92912 | 0.83075 | 0.26465 | 1.86737 | 6.47565 | -5.44471 | -3.63029 | -7.89888 | -0.41903 | CHK1, CHK2 |
| 262 | 260 | PD-0332991, PD-332991, PD-991, Palbociclib, Ibrance | -0.18197 | -0.77051 | -0.79771 | -3.59086 | -6.73522 | -4.55456 | -4.40427 | -5.26422 | -1.66813 | CDK4/Cyclin D1, CDK6/Cyclin D3 |
| 263 | 261 | Brivanib, BMS-540215 | 0.01174 | 0.74705 | 1.37774 | -0.44532 | -0.14964 | -0.78993 | -1.20031 | -1.12822 | -0.10002 | KDR (VEGFR2, VEGFR, FLK1), FGFR1 (FLT2), FGFR3, FLT4 (VEGFR3), FGFR2 |
| 264 | 262 | Janex-1, WHI-P131, DDE-9501, HI-P131 | -0.55738 | -0.15902 | 1.19013 | -0.39956 | -1.26828 | -1.13664 | -0.41362 | 0.59662 | -0.13623 | JAK3 |
| 265 | 263 | GSK-1904529A, GSK1904529A | -0.00296 | -0.51489 | 1.10118 | 1.33393 | -0.55213 | 1.80039 | 4.49677 | 1.97925 | 0.61155 | IGF1R (JTK13, IGFIR) |
| 266 | 264 | MK-2206, NSC-749607 | -0.21825 | 0.27146 | 0.23518 | 1.60555 | -0.59223 | 3.86879 | 1.05762 | 2.76571 | 0.57047 | AKT1 (PKBa) |
| 267 | 265 | SGX-523 | 0.08002 | 0.25747 | -1.07497 | 0.82361 | -1.13926 | -0.23311 | 0.17572 | -0.63964 | -0.11101 | MET (c-MET, HGFR) |
| 268 | 266 | XL-147, XL147, SAR-245408 | -0.91149 | -0.22600 | 1.60816 | -0.87064 | -1.08812 | 0.54355 | -0.85930 | -0.05635 | -0.11799 | PI3K |
| 269 | 267 | LY 333531 mesylate, Ruboxistaurin | 0.74378 | -0.64862 | 0.00583 | 1.65156 | -1.38514 | -0.70473 | -1.81491 | -0.92518 | -0.19520 | PKCb |
| 270 | 268 | Alvocidib, HMR-1275, L-868275, MDL-107826A, NSC-649890, L-868276, Flavopiridol | -1.90124 | -1.17827 | -1.62881 | -3.63006 | -4.71253 | -6.20594 | -2.51697 | -7.18158 | -1.83660 | CDC2 (CDK1, p34cdc2, CDC28p)/Cyclin B, CDK2/Cyclin A, CDK2/Cyclin E, CDK4/Cyclin D1, CDK6/Cyclin D3, CDK7, CDK9/Cyclin T1, Bcl-2, Mcl-1, Survivin, XIAP |
| 271 | 269 | JNJ-38877605 | -1.34177 | -1.77442 | -0.16330 | 0.71227 | -0.08741 | -0.28789 | -0.22388 | 0.23199 | -0.18613 | MET (c-MET, HGFR) |
| 272 | 270 | GSK-461364, GSK461364 | -0.86687 | 0.09558 | -0.85981 | -6.99966 | -9.42525 | -7.92421 | -3.49711 | -5.37329 | -2.21053 | PLK1 (PLK, STPK13) |
| 273 | 271 | BI-6727, volasertib | -0.19538 | 0.31774 | -1.40066 | -3.26032 | -8.53503 | -7.16762 | 1.36039 | -6.85323 | -1.63228 | PLK1 (PLK, STPK13) |
| 274 | 272 | AC220, AC-220, Quizartinib, AC010220 | 0.09306 | -0.25830 | -0.39351 | -1.97231 | -2.18681 | -1.27178 | -0.61948 | 2.87493 | -0.23686 | FLT3 (STK1, FLK2), FMS (c-FMS, CSF-1R, CSF1R), KIT (c-KIT), PDGFRb (PDGFR, PDGFR1), RET |
| 275 | 273 | AG13958, AG-013958 | -1.84386 | -0.50771 | -1.09857 | 0.38135 | -1.96725 | -0.48918 | -1.92144 | -0.97785 | -0.53436 | PDGFRb (PDGFR, PDGFR1), KDR (VEGFR2, VEGFR, FLK1) |
| 276 | 274 | Akt-I-1,2 | -0.32237 | -0.28483 | -0.96528 | 1.43368 | -1.08145 | 0.67270 | 0.19491 | 1.00934 | 0.04165 | AKT1 (PKBa), AKT2 (PKBb) |
| 277 | 275 | Akt-I-1 | -2.40780 | -1.16548 | 0.56858 | 1.21798 | -0.93014 | -1.90226 | 0.48680 | 0.03150 | -0.26011 | AKT1 (PKBa) |
| 278 | 276 | AMG-47a | -0.02424 | 0.48956 | -1.79924 | -2.03193 | -2.47412 | 1.35552 | -0.08002 | -6.12346 | -0.67792 | LCK (p56lck), SRC (c-SRC), KDR (VEGFR2, VEGFR, FLK1), p38a |
| 279 | 277 | AMG-Tie2-1, AMG-Tie-2-1 | -0.56503 | -0.77137 | 0.16391 | -0.47030 | -1.24350 | -2.64714 | -0.97426 | -3.16075 | -0.61326 | TIE2 (TEK) |
| 280 | 278 | AT9283, AT-9283 | -1.48562 | 0.24017 | 6.43135 | -5.99984 | -9.73031 | -4.52900 | -7.82599 | -12.61759 | -2.25279 | AurA (AuroraA), AurB (AuroraB), JAK2, JAK3, ABL1 mutant(ABL mutant), FLT3 (STK1, FLK2) |
| 281 | 279 | AT-7519 analogue, AT-7519M analogue | 0.65763 | -1.32418 | -0.07414 | 0.53304 | -0.94738 | 0.16595 | -0.37559 | 0.40492 | -0.06088 | Unknown |
| 282 | 280 | AV951, KRN951, Tivozanib, ASP-4130, AV-951, KRN-951, Kil-8951 | -0.52593 | -0.01596 | -1.03127 | -0.79008 | -0.76619 | 0.29683 | -1.44777 | -3.81091 | -0.51322 | FLT1 (VEGFR1), KDR (VEGFR2, VEGFR, FLK1), FLT4 (VEGFR3), KIT (c-KIT), PDGFRb (PDGFR, PDGFR1), p38 MAPK |
| 283 | 281 | AV-412, MP-412 | -0.97053 | 0.08003 | -0.42123 | 1.45818 | -3.19154 | -0.52472 | 0.85902 | -2.70090 | -0.34326 | EGFR (ERBB1, HER1), ErbB2 (TKR1, HER2, NEU) |
| 284 | 282 | AZ-960, AZD-960 | -0.55663 | -0.37152 | 0.72227 | 0.95165 | -0.89786 | 0.13249 | 0.76564 | -5.90763 | -0.32739 | JAK2 |
| 285 | 283 | BI-D1870 | 0.72715 | -0.60942 | -0.47133 | -0.01627 | -0.74889 | -0.74456 | 1.83539 | -3.50652 | -0.22419 | RSK1 (RSK, S6Ka, MAPKAPK1C) |
| 286 | 284 | Bisindoylmaleimide X, HCl salt | 0.28836 | -0.64166 | 0.78944 | 0.94573 | -1.52034 | 2.12281 | -10.53433 | -11.33748 | -1.26144 | PKC |
| 287 | 285 | BMS-2 | -0.87457 | 0.13829 | -0.62860 | 0.08402 | 0.33826 | 0.85700 | 1.12360 | -1.46861 | -0.02731 | MET (c-MET, HGFR) |
| 288 | 286 | BMS-3 | -0.12685 | 0.12855 | -0.63437 | 3.01179 | -2.32838 | -0.46681 | -3.55053 | -8.65277 | -0.80043 | LIMK1 (LIMK) |
| 289 | 287 | BMS-5 | -0.14181 | 0.25010 | 0.14274 | -0.06488 | 0.51341 | 1.45026 | -1.34409 | -0.24729 | 0.03542 | LIMK1 (LIMK) |
| 290 | 288 | BX912, BX-912 | 0.57447 | 1.35516 | -0.83897 | 0.76288 | -0.40749 | 0.10564 | 0.38832 | 9.17766 | 0.70518 | PDK1 (PDPK1) |
| 291 | 289 | CC-401, JNK-401 | -0.29791 | 0.26626 | -0.54598 | 0.35830 | 0.56842 | -0.76289 | 0.60752 | 1.56300 | 0.11143 | JNK1 (JNK), JNK2 |
| 292 | 290 | CP-724714 | 0.01716 | -1.74620 | 0.57738 | 0.64737 | -1.12275 | -1.15918 | -0.74776 | -1.52842 | -0.32110 | EGFR (ERBB1, HER1), ErbB2 (TKR1, HER2, NEU) |
| 293 | 291 | CYC-116 | -1.06975 | -2.87378 | -2.58981 | -4.40534 | -6.18069 | -5.49234 | -7.47330 | -8.64203 | -2.45641 | AurA (AuroraA), AurB (AuroraB), KDR (VEGFR2, VEGFR, FLK1) |
| 294 | 292 | CYT11387, CYT387, CYT-387, Momelotinib, GS-0387 | -0.11135 | -1.03567 | -0.08535 | 0.07341 | -2.02675 | -1.27011 | -1.91330 | -3.08224 | -0.59949 | JAK2 |
| 295 | 293 | E7080, E-7080, ER-203492-00, Lenvatinib | -1.27818 | 0.20237 | 1.01729 | -0.36017 | -2.09880 | 0.00035 | -1.43066 | -0.91342 | -0.30834 | FGFR1 (FLT2), KIT (c-KIT), PDGFRb (PDGFR, PDGFR1), FLT1 (VEGFR1), KDR (VEGFR2, VEGFR, FLK1), FLT4 (VEGFR3) |
| 296 | 294 | GSK690693, GSK-690693 | -0.87307 | 1.34636 | 0.97561 | 1.08534 | -0.33965 | 2.46274 | 1.01704 | 2.54784 | 0.52152 | AKT1 (PKBa) |
| 297 | 295 | Ki20227 (+/-), Ki-20227 | -0.30590 | -0.74783 | 1.44601 | -0.68399 | -1.15513 | 0.69812 | 1.19600 | -0.48748 | -0.00255 | FMS (c-FMS, CSF-1R, CSF1R), KDR (VEGFR2, VEGFR, FLK1), KIT (c-KIT), PDGFRb (PDGFR, PDGFR1) |
| 298 | 296 | KU0063794, KU-0063794, KU-63794 | 0.15271 | -0.29242 | 1.46480 | 0.25331 | -0.24681 | 0.85040 | -1.09172 | -0.59123 | 0.03165 | FRAP (MTOR, FRAP1), mTORC1, mTORC2 |

| | A | B | C | D | E | F | G | H | I | J | K | L |
|---|---|---|---|---|---|---|---|---|---|---|---|---|
| 299 | 297 | Merck-5, Mk-5, JAK Inhibitor I | -0.77457 | -0.43280 | 1.65659 | 0.13446 | -0.60596 | 0.10225 | 1.28423 | -0.22491 | 0.07226 | JAK1, JAK2, JAK3, TYK2, DAPK1 |
| 300 | 298 | MP-470, Amuvatinib, HPK-56 | -1.04372 | -0.96591 | 1.21588 | 0.84964 | -0.17200 | -1.40255 | 1.96411 | -0.23024 | 0.01365 | AXL (Ark), KIT (c-KIT), FLT3 (STK1, FLK2), MET (c-MET, HGFR), PDGFRa (PDGFR2), RET, RAD51 expression inhibitor |
| 301 | 299 | PD173955, PD-173955 | -0.32622 | -0.07951 | -1.33331 | -1.48072 | -3.91246 | -4.75085 | -7.92869 | -10.08056 | -1.89603 | BCR, ABL1 (ABL), SRC (c-SRC) |
| 302 | 300 | PD173955 analogue | -0.04245 | -0.83883 | -1.16887 | 0.60916 | -2.37715 | -1.81358 | 0.14567 | -1.10670 | -0.41817 | BCR, ABL1 (ABL) |
| 303 | 301 | PD04217903, PF-04217903, PF-4217903 | -0.51138 | -0.36979 | -0.15696 | 1.31203 | -0.33039 | 0.38048 | -0.23564 | -1.50171 | -0.08965 | MET (c-MET, HGFR) |
| 304 | 302 | PF562271, PF-562271 | -2.24261 | 0.23640 | -0.61917 | 2.07159 | -0.44677 | 0.40934 | 0.88575 | -0.25952 | 0.00222 | FAK (FAK1), PYK2 (FAK2, PTK2B, CAKB) |
| 305 | 303 | PHA690509, PHA-690509 | -0.46184 | -0.15897 | -1.03031 | 1.16122 | -1.44220 | 0.14129 | -0.00028 | -0.15366 | -0.12335 | CDK2/Cyclin A |
| 306 | 304 | PI-93, PIK-93 | -0.99777 | 0.08521 | 0.64048 | 0.81271 | 0.19316 | 0.69351 | 4.42558 | 3.59734 | 0.59942 | PI3Ka |
| 307 | 305 | PIK-294 | -0.77484 | 1.01166 | -1.22690 | 1.67553 | 0.66097 | 0.37382 | 1.84863 | 1.77894 | 0.33921 | PI3Ka |
| 308 | 306 | R1487, R-1487 | -0.54931 | 0.43509 | -0.92951 | 0.38099 | -1.62205 | 1.09270 | 0.18078 | -3.43118 | -0.28178 | p38 MAPK |
| 309 | 307 | RHO-15, GSK-429286 | 0.37556 | 0.14913 | 0.15016 | -0.07626 | -0.91996 | -0.75292 | -0.41953 | 0.74113 | -0.04774 | ROCK1, ROCK2 (ROCKa) |
| 310 | 308 | SB242235, SB-242235 | -1.46160 | 1.18666 | -0.29173 | 0.86763 | -1.80454 | 0.63263 | 1.34406 | 1.52992 | 0.12705 | p38 MAPK |
| 311 | 309 | SB590885, SB-590885 | -1.56716 | -2.00542 | -0.71911 | 1.87109 | 1.39908 | -0.19330 | -2.83290 | -3.21810 | -0.46086 | BRAF (B-raf) |
| 312 | 310 | SD-06, SD-0006 | -1.34155 | -0.36272 | -0.30201 | -1.21262 | -1.41004 | -1.77450 | -0.03826 | -4.00666 | -0.66273 | p38 MAPK |
| 313 | 311 | SNS-032, BMS387032, BMS-387032 | -0.08336 | 0.59621 | -0.45889 | 1.55908 | -0.08890 | 6.16921 | 3.49570 | -2.89937 | 0.52580 | CDK2/Cyclin E, CDK7, CDK9/Cyclin T1 |
| 314 | 312 | SNS-314 | -0.28614 | 0.60560 | 0.46490 | 2.42171 | -10.02855 | -9.77945 | -4.95229 | -10.38189 | -2.02567 | AurA (AuroraA), AurB (AuroraB), AurC (AuroraC) |
| 315 | 313 | SR3677, SR-3677 | 0.18885 | -0.08868 | -0.03567 | 0.09099 | -1.73203 | -1.74134 | -2.05604 | -0.70154 | -0.38536 | ROCK2 (ROCKa) |
| 316 | 314 | TAK-715 | -0.39936 | 0.64274 | 0.71644 | -0.33850 | -0.48063 | -0.59697 | -0.02209 | 0.26723 | -0.01339 | p38 MAPK |
| 317 | 315 | YM201636, YM-201636 | -1.58434 | -0.40307 | -0.44821 | 0.88065 | -0.24676 | -0.70086 | 0.46415 | -1.88129 | -0.24862 | PIKfyve (PIP5K) |
| 318 | 316 | MK-1775, MK1775 | -0.25663 | -0.08040 | -1.49649 | 0.29965 | 2.42414 | 0.53493 | -9.34917 | -8.01675 | -1.01110 | WEE1 |
| 319 | 317 | Lim2 Kinase Inhibitor | -1.67130 | -0.57396 | -1.07248 | 0.24333 | -1.50225 | 0.63683 | -0.84763 | -0.45352 | -0.33243 | LIMK2 |
| 320 | 318 | AZD-1152-HQPA, Barasertib-HQPA | -0.38263 | 1.44667 | 7.61756 | 11.60806 | -9.51711 | -2.67248 | -8.51675 | -9.31769 | -0.61744 | AurB (AuroraB) |
| 321 | 319 | ERK2 inhibitor | 0.33813 | 0.30502 | -0.62737 | -0.95269 | -0.13047 | 0.70065 | -0.68157 | -3.78005 | -0.30625 | Erk2 (ERK, p38) |
| 322 | 320 | PV-1019, NSC-744039 | -0.17921 | 0.52973 | 0.21015 | -0.56550 | -0.65917 | -1.78533 | -1.40608 | -0.01827 | -0.24570 | CHK2 |
| 323 | 321 | MAZ51, MAZ-51 | 2.09282 | -1.00669 | 0.04663 | -0.55724 | 0.00709 | -0.22492 | -1.96595 | -0.73511 | -0.14864 | FLT4 (VEGFR3) |
| 324 | 322 | IC86621, IC-86621 | 0.89392 | 0.11026 | -1.04467 | -0.04723 | 1.55219 | 1.12067 | 0.40976 | -0.64480 | 0.14906 | DNAPK (DNA-PKcs) |
| 325 | 323 | ST638, ST-638 | 0.51373 | 0.46465 | -0.85940 | 0.22408 | 0.77168 | -1.49721 | 1.18593 | -0.06885 | 0.04660 | Tyrosine kinase inhibitor |
| 326 | 324 | TG003, TG-003 | -0.72330 | 2.06723 | 0.66469 | 1.11826 | 1.00122 | 0.88312 | 0.05012 | 0.45861 | 0.35012 | CLK1, CLK4 |
| 327 | 325 | Cyclapolin 9, Cyclapolin9 | -0.44745 | 2.00767 | 0.22608 | -0.24292 | 0.76899 | -1.37312 | 0.29276 | -1.20806 | 0.00152 | PLK1 (PLK, STPK13) |
| 328 | 326 | PF431396, PF-431396 | 0.72488 | 1.96986 | 0.19072 | -0.88565 | 0.96478 | -0.66387 | 0.39213 | -7.91306 | -0.33111 | FAK (FAK1), PYK2 (FAK2, PTK2B, CAKB) |
| 329 | 327 | (+)-P276, P276-00, Riviciclib hydrochloride | -1.23733 | -0.82168 | -0.22726 | -0.05365 | -0.65020 | -2.10509 | -5.44732 | -9.83039 | -1.29223 | CDC2 (CDK1, p34cdc2, CDC28p)/Cyclin B, CDK4/Cyclin D1 |
| 330 | 328 | CX-4945, Silmitasertib | 1.93902 | 1.47437 | 1.21997 | 1.17579 | -0.23598 | -0.55948 | -0.71879 | 0.66969 | 0.31490 | CK2a1 (CK2, CKII, Casein kinase2) |
| 331 | 329 | Dianilinopyrimidine_01 | -0.59428 | 0.79324 | -1.15631 | -1.08855 | -2.66677 | -3.17864 | -8.44699 | -3.24516 | -1.24215 | Pan kinase |
| 332 | 330 | BIRB 796, Doramapimod | 0.62274 | 1.91534 | 0.91914 | -1.01113 | 2.66703 | -1.31773 | 0.29010 | 0.14422 | 0.26828 | p38 MAPK |
| 333 | 331 | NIK Kinase Inhibitor | -0.64649 | 0.22543 | -0.34250 | 0.28663 | 0.68932 | -0.06213 | -2.05982 | -3.47929 | -0.34181 | NIK |
| 334 | 332 | Neratinib, HKI-272, CPD-820 | 0.88281 | 1.02770 | -0.53018 | -0.12164 | 1.48772 | -3.03511 | -2.54119 | -8.06940 | -0.69133 | EGFR (ERBB1, HER1), ErbB2 (TKR1, HER2, NEU) |
| 335 | 333 | EMD-1214063, Tepotinib, MSC-2156119, MSC-2156119J | 1.85694 | 0.41159 | 0.20730 | -0.05920 | 2.31119 | 4.57982 | 5.28338 | -0.61961 | 0.88619 | MET (c-MET, HGFR) |
| 336 | 334 | SMG-1 Compound 467 | 0.41779 | 0.15386 | -0.08561 | -0.18789 | 3.80927 | 1.18244 | -0.68806 | -0.81804 | 0.24000 | SMG1 (ATX, LIP) |
| 337 | 335 | ENMD-2076, ENMD-981693 | -0.44088 | -0.71448 | 0.20908 | 0.75249 | -0.00138 | -1.49629 | 1.62344 | -8.32710 | -0.53249 | AurA (AuroraA) |
| 338 | 336 | AS-703026, MSC1936369B | 1.07282 | 1.07854 | 1.34644 | 0.99884 | 1.50903 | 0.03460 | -2.06735 | -2.62326 | 0.08561 | MAP2K1 (MEK1), MAP2K2 (MEK2) |
| 339 | 337 | AZD-8055, AZD 8055, AZD8055 | 0.60705 | 0.78090 | -0.66828 | -0.28146 | -1.10703 | -2.59965 | -0.35278 | 1.51925 | -0.13333 | FRAP (MTOR, FRAP1), mTORC1, mTORC2 |
| 340 | 338 | OSI-027 hydrochloride | 0.17181 | 0.84546 | -0.15581 | 1.50040 | 0.06157 | -0.61294 | 0.16459 | -3.50496 | -0.09704 | FRAP (MTOR, FRAP1), mTORC1, mTORC2 |
| 341 | 339 | AMG-900 | 0.57759 | 0.82868 | 0.10513 | -5.92230 | -5.25704 | -5.47460 | -5.90562 | -7.46239 | -1.80839 | AurA (AuroraA), AurB (AuroraB), AurC (AuroraC) |
| 342 | 340 | BKM120, BKM-120, NVP-BKM-120, Buparlisib | -0.30161 | -0.28620 | 1.16313 | 0.06070 | 2.43245 | -0.20587 | 2.81657 | 4.05093 | 0.61717 | PI3K |
| 343 | 341 | AB12134C3, PNK inhibitor | 0.85938 | 1.21399 | -0.33840 | 0.78062 | 1.35103 | -0.11013 | 0.26109 | -0.99084 | 0.19198 | PNK (PNKP) |
| 344 | 342 | PERK inhibitor, CP-22, CP-941222, PF-941222 | -0.69440 | 1.20051 | 0.38698 | 0.84489 | 1.07603 | 1.64860 | -4.38930 | -6.41517 | -0.40226 | PEK (PERK) |
| 345 | 343 | PKC theta inhibitor | 0.44532 | 0.56512 | -1.00453 | -0.58891 | 0.40519 | -0.20343 | 1.17218 | 0.75436 | 0.09802 | PKCt |
| 346 | 344 | ILK inhibitor | 1.21081 | 1.29101 | 0.36720 | -0.87294 | 0.15019 | 0.17606 | 0.61382 | 0.22979 | 0.20081 | ILK |
| 347 | 345 | PF-04691502, PF4691502, PF-502 | 0.52580 | 0.42067 | 0.75569 | 2.69079 | 3.48830 | 3.90520 | 6.79330 | 3.15488 | 1.37860 | PI3K, FRAP (MTOR, FRAP1) |
| 348 | 346 | ASC-033, APY 33 | -0.48516 | 0.72114 | -0.55363 | 0.06677 | -1.14551 | -3.30359 | -2.85984 | -4.73545 | -0.77987 | IRE1 (IRE1a, ERN1) |
| 349 | 347 | ASC-069, APY 69 | -1.21255 | -3.85769 | -10.85857 | -10.08551 | -7.97175 | -8.03832 | -3.13315 | -3.44030 | -3.08250 | IRE1 (IRE1a, ERN1) |
| 350 | 348 | ASC-081, APY 81 | -0.33483 | -0.42014 | -0.34431 | -1.66900 | -1.86310 | -4.59602 | -5.86315 | -9.26462 | -1.54482 | IRE1 (IRE1a, ERN1) |
| 351 | 349 | ASC-082, APY 82 | -3.24460 | -3.19164 | -3.63123 | -4.09844 | -3.99652 | -10.46328 | 0.42107 | -3.98688 | -2.04187 | IRE1 (IRE1a, ERN1) |
| 352 | 350 | ASC-086, APY 86 | -2.59721 | -3.40860 | -4.70232 | -3.34702 | -4.06715 | -3.26068 | -2.56220 | -0.36530 | -1.54198 | IRE1 (IRE1a, ERN1) |
| 353 | 351 | Quinalizarin | 0.21496 | -0.17364 | -0.56658 | 0.69142 | 0.84151 | 0.15417 | -0.16541 | -0.78420 | 0.01346 | CK2a1 (CK2, CKII, Casein kinase2) |
| 354 | 352 | WZ4002, WZ-4002 | 0.64830 | -0.18431 | 0.20773 | 1.10059 | 1.10489 | 0.21620 | 0.93798 | 2.07266 | 0.38717 | EGFR (ERBB1, HER1) |
| 355 | 353 | Pp242, Torkinib, PP-242 | -0.58192 | 0.31723 | 0.72403 | -0.90277 | 0.83016 | -0.77537 | 0.32746 | -0.91696 | -0.06204 | FRAP (MTOR, FRAP1) |
| 356 | 354 | OSI-906, Linsitinib, ASP-7487 | -0.44495 | 0.24048 | -1.26582 | -1.99589 | -2.63773 | -3.57023 | -3.36891 | -3.16788 | -1.02824 | IGF1R (JTK13, IGFIR) |
| 357 | 355 | INCB018424, INC-424, INCB-018424, INCB-424, INCB-18424, Ruxolitinib, Jakafi, Jakavi | 0.02988 | 1.03949 | 0.58917 | 0.24405 | -0.20678 | 2.17728 | -0.79403 | -2.67562 | 0.02559 | JAK1, JAK2, JAK3, TYK2 |
| 358 | 356 | PLX-4032, Vemurafenib, Zelboraf, RG-7204, RO-5185426 | -0.75253 | -0.21464 | 0.08261 | 0.22881 | 0.24766 | -1.34420 | -0.35603 | -0.80146 | -0.18456 | BRAF (B-raf) |
| 359 | 357 | BMS-754807 | 0.72127 | 0.74591 | 2.02609 | 3.05749 | 2.90388 | 2.58275 | 2.18703 | -8.89503 | 0.33804 | IGF1R (JTK13, IGFIR) |
| 360 | 358 | MK-8033 | -1.50676 | 0.16820 | 1.07238 | -0.18106 | -0.08253 | -0.03120 | -0.95025 | -0.53212 | -0.12961 | MET (c-MET, HGFR) |
| 361 | 359 | GSK-1120212, Trametinib, Mekinist, JTP-74057, NSC-763093 | -0.56156 | 3.02501 | 0.81453 | -1.41074 | -2.86671 | -7.88003 | -3.49266 | -2.87754 | -0.96727 | MAP2K1 (MEK1), MAP2K2 (MEK2) |
| 362 | 360 | BGT-226, NVP-BGT-226, NVP-BGT226 | 0.39205 | 1.27371 | 0.43759 | 0.48255 | -0.82592 | -0.03414 | -2.11604 | -2.14440 | -0.16077 | PI3K, FRAP (MTOR, FRAP1) |
| 363 | 361 | TG-101348 | -0.75986 | 0.43675 | 0.34049 | 0.43228 | 1.01170 | -0.01998 | -3.38837 | -6.17675 | -0.51528 | FLT3 (STK1, FLK2), JAK2 |
| 364 | 362 | Cdk1/2 Inhibitor III | 0.10909 | -2.17837 | -0.79626 | -0.19638 | -0.14653 | -6.24380 | -6.43761 | -8.07120 | -1.51982 | CDC2 (CDK1, p34cdc2, CDC28p)/Cyclin B, CDK2/Cyclin A |
| 365 | 363 | RO-5126766 | -0.18273 | 1.33507 | 0.67716 | 1.44627 | 1.62963 | -0.46072 | 1.37276 | 3.14937 | 0.56875 | MAP2K1 (MEK1), BRAF (B-raf), RAF1 (c-Raf) |
| 366 | 364 | BS-181 hydrochloride | 0.04316 | -0.17527 | -1.03273 | 0.28235 | -0.26369 | -1.34469 | -1.76155 | -3.72090 | -0.50574 | CDK7 |
| 367 | 365 | KU-60019, KU-0060019 | 0.47154 | 0.05848 | 0.43637 | -0.65776 | -0.51609 | -0.64769 | -0.29330 | 0.00574 | -0.07248 | ATM |
| 368 | 366 | XL-184, XL184, BMS-907351, Cabozantinib S-malate | -0.64548 | 0.63083 | -1.25864 | 1.05541 | 0.16097 | -0.67562 | 0.08201 | -0.20329 | -0.05416 | FLT3 (STK1, FLK2), MET (c-MET, HGFR), KIT (c-KIT), RET, TIE2 (TEK), KDR (VEGFR2, VEGFR, FLK1) |
| 369 | 367 | CP466722, CP-466722 | -0.38519 | 1.26935 | 0.43369 | -1.71645 | -0.38354 | -1.98102 | -1.22853 | -2.42533 | -0.40702 | ATM |
| 370 | 368 | XMD8-92, XMD-8-92 | 1.68532 | -0.16305 | -0.47423 | 1.09285 | 1.59142 | -0.98650 | -0.91387 | -4.66399 | -0.17963 | Erk5 (BMK1, PRKM7) |
| 371 | 369 | AZ628, AZ-628 | -0.75476 | -0.07334 | -0.99459 | -1.02365 | -1.70161 | -3.49970 | -3.16451 | -1.24027 | -0.78984 | BRAF (B-raf), RAF1 (c-Raf) |
| 372 | 370 | S-99 | -1.12803 | 1.77891 | -1.65835 | 0.60759 | -0.25171 | -1.65713 | 0.38731 | -0.82943 | -0.17448 | MAP3K5 (ASK1) |
| 373 | 371 | CHIR-124 | 0.03337 | 1.81499 | 3.15488 | 9.20932 | -8.40517 | -10.29549 | -6.38049 | -4.44501 | -0.97132 | CHK1 |
| 374 | 372 | AZD 7762, AZD-7762 hydrochloride | -0.61340 | 0.71270 | 0.32702 | 3.64242 | 6.46303 | -4.35017 | -3.97299 | -9.11361 | -0.43798 | CHK1, CHK2 |
| 375 | 373 | LY2784544, LY-2784544, Gandotinib | 0.54867 | 0.91120 | 1.51071 | -0.24461 | -0.00858 | -0.31549 | 0.57022 | -0.01037 | 0.18786 | JAK2 |
| 376 | 374 | AZ 3146, AZ-3146 | -0.36410 | 1.63345 | -1.05389 | 0.57972 | 0.18220 | 0.93219 | 2.50874 | 3.75852 | 0.51865 | TTK (ESK, MPS1L1, Monopolar spindle 1 (Mps1) kinase) |
| 377 | 375 | BMS 794833, BMS-794833 | -0.17056 | 1.02512 | 0.56066 | 0.29541 | 0.72221 | -1.20635 | -1.00162 | -0.53695 | -0.01979 | MET (c-MET, HGFR), KDR (VEGFR2, VEGFR, FLK1) |
| 378 | 376 | PI3-Kinase a Inhibitor 2 | 0.08056 | 1.31021 | -0.56320 | 0.57870 | -0.15722 | -0.15416 | -1.21439 | 2.66679 | 0.16157 | PI3K |
| 379 | 377 | BIX 02189, BIX-02189 | 1.03700 | 1.80552 | 0.69196 | 0.10733 | 0.69368 | 0.21737 | -0.78352 | -1.27629 | 0.15813 | MAP2K5 (MEK5), Erk5 (BMK1, PRKM7) |
| 380 | 378 | IRAK4 Inhibitor | -0.51566 | -0.57559 | -3.72854 | -4.48369 | -2.98768 | -4.02835 | -4.74621 | -9.75140 | -1.95469 | IRAK4 |
| 381 | 379 | Merck-22-6 | -0.47015 | 1.21514 | -0.05178 | 0.87765 | 0.05084 | 0.56830 | -0.61343 | 2.10474 | 0.23350 | AKT1 (PKBa), AKT2 (PKBb) |
| 382 | 380 | PF 4708671, PF-4708671, PF-04708671 | -0.32455 | 0.79020 | 0.41501 | -0.68985 | 1.96809 | 1.09755 | 3.42038 | 2.87051 | 0.60558 | p70S6K (S6K, S6K1) |
| 383 | 381 | PHA-793887 | 0.96671 | 1.48744 | 1.34577 | -0.63582 | -0.11601 | -1.06474 | -2.32730 | -0.00650 | -0.02223 | CDC2 (CDK1, p34cdc2, CDC28p)/Cyclin B, CDK2/Cyclin A, CDK2/Cyclin E, CDK4/Cyclin D1, CDK5/p25 |
| 384 | 382 | AZD5438, AZD-5438 | -0.27241 | 1.37582 | 0.01236 | -2.33994 | -3.11255 | -5.55757 | -6.62769 | -4.64620 | -1.34267 | CDC2 (CDK1, p34cdc2, CDC28p)/Cyclin B, CDK2/Cyclin A, CDK9/Cyclin T1 |
| 385 | 383 | R406, R-406, NSC-742317, Tamatinib | 0.33649 | 1.19557 | 0.34896 | 0.53874 | 0.83844 | -3.55372 | -4.75973 | -1.81023 | -0.43547 | FLT3 (STK1, FLK2), SYK, (IL-2, IL-6 and TNFa) production inhibitor |
| 386 | 384 | HMN-214, IVX-214 | -0.37422 | 1.65751 | -0.61634 | -0.26566 | -0.36085 | -1.18126 | -6.23656 | -7.50903 | -0.94423 | PLK1 (PLK, STPK13) |
| 387 | 385 | ON-01910Na, Estybon, Rigosertib sodium, ON-1910Na, Onc-01910, SyB C-1101, SyB L-1101, Novonex | -0.50088 | 0.95574 | 0.78309 | 0.52638 | -5.65224 | -9.35762 | 0.26514 | 0.24432 | -0.80783 | PLK1 (PLK, STPK13) |
| 388 | 386 | AZD8330, AZD-8330, ARRY-424704, ARRY-704 | 1.86692 | 2.95248 | 1.14522 | -1.85597 | -2.08375 | -3.88993 | -4.10984 | -4.71396 | -0.67798 | MAP2K1 (MEK1), MAP2K2 (MEK2) |
| 389 | 387 | RAF265, RAF-265, CHIR-265 | -0.33901 | -0.17666 | -0.67576 | -1.00087 | -2.79294 | -8.34576 | -9.59573 | -11.06535 | -2.15607 | KIT (c-KIT), PDGFRb (PDGFR, PDGFR1), BRAF (B-raf), RAF1 (c-Raf), KDR (VEGFR2, VEGFR, FLK1) |
| 390 | 388 | Tipifarnib, NSC-702818, R-115777, Zarnestra, IND 58359 | 0.24308 | 0.03807 | 1.22039 | -0.12132 | -0.83364 | -0.63227 | -2.05719 | -5.31391 | -0.47297 | Farnesyltransferase |
| 391 | 389 | CAL-101, GS-1101, Idelalisib, Zydelig | 1.10110 | 0.70118 | 2.13562 | 1.72947 | 2.11713 | 0.36816 | 2.55327 | 1.31239 | 0.76231 | PI3Kd |
| 392 | 390 | PIK-294 | 0.24664 | -0.18068 | -0.15169 | 1.16285 | 2.29529 | -0.77416 | -0.04695 | 0.30348 | 0.18107 | PI3Ka |
| 393 | 391 | WYE-354 | 0.14405 | 0.72836 | 1.03098 | -0.35896 | 0.33359 | -0.97400 | -0.55205 | -1.32612 | -0.06179 | FRAP (MTOR, FRAP1), mTORC1, mTORC2 |
| 394 | 392 | TC-A 2317 hydrochloride | 0.26523 | 0.91164 | -0.19163 | -0.36800 | 0.37927 | -1.23579 | -2.72922 | -8.82204 | -0.74786 | AurA (AuroraA) |
| 395 | 393 | ARQ-197, Tivantinib | -0.33516 | 1.43253 | -1.13877 | -0.56444 | -0.01075 | -2.04573 | 2.62927 | -8.40670 | -0.53532 | MET (c-MET, HGFR) |
| 396 | 394 | AZD-4547 | 0.37664 | 0.17474 | 0.47802 | -0.06940 | -0.30768 | -1.54229 | -0.48482 | 0.32833 | -0.06638 | FGFR1 (FLT2), FGFR2, FGFR3, FGFR4 |
| 397 | 395 | AZD-1480, AZD1480 | 1.96686 | 0.67902 | -0.13485 | 0.04967 | 1.07845 | -1.17244 | 1.29828 | -2.08874 | 0.10632 | JAK1, JAK2, JAK3 |
| 398 | 396 | BAY 73-4506, Regorafenib | -0.17688 | 0.29872 | -0.22401 | 1.80255 | -1.34188 | -4.17637 | -2.31951 | -2.29607 | -0.53492 | KIT (c-KIT), PDGFRb (PDGFR, PDGFR1), RET, RAF1 (c-Raf), BRAF (B-raf), TIE2 (TEK), KDR (VEGFR2, VEGFR, FLK1), p38 MAPK |
| 399 | 397 | JAK compound I (VI-53), R-333, R-932333 | 0.49834 | 0.38490 | 0.29656 | -0.64008 | 1.04680 | -1.03733 | 0.21265 | 0.48786 | 0.07927 | JAK1, JAK2, JAK3 |
| 400 | 398 | Rigel JAK Compound II, Prodrug of VI-53, R-348, R348, R-932348 | 1.18445 | -0.06522 | 1.44031 | -0.45938 | 0.72777 | -0.99480 | 0.14084 | -0.31874 | 0.10499 | JAK1, JAK2, JAK3 |
| 401 | 399 | TBK1 Inhibitor | 0.41570 | 1.39481 | -0.49357 | -0.50130 | 1.11117 | -1.71734 | -1.14171 | -3.85901 | -0.30390 | TBK1 |
| 402 | 400 | R788, R-788, Fostamatinib disodium, R935788, R-935788, NSC-745942 | -0.28480 | 0.15652 | 0.33461 | 1.35498 | -0.19012 | -1.17823 | -0.03178 | -3.27665 | -0.19761 | FLT3 (STK1, FLK2), SYK |
| 403 | 401 | PF 3644022, PF-3644022, MK2 inhibitor, compound 35 | -1.14310 | -0.34302 | -0.64697 | -1.96432 | -1.30325 | -3.00970 | -3.35987 | -3.77567 | -0.98606 | MAPKAPK2 (MK2) |
| 404 | 402 | GLPG-0259, Compound A | -0.83601 | 0.07388 | -0.89239 | -0.95479 | -1.59445 | -2.34964 | -3.53413 | -3.89733 | -0.88704 | MAPKAPK5 (PRAK) |
| 405 | 403 | ASP-3026 | -0.99540 | 0.82680 | -0.69977 | -0.00218 | 1.54094 | 1.91870 | 4.02635 | 10.76192 | 1.10222 | ALK, EGFR (ERBB1, HER1) |
| 406 | 404 | SJN 2511 | 1.05872 | 0.63413 | 1.24549 | 2.71835 | 2.60050 | 1.30751 | 0.35297 | -0.87037 | 0.57386 | TGFbR1 (ALK5) |
| 407 | 405 | BGJ-398, NVP-BGJ-398, NVP-BGJ398, Infigratinib | -0.63806 | 1.13399 | -1.06655 | -0.31128 | 1.63027 | 0.45959 | 3.75295 | 6.31258 | 0.71506 | FGFR1 (FLT2), FGFR2, FGFR3, FGFR4 |
| 408 | 406 | BMS 777607, BMS-777607 | 1.07356 | 1.04261 | 2.03117 | 0.67353 | -0.10017 | -0.94786 | -0.62751 | -1.02085 | 0.13475 | MET (c-MET, HGFR), AXL (Ark), RON (MST1R), TYRO3 (RSE, SKY, TIF, BYK) |
| 409 | 407 | Y-39983, RKI-983, SNJ-1656 | 1.17427 | 0.86049 | -0.62839 | -0.84997 | -0.61134 | -2.94162 | -1.02082 | -1.42544 | -0.34523 | ROCK1, ROCK2 (ROCKa) |
| 410 | 408 | GSK-1059615, GSK-615 | -0.23281 | 0.75761 | -0.23305 | 0.57488 | 0.30035 | 0.74523 | 1.34173 | 2.33278 | 0.35436 | PI3K |
| 411 | 409 | PCI-32765, Ibrutinib | -0.10068 | 1.45444 | -0.69726 | 0.93514 | 0.67172 | 0.23292 | -0.18127 | 0.96314 | 0.20793 | BTK (Bruton's tyrosine kinase), LCK (p56lck), LYN |
| 412 | 410 | Y-27632 | -0.74600 | 0.80118 | -0.34245 | 1.13340 | 0.56195 | -0.33311 | -1.12933 | -2.98888 | -0.19303 | LRRK2, ROCK1, ROCK2 (ROCKa), Calcium sensitizer |
| 413 | 411 | LDN193189, LDN-193189, DM-3189, LJPC-6417 | -0.58130 | 0.95283 | 1.39007 | 0.18596 | 1.72011 | 0.01376 | 1.25972 | 0.44341 | 0.34153 | ALK2, BMPR1A (ALK3) |
| 414 | 412 | A 769662, A-769662 | -0.59513 | -0.06869 | -0.24361 | 0.62566 | 1.85733 | -0.70267 | -0.04626 | -0.65602 | 0.01082 | AMPK |
| 415 | 413 | WZ8040, WZ-8040 | -0.69366 | 1.04110 | -0.34803 | 0.24374 | 0.69791 | -0.18035 | -0.66641 | 2.14284 | 0.14190 | EGFR (ERBB1, HER1) |
| 416 | 414 | Pelitinib, EKB-569, WAY-EKB-569 | -0.31552 | 1.31663 | -0.56899 | -0.35525 | 0.09687 | 0.54057 | -6.25658 | -7.87627 | -0.85112 | EGFR (ERBB1, HER1) |
| 417 | 415 | WZ3146, WZ-3146 | -0.32073 | 0.98733 | 0.01650 | 0.05157 | -0.50563 | 0.10410 | -0.03371 | -0.14212 | 0.00998 | EGFR (ERBB1, HER1) |
| 418 | 416 | KRN 633, KRN-633 | 0.19936 | 0.29872 | 0.64166 | 1.37197 | 0.26256 | 0.00716 | -0.70633 | -0.55025 | 0.09672 | FLT1 (VEGFR1), KDR (VEGFR2, VEGFR, FLK1), FLT4 (VEGFR3) |
| 419 | 417 | MGCD-265 | 0.54846 | 0.87704 | 0.20754 | 0.36190 | 0.85077 | 0.04225 | 1.33420 | 1.02905 | 0.33308 | MET (c-MET, HGFR), RON (MST1R), TIE2 (TEK), FLT1 (VEGFR1), KDR (VEGFR2, VEGFR, FLK1), FLT4 (VEGFR3) |
| 420 | 418 | OSI-930 | -0.69258 | -0.00381 | -0.08825 | -1.19118 | -1.88105 | -1.62014 | -2.72153 | 0.32864 | -0.49918 | KIT (c-KIT), KDR (VEGFR2, VEGFR, FLK1), FLT1 (VEGFR1), FMS (c-FMS, CSF-1R, CSF1R), RAF1 (c-Raf), LCK (p56lck), PDGFRb (PDGFR, PDGFR1) |
| 421 | 419 | NVP-ADW742, ADW-742, ADW, GSK-552602-A, NVP-ADW-742 | -0.49172 | -0.00710 | -1.07114 | 1.47503 | 1.15620 | 3.14497 | 4.15786 | 5.25446 | 0.86381 | IGF1R (JTK13, IGFIR) |
| 422 | 420 | KW 2449, KW-2449 | 0.01211 | 0.22779 | -0.60212 | -0.09847 | 2.06012 | -0.66102 | 1.12407 | -5.97516 | -0.24818 | FLT3 (STK1, FLK2), ABL1 (ABL), FGFR1 (FLT2), AurA (AuroraA) |
| 423 | 421 | SB-525334 | -0.08058 | -0.18313 | -0.91035 | 1.07211 | 3.39766 | 0.32751 | 2.35561 | 1.88797 | 0.49898 | TGFbR1 (ALK5) |
| 424 | 422 | CCT129202, CCT-129202 | 0.60983 | 0.62688 | 0.00602 | -1.13605 | 0.62303 | -0.44756 | -0.97792 | -1.27958 | -0.12530 | AurA (AuroraA), AurB (AuroraB), AurC (AuroraC) |
| 425 | 423 | Aurora A Inhibitor I, TC-S 7010 | 0.07709 | 0.83581 | -0.34708 | -0.49265 | 0.05889 | 0.07106 | 0.50996 | -5.07177 | -0.27647 | AurA (AuroraA) |
| 426 | 424 | Hesperadin | 0.22989 | 1.42697 | -1.53025 | -8.54234 | -4.98936 | -7.27062 | -4.71342 | -7.43915 | -2.08225 | AurB (AuroraB), Tyrosine Kinase inhibitor |
| 427 | 425 | PHA-680632 | 0.24927 | 0.91460 | -0.43135 | 0.33063 | 1.34405 | 1.29338 | 12.36003 | -0.85646 | 0.96438 | AurA (AuroraA), AurB (AuroraB), AurC (AuroraC) |
| 428 | 426 | PD318088, PD-318088 | -0.49851 | 1.32519 | -0.87480 | 2.86485 | 2.96703 | 0.38738 | 2.36419 | 1.87242 | 0.66015 | MAP2K1 (MEK1), MAP2K2 (MEK2) |
| 429 | 427 | LY2228820, LY-2228820, LSN-2322600, Ralimetinib mesylate | 0.32124 | 0.10909 | -0.78161 | -0.56209 | 1.07942 | -0.63807 | 1.34322 | -0.36784 | 0.03193 | p38a, p38b |
| 430 | 428 | AS605240, AS-605240 | 0.01207 | 0.88298 | 0.25412 | 0.62761 | -0.43591 | 0.83345 | 0.97496 | 2.73365 | 0.37315 | PI3K |
| 431 | 429 | VX-745 | -0.26031 | 1.45764 | 0.55680 | 0.90308 | 1.23397 | -0.91604 | 0.72826 | 0.01320 | 0.23574 | p38 MAPK |
| 432 | 430 | AZD6482, AZD-6482, KIN-193 | -0.57365 | 2.03904 | 0.24216 | -0.25282 | 2.55190 | 1.39368 | 2.00468 | 2.05750 | 0.60019 | PI3K |
| 433 | 431 | TG100-115, TG-100-115, TG-100115 | -0.17957 | 1.14411 | -0.01769 | -0.21570 | -0.07487 | -0.10897 | 0.77807 | 0.98356 | 0.14645 | PI3K |
| 434 | 432 | XL765, XL-765, SAR245409, Voxtalisib | -0.60984 | 0.21150 | -0.64909 | -1.81405 | 0.07930 | -0.77632 | -1.65671 | -3.43398 | -0.54861 | PI3K, mTORC1, mTORC2 |
| 435 | 433 | Indirubin, NSC-105327, Couroupitine B, Indigo red, Indigopurpurin | -2.48729 | -3.39287 | -5.52549 | -7.70916 | -7.02798 | -9.61157 | -9.67113 | -9.03587 | -3.45441 | CDC2 (CDK1, p34cdc2, CDC28p)/Cyclin B, CDK2/Cyclin A, CDK2/Cyclin E, CDK4/Cyclin D1, CDK5, GSK3B |
| 436 | 434 | AT7867, AT-7867, AT-867 | 0.26710 | 0.61633 | -0.92492 | -0.30654 | -0.20992 | -1.97986 | -1.90781 | -0.60173 | -0.32015 | AKT1 (PKBa), AKT2 (PKBb), AKT3 (PKBg), p70S6K (S6K, S6K1) |
| 437 | 435 | AEE 788, AEE-788, NVP-AEE-788 | -0.23890 | 1.71494 | 0.51054 | 0.35370 | 0.07321 | 0.97461 | 2.18034 | 4.34158 | 0.62858 | EGFR (ERBB1, HER1), ErbB2 (TKR1, HER2, NEU), KDR (VEGFR2, VEGFR, FLK1) |
| 438 | 436 | AZD-5363, AZD5363 | -0.75847 | 0.87918 | 1.19479 | 0.36890 | -0.66781 | 0.06664 | 1.89189 | 0.79025 | 0.23883 | AKT1 (PKBa), AKT2 (PKBb), AKT3 (PKBg), p70S6K (S6K, S6K1) |
| 439 | 437 | BMS-582664, Brivanib alaninate | 0.16330 | 2.16924 | -0.85789 | -0.46755 | 0.77581 | 0.17815 | 0.76195 | -1.75684 | 0.06128 | FGFR1 (FLT2), FGFR2, FGFR3, FLT1 (VEGFR1), KDR (VEGFR2, VEGFR, FLK1), FLT4 (VEGFR3) |
| 440 | 438 | S6K-18 | -0.56234 | 0.30787 | 1.00820 | -0.29545 | 0.73798 | -0.95951 | -0.41684 | -0.91210 | -0.06928 | p70S6K (S6K, S6K1) |
| 441 | 439 | NVP-BSK805, BSK-805, NVP-BSK-805 | 0.06533 | 0.54613 | -0.32872 | 1.00914 | 0.15751 | -0.71536 | -1.33535 | -0.58864 | -0.07548 | JAK2 |
| 442 | 440 | JNJ-38158471 | -0.07756 | 1.44248 | -0.39008 | 0.64367 | -0.08980 | 0.00911 | 0.95436 | 0.47615 | 0.18828 | KDR (VEGFR2, VEGFR, FLK1), RET, KIT (c-KIT) |
| 443 | 441 | OSI-TAK1 inhibitor | 1.58244 | 0.62448 | -0.06067 | -1.93487 | 1.02108 | -1.03239 | -3.63072 | -7.07217 | -0.66618 | TAK1 (TGF1a, MAP3K7) |
| 444 | 442 | SCH900776, SCH-900776, MK-8776 | 0.03841 | 0.71681 | -0.31668 | 1.25980 | 0.53959 | 3.74345 | -4.43274 | -6.25196 | -0.29833 | CHK1, CDK2/Cyclin A, CDK2/Cyclin E |
| 445 | 443 | GSK-2118436B, GSK2118436B, Dabrafenib mesylate | 0.24270 | -1.27564 | -0.20556 | -0.04617 | -0.59382 | -0.88456 | -1.92001 | -4.83334 | -0.60361 | RAF1 (c-Raf), BRAF (B-raf) |
| 446 | 444 | CH5424802, AF-802, CH-5424802, Alectinib, RG-7853, RO-5424802 | 0.04919 | -0.00658 | 0.58909 | -0.74029 | -0.39101 | -0.57160 | -0.75694 | 0.54146 | -0.08161 | ALK |
| 447 | 445 | AT-7519 hydrochloride, AT7519 hydrochloride | 0.37270 | -0.18213 | -1.52538 | -1.50999 | -1.53845 | -2.01471 | -0.35029 | -9.98646 | -1.06146 | CDC2 (CDK1, p34cdc2, CDC28p)/Cyclin B, CDK2/Cyclin A, CDK3/Cyclin E, CDK4/Cyclin D1, CDK5/p35, CDK6/Cyclin D3 |
| 448 | 446 | LY2835219 derivative, LY-2835219 derivative | 0.85897 | 1.36945 | 1.06617 | -1.02199 | 0.29409 | -1.83255 | -0.57933 | -0.42978 | -0.01744 | Unknown |
| 449 | 447 | AZ TAK1 Inhibitor | 0.48231 | 1.31063 | -0.42494 | 0.09546 | 0.02191 | -0.18914 | 2.10228 | -7.43900 | -0.25628 | TAK1 (TGF1a, MAP3K7) |
| 450 | 448 | GSK PERK Inhibitor | 1.68748 | 2.72231 | 2.04769 | 2.88905 | 3.94604 | -0.72247 | -0.05081 | 1.34381 | 0.87932 | PEK (PERK) |
| 451 | 449 | NQDI-1 | -0.54358 | -0.89120 | -2.71435 | -6.33974 | -8.23358 | -10.42967 | -8.80602 | -10.99558 | -3.10507 | MAP3K5 (ASK1) |
| 452 | 450 | PX-866, DJM-166, DJM-2-166, Sonolisib, NSC-722134 | 1.08376 | 0.31550 | 0.33355 | 0.04286 | 1.37800 | 0.34598 | 2.87005 | 1.60420 | 0.50577 | PI3K, PTEN inhibitor |
| 453 | 451 | TCS 2312, TCS-2312 | 0.39097 | 0.90255 | -0.72886 | 0.92284 | 1.11590 | 1.17082 | 0.04584 | 4.12394 | 0.50388 | CHK1 |

| | A | B | C | D | E | F | G | H | I | J | K | L |
|---|---|---|---|---|---|---|---|---|---|---|---|---|
| 454 | 452 | OICR0000857A, 421966 | -0.11812 | -0.25703 | 1.34863 | -0.91570 | 0.72317 | -0.43073 | -1.43926 | 0.38382 | -0.04473 | Histone Deacetylase SIRT1 inhibitor |
| 455 | 453 | NVP-BHG712, NVP-BHG-712 | 0.02160 | 0.02681 | -0.18671 | -0.54932 | -1.39860 | -3.24859 | -2.40900 | -3.82986 | -0.73410 | RAF1 (c-Raf), SRC (c-SRC), ABL1 (ABL), EphB4 |
| 456 | 454 | PF 3644022, PF-3644022 | -0.13535 | -0.57271 | -1.61883 | -2.09574 | -2.26725 | -4.19350 | -2.02856 | -3.61240 | -1.04812 | MAPKAPK2 (MK2) |
| 457 | 455 | GSK 2334470, GSK-2334470 | -0.50813 | 1.13135 | 1.36121 | -0.49941 | 0.88967 | 0.14111 | 4.14950 | 0.97736 | 0.48476 | PDK1 (PDPK1) |
| 458 | 456 | TTP 22 | -0.11102 | 0.77825 | -1.27418 | 0.41426 | 1.39497 | -0.36787 | -0.27511 | -0.37916 | 0.01143 | CK2a1 (CK2, CKII, Casein kinase2) |
| 459 | 457 | GW 788388, GW-788388 | -0.55352 | 0.72501 | -1.00080 | 1.92690 | 1.61678 | 1.03865 | 1.56618 | 0.49705 | 0.36892 | TGFbR1 (ALK5) |
| 460 | 458 | TC-DAPK 6 | 2.82276 | 0.29473 | -0.27549 | 0.77777 | 1.06334 | -0.06527 | 0.62914 | 0.63910 | 0.37335 | DAPK1, DAPK3 (ZIPK, ZIP) |
| 461 | 459 | PTK6 CompoundA | 1.00741 | 0.51343 | -0.10658 | 0.10645 | 0.01841 | -0.73740 | -0.85497 | 0.79052 | 0.04676 | BRK (PTK6) |
| 462 | 460 | PTK6 Inhibitor CompoundB | 0.42285 | 0.33830 | 0.85196 | 0.59151 | 1.10165 | -1.28005 | 0.97858 | 0.56780 | 0.22661 | BRK (PTK6) |
| 463 | 461 | OICR0000666A01 | -1.35372 | 0.49001 | -0.08134 | 0.17460 | -1.12544 | -1.43183 | 1.19790 | -1.75826 | -0.24662 | PIM1 |
| 464 | 462 | RET Inhibitor Example 8 (Free base) | 1.11888 | 0.89159 | -0.54505 | -0.00628 | 1.70219 | -0.51603 | -1.20738 | -2.27935 | -0.05337 | RET |
| 465 | 463 | CCT137690, CCT-137690 | 0.06333 | -0.07572 | -0.40712 | 0.12900 | -0.62008 | 0.97601 | -8.00834 | -6.32645 | -0.90509 | AurA (AuroraA), AurB (AuroraB), AurC (AuroraC), FLT3 (STK1, FLK2) |
| 466 | 464 | AMG458 | 0.15273 | 0.41830 | -0.52600 | 0.47898 | -0.52746 | 0.19001 | 0.37066 | -0.12414 | 0.02747 | MET (c-MET, HGFR) |
| 467 | 465 | GDC-0941, Pictilisib, Pictrelisib, RG-7321 | 0.90822 | 1.96794 | 0.32664 | 2.20819 | 2.78479 | 3.67361 | 4.82061 | 6.65642 | 1.48083 | PI3K |
| 468 | 466 | MK-5108, VX-689 | -0.50625 | 0.40857 | 0.96998 | 0.86442 | 0.67772 | 0.50579 | -0.14379 | -1.47889 | 0.08230 | AurA (AuroraA) |
| 469 | 467 | BAY 57-9352, Telatinib | 0.48859 | -0.30618 | -1.47102 | -0.63204 | -0.57899 | -1.43355 | -2.64484 | -3.38736 | -0.63209 | KDR (VEGFR2, VEGFR, FLK1), FLT4 (VEGFR3), KIT (c-KIT), PDGFRb (PDGFR, PDGFR1) |
| 470 | 468 | A 1070722, A-1070722 | -1.06175 | 1.38333 | -1.06000 | -1.77815 | -3.17510 | -2.98985 | -3.53728 | -7.99201 | -1.28194 | GSK3 |
| 471 | 469 | Dacomitinib hydrate, PF-00299804, PF-00299804-03, PF-299, PF-299804 | 0.65387 | 0.23419 | 0.92838 | 1.35207 | 0.88260 | -0.19464 | 0.72226 | -0.57376 | 0.25403 | EGFR (ERBB1, HER1), ErbB2 (TKR1, HER2, NEU), ErbB4 (HER4) |
| 472 | 470 | VE-821 | 1.13221 | 2.14040 | -0.00106 | -0.92592 | -0.84976 | -3.17077 | -4.77582 | -5.45085 | -0.75490 | ATR |
| 473 | 471 | BIX-RSK2 Inhibitor | 1.14857 | 0.45142 | -0.92079 | -0.24682 | -0.57505 | -0.92604 | -0.31127 | -1.28703 | -0.16917 | RSK1 (RSK, S6Ka, MAPKAPK1C), RSK2 (S6K-a3, p90-RSK2) |
| 474 | 472 | BYL-719, NVP-BYL-719, Alpelisib | -1.03117 | 0.13282 | 0.63505 | -0.11024 | 1.16553 | 0.93816 | 0.57559 | 3.56530 | 0.37239 | PI3Ka |
| 475 | 473 | PHA-848125, PHA-848125AC | -0.78200 | 0.78094 | -1.61449 | -1.09818 | -1.51793 | -6.43431 | -8.06332 | -5.33028 | -1.52607 | CDK2/Cyclin A, CDK7/Cyclin H, CDK4/Cyclin D1, CDK5/p35, CDK2/Cyclin E, CDC2 (CDK1, p34cdc2, CDC28p)/Cyclin B |
| 476 | 474 | LY-2157299, Galunisertib | 1.07348 | 0.08544 | 1.09817 | 0.35668 | 0.53335 | -0.76582 | 1.17733 | 0.41679 | 0.25216 | TGFbR1 (ALK5), TGFbR2 |
| 477 | 475 | WAY-600 | -0.13679 | 1.84529 | -1.53006 | 0.90574 | 0.27457 | -1.47087 | -0.91227 | -0.95543 | -0.12558 | FRAP (MTOR, FRAP1), mTORC1, mTORC2 |
| 478 | 476 | Crenolanib, ARO-002, CP-868596 | 0.76298 | -0.77311 | 0.68520 | 0.79369 | 0.19243 | -1.17448 | -5.54115 | -10.10691 | -0.96166 | PDGFRa (PDGFR2), PDGFRb (PDGFR, PDGFR1) |
| 479 | 477 | WYE-125132, WYE-132 | -1.13669 | 0.60519 | -0.05905 | 0.24903 | -1.16236 | -2.19463 | -1.72464 | -1.31827 | -0.42760 | PI3K, FRAP (MTOR, FRAP1), mTORC1, mTORC2 |
| 480 | 478 | Apatinib mesylate, YN-968D1 | 0.13085 | -0.31986 | -0.24973 | -0.44771 | 1.27546 | -1.80630 | -0.33930 | -2.57795 | -0.27494 | RET, KDR (VEGFR2, VEGFR, FLK1), FLT1 (VEGFR1), SRC (c-SRC), KIT (c-KIT) |
| 481 | 479 | MLN8054, MLN-8054 | 1.47522 | -0.58746 | -0.57254 | -0.16178 | 1.17838 | -0.18640 | -0.16923 | 0.17604 | 0.07309 | AurA (AuroraA) |
| 482 | 480 | Dinaciclib, MK-7965, NSC-727135, SCH-727965 | -1.48655 | -1.08108 | 0.53275 | -4.79662 | -10.32290 | -9.09399 | -2.03158 | -5.15552 | -2.12077 | CDK2/Cyclin A, CDK5/p25, CDC2 (CDK1, p34cdc2, CDC28p)/Cyclin B, CDK9/Cyclin T1 |
| 483 | 481 | CVT-313 | -0.45206 | 0.86960 | -0.44815 | 0.45220 | -0.23981 | -1.25386 | -0.80416 | -1.40172 | -0.20792 | CDC2 (CDK1, p34cdc2, CDC28p)/Cyclin B, CDK2/Cyclin A, CDK2/Cyclin E |
| 484 | 482 | Bohemine | -0.71106 | 1.49339 | -1.00601 | 0.51880 | -0.23376 | -0.69752 | -1.22975 | -2.54232 | -0.27961 | CDC2 (CDK1, p34cdc2, CDC28p)/Cyclin B, CDK2/Cyclin A, CDK2/Cyclin E |
| 485 | 483 | Indirubin-5-sulfonic acid | -0.04936 | 0.84735 | 0.22502 | 0.51184 | 1.38153 | -0.07033 | 0.18571 | -1.40134 | 0.10342 | CDC2 (CDK1, p34cdc2, CDC28p)/Cyclin B, CDK2/Cyclin A, CDK2/Cyclin E, CDK4/Cyclin D1, CDK5/p35, GSK3B |
| 486 | 484 | CAN-508 | -0.41941 | 1.01975 | -0.15548 | 0.71501 | -0.61761 | -0.75629 | 0.11491 | 0.37154 | 0.01728 | CDK9/Cyclin T1 |
| 487 | 485 | Masitinib, AB-1010, AB1010, Masiviera | 0.08341 | 1.40270 | -0.99672 | -0.34195 | 0.35262 | -1.06505 | -0.95537 | -1.17403 | -0.17090 | KIT (c-KIT), PDGFRb (PDGFR, PDGFR1), FGFR3 |
| 488 | 486 | AG-183, Tyrphostin 51 | 0.06033 | -0.26167 | 0.23531 | -0.31619 | 0.45947 | -0.54035 | -0.26594 | 0.77344 | 0.00916 | EGFR (ERBB1, HER1) |
| 489 | 487 | Trk Inhibitor III, GNF-5837 | -0.40572 | 0.65904 | -0.56105 | -0.45104 | 0.09833 | -3.84372 | -1.84277 | -12.19499 | -1.17609 | TRKA (TRK), TRKB |
| 490 | 488 | W-7 hydrochloride | -0.33997 | -0.16195 | 0.72728 | 1.48594 | 1.24982 | -1.88444 | -0.15361 | 0.00996 | 0.05918 | smMLCK (MLCK, myosin light chain kinase, MYLK) |
| 491 | 489 | W-13 hydrochloride | -0.48430 | -0.43517 | -0.74111 | -0.48709 | 0.12512 | -0.54260 | 0.28526 | -0.19277 | -0.15684 | smMLCK (MLCK, myosin light chain kinase, MYLK) |
| 492 | 490 | Casein Kinase II Inhibitor, TBCA | 0.17964 | 0.35757 | -0.03843 | 1.38432 | 1.04927 | -0.85605 | 0.55526 | -1.20014 | 0.09079 | CK2a1 (CK2, CKII, Casein kinase2) |
| 493 | 491 | HG-10-102-01 | -0.38148 | 1.18613 | 0.56858 | 0.34466 | -0.25999 | -0.65940 | 0.40467 | -1.88758 | -0.04341 | LRRK2 (Leucine-Rich Repeat Kinase 2, Dardarin) |
| 494 | 492 | SGI-1776 | -0.67175 | -0.79451 | -0.62040 | -0.19341 | -0.30289 | -1.35180 | 1.00820 | 1.48444 | -0.09147 | PIM1, PIM2, FLT3 (STK1, FLK2) |
| 495 | 493 | PF-3758309, PF-03758309 | 1.27990 | 3.19856 | 0.88854 | -1.57240 | -2.02952 | -5.03056 | -0.51532 | -1.71015 | -0.34828 | PAK4 |
| 496 | 494 | GSK-2636771 | -0.22009 | -0.63279 | 0.35725 | 0.47656 | 0.10821 | -1.16781 | 0.81576 | 2.03681 | 0.11252 | PI3K/p110b |
| 497 | 495 | JNJ-28312141 | 0.93771 | 1.89660 | 0.11640 | 0.69366 | 0.59475 | 0.16243 | 0.44120 | -0.41071 | 0.28112 | FMS (c-FMS, CSF-1R, CSF1R) |
| 498 | 496 | Compound 34 | 0.76149 | 0.24043 | 0.54371 | 0.87814 | 1.59444 | -0.36111 | -1.10055 | -1.55910 | 0.06327 | eEF2K |
| 499 | 497 | ARRY334543, ARRY-334543, ARRY-543, ASLAN-001, AR-00334543, Varlitinib | 0.47788 | 0.88128 | -0.71716 | -0.67700 | 0.04363 | -2.61174 | -0.87204 | -2.84145 | -0.40065 | EGFR (ERBB1, HER1), ErbB2 (TKR1, HER2, NEU) |
| 500 | 498 | GDC-0068, RG-7440, Ipatasertib | 0.16070 | 2.27666 | -0.91860 | 2.17757 | 0.44403 | -0.01058 | 0.94363 | 1.92901 | 0.44415 | AKT1 (PKBa), AKT2 (PKBb), AKT3 (PKBg) |
| 501 | 499 | Compound 10 | -0.79975 | 0.51498 | 0.19090 | -0.35061 | 0.51973 | 0.04604 | -0.38315 | 0.10638 | -0.00986 | Erk2 (ERK, p38) |
| 502 | 500 | Compound 70 | -1.31107 | -0.03082 | 0.67078 | -0.77553 | 0.92720 | 0.54079 | 1.71304 | -0.04971 | 0.10686 | Erk2 (ERK, p38) |
| 503 | 501 | Compound4 | -0.13219 | 0.37956 | -1.14782 | 0.61962 | -0.04379 | -3.44902 | -1.94070 | -0.80202 | -0.41332 | SRC (c-SRC) |
| 504 | 502 | BRD-8899 | 1.02360 | 0.72343 | 0.09025 | 0.89231 | -0.52651 | -0.63228 | 0.20274 | -1.91088 | -0.00871 | STK33, AurB (AuroraB) |
| 505 | 503 | Toceranib, PHA-291639, SU-11654 | -1.14028 | 1.42838 | -2.68743 | -0.42840 | -0.13099 | -1.41230 | -1.61275 | -8.06837 | -0.89131 | KDR (VEGFR2, VEGFR, FLK1), PDGFRb (PDGFR, PDGFR1), FGFR1 (FLT2), FGFR2, FGFR3, FGFR4 |
| 506 | 504 | Bafetinib, CNS-9, INNO-406, NS-187 | 0.32302 | 0.54146 | 0.72967 | -0.11962 | -0.53587 | -1.81969 | -0.93872 | -0.83317 | -0.16827 | BCR, LYN, ABL1 (ABL), FYN (p59fyn) |
| 507 | 505 | LY-2606368 | 1.11730 | 3.30198 | 8.31089 | -5.70376 | -5.45188 | -11.36442 | -6.64292 | -8.06249 | -1.55370 | CHK1 |
| 508 | 506 | CZC 24832, CZC-24832 | 1.35013 | 1.07867 | 1.05033 | 0.12316 | -0.20287 | -1.65021 | 1.54138 | 0.48608 | 0.23955 | Phosphatidylinositol 3-Kinase gamma (PI3Kgamma) Inhibitor |
| 509 | 507 | ACHP | 0.10123 | -0.19958 | -0.01303 | -0.28188 | -0.37591 | -1.26550 | 0.45857 | -2.12864 | -0.23499 | IKKb (IKK2) |
| 510 | 508 | AMG 548, AMG-548 | 1.15318 | 2.18457 | 0.20014 | 0.21240 | 0.84648 | -0.92134 | 1.96333 | -0.13052 | 0.34938 | p38 MAPK, TNF-alpha Production Inhibitor |
| 511 | 509 | Compound 16 | -0.94053 | 1.32758 | 1.19822 | 0.12740 | 2.50202 | -0.96726 | -1.28157 | -3.29163 | -0.08409 | ACK (ACK1, TNK2), LCK (p56lck) |
| 512 | 510 | Pacritinib, ONX-0803, SB-1518 | -0.20626 | 1.16823 | -1.88996 | -0.96898 | -0.23322 | -1.16744 | -2.82977 | -4.73577 | -0.68904 | JAK2, CDK2/Cyclin A, CDK2/Cyclin E, FLT3 (STK1, FLK2) |
| 513 | 511 | CGI-1746, G-182 | 1.37056 | -1.47116 | -0.24970 | 0.38630 | 1.07076 | -1.48776 | 1.00743 | -3.80001 | -0.20130 | BTK (Bruton's tyrosine kinase) |
| 514 | 512 | Compound 71 | -0.79657 | -0.82430 | -0.45554 | -0.01639 | 7.21499 | -0.86762 | -5.80490 | -5.81260 | -0.46702 | DYRK1A (DYRK) |
| 515 | 513 | CO-1686, AVL-301, Rociletinib | -0.42888 | 2.12755 | 0.32302 | 1.12507 | 0.56415 | -0.61951 | -0.35632 | 0.28466 | 0.19154 | EGFR (ERBB1, HER1) Thr790Met Mutant |
| 516 | 514 | A 484954, A-484954 | 0.72034 | 0.52541 | -0.36307 | 0.46476 | 0.28101 | -1.57488 | 0.96599 | -1.20770 | -0.01193 | eEF2K |
| 517 | 515 | CRT0066854 | 0.84615 | 1.89448 | 0.75712 | 0.75743 | -0.29804 | -1.45397 | -1.41245 | -5.16209 | -0.25824 | PKCi (PRKCI) |
| 518 | 516 | Compound 11j | 0.17738 | -0.22274 | 0.20550 | -0.11593 | -0.98704 | -3.35112 | -4.20287 | -7.04904 | -0.98605 | SMG1 (ATX, LIP) |
| 519 | 517 | XL-388 | 0.28739 | 0.50276 | 0.22275 | 0.61022 | 0.57578 | 0.47050 | 1.43883 | -0.44374 | 0.23243 | mTOR Complex 1 (mTORC1) Inhibitor, mTOR Complex 2 (mTORC2) Inhibitor |
| 520 | 518 | SB-505124 | -0.72751 | -0.01369 | -0.11331 | 1.07157 | 2.31519 | 1.95070 | 2.16459 | 2.41842 | 0.57504 | TGFbR1 (ALK5), ALK4 (ACVR1B, ACTRIB), ALK7 (ACVR1C) |
| 521 | 519 | AZ-20 | 0.22762 | 0.99142 | 0.96322 | 0.43867 | 0.81765 | 1.76170 | 0.32514 | -4.12847 | 0.08861 | ATR, FRAP (MTOR, FRAP1) |
| 522 | 520 | WZ-4003 | 0.02396 | 1.50857 | -0.15607 | -0.93038 | 0.47547 | 0.70945 | 1.51299 | 1.94667 | 0.32289 | EGFR (ERBB1, HER1) Thr790Met Mutant, NuaK1 (NUAK1, ARK5, KIAA0537), NuaK2 (NUAK2, SNARK) |
| 523 | 521 | AZD-2014 | -1.75341 | 0.05927 | -0.28137 | 0.81147 | -0.02787 | -2.69002 | -1.77749 | 0.20142 | -0.34619 | mTOR Complex 1 (mTORC1) Inhibitor, mTOR Complex 2 (mTORC2) Inhibitor |
| 524 | 522 | NMS-1286937, NMS-P937 | 0.88953 | 0.89966 | 0.05959 | 1.55974 | -6.53284 | -6.78393 | -7.73228 | -8.99516 | -1.68947 | PLK1 (PLK, STPK13) |
| 525 | 523 | MPI-0479605 | -0.93118 | 0.21281 | -0.64707 | 0.65820 | -0.16896 | 0.60534 | -3.68250 | -6.32766 | -0.65211 | TTK (ESK, MPS1L1, Monopolar spindle 1 (Mps1) kinase) |
| 526 | 524 | Chk2 Inhibitor II, 339253, BML-277 | 0.78059 | -0.40610 | -0.11792 | 0.25024 | 0.11401 | -2.20121 | -0.13242 | -0.63206 | -0.14873 | CHK2 |
| 527 | 525 | Compound 11d, Eph inhibitor | 1.00943 | 0.67862 | 0.44358 | -0.30258 | -0.51620 | -1.28622 | -0.94696 | -2.38437 | -0.20961 | EphB4 |
| 528 | 526 | UNC-569A, UNC-569, Mer inhibitor | -0.83549 | -0.18630 | -0.20932 | 0.77712 | 0.09898 | -0.70870 | 0.47805 | 2.17053 | 0.10053 | MER (MERTK, c-mer) |
| 529 | 527 | AZD-9291, Mereletinib, AZD9291, Osimertinib, Tagrisso | 1.05253 | 1.13277 | 0.84095 | -0.45889 | -0.71938 | -1.06101 | -0.01662 | -3.10207 | -0.14790 | EGFR (ERBB1, HER1) Thr790Met Mutant |
| 530 | 528 | RG-7842, GDC-0994 | 0.41353 | 2.03750 | 0.57163 | 0.86460 | 2.73499 | 0.34286 | 1.29048 | 1.25857 | 0.60347 | Erk1, Erk2 (ERK, p38) |
| 531 | 529 | FR-180204 | -0.49650 | 0.42249 | 0.00445 | -0.16749 | 1.09426 | -1.09908 | -0.39944 | 0.96762 | 0.02070 | Erk1, Erk2 (ERK, p38) |
| 532 | 530 | VE-822, VX-970 | -0.07842 | 0.63138 | -1.27434 | 0.28114 | 2.62092 | 2.51925 | -2.98497 | -2.11632 | -0.02546 | ATR |
| 533 | 531 | Filgotinib, G-146034, GLPG-0634 | -0.91952 | 0.06719 | -0.89785 | 1.93377 | 0.58945 | -1.18701 | -0.37069 | 0.84278 | 0.00369 | JAK1, JAK2 |
| 534 | 532 | LY2835219, LY-2835219 | -0.12998 | -0.59467 | -0.56193 | -0.92426 | -2.55681 | -4.83493 | -5.11183 | -4.88670 | -1.24327 | CDK4/Cyclin D1, CDK6/Cyclin D3, PIM1 |
| 535 | 533 | AMG-511 | -0.24484 | 0.49765 | 1.76886 | 5.75388 | 4.55006 | 5.15593 | 6.50428 | 4.53849 | 1.80926 | Phosphatidylinositol 3-Kinase alpha (PI3Kalpha) Inhibitor |
| 536 | 534 | | 0.90288 | -0.35739 | -0.13981 | -2.36701 | -2.76361 | -5.91718 | -5.20719 | -4.86954 | -1.31417 | CDK4/Cyclin D1, FLT3 (STK1, FLK2), CDK6/Cyclin D3 |
| 537 | 535 | AZ5104 | 0.63071 | 1.34414 | 0.03213 | 0.59833 | 0.71740 | -3.58818 | -1.55319 | -5.35288 | -0.45488 | |
| 538 | 536 | CEP-37440 | 0.37902 | 0.88486 | -0.05906 | -0.32793 | 0.31325 | 0.96248 | 4.27578 | 3.51537 | 0.63072 | ALK, FAK (FAK1) |
| 539 | 537 | GS-9973, Entospletinib | 2.11503 | 0.70674 | -0.67396 | 0.46625 | 0.18599 | 0.59371 | -1.21958 | -1.40257 | 0.04894 | SYK |
| 540 | 538 | SCH-772984 | 0.39103 | 1.62600 | 1.26931 | 2.16955 | 1.49730 | 1.33072 | -1.44646 | -4.78338 | 0.13029 | Erk1, Erk2 (ERK, p38) |
| 541 | 539 | PND-1186, VS-4718 | 0.17171 | 1.34313 | 1.33216 | -0.47837 | -0.59840 | -0.95685 | -0.73422 | -1.43751 | -0.08616 | FAK (FAK1) |
| 542 | 540 | JNK-IN-8 | -0.68406 | -0.78021 | -0.63951 | -0.19283 | 0.37162 | -1.28402 | 0.04466 | 0.37354 | -0.17702 | JNK1 (JNK), JNK3 |
| 543 | 541 | MPS1-IN-2 | 1.26483 | 0.10034 | -0.79895 | -0.43470 | 0.31453 | -2.12882 | -3.13289 | -4.81991 | -0.61117 | TTK (ESK, MPS1L1, Monopolar spindle 1 (Mps1) kinase) |
| 544 | 542 | WH-4-023, KIN001-112, KIN112 | 0.14600 | 0.04091 | -0.38007 | -2.00187 | 0.34710 | -0.71119 | 0.21493 | 1.09216 | -0.07941 | LCK (p56lck), SRC |
| 545 | 543 | GNE-0877 | -0.51613 | 0.40657 | -0.33410 | 0.85968 | -0.13486 | -0.01435 | -0.80584 | -0.06621 | -0.03839 | LRRK2 (Leucine-Rich Repeat Kinase 2, Dardarin) |
| 546 | 544 | TAK-632 | -0.50751 | 1.46743 | -0.46276 | -1.59764 | -1.75992 | -2.68982 | -1.65563 | -1.95244 | -0.58090 | BRAF (B-raf), RAF1 (c-Raf) |
| 547 | 545 | MRT 67307 dihydrochloride C26H36N6O2.2HCl | 0.19228 | 0.70856 | 1.56684 | 0.86324 | 0.47128 | 0.84590 | 1.00239 | -3.62514 | 0.12847 | SIK (MSK), TBK1, MARK1 (MARK), MARK2 (EMK-1, PAR-1), MARK3 (PAR1A), MARK4 (MARKL1), IKKe, NuaK1 (NUAK1, ARK5, KIAA0537) |
| 548 | 546 | LDN-209929 dihydrochloride | -1.33012 | 0.52302 | -0.90254 | -2.18463 | -1.20912 | -2.25013 | -0.44472 | -1.64510 | -0.59898 | Haspin (GSG2) |
| 549 | 547 | FRAX-486 | -0.41128 | 1.30433 | 0.47200 | 0.71679 | 3.84288 | 2.38315 | -1.15853 | -2.06550 | 0.32246 | PAK1 (PAKa) |
| 550 | 548 | SR-3576, SR 3576 | 1.45056 | -0.58763 | -0.49297 | 0.48257 | -0.18024 | -1.08316 | -0.97030 | -2.26467 | -0.23125 | JNK3 |
| 551 | 549 | AVL-292, CC-292, Spebrutinib | 0.56531 | 0.09350 | -2.27475 | -1.67168 | -4.33988 | -2.52711 | -6.45535 | -4.49030 | -1.33836 | ITK (EMT, LYK), JAK3, EGFR (ERBB1, HER1), BTK (Bruton's tyrosine kinase) |
| 552 | 550 | CNX-774 | 0.20925 | -0.13481 | 0.00370 | 1.85625 | 0.67966 | -0.10452 | -0.02253 | 0.42688 | 0.18482 | BTK (Bruton's tyrosine kinase) |
| 553 | 551 | PRT-2607, BIIB-057, P505-15, PRT-062607, | 0.38517 | 0.18799 | 0.19043 | 0.04146 | 1.54232 | 0.37510 | 1.83012 | 2.25050 | 0.43151 | SYK |
| 554 | 552 | SAR-131675 | -0.31537 | 1.21018 | 0.33870 | -0.34334 | 0.31650 | -0.50435 | 1.27614 | -1.93889 | 0.00251 | FLT4 (VEGFR3) |
| 555 | 553 | G-749 | 0.91141 | -0.17851 | 0.74272 | 0.58515 | -0.01754 | 1.06263 | 0.20334 | -0.40094 | 0.18447 | FLT3 (STK1, FLK2) |
| 556 | 554 | BMS-863233, XL-413 | 0.11936 | 0.90395 | -1.66359 | -0.00292 | 1.16498 | -1.87387 | 0.03722 | 2.93552 | 0.10280 | CDC7 |
| 557 | 555 | Losmapimod, 856553, GSK-AHAB, GW-856553, GW-856553X, SB-856553 | -1.81818 | 1.47695 | -0.33357 | 0.18183 | 0.16867 | -0.35364 | -0.11791 | 0.31393 | -0.03057 | p38a |
| 558 | 556 | Encorafenib, LGX-818, NVP-LGX-818 | 0.65219 | 0.04203 | -0.00684 | 1.01256 | 1.33524 | -0.06868 | -3.59153 | -1.72753 | -0.14922 | BRAF (B-raf) |
| 559 | 557 | Binimetinib, ARRY-162, ARRY-438162, MEK-162 | 0.15119 | -0.05768 | 1.27216 | 1.13628 | 3.42148 | -0.43693 | 1.76416 | 1.29244 | 0.54188 | MAP2K1 (MEK1), MAP2K2 (MEK2) |
| 560 | 558 | Ketohexokinase Inhibitor, JNJ-42065426 | 1.15520 | 1.08148 | 0.12547 | 0.39147 | 0.73615 | 0.89283 | -0.16247 | -0.51170 | 0.23522 | KHK (Ketohexokinase) |
| 561 | 559 | SB-706504 | 0.32460 | -0.55394 | -0.35710 | 0.07815 | 0.23189 | -0.40009 | 0.31998 | 0.54066 | 0.01168 | p38 MAPK |
| 562 | 560 | AL 8697 | 0.16771 | 0.22323 | 0.31115 | -0.54555 | 1.35835 | -0.19143 | 1.01119 | 0.93238 | 0.20722 | p38a, TNF-alpha Production Inhibitor, RET |
| 563 | 561 | 17-(Allylamino) Geldanamycin, 17-AAG, Tanespimycin, 17-(Allylamino)-17-demethoxygeldanamycin, BMS-722782, CNF-101, CP-127374, KOS-953, NSC-330507, NSC-704057 | -0.51116 | 1.37981 | 0.40296 | 0.45487 | 1.15748 | 3.70276 | -0.25506 | -5.26864 | 0.06743 | Heat Shock Protein 90 (hsp90) Inhibitor, CHK1 Expression Inhibitor |
| 564 | 562 | Bortezomib, Velcade, MG-341, PS-341, DPBA, JNJ-26866138, LDP-341, MLN-341, NSC-681239 | 0.35436 | -0.99125 | 0.74472 | -4.87992 | -9.54564 | -11.74272 | -5.45735 | -5.62357 | -2.35583 | Proteasome inhibitor, Caspase 3 activators, NF-kappaB (NFKB) Activation Inhibitor |
| 565 | 563 | KRIBB3 | -0.39004 | -0.10054 | 0.53302 | 0.62220 | 1.57174 | -2.36226 | -5.70606 | -6.20580 | -0.76354 | Heat Shock Protein 27 (hsp27) Inhibitor, Tubulin polymerization inhibitor |
| 566 | 564 | SB 334867, SB-334867 | 1.24985 | 1.00194 | -0.38468 | 0.69756 | 0.20244 | -1.22320 | -0.39524 | -0.33283 | 0.05175 | Orexin (OX-1) receptor antagonist |
| 567 | 565 | SB-408124 | 0.66903 | -0.09394 | -0.08297 | 0.86501 | 0.54732 | -1.60398 | -1.44937 | 0.26906 | -0.05581 | Orexin (OX-1) receptor antagonist |
| 568 | 566 | AZD-2281, Olaparib, KU0059436, KU-0059436 | -0.37397 | 0.11021 | 1.06294 | 1.46375 | 0.72997 | 0.66671 | 1.50834 | 1.38555 | 0.41568 | Poly(ADP-ribose)polymerase-1 (PARP-1) Inhibitor, Poly(ADP-ribose)polymerase-2 (PARP-2) Inhibitor |
| 569 | 567 | ABT-888, Veliparib, NSC-737664 | 0.86874 | -0.41581 | 0.70609 | 0.27984 | 0.11175 | 0.16133 | -0.15675 | 0.65187 | 0.13999 | Poly(ADP-ribose)polymerase-1 (PARP-1) Inhibitor, Poly(ADP-ribose)polymerase-2 (PARP-2) Inhibitor |
| 570 | 568 | Ouabain, g-strophanthin, BNC-1 | 0.01378 | 0.91311 | 0.08913 | -4.79979 | -5.87656 | -6.47711 | -5.80851 | -7.11486 | -1.84329 | Na+, K+ ATPase inhibitor |
| 571 | 569 | 4-HQN | 0.57835 | 0.38606 | 0.00406 | 0.34219 | -0.06245 | -0.06903 | -0.41394 | 0.35787 | 0.07124 | Poly(ADP-ribose)polymerase-1 (PARP-1) Inhibitor, Poly(ADP-ribose)polymerase-2 (PARP-2) Inhibitor |
| 572 | 570 | NU 1025, NU-1025 | -0.25943 | 1.42801 | -0.82687 | 1.45740 | 0.36276 | 0.20912 | -0.57075 | -0.43249 | 0.08676 | Poly(ADP-ribose)polymerase-1 (PARP-1) Inhibitor, DNA-Dependent Protein Kinase (DNA-PK) Inhibitor |
| 573 | 571 | 3-Aminobenzamide, 3-ABA, INO-1001 | -1.26475 | 2.34610 | -0.70909 | 1.21595 | 0.23848 | -0.75646 | -0.34004 | 0.37889 | 0.07035 | Poly(ADP-ribose)polymerase-1 (PARP-1) Inhibitor, Poly(ADP-ribose)polymerase-2 (PARP-2) Inhibitor, NAD+ ADP-Ribosyltransferase Inhibitor |
| 574 | 572 | DR2313, PARP Inhibitor XI | -0.62032 | 1.22096 | 0.26261 | -0.44092 | 0.96668 | -0.47552 | -0.67500 | 0.08655 | 0.02062 | Poly(ADP-ribose)polymerase-1 (PARP-1) Inhibitor, Poly(ADP-ribose)polymerase-2 (PARP-2) Inhibitor, NAD+ ADP-Ribosyltransferase Inhibitor |
| 575 | 573 | PAC 1, PAC-1 | 0.41058 | 0.82137 | 0.51708 | 1.00708 | 0.37256 | -1.27388 | -0.24045 | 0.81487 | 0.15408 | Procaspase 3 Activator |
| 576 | 574 | INH2BP, AntiPARP-1 | -0.04411 | 0.90713 | 0.31767 | 0.70104 | -0.01630 | -0.98086 | -0.26376 | -0.77736 | -0.00993 | Poly(ADP-ribose)polymerase-1 (PARP-1) Inhibitor, Poly(ADP-ribose)polymerase-2 (PARP-2) Inhibitor, NAD+ ADP-Ribosyltransferase Inhibitor |
| 577 | 575 | DPQ | 1.38810 | 1.31465 | -0.14923 | 0.60403 | 1.19223 | -1.10419 | -0.12676 | -1.13940 | 0.12555 | Poly(ADP-ribose)polymerase-1 (PARP-1) Inhibitor, Poly(ADP-ribose)polymerase-2 (PARP-2) Inhibitor, NAD+ ADP-Ribosyltransferase Inhibitor |
| 578 | 576 | 1,5-Isoquinolinediol, IQD | -0.67945 | -0.39807 | -1.68284 | 1.28537 | -0.10403 | 0.03550 | 0.22654 | -0.30522 | -0.10289 | Poly(ADP-ribose)polymerase-1 (PARP-1) Inhibitor, Poly(ADP-ribose)polymerase-2 (PARP-2) Inhibitor, NAD+ ADP-Ribosyltransferase Inhibitor |
| 579 | 577 | PJ-34 hydrochloride hydrate | -0.47187 | -1.22707 | -0.11773 | -0.07415 | -0.78613 | -2.58735 | -2.98247 | -3.53908 | -0.74756 | Poly(ADP-ribose)polymerase-1 (PARP-1) Inhibitor, Poly(ADP-ribose)polymerase-2 (PARP-2) Inhibitor, NAD+ ADP-Ribosyltransferase Inhibitor |
| 580 | 578 | EB-47 | -0.29503 | 1.43657 | 1.53355 | -0.20941 | -0.52317 | 1.39322 | -0.03594 | 2.48970 | 0.36722 | Poly(ADP-ribose)polymerase-1 (PARP-1) Inhibitor |
| 581 | 579 | 5-AIQ hydrochloride | 0.03259 | 1.06789 | 0.62734 | 0.80592 | -1.15738 | -0.12381 | -1.51919 | -1.18631 | -0.09216 | Poly(ADP-ribose)polymerase-1 (PARP-1) Inhibitor, Poly(ADP-ribose)polymerase-2 (PARP-2) Inhibitor, NAD+ ADP-Ribosyltransferase Inhibitor |
| 582 | 580 | TIQ-A, UPF-734 | 0.92734 | 1.65592 | 0.39719 | -0.58622 | -0.15230 | -1.60976 | -1.52954 | -0.84860 | -0.11074 | Poly(ADP-ribose)polymerase-1 (PARP-1) Inhibitor |
| 583 | 581 | 4-ANI, PARP inhibitor V, ANI | 0.06264 | 1.74352 | -0.24488 | -0.55602 | 0.55006 | -0.77205 | 0.42380 | -3.28032 | -0.13150 | PARP inhibitor |
| 584 | 582 | YM-155, YM155, Sepantronium bromide | 1.59735 | 3.96258 | 8.70981 | 7.16492 | -8.68574 | -6.90119 | -8.25801 | -8.53437 | -0.69421 | BIRC5 Expression Inhibitor, Survavin inhibitor |
| 585 | 583 | Loperamide hydrochloride, ADL-2-1294, R-18553, Dissenten, Fortasec, Imodium, Lopemin, Salvacolina | 1.33515 | 1.40807 | -0.53698 | -0.68841 | -0.04042 | 0.38379 | -0.46845 | -0.38746 | 0.06376 | mu-Opioid Agonist |
| 586 | 584 | Anisomycin, Flagecidin, NSC-76712 | 0.04097 | 0.54029 | 1.51663 | 4.91103 | 1.55964 | -7.00077 | -2.57203 | -7.91390 | -0.56567 | Protein synthesis inhibitor |
| 587 | 585 | Lopinavir, ABT-378 | -1.73589 | 1.15191 | 0.31400 | 0.09785 | 0.97204 | -1.62167 | -0.62069 | -1.14194 | -0.16392 | HIV Protease Inhibitor |
| 588 | 586 | IWP-3 | 0.46200 | -0.30434 | 1.36667 | 0.07242 | 1.25388 | -0.49141 | -0.18091 | -1.67199 | 0.03212 | Wnt inhibitor |
| 589 | 587 | Cyclosporin A, Ciclosporin A, ATH-002; CB-01-09 MMX; CSA; CyA; DE-076; Nova-22007; OL-27400; OLO-400; ST-603; Sang-2000; Sang-35 | -0.06537 | 0.29065 | 0.11220 | 0.53377 | 1.05719 | 0.83917 | 1.44047 | 2.87257 | 0.44912 | Cytochrome P450 CYP3A4 Inhibitors; Cyclophilin D Inhibitors; INS Expression Inhibitors; Calcineurin (PP-2B) Inhibitors |
| 590 | 588 | Cyclopamine, 11-deoxojervine | -0.44032 | 0.48222 | -0.73855 | 0.71938 | -1.02092 | -0.89246 | -0.47028 | 0.05658 | -0.14616 | Smoothened (Smo) Receptor Antagonist, Hedgehog inhibitor, Hh inhibitor |
| 591 | 589 | Bexarotene, Targretin, Targretyn, Targrexin, LGD-1069, LG 1069, LG100069, LG-100069, SR-11247 | 0.88333 | 1.48171 | -0.21283 | -1.69862 | -3.66411 | -7.38720 | -10.14258 | -12.16932 | -2.08742 | Retinoid RXR Agonists; Extracellular-Regulated Kinase (ERK) Inhibitors; SAPK1 (JNK) Inhibitors; ABCA1 Expression Enhancers |
| 592 | 590 | Docetaxel, Taxotere, Docetaxol, ANX-514; EmDOC; ML-061; NSC-628503; RP-56976; SD-009; SDP-014; SID-530; SYP-0704A; XRP-6976L | 0.14245 | 5.22362 | -0.80557 | -4.43166 | -7.71595 | -5.73739 | -8.13038 | -5.78531 | -1.72781 | Microtubule-Stabilizing Agents; Antimitotic Drugs |
| 593 | 591 | Paclitaxel, Taxol | 0.43727 | 1.75846 | 6.22316 | 0.50813 | -3.85679 | -9.52512 | -9.59257 | -7.44159 | -1.36302 | P-Glycoprotein (MDR-1; ABCB1) Inhibitors; Angiogenesis Inhibitors; Microtubule-Stabilizing Agents; Antimitotic Drugs |

| | A | B | C | D | E | F | G | H | I | J | K | L |
|---|---|---|---|---|---|---|---|---|---|---|---|---|
| 594 | 592 | CAY10594, VU-0155072 | -0.60033 | 0.85468 | 0.08287 | 0.11261 | 0.38004 | -0.02628 | -0.26890 | 0.12589 | 0.04190 | Phospholipase D2 (PLD2) Inhibitor |
| 595 | 593 | ABT-263, Navitoclax, ABT263, Navitodax, RG-7433 | -0.82560 | 1.35610 | 0.46021 | 1.44482 | -0.97564 | -7.28049 | -6.80350 | -7.88766 | -1.30103 | Bcl-2, Bcl-XL, Bcl-w inhibitors |
| 596 | 594 | CUR-691; GDC-0449; GDC-449 ; Hh-Antag691; HhAntag; R-3616; RG-3616, Vismodegib, Erivedge | 0.44977 | 0.82697 | 0.00564 | -0.30678 | 0.91154 | -0.34904 | 1.23933 | 1.18372 | 0.25125 | P-Glycoprotein (MDR-1; ABCB1) Inhibitor; Hedgehog (Hh) Signaling Inhibitor; Breast Cancer-Resistant Protein (BCRP; ABCG2) Inhibitor; Smoothened (Smo) Receptor Antagonist |
| 597 | 595 | Raltegravir, MK-0518, Isentress | -0.98779 | 0.63829 | -0.80112 | 1.39091 | 0.56420 | -1.43151 | 1.62318 | 0.54405 | 0.09769 | HIV integrase inhibitor |
| 598 | 596 | Odanacatib, MK-0822 | 0.16213 | 0.94839 | -0.68951 | 0.75952 | 0.09685 | -0.96668 | 0.51043 | -0.73457 | 0.00549 | Cathepsin K inhibitor |
| 599 | 597 | Rofecoxib, Vioxx, Ceoxx, Ceeoxx, MK-0966, MK-966 | 0.19716 | 1.07202 | 0.56009 | 0.38958 | 1.38009 | 0.02020 | 0.74523 | -0.46487 | 0.24734 | Cyclooxygenase (COX-2) inhibitor |
| 600 | 598 | D07001-F4; Inno-D07001; LY-188011; dFdC, Gemcitabine hydrochloride, Gemzar | 1.57739 | 0.32533 | -9.28124 | -7.86511 | -8.78731 | -8.99596 | -7.20444 | -7.30576 | -3.01522 | Ribonucleoside-Diphosphate Reductase Inhibitor; Pyrimidine Antagonist |
| 601 | 599 | 17-DMAG, Alvespimycin, KOS-1022, NSC 07545, 17-(Dimethylaminoethylamino)-17-demethoxygeldanamycin Hydrochloride, BMS-826476 | 0.00269 | 0.78414 | 0.62875 | 3.33798 | -0.84859 | -10.23558 | -9.68517 | -7.43915 | -1.48772 | Heat Shock Protein 90 (hsp90) Inhibitor |
| 602 | 600 | Ispinesib, SB-715992, CK0238273, CK-0238273, 715992 | 0.64485 | 1.58983 | -7.36130 | -8.03486 | -6.30954 | -8.85895 | -7.42233 | -6.32738 | -2.66906 | Kinesin-Like Spindle Protein (KSP, Eg5) Inhibitor; Antimitotic Drug |
| 603 | 601 | Cercosporin from Cercospora hayii | 0.81303 | 0.50300 | 0.20728 | 0.65475 | -0.25793 | -0.45379 | 3.63453 | 0.16976 | 0.33431 | Phytotoxin, Protein Kinase C (PKC) Inhibitor |
| 604 | 602 | Metformin hydrochloride, Glucophage, Riomet, Diabex, ADX-155; La-6023; SMP-862, Fortamet, Glumetza; Glycoran; Melbin; MetControl, Metgluco, Siofor | -0.48938 | 0.28166 | 0.18821 | -0.42757 | 1.17447 | 0.05725 | -0.09034 | 0.82144 | 0.09614 | Insulin Sensitizers; AMP-Activated Protein Kinase (AMPK) Activator, Antihyperglycemic agent |
| 605 | 603 | Trichostatin A, TSA, Tricostatin A | -0.11481 | -0.05198 | 0.47196 | -0.34071 | 0.18891 | -0.22855 | 0.76619 | 0.76940 | 0.09263 | HDAC inhibitor, CDKN1A Expression Enhancer, ID1 Expression Inhibitor |
| 606 | 604 | IWR-1 | -0.62383 | 0.71044 | 0.40997 | -0.96679 | -0.27426 | -0.38877 | 0.06944 | 0.36577 | -0.04428 | Tankyrase 2 (TNKS2) Inhibitor, Tankyrase 1 (TNKS1) Inhibitor |
| 607 | 605 | XAV-939, NVP-XAV-939 | -0.07623 | 0.03802 | -2.93763 | -2.26163 | -3.38507 | -5.52629 | -4.83810 | -5.21309 | -1.53497 | Tankyrase 2 (TNKS2) Inhibitor, Tankyrase 1 (TNKS1) Inhibitor |
| 608 | 606 | 2,3-Dibromo-6-hydroxy-5-methoxy-benzaldehyde | 0.66742 | 0.58306 | 0.13456 | 1.73166 | 1.92228 | -0.39977 | 0.82786 | -0.47867 | 0.31641 | IRE1 inhibitor |
| 609 | 607 | TW-37, TW37 | 0.31445 | 1.45447 | -0.03424 | 0.12574 | 0.24531 | -1.06898 | -1.41151 | -5.22373 | -0.35510 | Bcl-2, Bcl-XL, Mcl-1 inhibitor |
| 610 | 608 | ABT 737, ABT-737 | 1.38366 | 0.58106 | 0.14947 | 1.80582 | 0.36928 | -0.35053 | -2.75979 | -5.33485 | -0.26360 | Bcl-2, Bcl-XL, Bcl-w inhibitors |
| 611 | 609 | GX15-070 mesylate, GX-015-070 mesylate, Obatoclax Mesylate, CEP-41601 | -1.05318 | 0.53217 | -0.15259 | -0.03032 | 0.01496 | -4.93058 | -9.23906 | -7.61668 | -1.42558 | Bcl-2, Bcl-XL, Mcl-1 inhibitor |
| 612 | 610 | CI 994, CI-994, Tacedinaline, Acetyldinaline, Go 5549, Goe 5549, PD 123654, PD-123654 | 0.55703 | 0.22033 | 1.23206 | 1.04011 | 1.26954 | 1.79349 | 1.19547 | 1.08870 | 0.53259 | Histone Deacetylase (HDAC) Inhibitor |
| 613 | 611 | Deferoxamine mesylate salt, Desferrioxamine mesylate salt, DFOM, Ba-33112 | 0.01075 | 1.09047 | -0.05122 | -0.02385 | 0.25988 | -0.85001 | -1.02999 | -1.23796 | -0.11620 | APP Expression Inhibitor, Antimetabolite, Chelating Agent |
| 614 | 612 | Deferiprone, Ferriprox | 0.80154 | 1.29377 | 0.01922 | 0.79918 | 0.28946 | -0.31931 | 0.37749 | -1.57911 | 0.10670 | Chelating Agent |
| 615 | 613 | Pyridoxal isonicotinoyl hydrazone, PIH, NSC-77674 | -0.18604 | 0.91773 | 1.49346 | -0.34903 | 0.59546 | -0.82204 | -0.26318 | -0.71886 | 0.04234 | Chelating Agent |
| 616 | 614 | NIH, 311, NSC-51355 | 0.26930 | 0.00579 | -0.60187 | -0.65485 | -1.08961 | -3.33645 | -3.87061 | -1.23794 | -0.66703 | Chelating Agent |
| 617 | 615 | PCIH | 0.21175 | 0.86460 | 0.72324 | 1.55645 | -0.07505 | -0.90565 | -1.52178 | -1.72876 | -0.05551 | Chelating Agent |
| 618 | 616 | PBT-1, Clioquinol; Iodochlorhydroxyquin, Chinoform; Linolasept; Vioform | -0.31334 | 0.22158 | 0.55099 | -0.21267 | 0.21940 | -1.10661 | 0.05058 | -0.85692 | -0.09178 | Chelating Agents; Antiamyloidogenic Agents; Carbonic Anhydrase Inhibitors |
| 619 | 617 | Deferasirox, Exjade, CGP-72670, ICL-670 | -0.65727 | 1.53580 | -0.43514 | 0.02858 | 1.30057 | -0.50817 | 0.46787 | 0.76879 | 0.15864 | Chelating Agent |
| 620 | 618 | Propiconazole | 1.58908 | 0.55399 | 0.47004 | -0.36851 | -0.15778 | -0.01370 | -1.15417 | 1.01990 | 0.12298 | Fungicide |
| 621 | 619 | Fluconazole, Diflucan, Trican, Alfumet, Candivast | -0.39360 | 1.43339 | -0.59978 | -0.31374 | 1.25661 | -0.96196 | -0.52311 | 0.07712 | -0.00159 | Antifungal, Cytochrome P450 CYP3A4 Inhibitor, Cytochrome P450 CYP1A2 Inhibitor, Cytochrome P450 CYP2C9 Inhibitor |
| 622 | 620 | Econazole nitrate salt, R-14827, SQ-13050, Ecostatin, Epi-Pevaryl, Gino-Pevaryl, Spectazole | -0.00253 | 1.56372 | -1.07012 | 0.44456 | 1.30326 | 0.90918 | 2.27680 | 1.38892 | 0.43219 | Antifungal |
| 623 | 621 | L-741,626, L-741626 | 0.19626 | 0.36750 | 0.11026 | 0.09901 | 0.73092 | -0.20342 | 0.04164 | 2.15872 | 0.22206 | Dopamine D2 Antagonist |
| 624 | 622 | MK-0431; MK-431; ONO-5435, Sitagliptin; Sitagliptin phosphate, Glactiv; Januvia; Tesavel; Xelevia | 0.58857 | 0.06037 | -0.28278 | -0.32861 | 0.37479 | -0.76548 | 1.26460 | -0.12478 | 0.04990 | Dipeptidyl Peptidase IV (CD26; DPP-IV; DP-IV) Inhibitor; HMG-CoA Reductase Inhibitor; TNFSF6 Expression Inhibitor |
| 625 | 623 | BC-105, Pizotifen, Pizotyline, Sandomigran, Sanmigran, Sanomigran | -0.81468 | 0.02816 | -0.52239 | 0.35980 | -0.12484 | -1.25742 | -0.03405 | -0.34084 | -0.17165 | Serotonin antagonist |
| 626 | 624 | MS-275, Entinostat, SNDX-275, MS-27-275, NSC-706995 | 0.08106 | 1.16963 | 0.47907 | -0.69907 | -0.09222 | 0.15347 | 1.16643 | -0.78184 | 0.09365 | Histone Deacetylase (HDAC) Inhibitor |
| 627 | 625 | Calmidazolium chloride, R 24571 | -0.31904 | -0.01457 | -0.16119 | 0.46901 | 1.13191 | -0.37879 | 0.85037 | 0.36415 | 0.12317 | Calmodulin antagonist |
| 628 | 626 | Tebuconazole | -0.43960 | 0.68562 | -0.08430 | 1.50303 | 0.85996 | 0.65047 | 3.21582 | 4.91788 | 0.71731 | Antifungal |
| 629 | 627 | Ketotifen fumarate salt, Zaditen, Zaditor | 0.59423 | 0.98247 | -0.24758 | -0.47638 | 1.25334 | -0.60447 | 0.34159 | 0.21121 | 0.13031 | Histamine H1 Receptor Ligand |
| 630 | 628 | Butoconazole, RS-35887, Femstat, Gynazole-1, Gynomyk | 0.44835 | 0.20439 | -0.48885 | 0.97776 | 3.69531 | 3.02707 | -0.23031 | -1.16280 | 0.41044 | Antifungal |
| 631 | 629 | Voriconazole, UK-109496, Vfend | -1.32982 | 0.52914 | -0.52940 | 0.57509 | 1.47381 | -0.30486 | -0.52528 | 0.34422 | 0.01477 | Antifungal, Lanosterol 14alpha-demethylase Inhibitor |
| 632 | 630 | ARQ-621 | -0.10917 | 1.26456 | -0.14260 | 1.31694 | -0.81162 | 3.65624 | 18.15572 | -6.04025 | 1.09667 | Kinesin-Like Spindle Protein (KSP, Eg5) Inhibitor |
| 633 | 631 | Sertaconazole nitrate, Dermofix, Dermoseptic, Ertaczo, Extens, Fisderm, Ginedermofix, Onabet, Zalain | 0.57963 | 0.40952 | 0.60106 | 1.18921 | -0.12031 | -0.83653 | 1.32500 | 2.62224 | 0.36597 | Antifungal |
| 634 | 632 | FTY720, FTY-720; FTY-720A; TDI-132, Fingolimod, Gilenia, Gilenya; Imusera | -0.23524 | -1.25867 | -0.06445 | 0.04601 | 0.62471 | -2.19060 | 1.16580 | 1.16620 | -0.04733 | Lysophospholipid edg8 (S1P5) Receptor Agonist; Lysophospholipid edg6 (S1P4) Receptor Agonist; Sphingosine Kinase 1 (SphK1) Inhibitor; Apoptosis Inducer; Lysophospholipid edg3 (S1P3) Receptor Agonist; Lysophospholipid edg1 (S1P1) Receptor Agonist |
| 635 | 633 | JTE-013 | 0.99775 | 0.15070 | -1.19314 | 0.25440 | -0.38142 | -2.62840 | -1.51403 | -2.98075 | -0.46271 | Lysophospholipid edg5 (S1P2) Receptor Antagonist |
| 636 | 634 | AGK2, AGK-2 | -0.38404 | 0.73000 | 0.10308 | 0.21635 | 1.09041 | -0.04338 | 0.33324 | 1.37784 | 0.21715 | Histone Deacetylase SIRT2 Inhibitor |
| 637 | 635 | IWR-1-exo | 2.20644 | 0.71758 | 0.01241 | -0.79823 | -0.85337 | 0.95373 | -0.90171 | 0.06532 | 0.08894 | Tankyrase 2 (TNKS2) Inhibitor, Tankyrase 1 (TNKS1) Inhibitor |
| 638 | 636 | IWR-1-endo | -0.51347 | 0.40399 | 1.30525 | 0.81775 | 0.60729 | -0.82732 | 0.34013 | -1.58116 | 0.03504 | Tankyrase 2 (TNKS2) Inhibitor, Tankyrase 1 (TNKS1) Inhibitor |
| 639 | 637 | Nutlin-3 | 1.12298 | 1.37758 | -0.26281 | -0.70149 | -0.09391 | 1.08431 | 0.09169 | -0.00691 | 0.16564 | MDM2 (hdm2) Inhibitor |
| 640 | 638 | 6-Anilinoquinoline-5,8-quinone, LY-83,583, LY-83583 | 0.03623 | -0.39485 | -0.40143 | 0.74179 | -0.19092 | 0.16498 | 5.17929 | 7.50111 | 0.80150 | Guanylate cyclase and cGMP production inhibitor, Leukotriene Synthesis Inhibitor |
| 641 | 639 | IWP-2 | -0.33232 | -0.07814 | 1.42143 | 0.08813 | 1.96552 | -0.30500 | -0.51720 | -0.90634 | 0.08475 | Wnt Signaling Inhibitor |
| 642 | 640 | IWP-4 | 0.28578 | 0.65293 | -0.49862 | -0.50109 | 0.93891 | -0.07384 | -0.12846 | 1.19659 | 0.11875 | Wnt Signaling Inhibitor |
| 643 | 641 | PARP Inhibitor XII, 3-(4-Chlorophenyl)quinoxaline-5-carboxamide, 334876 | -0.97949 | -1.48581 | -2.26275 | -0.99180 | -1.44194 | -2.52602 | -2.74718 | -2.61556 | -0.95464 | Poly(ADP-ribose)polymerase-2 (PARP-2) Inhibitor, Poly(ADP-ribose)polymerase-1 (PARP-1) Inhibitor |
| 644 | 642 | L-685,458, L685458, L-685458 | -0.34448 | 0.59576 | 0.12523 | -0.42489 | -0.07803 | 0.09383 | 0.78629 | -0.47431 | 0.01772 | gamma-Secretase inhibitor |
| 645 | 643 | STX-0119 | 0.12354 | -0.34559 | 0.07637 | -0.26518 | 0.15478 | -0.91442 | 2.70719 | -0.17395 | 0.08644 | STAT3 inhibitor |
| 646 | 644 | UPF 1069, UPF-1069 | -0.61171 | 0.21446 | -0.95589 | -0.71837 | 0.96453 | -0.97654 | -0.21586 | 0.69588 | -0.10171 | Poly(ADP-ribose)polymerase-2 (PARP-2) Inhibitor, Poly(ADP-ribose)polymerase-1 (PARP-1) Inhibitor |
| 647 | 645 | BYK 204165, BYK-204165 | -0.37458 | -0.68896 | -0.69179 | 3.70105 | 0.16915 | 0.78666 | 0.01365 | -0.76745 | 0.13623 | Poly(ADP-ribose)polymerase-2 (PARP-2) Inhibitor, Poly(ADP-ribose)polymerase-1 (PARP-1) Inhibitor |
| 648 | 646 | Cilostazol, OPC-13013, OPC-21, Pletaal, Pletal | 1.89788 | 0.91044 | 0.31726 | 0.41157 | -2.40945 | -1.23358 | 0.90444 | 1.51093 | 0.14649 | Phosphodiesterase III A (PDE3A) inhibitor |
| 649 | 647 | NCG30335, NCGC00031955 | 0.17865 | -0.80636 | -0.81955 | 0.08656 | -1.55684 | -0.49981 | 1.47139 | 1.69717 | -0.01578 | PKM2 activator |
| 650 | 648 | NCG31955 | -0.93812 | 0.26548 | -0.75707 | 1.01213 | -0.37293 | 0.50707 | -0.07423 | 0.66870 | 0.01973 | PKM2 activator |
| 651 | 649 | Carboplatin, cis-Diammine(1,1-cyclobutanedicarboxylato) platinum, Ercar, Paraplatin, CBDCA, JM-8, | -0.50071 | 0.68063 | -0.92978 | -0.16777 | -1.44869 | -0.77626 | 1.03292 | -0.90365 | -0.19113 | DNA alkylating agent, NAD+ ADP-Ribosyltransferase (poly(ADP-ribose)polymerase; PARP) Inhibitors |
| 652 | 650 | Irinotecan hydrochloride; Camptosar; DQ-2805; Campto; U-101440E; NSC-616348; CPT-11; Topotecin | -0.05107 | -0.56087 | 0.85000 | 3.28474 | 3.25612 | 3.82842 | 3.64146 | -3.01613 | 0.71247 | DNA Topoisomerase I Inhibitors |
| 653 | 651 | Pemetrexed disodium heptahydrate, Alimta, LY-231514, MTA, Rolazar, Tifolar | 1.00178 | -0.47819 | -1.36330 | 0.67768 | -4.86709 | -7.34096 | -5.29661 | -3.68981 | -1.35461 | Antimetabolites; Apoptosis Inducers; Dihydrofolate Reductase (DHFR) Inhibitors; Glycinamide Ribonucleotide Formyltransferase (GARTFase) Inhibitors; Thymidylate Synthase Inhibitors |
| 654 | 652 | Lenalidomide, CC-5013, Revlimid, CDC-501, CDC-5013, ENMD-0997, IMID-5013, IMiD3, Revamid, Revimid | -0.30895 | -0.22638 | 0.48915 | 0.02389 | 0.53634 | 4.04717 | 2.71126 | 2.75184 | 0.63583 | Angiogenesis Inhibitors; TNF-alpha Production Inhibitors |
| 655 | 653 | Vorinostat, SAHA, Suberoylanilide Hydroxamic Acid, Zolinza, MK-0683, Suberanilohydroxamic Acid, NSC-701852, SW-064652 | -1.78014 | 0.74704 | 0.09051 | 2.54377 | 1.28931 | -0.25140 | -2.48302 | -8.20876 | -0.51077 | Histone Deacetylase (HDAC) Inhibitor |
| 656 | 654 | TM30089, TM-30089, CAY-10471 | 1.15174 | 1.86227 | -0.41545 | 1.23424 | -0.38019 | 0.80375 | 0.81919 | -0.40415 | 0.29630 | CRTH2 receptor antagonist |
| 657 | 655 | NVP-AUY922, AUY922, VER-52296 | -0.87613 | 0.02892 | -9.50049 | -8.22240 | -4.97992 | -0.98404 | -5.59116 | -8.67220 | -2.46087 | Hsp90 inhibitor |
| 658 | 656 | Maraviroc, MVC, UK-427857, Selzentry, Celsentri | -0.78531 | -1.11630 | -1.04624 | 0.94183 | -0.08576 | -0.91721 | -0.15483 | -0.45405 | -0.22948 | Chemokine CCR5 Antagonists; HIV Attachment Inhibitors; Reverse Transcriptase Inhibitors |
| 659 | 657 | LBH-589, Panobinostat, NVP-LBH589, LBH589 | -0.00333 | 2.19175 | 4.10789 | 2.76669 | -9.83468 | -7.59140 | -8.92487 | -10.82170 | -1.78296 | Histone Deacetylase (HDAC) Inhibitor |
| 660 | 658 | Laropiprant, L-888839, MK-0524 | 0.22193 | -1.07106 | 0.72081 | -0.85270 | -0.81947 | -0.46248 | 1.41722 | -1.39613 | -0.14220 | Prostanoid DP (DP1) Antagonists |
| 661 | 659 | Cisplatin, cis-Diammineplatinum(II) dichloride, cis-Dichlorodiammine platinum(II), cis-Platinum(II) diammine dichloride | -0.18966 | 0.20851 | -1.79175 | 0.18434 | 0.00653 | -1.92656 | -1.03876 | -0.74830 | -0.33590 | DNA alkylating drug, BIRC4 expression enhancer, TNFSF6 expression inhibitor, Thioredoxin Reductase inhibitor |
| 662 | 660 | Oxaliplatin, Eloxatin | -0.58550 | -1.74660 | -1.99903 | -0.14740 | 0.28295 | -1.53448 | -0.57742 | -0.38685 | -0.42461 | DNA Alkylating Drug |
| 663 | 661 | Retinoic acid, ATRA, Tretinoin, Vitamin A acid, all-trans-Retinoic acid, NSC-122758, Airol, Atralin, Avita, Nexret, Renova, Retacnyl, Retin-A Micro, Retinova, Vesanoid | 0.32135 | -3.16670 | -9.62561 | -12.22145 | -12.65746 | -13.78069 | -13.32233 | -13.44718 | -4.94110 | retinoic acid receptor (RAR) and the retinoid X receptor (RXR) agonist |
| 664 | 662 | Prasugrel, Effient, Efient, Prasita | -2.03646 | 1.18311 | 1.01223 | -0.44465 | -0.37047 | 0.32763 | 0.57533 | 0.41789 | 0.04216 | Platelet inhibitor, P2Y12 (P2T) antagonist |
| 665 | 663 | Niclosamide | -0.63813 | -0.97449 | -1.08337 | -1.16185 | -2.04912 | -2.05873 | -2.46417 | -2.81069 | -0.83983 | STAT3 inhibitor |
| 666 | 664 | MDV 3100, MDV-3100, NC-54 | 0.21563 | -0.20813 | 0.82303 | 0.04453 | 0.15234 | 0.74063 | 0.69500 | -0.34527 | 0.13433 | Androgen receptor (AR) antagonist |
| 667 | 665 | SB939, SB-939, Pracinostat | -0.89808 | 1.31144 | -1.09384 | 1.80015 | 1.96121 | -2.28131 | -5.09183 | -9.00311 | -0.84331 | HDAC inhibitor |
| 668 | 666 | Daunorubicin hydrochloride, Rubidomycin hydrochloride | 0.34557 | 2.28728 | 0.67893 | -5.04528 | -8.75878 | -9.95111 | -3.74881 | -11.76995 | -2.28103 | DNA Topoisomerase inhibitor |
| 669 | 667 | Idarubicin hydrochloride, Idamycin, Zavedos, 4-demethoxydaunorubicin | 0.26381 | -3.83384 | -6.69598 | -7.64500 | -10.56945 | -11.59661 | -3.67847 | -5.71762 | -3.13802 | DNA Topoisomerase inhibitor |
| 670 | 668 | Epirubicin hydrochloride, Ellence, Pharmorubicin, Epirubicin Ebewe | 0.08907 | -0.14327 | -1.45865 | 2.98572 | -5.77017 | -10.36371 | -8.89962 | -12.62604 | -2.29527 | DNA Topoisomerase II inhibitor |
| 671 | 669 | ERB 041, ERB-041, WAY-202041, Prinaberel | 0.46641 | -0.95291 | 0.08850 | -0.17545 | 0.07002 | 0.41944 | 0.59722 | -0.36255 | 0.00956 | Estrogen Receptor (ER) beta agonist |
| 672 | 670 | Irestatin 9389 | 0.31740 | -0.98376 | -1.22097 | -0.47782 | -0.15377 | -3.03227 | -1.07102 | -2.77387 | -0.59598 | IRE1 inhibitor |
| 673 | 671 | GSK 4716 | -0.49673 | 0.64317 | -0.09714 | -1.07511 | -0.09997 | 0.01835 | -0.01627 | -1.04559 | -0.13760 | Estrogen-Related Receptor beta (ERRbeta, ERR2) agonist, Estrogen-Related Receptor gamma (ERRgamma) agonist |
| 674 | 672 | Carfilzomib, PR-171, ONO-7057 | -0.81471 | 5.94292 | -6.82815 | -8.12260 | -0.97765 | -9.26634 | -4.89481 | -9.25880 | -2.17054 | Proteasome Inhibitor |
| 675 | 673 | SB 225002, SB-225002 | -0.13612 | -0.79501 | 1.35184 | 0.54854 | -1.24073 | 0.36272 | 1.47757 | -4.43582 | -0.18185 | Chemokine CXCR2 (IL-8 beta Receptor) antagonist |
| 676 | 674 | MGCD0103, MGCD-0103, MG-0103, Mocetinostat | -1.24158 | -0.03670 | -1.42781 | -1.43383 | -0.13007 | -3.41140 | -3.92163 | -8.26959 | -1.26049 | HDAC 1 inhibitor |
| 677 | 675 | LAQ824, LAQ-824, NVP-LAQ-824, Dacinostat | 2.35638 | 2.43701 | 2.76261 | -3.76171 | -8.14347 | -9.26799 | -5.89921 | -8.75037 | -1.79292 | HDAC inhibitor |
| 678 | 676 | Wnt-C59 | -0.19256 | 0.16546 | -1.61864 | 0.14254 | -1.48275 | 0.84762 | 1.31607 | -0.62952 | -0.09209 | Wnt inhibitor |
| 679 | 677 | ITF2357, ITF-2357, Givinostat hydrochloride | 0.53535 | -0.57211 | 0.31065 | 3.11198 | 3.35904 | -4.03081 | -9.06971 | -7.71900 | -0.89273 | HDAC Inhibitor, IL-1beta production inhibitor, TNF-alpha release inhibitor |
| 680 | 678 | DAPT, AN-37124, GSI-9, LY-374973 | -0.19513 | -0.76732 | 0.29150 | -0.43772 | -0.15576 | 1.65949 | -0.72300 | 0.45189 | 0.00786 | Antiamyloidogenic agent, gamma-secretase inhibitor, Notch signaling inhibitor |
| 681 | 679 | Azacitidine, 5-aza-CR, NS-17, NSC-102816, U-18496, 5-Azacytidine, Azacytidine, Ladakamycin, Mylosar, Vidaza | -0.59823 | 1.18838 | -1.88864 | -0.26064 | 0.15635 | -0.28388 | 0.41445 | 1.02557 | -0.01564 | Antimetabolite, DNA methyltransferase (DNMT) inhibitor |
| 682 | 680 | Decitabine, 5-aza-CdR, AzadC, DAC, E-7373, NSC-127716, 5-Aza-2'-deoxycytidine, Dezocitidine, Dacogen | 0.71868 | 0.87348 | 0.22397 | 0.79346 | -0.99990 | -1.65313 | -1.78450 | -2.43077 | -0.27013 | Antimetabolite, DNA methyltransferase (DNMT) inhibitor |
| 683 | 681 | MC1568, MC-1568 | -0.00378 | -0.64341 | -0.87057 | 0.30082 | -0.63312 | -0.92996 | 1.06821 | 1.59001 | -0.00773 | HDAC2 inhibitor |
| 684 | 682 | BSI-201, Iniparib, NSC-746045, IND-71677, INO2BA, NIBA, SAR-240550 | -0.70353 | -0.02023 | 0.05153 | 1.41224 | -0.09355 | 1.15637 | -1.53187 | 0.70461 | 0.06188 | PARP1 inhibitor |
| 685 | 683 | PCI-24781, CG-781, CRA-024781, S-78454, Abexinostat | 1.08661 | 1.39300 | -1.58161 | 5.12476 | 6.15355 | -3.37507 | -11.70138 | -7.32403 | -0.64851 | HDAC inhibitor |
| 686 | 684 | EX 527, EX-527, SEN-0014196, Selisistat | -0.82008 | -1.24399 | -0.22499 | -0.37694 | 0.43464 | -1.46394 | 0.87144 | -1.04845 | -0.24562 | NAD-Dependent Protein Deacetylase Sirtuin-1 (SIRT1) inhibitor |
| 687 | 685 | JNJ-26481585, Quisinostat | 4.37221 | 4.18773 | -2.87239 | -7.43015 | -4.81506 | -4.67815 | -6.91469 | -6.31329 | -1.55171 | HDAC1 inhibitor |
| 688 | 686 | Belinostat, PX-105684, PXD-101, Beleodaq | -0.77481 | -0.42715 | -0.17065 | -0.43303 | 1.72166 | -1.27722 | -2.16552 | -7.02117 | -0.66904 | HDAC1, HDAC2, Angiogenesis inhibitor, Apoptosis inducer |
| 689 | 687 | Thiazovivin | -1.73573 | 0.75543 | -0.72608 | 0.63658 | -0.83161 | -1.96940 | -2.77352 | -3.32785 | -0.63252 | Stem cell signaling |
| 690 | 688 | Phenylbutyrate Sodium, EL-532, NaPb, VP-101, Phenylbutyric acid sodium salt, Ammonaps, Buphenyl, Lunaphen, TriButyrate, Ammonapse | 0.41150 | 0.13349 | -4.21910 | 0.46070 | -0.02490 | -0.81055 | 1.31378 | -0.08892 | -0.17912 | HDAC inhibitor |
| 691 | 689 | Garcinol (BML-GR343) | -1.43740 | 1.08816 | -0.23650 | -0.85320 | 0.53824 | -0.74793 | -2.29140 | -0.80592 | -0.30103 | Histone N-Acetyltransferase (HAT) inhibitor |
| 692 | 690 | Pifithrin-alpha | -1.53682 | -2.04613 | -2.43582 | -3.03916 | -4.79587 | -5.15560 | -5.69418 | -5.94975 | -1.94430 | p53 inactivator |
| 693 | 691 | DMOG, Dimethyloxallyl Glycine | 0.12745 | 1.16989 | -1.38783 | -0.51783 | -0.44804 | -0.75712 | 0.54400 | -7.35424 | -0.54699 | HIF-alpha prolyl hydroxylase (HIF-PH) inhibitor |
| 694 | 692 | RG-108, NSC-401077 | -0.04688 | 2.39182 | 0.01540 | 0.09061 | -0.04294 | -1.29737 | -0.53421 | -0.90472 | -0.02082 | DNA methyltransferase (DNMT) inhibitor |
| 695 | 693 | S-Adenosylhomocysteine, SAH, AdoHcy | 0.23674 | -2.17026 | -1.47842 | 2.42619 | -0.17948 | -0.15287 | 1.85187 | 1.13021 | 0.10554 | Histone-Lysine N-Methyltransferase, H3 Lysine-79 Specific (DOT1L) Inhibitor |
| 696 | 694 | Anacardic Acid | 0.13878 | 0.80846 | -2.30730 | -0.66203 | -0.69643 | -1.45766 | 0.30372 | 1.66960 | -0.13972 | Histone acetyltransferase (HAT) inhibitor |
| 697 | 695 | Cyclic Pifithrin-alpha hydrobromide | 0.56343 | -2.36164 | -2.41432 | -3.31220 | -4.65800 | -4.45108 | -5.10433 | -7.25761 | -1.83916 | p53 inhibitor |
| 698 | 696 | BIX01294, BIX-01294 | 0.42626 | -0.28670 | 0.06955 | 1.86167 | 1.14122 | 1.18942 | 0.28599 | -0.32453 | 0.27673 | DNA methyltransferase (DNMT) inhibitor, Histone-lysine N-methyltransferase EHMT2 (H3-K9-HMTase 3; G9a) inhibitor |
| 699 | 697 | Vinorelbine, ANX-530, KW-2307, NVB, SDP-012, Navelbine, Eunades, Exelbine | 0.76691 | 6.32249 | 2.05944 | -1.07390 | -5.79038 | -5.36719 | -3.49018 | -3.58781 | -0.64448 | Tubulin polymerization inhibitor |
| 700 | 698 | Zebularine, NSC-309132 | -0.88535 | -0.12803 | 0.47548 | 0.15025 | -0.73654 | -0.84244 | 0.27146 | 0.31829 | -0.08733 | Cytidine deaminase inhibitor, DNA methyltransferase (DNMT) inhibitor |
| 701 | 699 | IDE1, IDE-1 | 1.30753 | -1.47638 | 0.10519 | 0.05081 | -0.14426 | -0.05763 | 0.60123 | 0.45548 | 0.05341 | Stem cell differenciation |
| 702 | 700 | trans-2Phenylcyclopropylamine hydrochloride, Tranylcypromine hydrochloride, 2-PCPA, Parnate | 0.54707 | 0.05687 | 0.03918 | 1.17262 | 0.00950 | -1.90105 | -0.35718 | 1.21102 | 0.04935 | Lysine-specific demethylase 1 (LSD1) inhibitor |
| 703 | 701 | Dexamethasone, ISV-305, OTO-104, Decadron, DexaSite, Maxidex, ProDex | -2.74067 | -2.60968 | -6.63230 | -7.64003 | -11.16445 | -11.92807 | -11.18704 | -11.50058 | -4.14842 | Antiinflammatory drug |
| 704 | 702 | Doxycycline hyclate, ALTY-0501, Cil-101, Atridox, Doryx, NanoDOX, Oracea, Periostat, Vibramycin hyclate | -1.15985 | -0.76918 | -0.79831 | -0.64492 | -0.58447 | 0.42160 | -2.40453 | -1.37578 | -0.46401 | 30S Ribosomal protein inhibitor, Matrix metalloproteinase inhibitor |
| 705 | 703 | SHH Antagonist, JK 184 | 1.98349 | -0.02814 | -0.81713 | -0.85637 | 0.31100 | -4.59722 | -10.11848 | -7.76217 | -1.38814 | SHH antagonist |
| 706 | 704 | SMO Antagonist, CUR-1 | -0.27580 | -0.08466 | -3.07119 | 0.75276 | 1.03523 | -2.06438 | -0.54771 | -1.72502 | -0.37935 | SMO antagonist |
| 707 | 705 | Purmorphamine | 0.57451 | -0.79115 | -1.07347 | 0.59815 | 0.34273 | 0.67060 | 2.15609 | 1.37156 | 0.24414 | hedgehog signaling pathway activator |
| 708 | 706 | Sodium butyrate, NaB | -0.98035 | -0.43797 | -0.27738 | 0.81556 | 0.26504 | 1.60043 | 0.88234 | -0.61147 | 0.07968 | HDAC inhibitor |
| 709 | 707 | Valporic Acid Sodium Salt | 0.02126 | 0.08589 | 0.13091 | -1.36847 | 0.08258 | -0.31537 | 1.79814 | 1.46144 | 0.12029 | Etoposide cytotoxic Enhancer in human glioblastoma cells |
| 710 | 708 | Procaine hydrochloride | 0.06511 | -0.17374 | 0.08890 | 0.90719 | 0.40558 | 0.50727 | -0.39862 | -0.65619 | 0.04729 | HGM-CoA reductase mRNA expression inhibitor |
| 711 | 709 | Pyrintegrin | -0.11923 | -0.86657 | -0.57924 | 0.67888 | 0.33353 | 1.63780 | 0.14646 | -0.04184 | 0.07547 | Promote human embryonic stem cell (hESC) survival |
| 712 | 710 | Sinefungin, A-9145, Antibiotic 32232RP, Compound-57926 | -0.63784 | -2.19847 | -0.26329 | -0.21653 | 1.54905 | 1.44473 | 0.38817 | -1.28560 | -0.07737 | Coactivator Associated Arginine Methyltransferase 1 (CARM1, PRMT4) inhibitor |
| 713 | 711 | IDE-2, IDE2 | -0.00916 | -0.28441 | -0.11800 | 1.20035 | 0.65547 | -0.41500 | 1.40389 | 1.81795 | 0.26964 | Stem cell differenciation |
| 714 | 712 | UNC0638 | 0.30448 | -0.14044 | -1.55223 | 0.26879 | -0.45123 | -0.62410 | -0.13639 | 1.04113 | -0.08182 | Histone-lysine N-methyltransferase EHMT2 (H3-K9-HMTase 3; G9a) Inhibitor |
| 715 | 713 | S-HDAC-42, AR-42, (S)-HDAC-42, NSC-736012, OSU-42, OSU-HDAC42 | 0.19412 | 0.93681 | -0.19846 | 3.19010 | 1.37537 | -1.68286 | -7.10456 | -9.84626 | -0.83318 | HDAC1, HDAC2 inhibitor |
| 716 | 714 | SC-1, Pluripotin | -0.71777 | -0.64004 | -1.45056 | -1.03220 | -0.62522 | -6.37575 | -8.47912 | -7.70305 | -1.71408 | Activator of stem cell renewal, RasGAP and ERK1 inhibitors |

| | A | B | C | D | E | F | G | H | I | J | K | L |
|---|---|---|---|---|---|---|---|---|---|---|---|---|
| 717 | 715 | (-)-Indolactam V | -0.18380 | -1.64624 | -2.08013 | -0.30967 | -0.94787 | 2.20754 | 3.10907 | 9.30385 | 0.59958 | PKC activator, human embryonic stem cells (ESCs) differentiation |
| 718 | 716 | Trapoxin A, RF-1023-A | -6.34249 | -11.89892 | -11.44047 | -10.63967 | -5.28134 | -7.04905 | -7.81486 | -7.38543 | -4.30378 | HDAC inhibitor |
| 719 | 717 | Chaetocin, Chetocin | 0.12874 | 1.79851 | -1.37050 | -4.29058 | -12.02719 | -3.90641 | -8.45286 | -6.59746 | -2.20210 | Apoptosis inducer |
| 720 | 718 | RITA, NSC-652287, NSC-652287D | -0.08156 | -1.42606 | -1.28878 | -0.95545 | -1.73840 | -0.80735 | -5.27412 | -7.73682 | -1.22472 | MDM2 (hdm2) inhibitor, Thioredoxin reductase inhibitor |
| 721 | 719 | OICR0000858A01, keto-2k, NCS-12k | 0.41127 | -0.96463 | -0.14822 | -1.93162 | 0.73554 | 1.34093 | 0.94831 | 0.35358 | 0.04726 | SIRT1 inhibitor |
| 722 | 720 | OICR0000860A01, SRT2183, SRT-2183 | -1.05005 | 0.22852 | -0.51492 | -1.23976 | -1.67207 | -4.15494 | -12.89978 | -6.59919 | -1.76980 | SIRT1 activator |
| 723 | 721 | Daunomycinone, Daunomycin Aglicone, Daunomycinon, Daunorubicin Aglycone, Daunorubicinone, Leukaemomycinone C, NSC-109351 | -1.54516 | -0.41408 | -0.32925 | -0.11397 | -0.27211 | -1.69672 | -0.35156 | -1.34218 | -0.38470 | Daunorubicin primary metabolite |
| 724 | 722 | WDR5 Inhibitor Compound 1 | -0.30111 | 0.45498 | 0.65707 | 0.98183 | -0.00509 | 0.02183 | -0.70084 | 0.42436 | 0.09724 | WDR5 Inhibitor |
| 725 | 723 | N-(Trifluoroacetyl)daunorubicin | 0.16942 | 0.47150 | 1.29375 | 2.11491 | -2.36622 | -6.32569 | -6.16957 | -5.67186 | -1.04554 | Analogue of Valrubicin |
| 726 | 724 | Etoposide, EPEG, NK-171, NSC-141540, VP-16, VP-16-213, Etopol, Etosid, Lastet, Vepesid, Vepeside | 0.02138 | -1.26026 | -1.04633 | 2.41167 | 1.86431 | 1.87856 | -1.14966 | -4.74354 | -0.12837 | DNA Topoisomerase II alpha Inhibitor, DNA-Intercalating Drug |
| 727 | 725 | Capecitabine, R-340, RG-340, Ro-09-1978, Xeloda | -1.92359 | 0.32363 | 0.43624 | 1.12231 | -1.16169 | 0.16663 | 2.90629 | 0.33368 | 0.13977 | Pyrimidine Antagonist |
| 728 | 726 | Vincristine sulphate, NSC-67574, Oncovin | 1.02573 | 3.71364 | 5.65059 | -0.09568 | -6.50562 | -4.92951 | -7.90497 | -3.66005 | -0.80592 | Tubulin polymerization inhibitors, RNA synthesis inhibitor, Cell cycle inhibitor |
| 729 | 727 | Cytarabine, Ara-C, Cytosine b-D-arabinofuranoside, AraC, CHX-3311, NSC-63878, U-19920A, Aracytin, Cylocide, Cytosar, Cytosar-U | -0.17319 | -4.65011 | -5.41657 | -6.15155 | -9.99345 | -5.26304 | -9.40131 | -9.59528 | -3.21232 | Antimetabolite, DNA Polymerase Inhibitor |
| 730 | 728 | Bleomycin sulfate | -1.23211 | -1.20583 | -0.71706 | 1.00444 | 0.69299 | 1.25339 | 1.24875 | -5.98731 | -0.31351 | Antimitotic Drug |
| 731 | 729 | Amsacrine Hydrochloride, CI-880, NSC-156303, NSC-249992, SN-11841, m-AMSA, NSC-141549 (HCl), Amecrin, Asidine, Amsidyl | 0.50304 | -0.91586 | -0.87588 | 2.12604 | 3.47185 | -3.37237 | -9.05590 | -9.59313 | -1.12346 | DNA Topoisomerase II Inhibitor, DNA-Intercalating Drug |
| 732 | 730 | Busulfan; Myleran; KRN-246; Busilvex; Partaject; Busulfex; Busulphan; NSC-750; Spartaject; Busulfanex | 0.34563 | -0.52035 | -4.08044 | 0.16122 | -0.54646 | 0.68271 | 2.71617 | 0.37121 | -0.05520 | DNA-Damaging Drug, Antimitotic Drug, DNA alkylating agent |
| 733 | 731 | Bendamustine HCl; Levact; Innomustine; Treakisym; Symbenda; Levact; Benda; Inno-P08001; Ribomustin; SyB L-0501; Treanda; SDX-105; Cytostasan | -0.42261 | -0.53098 | -1.64636 | 2.05209 | 0.65360 | 1.16801 | 1.84492 | 1.21121 | 0.27464 | Antimetabolites; Apoptosis Inducers; DNA Alkylating Drugs |
| 734 | 732 | Carmustine; NPC-08; SK-27702; NSC-409962; BCNU; Gliadel; Nitrumon; Carmubris; BiCNU; Becenum | -0.37531 | -0.37777 | -1.25450 | 1.03586 | 0.04165 | -1.50801 | 0.57903 | 0.52750 | -0.08446 | Apoptosis Inducers; DNA Alkylating Drugs; Glutathione Reductase (NADPH) Inhibitors; Thioredoxin Reductase Inhibitors |
| 735 | 733 | Chlorambucil; NSC-3088; Leukeran | 0.51405 | -0.76357 | -0.35086 | -0.31050 | -0.16775 | -0.68063 | -0.29338 | 0.22745 | -0.11577 | DNA-Damaging Drugs, DNA Alkylating Drug |
| 736 | 734 | Cladribine; Movectro; NSC-105014; Leustatin; Litak; Leustat; Mylinax; RWJ-26251-000; RWJ-26251; 2-CdA | -1.07524 | -1.30566 | -6.66152 | -10.05544 | -3.93433 | -8.71526 | -8.73595 | -10.22799 | -3.21656 | Adenosine Deaminase (ADA) Inhibitors; Antimetabolites; Apoptosis Inducers |
| 737 | 735 | Clofarabine; SAR-393590; Clofarex; CAFdA; Evoltra; Clolar; 2-Cl-2'-F-araA | -2.98001 | -8.62159 | -10.84804 | -9.07349 | -8.87687 | -11.18342 | -6.72450 | -7.82357 | -4.19464 | DNA Polymerase Inhibitors; Ribonucleoside-Diphosphate Reductase Inhibitors |
| 738 | 736 | Clafen, Revimmune; Cytoxan; Neosar; Endoxana; NSC-26271; Endoxan; Cyclophosphamide | -0.22242 | 0.08860 | -3.27382 | -0.07995 | -0.05463 | -0.84960 | 1.88854 | 0.79097 | -0.10861 | Apoptosis Inducers; BCL2 Expression Inhibitors; LGALS1 Expression Inhibitors |
| 739 | 737 | Fludarabine, F-ara-A, NSC 118218 | -0.44058 | -1.19273 | 0.80278 | -0.91476 | -1.51498 | -4.62670 | -9.45912 | -6.14220 | -1.48983 | STAT1 activation inhibitor, DNA synthesis inhibitor |
| 740 | 738 | Dacarbazine; Deticene; NSC-45388; DTIC-Dome | -0.28286 | 0.21245 | 0.07895 | 0.57468 | -0.81972 | -0.45625 | 0.79569 | -0.95939 | -0.05432 | DNA Alkylating Drug |
| 741 | 739 | Actinomycin D, Dactinomycin, Cosmegen, NSC-3053 | -8.96955 | -6.99667 | -10.43955 | -2.73320 | -6.83517 | -7.23006 | -6.42550 | -8.87652 | -3.71097 | DNA-Directed RNA Polymerase Inhibitor |
| 742 | 740 | 5-FU; 5-Fluorouracil; Carac; Flurobastin; Efudix; Adrucil; Fluorouracil; Ro-29757; NSC-19893; Efudex | 0.75568 | 1.17407 | -0.81402 | 0.15640 | -0.15318 | -2.34397 | 0.61899 | -2.47679 | -0.19554 | Antimitotic Drugs; Apoptosis Inducers; Pyrimidine Antagonists |
| 743 | 741 | Hydroxyurea; Onco-Carbide; Siklos; Hydroxycarbamide; Litalir; Mylocel; Droxia; Hydrea | 0.77849 | -0.03601 | -1.38965 | -1.27735 | -0.80089 | -0.65612 | 1.55736 | -1.65676 | -0.22079 | Antimetabolites; Nitric Oxide Donors; Ribonucleoside-Diphosphate Reductase Inhibitors |
| 744 | 742 | Ifosfamide; Ifomide; Z-4942; Mitoxana; Holoxan; Ifex; NSC-109724; MJF-9325 | 0.35514 | -0.33123 | -0.87408 | 0.91997 | -0.03813 | 0.76186 | 0.18036 | -0.38633 | 0.03727 | DNA Alkylating Drugs |
| 745 | 743 | Vindesine Sulfate Salt; Eldisine; Compound 112531; Vinca Alkaloids | 0.83148 | 4.77329 | 9.49905 | 8.17467 | -5.03890 | -2.29935 | -7.63696 | -3.09312 | 0.33047 | Antimitotic Drugs; Tubulin polymerization inhibitors; Cell cycle inhibitors |
| 746 | 744 | Melphalan; CDX-353; NSC-8806; CB-3025; Alkeran | 0.87731 | -1.98371 | -1.22943 | 1.77147 | -6.86803 | 2.26306 | 1.45431 | -2.55653 | -0.39780 | DNA Alkylating Drugs |
| 747 | 745 | Xaluprine; DR-6MP; NCS-755; 6-Mercaptopurine; Purinethol; Mercaptopurine | -1.29454 | -1.22687 | -1.62095 | -1.56665 | -4.07898 | -4.82486 | -5.92297 | -6.42095 | -1.70983 | Purine Antagonists |
| 748 | 746 | Sodium 2-mercaptoethanesulfonate; Mesna; Uromitexan; S-7878; D-7093; Mesnex | 0.31705 | 0.08681 | 0.40995 | -0.21920 | 0.88634 | -0.58635 | -1.20426 | 0.16888 | -0.00893 | Antioxidants |
| 749 | 747 | Mitomycin C; Mitosol; Mitomycin; MMc; Ametycine; Mito-medac; Mutamycin | 0.57715 | 0.62311 | 0.53583 | 0.39144 | -1.87777 | -5.41978 | -6.91477 | -9.86468 | -1.39223 | Apoptosis Inducers; DNA Alkylating Drugs |
| 750 | 748 | 2,4'-DDD; Lysodren; o,p'-DDD; NSC-38721; CB-313; Mitotane | 0.12238 | -0.98137 | -0.90788 | 1.47578 | -1.33922 | -1.53362 | -0.66171 | 1.26350 | -0.16251 | Antineoplastic Drug |
| 751 | 749 | Mitoxantrone diHCl; CL-232315; Ralenova; Eslep; Onkotrone; DAD; NSC-301739; Novantrone; Novantron | 0.79347 | 0.81446 | -0.84540 | 0.97023 | -4.26393 | 11.45760 | 43.02091 | 52.19130 | 6.60538 | DNA Topoisomerase II Inhibitors; DNA-Intercalating Drugs |
| 752 | 750 | Pentostatin; CI-825; Nipent; CL-67310465; Covidarabine; Deoxycoformycin; PD-81565; YK-176; NSC-218321; Coforin; Oncopent | 1.12914 | -0.17801 | -0.71334 | 0.40119 | -1.77593 | -0.05006 | -2.35825 | -0.00993 | -0.22550 | Adenosine Deaminase (ADA) Inhibitors; Antimetabolites |
| 753 | 751 | Caffeic acid phenethyl ester, CAPE | -0.52691 | -0.17871 | 0.22042 | 0.70167 | 0.32632 | 0.46121 | -0.23552 | 0.24761 | 0.06445 | Antiinflammatory Drugs; Antioxidants; Apoptosis Inhibitors; HIV Integrase Inhibitors; NF-kappaB (NFKB) Activation Inhibitors |
| 754 | 752 | Ro 106-9920 | 0.06270 | -1.38956 | -1.20274 | 1.25502 | 0.76978 | -1.31394 | -0.31909 | 2.72543 | 0.03727 | NF-kB activation Inhibitor |
| 755 | 753 | Procarbazine HCl; Natulan; Matulane; NSC-77213; Ro-04-6467/1 | 1.02926 | -0.61840 | -0.27039 | -0.07537 | -0.86431 | -0.06980 | -0.56901 | -0.49607 | -0.12268 | DNA Alkylating Drugs |
| 756 | 754 | Raltitrexed; ICI-D-1694; NSC-639186; ZN-D1694; ZD-1694; D-1694; Tomudex | -0.94980 | 0.18838 | -1.82264 | -6.10305 | -7.88370 | -2.75994 | -7.88840 | -6.28684 | -2.12524 | Pyrimidine Antagonists; Thymidylate Synthase Inhibitors |
| 757 | 755 | NPC-10; Streptozotocin; Streptozoticin; MNGU; Zanosar; U-9889; NSC-85998; Streptozocin | -0.55970 | 0.41468 | -0.83161 | 0.08376 | 0.43935 | -1.63223 | 0.32503 | 0.02499 | -0.11010 | DNA Alkylating Drugs |
| 758 | 756 | Topotecan Hydrochloride Hydrate; Nogitecan hydrochloride; NSC-609699; Hycamtin; NK-211; E-89/001; SK&F-S-104864-A; Evotropin | 2.24529 | 3.52635 | -2.18360 | -6.92348 | -4.88887 | -9.95270 | -6.07057 | -10.24102 | -2.18757 | DNA Topoisomerase I Inhibitors |
| 759 | 757 | Thioguanine; Tabloid; NSC-752; Tioguanine; 6-Thioguanine; Lanvis | -0.81790 | -1.56242 | -1.09069 | -1.51582 | -4.51244 | -8.89024 | -4.17076 | -5.28888 | -1.76644 | Purine Antagonists, disrupts cytosine methylation |
| 760 | 758 | Thio-TEPA; Tepadina; NSC-6396; ThioTEPA; Thioplex | -0.16366 | -0.22489 | 0.26121 | 0.47725 | -0.58440 | -0.29586 | -0.93595 | -1.94729 | -0.21652 | Cytochrome P450 Oxidase Inhibitors; DNA alkylating drugs |
| 761 | 759 | Temozolomide; NSC-362856; M&B-39831; Temodar; Sch-52365; CCRG-81045; Temodal | 1.12690 | -0.27767 | -3.46751 | -0.18836 | 0.47129 | -0.50162 | 1.32803 | 1.14998 | -0.02277 | DNA Alkylating Drugs; DNA Topoisomerase II alpha Inhibitors |
| 762 | 760 | Tegafur; Fental; FT; Utefos; Fulaid; Fluorofur; Ftorafur; FR-207; MJF-12264 | -0.05048 | 0.53304 | 0.85019 | 1.44072 | -0.65858 | 0.52662 | -0.34739 | 0.86424 | 0.20033 | Antimitotic Drugs; Apoptosis Inducers; Pyrimidine Antagonists |
| 763 | 761 | Chidamide; HBI-8000; CS-02100055; CS-055 | -0.54060 | 1.68156 | -3.81537 | -0.93566 | 0.30627 | -0.38676 | -1.54609 | -2.59070 | -0.49648 | Histone Deacetylase (HDAC) Inhibitors |
| 764 | 762 | UNC0321, UNC-0321 | 0.07077 | 0.03156 | -1.78966 | -0.58991 | 1.54580 | -4.52212 | 0.29015 | -0.61644 | -0.35392 | Histone-Lysine N-methyltransferase EHMT2 (H3-K9-HMTase 3; G9a) Inhibitors |
| 765 | 763 | CAY10603; YL-2-88; ST-2-92 | 1.01548 | 1.15114 | 0.29229 | -0.92442 | -0.25062 | -0.26550 | -3.94975 | -8.26959 | -0.71046 | Histone Deacetylase 6 (HDAC6) Inhibitors |
| 766 | 764 | TC-H 106 | 0.98035 | 0.15700 | -2.29881 | -2.02192 | -0.12378 | 0.37018 | 0.88728 | 1.31073 | -0.04687 | Histone Deacetylase (HDAC) Inhibitors |
| 767 | 765 | Tubastatin A (trifluoroacetate salt) | -0.27895 | -1.20383 | -0.13078 | 1.42401 | 0.12450 | 1.96814 | 4.62825 | 10.16260 | 1.05888 | Histone Deacetylase 6 (HDAC6) Inhibitors |
| 768 | 766 | Splitomicin | 0.01883 | -0.89617 | -2.82937 | -0.72040 | -0.28711 | -0.55716 | 0.44161 | -0.58256 | -0.34330 | Histone Deacetylase SIRT2 Inhibitors |
| 769 | 767 | Tenovin-1 | 0.07197 | -0.04676 | -0.87367 | -0.82861 | -2.27339 | 0.65501 | 0.67259 | -3.63135 | -0.39670 | SIRT1 and SIRT2 inhibitors |
| 770 | 768 | CBHA | 0.12072 | 1.10571 | -1.44629 | 0.96830 | 0.20361 | 3.43183 | 2.33362 | -3.95364 | 0.17531 | Histone Deacetylase (HDAC) Inhibitors; Apoptosis Inducers |
| 771 | 769 | Histone Deacetylase Inhibitor VI, HNHA | 0.79808 | -2.00970 | -0.88393 | 0.48146 | -0.67713 | 1.86071 | -1.88206 | -3.27112 | -0.35417 | Angiogenesis Inhibitors; Histone Deacetylase (HDAC) Inhibitors |
| 772 | 770 | Oxamflatin; 107-0130 | 2.26166 | 0.04173 | 1.16381 | 8.87694 | 5.96325 | 1.62239 | -3.68630 | -1.79916 | 0.91619 | Histone Deacetylase (HDAC) Inhibitors |
| 773 | 771 | Apicidin, OSI-2040 | -0.41615 | -2.03433 | -1.43232 | 1.52100 | 0.15760 | -4.94451 | -5.46069 | -10.27378 | -1.45145 | Histone Deacetylase (HDAC) Inhibitors |
| 774 | 772 | UNC0224, UCN-0224 | 0.89274 | -0.69939 | 2.53702 | 0.84673 | 0.10982 | -0.39150 | 0.07161 | 0.27315 | 0.23089 | Histone-Lysine N-methyltransferase EHMT2 (H3-K9-HMTase 3; G9a) Inhibitors |
| 775 | 773 | CCG-100602 | -0.23785 | 0.31997 | -0.27529 | 0.40654 | -0.62162 | 0.30896 | 0.96537 | 0.37852 | 0.07894 | Rho pathway inhibitor |
| 776 | 774 | Tamoxifen; Istubal; Valodex; Soltamox; ICI-46474; Nolvadex | -0.95133 | 0.08451 | -2.44823 | 0.76262 | 0.50033 | 2.30507 | 3.70450 | 4.16869 | 0.51543 | Protein Kinase C (PKC) Inhibitors; Selective Estrogen Receptor Modulators (SERM) |
| 777 | 775 | Abiraterone, CB-7598 | 0.40630 | -0.91595 | 0.79345 | -1.54195 | -1.58744 | 0.55058 | 1.97852 | 1.03327 | 0.04546 | Androgen Biosynthesis Inhibitors; Cytochrome P450 CYP17 (17alpha-Hydroxylase/C17-20 Lyase) Inhibitors |
| 778 | 776 | Anagrelide; Agrelin; Agrylin; BL-4162A; SPD-422; Thromboreductin; KRN-654; Xagrid; BMY-26538-01 | 2.17684 | 0.08007 | -1.45004 | 0.99558 | 0.42226 | -0.03130 | 0.89276 | 0.99926 | 0.25913 | Phosphodiesterase III Inhibitors |
| 779 | 777 | Anastrozole; Arimidex; ZD-1033; ICI-D1033 | -1.14978 | -0.24287 | -0.76178 | 0.00708 | -1.15650 | 1.35657 | 2.36703 | -0.40191 | 0.00113 | Aromatase Inhibitors |
| 780 | 778 | Bicalutamide; Casodex; Raffolutil; ICI-176334 | 1.46072 | 0.40584 | -0.00908 | 0.60248 | -1.54908 | 0.00854 | 0.27504 | 0.45970 | 0.10492 | Androgen Receptor Antagonists |
| 781 | 779 | EXE; FCE-24304; Aromasin; Exemestane; Nikidess; (8R,9S,10R,13S,14S)-10,13-Dimethyl-6-methylene-7,8,9,11,12,13,15,16-octahydro-6H-cyclopenta[a]phenanthrene-3,17(10H,14H)-dione | 0.06152 | -0.26122 | -1.29185 | 0.62720 | -2.02751 | -0.17098 | 0.08268 | -1.17045 | -0.26327 | Aromatase Inhibitors |
| 782 | 780 | Flutamide; NFBA; Odyne; Chimax; Prostandril; NK-601; FTA; Sch-13521; Eulexine; Eulexin; Drogenil | -1.56327 | 0.15608 | -0.36956 | 0.18688 | -0.66417 | 0.04844 | 0.63221 | 0.47668 | -0.06956 | Androgen Receptor Antagonists |
| 783 | 781 | Goserelin acetate; ZD-9393; ICI-118630; Zoladex | -0.06375 | -0.41475 | 0.29260 | -0.06115 | -0.05569 | 1.65536 | -0.10421 | 0.75574 | 0.12712 | GnRH (LHRH) Agonists |
| 784 | 782 | Hydrocortisone; Corhydron; Cortef; Chronocort; Dermacort; Cortisol; Alphaderm; Aeroseb-HC; Acticort; Plenadren; Proctocort; Anusol-HC; Cetacort; NSC-10483; DuoCort | 0.03790 | -1.41943 | 0.21405 | -2.39590 | -7.35875 | -8.20939 | -9.26627 | -9.82297 | -2.42429 | Glucocorticoid; Steroids |
| 785 | 783 | Letrozole; FEM-345; CGS-20267; Femara | 1.18454 | -0.36372 | -1.84825 | -0.01682 | -0.01480 | 0.57766 | -0.09167 | 0.87860 | 0.01938 | Aromatase Inhibitors |
| 786 | 784 | Mechlorethamine Hydrochloride; Clearazide; Mustargen; Mustine hydrochoride; Chlormethine hydrochloride; NSC-762; Nitrogen Mustard | 0.63944 | -0.02313 | -3.72833 | -0.24796 | -0.26694 | -0.76203 | 0.50717 | 0.03015 | -0.24430 | DNA Alkylating Drug |
| 787 | 785 | Medroxyprogesterone 17-acetate; Gestoral; Farlutal; MAP; Depo-subQ provera 104; Amen; Medroxyprogesterone acetate; NSC-26386; Depo-Provera; Provera; Cycrin | -0.48518 | -0.17170 | -2.11392 | -0.88046 | -4.01818 | -6.32284 | -6.77398 | -7.07542 | -1.76596 | Progesterone, androgen and glucocorticoid receptor agonist |
| 788 | 786 | Nilutamide; Nilandron; RU-23908; Anandron | 0.15636 | 0.67231 | 0.50221 | 0.44679 | 0.34442 | -0.95181 | 0.27499 | -0.35615 | 0.06908 | Androgen Receptor Antagonists |
| 789 | 787 | Prednisolone; Prelone | -0.70013 | -1.12636 | -4.80510 | -4.89336 | -8.23362 | -10.38613 | -10.74401 | -10.95601 | -3.28844 | Glucocorticoid receptors (GR) alpha and beta agonist - Irreversible |
| 790 | 788 | Prednisone; Rayos; Meticorten; NSC-10023; Lodotra; Deltadehydrocortisone; Deltacortisone; Decortin; Deltasone | 1.04350 | 0.12367 | 0.40250 | -0.49944 | -0.14670 | -0.42870 | -0.41774 | -0.84535 | -0.04873 | glucocorticoid receptor a(GR) gonist prodrug |
| 791 | 789 | LB 42708, LB-42708 | -0.31087 | -0.26259 | -2.42684 | 0.09364 | 0.42466 | 1.08470 | -2.24769 | -4.18808 | -0.49684 | Farnesyltransferase (geranylgeranyl pyrophosphate synthase) Inhibitors; Inhibitors of Signal Transduction Pathways |
| 792 | 790 | Vinblastine Sulfate; NSC-49842; Velban; Exal; Velsar; Velbe; 29060-LE, Vinca Alkaloids | 1.38226 | 9.04680 | -1.12392 | -3.53515 | -2.71639 | -2.25563 | -6.88672 | -3.52931 | -0.61006 | Antimitotic Drugs; Tubulin polymerization inhibitors |
| 793 | 791 | Lomustine; ; Cecenu; Belustine; CeeNU; NSC-79037; CCNU | -0.03368 | 0.45451 | -0.89896 | -0.28171 | 0.15116 | -1.37332 | 4.19810 | -1.25156 | 0.06118 | DNA Alkylating Drugs |
| 794 | 792 | Methotrexate; Amethopterin; MPI-5004; Intradose-MTX; Metolate; Maxtrex; G-301; APC-2002; CL-14377; Rheumatrex; Folex | 0.32212 | 1.09439 | 0.13730 | 1.69383 | -0.55710 | -3.21367 | -7.49855 | -6.34947 | -0.91154 | Dihydrofolate Reductase (DHFR) Inhibitors; Folate Antagonists |
| 795 | 793 | Satraplatin; JM-216; YHI-601; BMY-45594; BMS-182751 | 0.38110 | -0.19860 | 0.07236 | 0.82223 | -1.88578 | 0.25747 | -3.42990 | -8.74315 | -0.80708 | DNA Alkylating Drugs |
| 796 | 794 | Buserelin; Suprefact; Suprecur; Profact; Hoe-766 MP | -0.34274 | 0.82773 | -0.28886 | 0.35768 | -0.44373 | -0.50329 | -0.91508 | 0.45113 | -0.05437 | GnRH (LHRH) Agonists |
| 797 | 795 | Leuprolide; Lucrin; Onectyl; Enanton; Abbott-43818; Lupron; Prostap; A-43818; TAP-144; Enantone; Carcinil | -0.19385 | 1.11010 | -1.23838 | 0.71030 | 0.25198 | -0.44708 | 0.80523 | -0.69691 | 0.01912 | GnRH (LHRH) Agonists |
| 798 | 796 | Megestrol acetate; DW-ES(B); PAR-100.2; Megace ES; Megestat; Megace | -0.42834 | -0.70000 | -1.05653 | -0.46622 | -2.52500 | -3.73692 | -4.15813 | -5.27292 | -1.16354 | Progesterone Receptor Agonists |
| 799 | 797 | Teniposide; VM-26; Vumon; Podophyllotoxins | 0.04455 | 0.34592 | -1.45637 | 4.02527 | 0.11213 | -4.96726 | -10.33878 | -9.20776 | -1.36006 | DNA Topoisomerase II Inhibitors |
| 800 | 798 | Calcium Folinate Hydrate; Citrovorum factor; Folidan; Foliben; Emovis; Lederfolin; Leucovorin-25; Uzel; Leucosar; Lederfolat; Folaren; NSC-3590; Wellcovorin; Refolinon; Folinic acid calcium salt; Leucovorin calcium | -0.64495 | -0.21181 | -0.58264 | 1.46583 | 1.75143 | -0.51531 | -0.25908 | -0.68428 | 0.02025 | Thymidylate synthase inhibitor |
| 801 | 799 | EPZ004777, EPZ-004777 | -1.35395 | 1.23462 | -2.25135 | -0.78516 | -0.36214 | -0.66743 | -0.99201 | -0.85414 | -0.38257 | Histone-Lysine N-Methyltransferase, H3 Lysine-79 Specific (DOT1L) Inhibitors |
| 802 | 800 | I-BET151, GSK-1210151A, GSK1210151A | -0.52890 | 1.05102 | -1.39277 | -0.30615 | -0.81822 | -1.59170 | -3.35294 | -2.93202 | -0.62615 | Bromodomain-Containing Protein 2 (Brd2) Inhibitors; Bromodomain-Containing Protein 3 (Brd3, RING3-like protein) Inhibitors; Bromodomain-Containing Protein 4 (Brd4, HUNK1) Inhibitors |
| 803 | 801 | Lomeguatrib, 4-BTG, AZD-5896, PaTrin-2, Patrin | 0.76811 | 0.41543 | -0.39245 | 0.59218 | 0.43268 | -0.98365 | -1.24663 | -3.37472 | -0.24034 | DNA Repair Inhibitors; O6-Alkylguanine-DNA Alkyltransferase (MGMT) Inhibitors |
| 804 | 802 | MG-132, MG 132, ZLLL | -1.26343 | -0.37466 | -0.17457 | -1.02839 | -1.10710 | -1.90935 | 5.85093 | -6.30677 | -0.40045 | Proteasome Inhibitor |
| 805 | 803 | UNC 0646, UNC-0646, UNC0646 | -0.76593 | -0.18578 | 0.14742 | 1.10783 | -0.24685 | 0.64388 | 2.81820 | 1.98057 | 0.34882 | Histone-Lysine N-methyltransferase EHMT2 (H3-K9-HMTase 3; G9a) Inhibitor |
| 806 | 804 | NG38 | -0.52680 | -0.23535 | -3.21817 | -0.60827 | -1.41331 | 1.05859 | -0.28338 | -1.01345 | -0.39580 | Estrogen Sulfotransferase (EST) Inhibitor, CDK1 |
| 807 | 805 | Ftorafur, FR-207; FT; MJF-12264, Fluorofur; Ftorafur; Tegafurÿ(BAN; USAN; Rec INN), Fental; Fulaid; Utefos | -0.94278 | 0.48709 | -1.92378 | -0.74501 | -0.79228 | -0.83285 | -0.41484 | 0.87038 | -0.27237 | Thymidylate Synthase Inhibitor, antimetabolite |
| 808 | 806 | PFI 1, PFI-1 | 0.38665 | 1.25612 | -0.06565 | 0.26455 | -0.05789 | 0.30972 | 0.36742 | -1.38234 | 0.06841 | Bromodomain-Containing Protein 4 (Brd4, HUNK1) Inhibitor; Bromodomain-Containing Protein 2 (Brd2) Inhibitor |
| 809 | 807 | IOX 2 | -0.52659 | 0.27166 | -0.50259 | -1.19334 | -0.15649 | -0.56193 | 0.81248 | 0.30798 | -0.09824 | HIF-1a prolyl hydroxylase-2 (PHD2) |
| 810 | 808 | SCH 79797 dihydrochloride, SCH-79797 dihydrochloride | -0.64152 | -1.76258 | -0.37890 | 0.51218 | -0.34653 | -1.03163 | -3.24520 | -9.55725 | -1.04349 | PAR1 receptor antagonist |
| 811 | 809 | CX-5461 | -1.03679 | -0.13605 | 1.55805 | 1.40381 | -2.44341 | -7.59131 | -8.73251 | -8.77225 | -1.63332 | RNA Polymerase Inhibitor |
| 812 | 810 | Vinflunine ditartrate, BMS-710485; F-12158; L-0070, Vinflunineÿ(Prop INN; USAN), Javlor | 0.11542 | -0.24965 | -0.87193 | 1.41644 | 1.81632 | 1.70442 | -7.19271 | -9.54884 | -0.81256 | Antimitotic Drugs; Tubulin polymerization inhibitors |
| 813 | 811 | MLN2238, MLN-2238, Ixazomib | 0.12292 | -0.39402 | -0.98390 | -0.93951 | 0.34759 | -4.80213 | -8.87046 | -1.88984 | -1.10425 | Proteasome Inhibitor |
| 814 | 812 | C 646, C-646 | 0.88648 | -0.41783 | -0.21733 | -0.86958 | -0.76964 | 0.58516 | 0.62835 | 0.30126 | 0.00805 | p300/CREB-binding protein (CBP) inhibitor |
| 815 | 813 | A-966492 | -2.39777 | 0.06643 | -2.31605 | -0.95093 | -1.35718 | -0.64814 | -1.43709 | -1.48656 | -0.66773 | Poly(ADP-ribose)polymerase-1 (PARP-1) Inhibitor |
| 816 | 814 | AT-406, Debio-1143; SM-406 (former code) | -0.57714 | 0.38451 | -0.18733 | 0.97973 | 0.90903 | -0.25092 | 1.09876 | -0.78696 | 0.09956 | Inhibitor of Apoptosis Protein 2 (IAP2) Inhibitor; Apoptosis Inducers; Inhibitor of Apoptosis Protein 1 (IAP1) Inhibitor; X-Chromosome-Linked Inhibitor of Apoptosis Protein (XIAP) Inhibitor |
| 817 | 815 | TCS 401, NNC-52-0956 | -0.41297 | -2.92714 | -0.92374 | 0.90682 | 0.12878 | 0.00537 | 0.81623 | 0.50525 | -0.12060 | Protein Tyrosine Phosphatase PTP-1B Inhibitor |
| 818 | 816 | 5-Aminoimidazole-4-carboxamide-1-D-ribofuranoside, AICAR, AR-100; ATH-001; GP-1-110-0; SCH-900395, Acadesineÿ(BAN; USAN; Rec INN); Aica-riboside, Acadra; Protara; Arasine (former TM) | -1.11167 | -0.78592 | -0.47018 | 0.99839 | -0.94396 | -1.24799 | 0.20396 | -0.12633 | -0.22097 | AMP-Activated Protein Kinase (AMPK) Activators; Apoptosis Inducers |
| 819 | 817 | DMXAA, 5,6-MeXAA; AS-1404; ASA-404; DMXAA; NSC-640488; R-1564, Vadimezanÿ(Prop INN) | -0.01126 | 1.04653 | -1.51042 | -0.82766 | -0.24263 | -0.50078 | -0.07316 | -2.35679 | -0.28392 | Interferon Inducer, Vascular Disrupting Agents (VDA) |
| 820 | 818 | PPT | -0.84214 | -0.77647 | -0.53874 | 0.41196 | -0.97881 | 0.85920 | -0.11636 | -0.19156 | -0.13783 | Estrogen Receptor (ER) alpha Agonist |
| 821 | 819 | IDH-C227 | -0.55961 | -0.62791 | -1.81235 | 2.01277 | -0.01490 | -1.73386 | -0.90751 | 0.36370 | -0.20802 | Isocitrate dehydrogenase 1 (IDH1) inhibitor |
| 822 | 820 | Diphenyleneiodonium chloride, DPI | -0.09514 | 0.14600 | -0.59811 | -1.04674 | -2.31300 | -2.86800 | -1.70617 | -9.53868 | -1.14298 | Cytochrome P450 Inhibitor; Xanthine Oxidase Inhibitor; NADPH Oxidase (NOX) Inhibitor; Nitric Oxide Synthase Inhibitor |

| | A | B | C | D | E | F | G | H | I | J | K | L |
|---|---|---|---|---|---|---|---|---|---|---|---|---|
| 823 | 821 | Dp44mT, HDp44mT | -0.38215 | 1.86874 | -1.83238 | -0.72414 | -5.04701 | -6.14906 | -0.84309 | -7.38428 | -1.29987 | Chelating agent, Apoptosis Inducer |
| 824 | 822 | ABT-199, GDC-0199, RG-7601, Venetoclax | -0.36788 | 0.98645 | 1.86840 | 2.55581 | 1.26603 | 0.14799 | 0.75595 | -1.80663 | 0.34290 | Bcl-2 Inhibitor |
| 825 | 823 | MK-571 sodium salt hydrate, L-660711 | -2.06550 | 1.46891 | -1.18272 | 0.11575 | -0.22674 | -1.59113 | 0.97374 | 0.65893 | -0.11726 | Leukotriene CysLT1 (LTD4) Antagonist, Leukotriene Synthesis Inhibitor |
| 826 | 824 | Y-29794 oxalate | 0.45546 | -0.89723 | 0.26029 | -0.52393 | 0.95310 | 3.61152 | 5.11436 | -6.27375 | 0.17125 | Prolyl Endopeptidase (prolyl oligopeptidase; POP) Inhibitor |
| 827 | 825 | Ivermectin B1a, MK-933, Ivermectin, Cardomec; Mectizan; Sklice; Stromectol | -1.53328 | 1.02216 | 0.26566 | -0.24392 | 0.26018 | 0.21139 | 0.63821 | -4.29184 | -0.23287 | Drugs Acting on Chloride Channels |
| 828 | 826 | Gabexate mesylate, FOY | 0.44544 | -3.06290 | -0.86646 | 1.90128 | -0.26603 | -0.76813 | 0.40500 | -1.12435 | -0.21161 | Serine Protease Inhibitor; NF-kappaB (NFKB) Activation Inhibitor; AP-1 Inhibitor |
| 829 | 827 | MK-0752, L-000891675 | 0.31317 | 0.08118 | -1.14424 | 1.73874 | -0.55132 | -0.49837 | -0.65820 | -0.58766 | -0.08288 | gamma-Secretase Inhibitor, Notch Signaling Inhibitor |
| 830 | 828 | KD 5170, KD-5170 | -1.06535 | 2.33064 | -2.94962 | 0.63786 | -0.32318 | -0.27587 | 2.24222 | 2.56217 | 0.20036 | Histone Deacetylase (HDAC) Inhibitor, Apoptosis Inducer |
| 831 | 829 | ACY-1215, Rocilinostat | 0.01968 | 0.16366 | -3.56720 | 1.75237 | -0.81588 | 1.01244 | 1.33599 | 2.39486 | 0.14563 | Histone Deacetylase 6 (HDAC6) Inhibitor |
| 832 | 830 | ML-3H2 | -0.25027 | 0.95289 | -0.14266 | 0.45228 | -0.37976 | -1.16654 | -3.01403 | -2.75500 | -0.39980 | Human Porphobilinogen Synthase Inhibitor |
| 833 | 831 | Daminozide | -0.00343 | 0.40873 | -1.22387 | -0.69694 | -0.08813 | 1.77991 | 0.91417 | 0.67362 | 0.11189 | Histone demethylase KDM2, KDM7 Inhibitors, Histone Lysine Demethylase PHF8 (KIAA1111; ZNF422) inhibitor |
| 834 | 832 | Anacetrapib, MK-0859 | 0.40881 | 0.00650 | -1.01854 | -1.40393 | -0.32116 | -1.01050 | 0.22957 | -0.57866 | -0.23392 | Cholesteryl Ester Transfer Protein (CETP) Inhibitor, HDL-Cholesterol Increasing Agent |
| 835 | 833 | Example 1F | 1.31566 | 1.60381 | -1.33294 | -0.82034 | -0.42135 | 0.31149 | 0.02184 | 0.65195 | 0.08437 | Mcl1 Inhibitor |
| 836 | 834 | Sanguinarine chloride, Pseudochelerythrine; Sanguinarine chloride; Sanguinarium chlorideÿ(USAN; Rec INN) | -0.71088 | 1.49483 | 0.40864 | -0.51581 | -0.67151 | -0.09310 | -0.34663 | -0.12879 | -0.03573 | Rho GTPase Inhibitor; Caspase 3 Activator; Apoptosis Inducer; FtsZ Inhibitor; Acetylcholinesterase Inhibitor |
| 837 | 835 | Scriptaid, SB-556629 | 0.09547 | -0.62934 | -1.02072 | -0.74415 | 0.92035 | 0.67869 | 4.50364 | -3.38489 | 0.02658 | Histone Deacetylase (HDAC) Inhibitor |
| 838 | 836 | (-)-YK-4-279, YK-4-279 | -0.13262 | -0.13652 | -3.43890 | -0.40715 | 0.22786 | -0.51054 | 1.65419 | -3.84093 | -0.41765 | EWS-FLI1 Inhibitor |
| 839 | 837 | Doxorubicin hydrochloride, NSC-123127, Doxorubicin hydrochlorideÿ(Rec INNM; BAN; USAN; USP), Adriacin; Adriamycin; Adriblastina; Resmycin; Rubex | 0.66501 | -0.11871 | 1.24216 | -0.52738 | -7.91136 | -10.88935 | -12.67438 | -6.27784 | -2.31463 | DNA-Intercalating Drug; DNA Topoisomerase II alpha Inhibitor |
| 840 | 838 | Pyroxamide, NSC-696085 | 0.18289 | -0.81149 | 0.04342 | 1.15208 | 1.34245 | 1.82997 | 1.81572 | 1.97902 | 0.47788 | Histone Deacetylase (HDAC) Inhibitor |
| 841 | 839 | MKC204, MKC-204 | 0.00414 | 0.42572 | -3.38598 | 0.74884 | 0.44991 | -0.38487 | 1.47105 | -1.11369 | -0.11321 | IRE1 inhibitor |
| 842 | 840 | Pyrrolidinedithiocarbamate ammonium | 0.49900 | -0.11746 | -2.32918 | 1.33974 | 0.53543 | 0.73336 | -0.41978 | -2.92570 | -0.17028 | Nuclear factor KB (NF-kB) inhibitor |
| 843 | 841 | GSK J1, GSK-J1 | 0.29541 | -3.36322 | 0.68070 | -0.14452 | -0.42712 | -0.20077 | -0.50274 | 0.71898 | -0.18669 | Lysine-Specific Demethylase 6A (UTX) Inhibitor, TNF-alpha Production Inhibitor, Lysine-Specific Demethylase 6B (JMJD3) Inhibitor |
| 844 | 842 | Calpain Inhibitor I, ALLN, ALLnL, SIB-1980 | -0.33527 | -1.61441 | -1.95301 | 0.31457 | -1.25268 | -0.77673 | -1.63236 | -2.21834 | -0.60056 | mu-Calpain (Calpain-1) Inhibitor, Proteasome Inhibitor |
| 845 | 843 | 2-Methoxyestradiol, 2-ME, 2-MeOE2, 2ME2, NSC-659853, Panzem, Panzem NCD, PulmoLAR | 1.14266 | -0.84220 | 0.46582 | 1.36521 | -1.27373 | -1.88643 | 1.67847 | 2.19424 | 0.18039 | Hypoxia Inducible Factor 1-alpha (HIF-1alpha) Inhibitor, Apoptosis Inducer, Angiogenesis Inhibitor |
| 846 | 844 | (+)-JQ-1, (+)-SGCBD-01, JQ1 | 0.07651 | -1.17565 | -0.10092 | -0.11692 | -1.51273 | -3.48141 | -5.14991 | -5.67203 | -1.08673 | Bromodomain-Containing Protein 4 (Brd4, HUNK1) Inhibitor; Bromodomain-Containing Protein 2 (Brd2) Inhibitor; Bromodomain-Containing Protein 3 (Brd3, RING3-like protein) Inhibitor |
| 847 | 845 | Example 6 | 0.83263 | -0.67079 | 0.81339 | -1.07327 | -1.37116 | -0.82190 | 0.66978 | -0.26990 | -0.11996 | Mcl1 inhibitor |
| 848 | 846 | Convallatoxin | -0.66660 | -1.38484 | -9.12864 | -1.03118 | 2.32235 | 5.55738 | -4.48820 | -7.50220 | -1.03528 | Hypoxia Inducible Factor 1-alpha (HIF-1alpha) Inhibitor |
| 849 | 847 | OICR0003752A01, 676205 | 0.30243 | -0.91575 | -0.57535 | -0.05309 | 0.05147 | -0.17189 | -0.52595 | -0.31010 | -0.13943 | Reverse Transcriptase Inhibitor |
| 850 | 848 | Guanabenz acetate salt, WY-8678 acetate, Guanabenz acetateÿ(Rec INNM; USAN; JAN), Wytensin | -1.02979 | 0.02032 | -3.42134 | 0.62604 | -1.09836 | -2.35642 | -0.51425 | -1.82159 | -0.60862 | a2-adrenergic receptor agonist |
| 851 | 849 | Gossypol from cotton seeds, NSC-19048 | 0.25244 | 0.38570 | 0.01718 | 1.26043 | 0.72198 | 0.19953 | 0.29334 | -0.56308 | 0.16286 | Steroid 5alpha-Reductase Inhibitors; Bcl-xl Inhibitors; Apoptosis Inducers; 11beta-Hydroxysteroid Dehydrogenase (11beta-HSD) Inhibitors; Antimitotic Drugs |
| 852 | 850 | Isoliquiritigenin, ISLQ | 0.60456 | 1.02446 | -1.60246 | 0.82146 | 0.54176 | 0.28881 | 0.72824 | -0.28445 | 0.13462 | NMDA Antagonists; Histamine H2 Receptor Antagonists; Apoptosis Inducers; Guanylate Cyclase Activators; Aldose Reductase Inhibitors; Angiogenesis Inhibitors; Histone Deacetylase SIRT1 Activators |
| 853 | 851 | Ciclopirox, HOE-296, Ciclopiroxolamine, Batrafen; Ciclochem; Loprox; Mycoster; Penlac nail lacquer | 1.38231 | 0.51442 | -0.46629 | -1.29666 | -8.18420 | -0.62841 | -0.55462 | -0.17511 | -0.59677 | antimycotic drug |
| 854 | 852 | (R)-Ibuprofen, ST-1482; ZAG-1701, Ibuprofenÿ(BAN; USAN; Rec INN), Advil; Aktren; Amelior; Brufen; Brufen Retard; CDT-ibuprofen; Caldolor; Dolgit; Ibuprox; Motrin; Nurofen Meltlets; Pedea; Spedifen | -1.54669 | -1.20511 | -0.23009 | 0.70765 | -0.36932 | 0.08934 | 1.09248 | 0.25384 | -0.07661 | Cyclooxygenase-2 Inhibitors; Non-Steroidal Antiinflammatory Drugs ; NF-kappaB (NFKB) Activation Inhibitors; Cyclooxygenase-1 Inhibitors; Cyclooxygenase-3 Inhibitors |
| 855 | 853 | APHA-8, APHA Compound 8 | 2.23180 | -1.39301 | -0.69698 | 0.98936 | -0.44671 | -0.35710 | 1.31091 | 7.01555 | 0.54890 | Histone Deacetylase (HDAC) Inhibitor |
| 856 | 854 | LY2608204, LY-2608204 | -0.40640 | 0.35190 | -0.07917 | 0.14556 | -0.49478 | 0.15519 | 0.20840 | 1.37663 | 0.07975 | Glocokinase (GK) activator |
| 857 | 855 | PHAH (DC-002263) | 0.73791 | 1.50478 | -0.52787 | -1.50649 | -1.08760 | -0.67913 | 0.48977 | 3.48527 | 0.15328 | Enolase (ENO) inhibitor |
| 858 | 856 | KY 02111 | 0.28472 | 0.34951 | -0.81987 | 0.47445 | -0.75234 | -0.20975 | 0.78357 | -1.54078 | -0.09073 | Wnt signaling inhibitor |
| 859 | 857 | MALT1-MI-2, MI-2, MI 2 | -1.17544 | 1.18525 | -1.58748 | -0.46755 | 0.78055 | 1.46885 | 1.68697 | -1.36172 | 0.03358 | Mucosa Associated Lymphoid Tissue Lymphoma Translocation Protein 1 (MALT1) Inhibitor |
| 860 | 858 | IDH-C35 | -0.41665 | -1.17710 | -0.65980 | -0.56680 | -0.62085 | -0.13243 | -0.01285 | 1.21041 | -0.15071 | Isocitrate dehydrogenase (IDH) Inhibitor |
| 861 | 859 | GSK126, GSK-126 | 0.21287 | -0.02168 | -2.44995 | 0.72452 | -0.75544 | 0.29521 | 1.14136 | 2.24143 | 0.08806 | Histone-Lysine N-Methyltransferase EZH2 Inhibitor |
| 862 | 860 | CUDC-907 | 3.91968 | -5.60237 | -11.70466 | -9.59183 | -11.51278 | 0.03245 | -1.14082 | -5.20944 | -2.58851 | Histone Deacetylase (HDAC) Inhibitor, PI3K inhibitor |
| 863 | 861 | UNC 1215 | -1.66311 | 1.04214 | -2.58268 | -0.67728 | -0.56460 | -0.19627 | 4.31025 | 6.53232 | 0.39331 | L3MBTL3 inhibitor |
| 864 | 862 | BAX Activator BAM7 | -0.38454 | 0.09475 | -1.24209 | 0.75946 | 0.04697 | -0.55685 | -0.40124 | -0.13969 | -0.11564 | Bax activator |
| 865 | 863 | Tubacin, 415N3, JCWII-114 | 0.14054 | 0.69972 | -0.74456 | 2.85582 | -1.16764 | 1.83488 | 1.59230 | 3.34576 | 0.54275 | Histone Deacetylase 6 (HDAC6) Inhibitor |
| 866 | 864 | MLN-4924, Pevonedistat | 0.13568 | -1.44595 | 0.46569 | 0.84590 | 3.73762 | 14.78516 | 4.67764 | -5.88916 | 1.09811 | NEDD8-Activating Enzyme (NAE) Inhibitor |
| 867 | 865 | MI-3 | -0.41556 | -0.79681 | 0.22492 | 1.22588 | 5.23900 | 9.84762 | 9.85632 | 4.81744 | 1.90279 | Menin/MLL Interaction Inhibitor |
| 868 | 866 | Ubiquitin Protein Ligase Inhibitor | -0.26580 | -0.89562 | -0.98956 | 0.24207 | 0.73273 | -0.41007 | 0.64914 | 0.84688 | -0.00572 | Ubiquitin Protein Ligase Inhibitor |
| 869 | 867 | FH 535, FH-535 | -0.46332 | -0.54796 | -2.29624 | 0.14088 | 1.57250 | -0.24755 | 1.26815 | -4.81336 | -0.34168 | PPARgamma Antagonist, PPARdelta Antagonist |
| 870 | 868 | AMD 3100 octahydrochloride, AMD-3100, JM 3100, JM-3100, Plerixafor, SDZ-SID-791, Mozobil | 0.65297 | -0.24776 | -1.61542 | 1.66323 | -0.01057 | -0.29604 | 1.07057 | 0.73355 | 0.12372 | Chemokine CXCR4 (SDF-1 Receptor) Antagonist, Chelating Agent |
| 871 | 869 | SGC 0946, SGC-0946, SGC-946, SGC 946 | -0.06375 | -1.12358 | -0.51554 | -0.84165 | 0.88718 | -0.44568 | 3.21908 | 2.23663 | 0.21266 | DOT1L inhibitor |
| 872 | 870 | NCH 51, NCH-51, PTACH | -0.72732 | 1.15766 | 0.71280 | 0.11002 | -0.20477 | 3.19036 | -5.20500 | -3.89375 | -0.30826 | Histone Deacetylase (HDAC) Inhibitor |
| 873 | 871 | M 344, M-344, D-237, MS-344 | 0.60739 | -0.19475 | 1.07885 | 3.24434 | 2.90847 | 1.95867 | 1.51120 | -1.75813 | 0.59344 | Histone Deacetylase (HDAC) Inhibitor |
| 874 | 872 | GSK J4, GSKJ4 | -0.78228 | -1.06407 | 1.24283 | -0.23773 | -1.87670 | -0.80066 | -1.08540 | -9.38409 | -0.88725 | Lysine-Specific Demethylase 6A (UTX) Inhibitor, TNF-alpha Production Inhibitor, Lysine-Specific Demethylase 6B (JMJD3) Inhibitor |
| 875 | 873 | IOX 1, 5-carboxy-8HQ, IOX-1 | -1.33639 | 0.27277 | -0.83985 | -1.67298 | 0.37323 | -0.55341 | 3.88850 | -8.41868 | -0.52562 | Lysine-Specific Demethylase (JMJD) Inhibitor |
| 876 | 874 | 3-Deazaneplanocin A hydrochloride, DZNep, NSC 617989, NSC-617989, 3-DNPA, c3Nep | -1.44878 | 0.28493 | -0.50680 | 2.57487 | 3.58959 | 10.79924 | 5.96789 | 0.61782 | 1.38774 | S-Adenosyl-L-homocysteine Hydrolase Inhibitor, Histone-Lysine N-Methyltransferase EZH2 Inhibitor |
| 877 | 875 | KC7F2 | 1.33131 | 1.11721 | -1.09210 | 0.21190 | -5.77917 | -0.56024 | -0.07957 | 1.64851 | -0.20311 | Hypoxia Inducible Factor 1-alpha (HIF-1alpha) Inhibitor |
| 878 | 876 | BVT948, BVT-948 | -0.83478 | 0.80458 | -1.69893 | -0.09441 | -1.42829 | -0.42103 | 0.97692 | 3.57682 | 0.05587 | Protein Tyrosine Phosphatase (PTP) Inhibitor |
| 879 | 877 | NSC 87877, NSC-87877 | -0.23738 | -0.84488 | -3.18834 | 0.30773 | -0.13465 | -0.16372 | 1.43946 | 0.90557 | -0.12154 | Protein Tyrosine Phosphatase SHP-2 (PTP2C) Inhibitor, CDC25A Inhibitor, CDC25B Inhibitor |
| 880 | 878 | CS 2100, CS-2100 | -0.47535 | 0.39668 | -2.41864 | 1.51613 | -0.51590 | -1.31524 | -0.13270 | 4.52381 | 0.10014 | Lysophospholipid edg1 (S1P1) Receptor Agonist |
| 881 | 879 | ARRY 520 trifluoroacetate, ARRY-520, Filanesib | 0.98425 | -0.22311 | -9.20792 | -6.18357 | -9.30838 | -4.67123 | -8.78489 | -8.65441 | -2.92084 | Kinesin-Like Spindle Protein KIF11 (KSP, Eg5) Inhibitor |
| 882 | 880 | CCT 031374 hydrobromide | -0.15911 | 2.76064 | -0.09295 | -1.23428 | 0.01261 | -0.92701 | -0.48190 | -1.69605 | -0.11532 | TCF-dependent transcription inhibitor |
| 883 | 881 | Tazarotene, AGN-190168, Avage, Fabior, Suretin, Tazorac, Tazoral, Zorac | 0.63891 | -0.48787 | -2.58132 | -2.41308 | -5.63123 | -9.96827 | -12.64322 | -13.49301 | -2.95445 | Ornithine Decarboxylase Inhibitor, Retinoid RARgamma Agonist, Retinoid RARbeta Agonist |
| 884 | 882 | AHPN, AGN-192837, CD-437, CD 437 | -0.58348 | 0.83599 | -1.23761 | -1.06301 | -2.64481 | -7.28175 | -11.08024 | -8.21815 | -1.98361 | Retinoid RARgamma Agonist, Signal Transduction Modulator, Apoptosis Inducer |
| 885 | 883 | Am-580, AM 580, CD-336, Ro-40-6055 | -12.93181 | -13.92472 | -13.56619 | -13.14120 | -13.54302 | -12.33867 | -13.07303 | -13.50845 | -6.72516 | Retinoid RARalpha Agonist |
| 886 | 884 | CD 2314, CD-2314 | -0.29843 | 0.18623 | 0.67764 | 0.13408 | -0.80865 | -0.83965 | -2.72895 | -8.13851 | -0.74949 | Retinoid RARbeta Agonist |
| 887 | 885 | Cilengitide, EMD-121974, EMD-85189, NSC-707544 | 0.18626 | 0.85289 | 0.31811 | 1.11883 | -1.13169 | 1.35545 | 0.71256 | -0.32547 | 0.19580 | Integrin alphavbeta3 (Vitronectin) Antagonists |
| 888 | 886 | AC-261066 | -0.29825 | 0.54878 | -1.12908 | -0.91244 | -4.25469 | -7.54479 | -10.85651 | -13.55374 | -2.41034 | Signal Transduction Modulator, Retinoid RARbeta2 Agonist |
| 889 | 887 | BMS-189961, BMS 961 | -0.50114 | -1.91356 | 0.42242 | 0.22055 | -0.43221 | -2.12576 | -2.17541 | -3.32652 | -0.62361 | Retinoid RARgamma Agonists Signal Transduction Modulator |
| 890 | 888 | BMS 753 | -1.14674 | -2.88903 | -4.35784 | -8.91536 | -12.13168 | -12.82939 | -12.95294 | -13.50540 | -4.35935 | Selective Retinoid RARapha Agonist |
| 891 | 889 | ID-8 | -1.61165 | -1.08957 | -1.01396 | 0.10544 | -0.66486 | -1.73548 | -1.37382 | -1.56402 | -0.56756 | DYRK inhibitor |
| 892 | 890 | BMS-708163, Avagacestat | 0.29508 | 0.02614 | -0.66750 | 0.38062 | 1.53863 | -0.52432 | 1.90430 | -0.36813 | 0.16395 | Gamma-Secretase Inhibitor |
| 893 | 891 | WRN helicase, NSC19630, MIRA-1 | -0.51194 | -1.66650 | -0.20231 | -0.03211 | -0.03126 | 0.86669 | -0.14253 | -0.56943 | -0.14521 | DNA-Damaging Drug, Apoptosis Inducer |
| 894 | 892 | CD 2665 | -0.77007 | 1.20158 | -1.79765 | -1.45104 | -1.58143 | -7.02875 | -7.97942 | -10.20463 | -1.87821 | RARbeta antagonist, RARgamma antagonist |
| 895 | 893 | BMS 195614, BMS-614 | -0.21769 | -1.94727 | -1.73156 | -1.71597 | -3.07590 | -5.33424 | -5.93155 | -5.80453 | -1.63384 | Signal Transduction Modulators Retinoid RARalpha Antagonist |
| 896 | 894 | LE 135, LE-135 | -0.17685 | 1.79505 | 0.67690 | 0.26026 | -1.35435 | 0.99105 | 0.79884 | 0.06466 | 0.19381 | RARgamma antagonist, RXRalpha antagonist, RARbeta antagonist |
| 897 | 895 | MM 11253 | -1.41516 | -1.49580 | 0.31916 | -0.05377 | -1.66745 | -1.51577 | -1.44710 | -5.14450 | -0.78781 | RARgamma antagonist |
| 898 | 896 | RGFP966, RGFP-966 | -1.00946 | -1.01513 | -2.76009 | -0.86532 | 1.82743 | 1.51260 | -1.70069 | -5.59195 | -0.60908 | Histone Deacetylase 2 (HDAC2) Inhibitor, Histone Deacetylase 1 (HDAC1) Inhibitor, Histone Deacetylase 3 (HDAC3) Inhibitor |
| 899 | 897 | ER 50891 | -1.23124 | -2.11766 | -3.90441 | -2.28489 | -3.32396 | -3.93089 | -3.63407 | -2.93631 | -1.48191 | RARalpha antagonist, RARdelta antagonist, RARgamma antagonist |
| 900 | 898 | BMS 453, BMS-189453 | -0.08304 | -1.62851 | -4.45627 | -9.47511 | -12.80865 | -13.40039 | -12.48908 | -13.34226 | -4.29306 | Signal Transduction Modulator, Retinoid RAR Antagonist |
| 901 | 899 | Mirin | 0.37115 | -0.37233 | 0.17674 | 0.77095 | 0.52288 | -0.47068 | 1.02315 | 2.31121 | 0.27484 | Antineoplastic Enhancing Agent |
| 902 | 900 | GSK-343 | 0.03812 | 0.00820 | -0.71908 | 2.58035 | 1.08443 | 1.52943 | 3.58256 | 3.38908 | 0.72899 | Histone-Lysine N-Methyltransferase EZH2 Inhibitor, Apoptosis Inducer |
| 903 | 901 | EPZ-005687 | 0.01210 | 0.86586 | -0.42176 | 1.26617 | -0.78151 | -0.06385 | -0.01076 | -0.25899 | 0.03852 | Histone-Lysine N-Methyltransferase EZH2 Inhibitor |
| 904 | 902 | EPZ-6438, Tazemetostat, E-7438, EPZ6438 | -0.23337 | 0.60264 | -2.27803 | 2.59371 | -1.44556 | 1.72810 | 2.83933 | 3.37236 | 0.45537 | Histone-Lysine N-Methyltransferase EZH2 Inhibitor |
| 905 | 903 | SCR7 | -0.60567 | 0.55374 | -3.19144 | -0.13144 | 0.34608 | 0.49587 | -0.25999 | 0.07714 | -0.17225 | Ligase Inhibitor |
| 906 | 904 | AR-7 | 0.54562 | 1.06867 | 0.26622 | -1.00079 | -0.86520 | 0.75569 | 0.33511 | -2.05546 | -0.06027 | Signal Transduction Modulator, Retinoid RARalpha Antagonist |
| 907 | 905 | TMP269, TMP-269 | -0.52100 | 0.57181 | 0.38671 | -0.00043 | 0.06839 | -0.00570 | 0.56198 | 0.65129 | 0.10866 | Histone Deacetylase 5 (HDAC5) Inhibitor, Histone Deacetylase 4 (HDAC4) Inhibitor, Histone Deacetylase 7 (HDAC7) Inhibitor, Histone Deacetylase 9 (HDAC9) Inhibitor |
| 908 | 906 | Fluorobexarotene | -0.92795 | -0.59546 | -1.13612 | -0.54228 | -2.88172 | -6.98289 | -10.07956 | -12.78782 | -2.27924 | Retinoid RXR Agonist, Signal Transduction Modulator |
| 909 | 907 | UVI 3003 | -0.30991 | -0.67927 | 0.22699 | 0.96719 | -0.94243 | -0.27706 | -0.21229 | -2.74575 | -0.25197 | RXR antagonist |
| 910 | 908 | Compound BrBzGHSCP2 | 1.74786 | -0.63919 | -1.66916 | 0.88259 | -0.34542 | -0.74700 | 1.12477 | -0.83913 | -0.03074 | Glyoxalase I inhibitor |
| 911 | 909 | Tamsulosin HCl | 0.18424 | 0.92935 | -0.94782 | 0.67934 | -0.02729 | -0.11218 | 2.06752 | 1.34077 | 0.26094 | Signal Transduction Modulator, alpha1-Adrenoceptor Antagonist |
| 912 | 910 | MG149 | -0.79865 | -0.42796 | -0.68769 | 0.36475 | -0.11200 | 0.70565 | -0.39626 | 0.62966 | -0.04583 | Histone Acetyltransferase KAT5 (Tip60) Inhibitor |
| 913 | 911 | ARRY-520, Filanesib | 0.75961 | -3.13771 | -5.83564 | -3.97744 | -8.76152 | -7.30047 | -8.65014 | -6.11633 | -2.72868 | Kinesin-Like Spindle Protein KIF11 (KSP, Eg5) Inhibitor, Antimitotic Drug |
| 914 | 912 | STK549836 | 0.18278 | -1.34207 | 0.07249 | 1.31635 | -0.56877 | 1.94655 | 0.14602 | -12.83861 | -0.70312 | Bcl-2 Inhibitor, Signal Transduction Modulator, Apoptosis Inducer |
| 915 | 913 | Colchicine | -0.11100 | 1.18503 | 1.34693 | -2.64259 | 0.79963 | 1.00576 | -5.03303 | -1.17369 | -0.29323 | |
| 916 | 914 | LGK974, LGK-974, WNT-974 | 1.56397 | 0.20400 | -0.37137 | -0.66697 | -0.88909 | 0.71218 | 1.57982 | 0.20607 | 0.14834 | Protein-Serine O-Palmitoleoyltransferase Porcupine (PORCN) Inhibitor, Wnt Signaling Inhibitor |
| 917 | 915 | PBIT | -0.06384 | -0.18997 | -0.44726 | 1.00125 | -1.76555 | -0.38531 | -2.63672 | -0.27626 | -0.30215 | Lysine-Specific Demethylase 5B (KDM5B; JARID1B) Inhibitor, Mannose-6-Phosphate Isomerase Inhibitor |
| 918 | 916 | Piperlongumine, Piplartine | 0.27936 | -0.92553 | -2.91288 | -0.27601 | 0.28246 | -0.62707 | 0.28829 | -0.17259 | -0.25777 | Apoptosis Inducer |
| 919 | 917 | BRD-73954 | 1.05799 | 0.70806 | -0.31182 | -0.67106 | -0.70118 | 0.27994 | 0.11450 | 0.56352 | 0.06596 | Histone Deacetylase 8 (HDAC8) Inhibitor, Histone Deacetylase 6 (HDAC6) Inhibitor |
| 920 | 918 | OTX015, OTX-015, Y-803 | 0.05584 | -0.27584 | -1.38982 | -0.28450 | -2.02608 | -5.29069 | -4.82618 | -5.67858 | -1.25055 | Bromodomain-Containing Protein 2 (Brd2) Inhibitor, Bromodomain-Containing Protein 4 (Brd4, HUNK1) Inhibitor, E-Selectin Inhibitor, Bromodomain-Containing Protein 3 (Brd3, RING3-like protein) Inhibitor, VCAM-1 Antagonist |
| 921 | 919 | Rifabutin | 0.76327 | 0.05407 | -0.72448 | -0.19763 | 0.67125 | -0.06734 | -0.47095 | -0.09346 | -0.00414 | DNA-Directed RNA Polymerase Inhibitor |
| 922 | 920 | PTC-209 | -0.66965 | -0.78285 | -1.18491 | -0.26347 | -0.02285 | 1.92506 | -0.11644 | 1.11794 | 0.00018 | Polycomb Complex Protein BMI-1 Inhibitor |
| 923 | 921 | PFI-3 | 0.67726 | 0.45964 | -1.64298 | -0.31581 | 0.52009 | 0.19003 | 0.66944 | -0.27457 | 0.01796 | Drugs Targeting Bromodomain-Containing Protein |
| 924 | 922 | PFI-2 | -0.62783 | -0.55652 | -0.83474 | -0.48575 | 0.46231 | 0.12921 | -1.19325 | -0.56163 | -0.23267 | SETD7 Histone Lysine Methyltransferase Inhibitor |
| 925 | 923 | LCL161, LCL-161 | 0.55097 | -0.85089 | -0.07359 | 0.20292 | 0.45120 | -0.51786 | -0.49999 | -3.75356 | -0.28485 | Inhibitor of Apoptosis Proteins (IAP) Inhibitor, Signal Transduction Modulator, Apoptosis Inducer |
| 926 | 924 | PD151746, PD-151746 | -0.75988 | -0.56368 | 1.21151 | 1.15728 | 0.29917 | -0.98452 | 0.08507 | -1.29842 | -0.05413 | Calpain Inhibitor |
| 927 | 925 | 6H05 | 0.74575 | -0.84128 | -1.73796 | 0.49682 | 0.45119 | 0.65097 | 0.08726 | -0.16248 | -0.01965 | K-Ras(G12C) allosteric inhibitor |
| 928 | 926 | HSP990, NVP-HSP990, HSP-990 | 0.04729 | 0.88459 | 0.98449 | -5.72003 | -1.31197 | -3.86089 | -7.76771 | -9.62978 | -1.67287 | Heat Shock Protein 90 (HSP90) Inhibitor |
| 929 | 927 | BV-6 | 0.67890 | 0.11747 | -0.26719 | 0.48708 | 1.01233 | -0.61284 | 1.41143 | 11.52053 | 0.91006 | Inhibitor of Apoptosis Proteins (IAP) Inhibitor, Signal Transduction Modulator, Apoptosis Inducer |
| 930 | 928 | Santacruzamate A (CAY10683) | 0.78937 | 0.91708 | -0.07045 | 0.38195 | -1.24546 | -1.08111 | 0.83802 | -0.72398 | -0.01234 | Histone Deacetylase 2 (HDAC2) Inhibitor |
| 931 | 929 | INCB024360 | -0.44568 | 0.88237 | 0.43722 | -0.41239 | 0.17364 | 0.04935 | 0.91304 | -1.04721 | 0.03491 | Indoleamine 2,3-dioxygenase Inhibitor, Tryptophan 2,3-Dioxygenase Inhibitor |
| 932 | 930 | GSK923295, GSK-295, GSK-923295, GSK-923295A | -0.86956 | -1.75636 | -0.01530 | 0.72145 | 2.93580 | -8.90164 | -5.79165 | -7.80986 | -1.36290 | Centromere Associated Protein (CENP) Inhibitor, Antimitotic Drug |
| 933 | 931 | ML323, ML-323 | -1.42725 | -0.24734 | 0.51988 | -0.34099 | -0.23367 | 0.28208 | 0.86227 | -0.48951 | -0.06816 | DNA Repair Inhibitor, Ubiquitin Carboxyl-terminal Hydrolase Inhibitor |
| 934 | 932 | HS-173 | 0.34995 | -0.16008 | -0.29167 | 0.85400 | -0.32652 | -0.01686 | 1.24635 | 2.74799 | 0.27929 | Phosphatidylinositol 3-Kinase alpha (PI3Kalpha) Inhibitor, Signal Transduction Modulator, Apoptosis Inducer, Angiogenesis Inhibitor |
| 935 | 933 | HDAC inhib.26 | -0.25522 | -1.41723 | -0.58587 | -2.41687 | -0.82328 | 0.75512 | 4.49928 | 7.00205 | 0.42865 | Histone Deacetylase 2 (HDAC2) Inhibitors, Histone Deacetylase 1 (HDAC1) Inhibitors |
| 936 | 934 | RG-7388, RO-5503781, Idasanutlin | -0.40838 | 0.16364 | -1.38511 | -0.33486 | 0.30261 | 0.17915 | 0.11545 | -0.79095 | -0.13691 | MDM2 (hdm2) Inhibitor |
| 937 | 935 | BAZ2-ICR | 0.22039 | -0.52562 | -2.57929 | 0.64930 | -0.07239 | 0.27873 | 0.75163 | 0.79372 | -0.03067 | BAZ2 bromodomain inhibitor |
| 938 | 936 | PC190723, PC-190723 | 0.83543 | -2.37067 | 0.33985 | 1.28836 | 0.04178 | -0.82256 | -0.25164 | -0.02467 | -0.06115 | FtsZ Inhibitor |
| 939 | 937 | RVX-208, RVX-000222, Apabetalone | -1.37085 | 0.79528 | -0.14774 | 1.32936 | 0.32523 | -0.81041 | -0.58349 | -2.10405 | -0.16280 | Bromodomain-Containing Protein 4 (Brd4, HUNK1) Inhibitor, APOA1 Expression Enhancer, HDL-Cholesterol Increasing Agent |
| 940 | 938 | I-BET-762, GSK-525762, GSK-525762A, I-BET762 | -1.09114 | -1.00069 | -0.34585 | -0.82669 | -3.58366 | -3.16996 | -3.31567 | -3.69509 | -1.08011 | Bromodomain-Containing Protein 4 (Brd4, HUNK1) Inhibitor, Bromodomain-Containing Protein 2 (Brd2) Inhibitor, APOA1 Expression Enhancer, Bromodomain-Containing Protein 3 (Brd3, RING3-like protein) Inhibitor |
| 941 | 939 | Sabutoclax | -0.03045 | 0.51547 | 0.47670 | 0.84571 | -0.52845 | 0.49693 | 6.68998 | 6.50648 | 0.94968 | Mcl-1 Inhibitor, Bcl-2 Inhibitor, Bcl-xl Inhibitor, Signal Transduction Modulator, Apoptosis Inducer |
| 942 | 940 | Tasquinimod | -0.25624 | -1.14469 | -1.22380 | 0.14836 | -2.01476 | -2.33787 | -0.94636 | -3.09736 | -0.68964 | Histone Deacetylase 4 (HDAC4) Inhibitor, Angiogenesis Inhibitor, S100A9 Inhibitor |
| 943 | 941 | Compound 24a | -0.19023 | 1.65793 | 0.31977 | 0.13211 | -1.32832 | -0.09411 | -1.77519 | -5.10345 | -0.40477 | Histone Deacetylase 2 (HDAC2) Inhibitor, Apoptosis Inducer, Angiogenesis Inhibitor, Histone Deacetylase 3 (HDAC3) Inhibitor |
| 944 | 942 | OICR10295 | 0.26037 | -0.67987 | -1.04498 | 0.14750 | 0.46870 | 1.17934 | 2.17793 | -7.75474 | -0.33273 | Histone Deacetylase 2 (HDAC2) Inhibitors, |
| 945 | 943 | OICR9784, MI-192 | -0.00565 | 1.94309 | 0.11871 | -1.15901 | -0.36083 | 0.28352 | 1.18877 | -2.64295 | -0.04024 | Histone Deacetylase 2 (HDAC2) Inhibitor, Histone Deacetylase 3 (HDAC3) Inhibitor |
| 946 | 944 | Abbvie Mcl-1 Cpd 30b | -1.24099 | 0.60153 | -0.83275 | -0.44717 | 0.09464 | 0.90658 | 0.26982 | 0.35918 | -0.01834 | Mcl-1 inhibitor |
| 947 | 945 | AMG 232, AMG-232 | 0.62485 | -1.72708 | -2.09414 | 0.97668 | 0.18848 | -1.90276 | -8.31039 | -10.03763 | -1.41332 | MDM2 (hdm2) Inhibitor |
| 948 | 946 | PCI-27483 | -0.27084 | 1.88318 | -2.14140 | -0.67188 | 1.41655 | 1.17217 | 0.74614 | -0.32125 | 0.11498 | Coagulation Factor VIIa Inhibitor, Inhibitors of Blood Coagulation Pathway |
| 949 | 947 | BMS 833923, BMS-833923, BMS833923, XL-139, XL139, XL 139 | -0.56502 | -0.14596 | -0.60853 | -0.51669 | 0.10123 | -1.83606 | 0.67241 | 0.01859 | -0.18268 | Hedgehog Signaling Inhibitor |
| 950 | 948 | Abbvie Mcl-1 Cpd 30e, A-1210477 | -0.77842 | 0.82976 | 1.31647 | 0.76451 | -0.36324 | -0.18647 | 0.14687 | 1.29315 | 0.19172 | Mcl-1 Inhibitor, Signal Transduction Modulator, Apoptosis Inducer |
| 951 | 949 | Nexturastat A | -0.64867 | -0.16424 | -0.27027 | 1.03998 | -0.26079 | 1.17138 | 4.64892 | 8.00178 | 0.85743 | Histone Deacetylase 6 (HDAC6) Inhibitor |
| 952 | 950 | S3I 201, GLG-302, NSC-74859, S31-201, S3I-201 | -0.12222 | -2.50775 | -1.09550 | -0.23089 | -0.88037 | 0.15488 | -0.42874 | 0.16405 | -0.31375 | Signal Transducer and Activator of Transcription 3 (STAT3) Inhibitor, Signal Transduction Modulator |
| 953 | 951 | BG45, BG-45 | -1.45550 | -0.27309 | -0.88888 | 0.15177 | 0.68610 | 0.36706 | -0.04788 | 2.03511 | 0.03645 | Histone Deacetylase 3 (HDAC3) Inhibitor |
| 954 | 952 | AZ 505, AZ-505 | 0.13688 | -2.49257 | -0.63658 | 0.63454 | 0.21529 | 2.54858 | 0.78915 | 0.17127 | 0.08668 | N-Lysine Methyltransferase SMYD2 (HSKM-B) Inhibitor |
| 955 | 953 | SMBA 1, NSC-408730, CYD-1-87 | -0.25411 | 0.74214 | -0.95488 | 1.20155 | 0.69361 | -1.70034 | -1.46339 | -0.94851 | -0.17024 | BAX Activator, Signal Transduction Modulator, Apoptosis Inducer |
| 956 | 954 | SP 100030, SP-100030 | 1.57279 | -0.46527 | -2.06612 | -1.09251 | -1.02725 | 1.31045 | 0.20585 | 1.21441 | -0.02205 | NF-kappaB (NFKB) Activation Inhibitor, Signal Transduction Modulators Antiinflammatory Drugs |
| 957 | 955 | GSK503 | -1.13858 | 0.78064 | 1.58131 | 0.36725 | -1.70483 | 0.29020 | 1.68951 | 2.87349 | 0.30059 | Histone-Lysine N-Methyltransferase EZH2 Inhibitor |
| 958 | 956 | MN 64, MN-64 | -0.16317 | 0.51866 | -0.53709 | -0.11345 | -2.66810 | -2.50936 | -3.80781 | -4.28718 | -0.86057 | Signal Transduction Modulator, Tankyrase 1 (TNKS1; PARP5A) Inhibitor, Wnt Signaling Inhibitor |

| | A | B | C | D | E | F | G | H | I | J | K | L |
|---|---|---|---|---|---|---|---|---|---|---|---|---|
| 959 | 957 | OXF BD 02, OXFBD-02 | -0.95629 | -0.45325 | -1.06232 | -0.24199 | -0.69414 | 0.35070 | -0.79237 | -2.25202 | -0.38702 | Signal Transduction Modulator, Cyclic AMP Response Element-Binding Protein (CREB) Inhibitor, Bromodomain-Containing Protein 4 (Brd4, HUNK1) Inhibitor |
| 960 | 958 | SCH 51344 | -1.19152 | -2.43148 | -0.02946 | 1.65207 | -0.39247 | -2.08303 | -3.06258 | -2.57399 | -0.64142 | DNA-Damaging Drug, 7,8-Dihydro-8-Oxoguanine Triphosphatase (NUDT1; MTH1) Inhibitor |
| 961 | 959 | A 366, A-366 | -0.08414 | 0.27550 | -0.92341 | 2.17093 | -0.30356 | -0.63256 | -0.14690 | 0.65287 | 0.06398 | Histone-Lysine N-Methyltransferase EHMT2 (H3-K9-HMTase 3; G9a) Inhibitor |
| 962 | 960 | SGI 1027, MC-2841, S-1027, SGI-1027 | -0.22823 | -2.48228 | -0.39786 | -0.66943 | -0.44495 | 0.24132 | -1.68263 | -1.31618 | -0.44275 | DNA Methyltransferase 3A (DNMT3A) Inhibitor, DNA Methyltransferase 1 (DNMT1) Inhibitor |
| 963 | 961 | MS436, MS-436 | -1.32553 | 0.36742 | -1.18487 | -0.08684 | -1.81104 | 0.76847 | -0.69967 | -1.06339 | -0.31939 | Bromodomain-Containing Protein 4 (Brd4, HUNK1) Inhibitor |
| 964 | 962 | GSK-2830371 | -0.66360 | 0.67544 | -1.54784 | 1.93696 | 1.27376 | 1.55001 | -1.08766 | -9.11507 | -0.44261 | Protein phosphatase 1D (PP-2Cdelta; WIP1) Allosteric Inhibitor |
| 965 | 963 | (-)-Nutlin-3, Nutlin-3-(-), Nutlin-3-enantiomer A, Nutlin-3A | -0.29537 | 0.16957 | 0.26332 | 0.24581 | -1.66231 | 1.08466 | -0.60527 | 0.18994 | -0.03867 | MDM2 (hdm2) Inhibitor, Cellular Tumor Antigen p53 (TP53) Activator |
| 966 | 964 | Doxorubicin, NSC-123127, SA-033, Adriacin, Adriamycin, Adriblastina, Resmycin, Rubex | -0.98244 | 1.87293 | -0.27435 | -1.96927 | -9.24063 | -11.37554 | -10.42765 | -1.23374 | -2.13315 | DNA-Intercalating Drug, DNA Topoisomerase II alpha Inhibitor |
| 967 | 965 | FR-901464, WB-2663B, FR901464 | 0.22215 | -0.02746 | 2.18352 | -0.36406 | 1.38454 | 2.29996 | 4.51274 | 1.87060 | 0.76635 | Splicing Factor SF3b Modulator |
| 968 | 966 | Tenovin-6 | 0.23531 | 0.18445 | 0.65440 | -0.26891 | 0.39612 | 5.90142 | 6.26478 | 9.23308 | 1.43353 | NAD-Dependent Protein Deacetylase Sirtuin-2 (SIRT2) Inhibitor, NAD-Dependent Protein Deacetylase Sirtuin-1 (SIRT1) Inhibitor |
| 969 | 967 | G007-LK, G-007-LK | -0.55923 | 0.00310 | 1.47233 | 0.67582 | -1.38428 | 2.13786 | -1.12075 | -1.40985 | -0.01173 | Tankyrase 2 (TNKS2; PARP5B) Inhibitor, Tankyrase 1 (TNKS1; PARP5A) Inhibitor |
| 970 | 968 | Amgen IRE1 Inhibitor | 0.07681 | 1.22593 | 1.30843 | 1.16198 | 0.12120 | -0.20414 | 1.85235 | -1.27168 | 0.27090 | X-Box-Binding Protein 1 (XBP1) Inhibitor, Serine/Threonine Kinase/Endoribonuclease IRE1 (ERN1) Inhibitor |
| 971 | 969 | UMI-77 | 0.25712 | 0.52637 | -1.01756 | -1.85706 | 0.56446 | -0.51679 | 1.02767 | -1.25907 | -0.14429 | Mcl-1 Inhibitor |
| 972 | 970 | Maritoclax, Marinopyrrole A, (-)-Marinopyrrole A | 0.41643 | -3.97042 | -0.65859 | 1.59586 | 0.37325 | 0.08980 | 0.57400 | -0.73877 | -0.14706 | Mcl-1 Inhibitor |
| 973 | 971 | MDM2/MDMX Dual inhib., RO-5963 | 1.96841 | 0.20060 | 0.50186 | -2.33724 | -1.22602 | -1.05939 | -1.42359 | -0.83771 | -0.26723 | MDM4 (MDMX) Inhibitor, MDM2 (hdm2) Inhibitor |
| 974 | 972 | Diisopropyl Fluorophosphate (DFP), Diisopropyl Fluorophosphate, Difluorophate, Diflupyl, Diflurphate, Dyflos, Dyphlos, Fluropryl, Fluostigmine, isofluorophate, isofluorphate, Neoglaucit, PF-3, PF3, T-1703, TL 466 | 0.56670 | 1.24876 | -0.79722 | -1.68779 | 1.42636 | -0.06027 | 0.42685 | 1.18609 | 0.14649 | Acetylcholinesterase inhibitor |
| 975 | 973 | 3,4-Dichloroisocoumarin | 0.25501 | 0.67309 | -0.21251 | 0.65115 | -2.24918 | 0.63960 | -1.08258 | 0.33099 | -0.06308 | Serine protease inhibitor |
| 976 | 974 | Z-Leu-Tyr-chloromethylketone, Z-LY-CMK | 0.56253 | 1.06543 | -0.91717 | -0.18750 | -0.90403 | -0.11987 | -0.19055 | 0.02511 | -0.04225 | Calpain inhibitor, E.coli ClpP inhibitor |
| 977 | 975 | N-p-Tosyl-L-phenylalanine chloromethyl ketone, TPCK | 0.38904 | -0.65514 | -2.86141 | -2.72839 | -0.29431 | 0.05166 | 1.66627 | 1.09508 | -0.21167 | Serine Protease Inhibitor |
| 978 | 976 | UNC0642, UNC-0642 | 0.19764 | 0.91378 | -1.04354 | 0.67817 | -0.07073 | 0.42478 | -0.54654 | -0.42674 | 0.00804 | Histone-Lysine N-Methyltransferase EHMT1 (H3-K9-HMTase 5; GLP) Inhibitor, Histone-Lysine N-Methyltransferase EHMT2 (H3-K9-HMTase 3; G9a) Inhibitor |
| 979 | 977 | OICR0009429A3, OICR-9429 | 0.08393 | -0.53520 | -0.83726 | -0.36622 | 0.98818 | -0.52663 | -1.42216 | 1.70982 | -0.05744 | WD repeat-containing protein 5 (WDR5; BIG3)/MLL1 interaction Inhibitor |
| 980 | 978 | OICR0010547A1 | 0.14379 | -1.97897 | -12.81041 | 0.18588 | -0.53540 | -0.15019 | 0.00127 | 0.82875 | -0.90800 | Negative control |
| 981 | 979 | GSK2801, GSK-2801 | 0.76604 | -0.64687 | 1.77525 | -1.63140 | 2.36503 | 0.13614 | -1.51134 | -1.20824 | 0.00283 | Bromodomain Adjacent To Zinc Finger Domain Protein 2B (BAZ2B) Inhibitor, Bromodomain Adjacent To Zinc Finger Domain Protein 2A (BAZ2A) Inhibitor |
| 982 | 980 | GSK-J4 | -0.92272 | 0.98929 | -1.42744 | 0.11248 | -0.76388 | -1.95173 | -4.11531 | -9.35502 | -1.10584 | Lysine-Specific Demethylase 6A (UTX) Inhibitor, TNF-alpha Production Inhibitor, Lysine-Specific Demethylase 6B (JMJD3) Inhibitor |
| 983 | 981 | SGC0946a, SGC-946a | -0.13719 | -1.79338 | -1.67517 | -0.99045 | -0.45714 | -0.37564 | 2.56654 | 0.75309 | -0.13379 | Histone-Lysine N-Methyltransferase, H3 Lysine-79 Specific (DOT1L) Inhibitor |
| 984 | 982 | Bromosporine | -0.99506 | -2.18175 | -0.50346 | -1.54680 | -1.39233 | -1.58183 | -4.49905 | -4.94966 | -1.11951 | Cat eye syndrome critical region protein 2 (CECR2) Inhibitor, Bromodomain-Containing Protein 9 (Brd9) Inhibitor, Bromodomain-Containing Protein 4 (Brd4) Inhibitor, Bromodomain-Containing Protein 2 (Brd2) Inhibitor |
| 985 | 983 | SGC-CBP30 | 0.48132 | 0.46916 | -0.17637 | -2.53880 | 1.12703 | -1.17534 | -2.09567 | -12.00985 | -1.00969 | Cyclic AMP Response Element-Binding Protein (CREB) Inhibitor, Platelet-Activating Factor Receptor (PAFR) Ligand, alpha2C-Adrenoceptor Antagonist, alpha2A(2D)-Adrenoceptor Antagonist, Phosphodiesterase PDE5A Inhibitor |
| 986 | 984 | CPI 203, CPI-203, CPI-'203 | 0.06958 | -4.03769 | -1.82021 | -2.91395 | -7.84127 | -7.32001 | -9.17364 | -3.29079 | -2.30424 | Bromodomain-Containing Protein 4 (Brd4, HUNK1) Inhibitor, IL-6 Production Inhibitor |
| 987 | 985 | GSK 5959 | 0.83690 | -0.92068 | 1.07586 | 1.26437 | 0.05144 | 0.16930 | -1.02445 | -0.04228 | 0.08946 | Drugs Targeting Bromodomain-Containing Protein |
| 988 | 986 | I-CBP 112, I-CBP-112 | 0.61154 | 0.73227 | 0.16886 | 1.02954 | 0.21613 | -2.17076 | -0.59999 | -3.10600 | -0.19780 | Drugs Targeting specific Bromodomain-Containing Protein |
| 989 | 987 | LY 303511, LY-303511, EM-101 | 0.61650 | -0.55947 | 0.58228 | -0.52026 | 0.35331 | -0.46522 | -1.22512 | -2.90775 | -0.26169 | Casein Kinase II (CK2) Inhibitor, Mammalian Target of Rapamycin (mTOR; FRAP1) Inhibitor |
| 990 | 988 | OF 1 | 0.30621 | 0.09795 | 0.14818 | 1.26376 | -0.30941 | 1.12138 | 0.75174 | -1.33266 | 0.12985 | Drugs Targeting specific Bromodomain-Containing Protein |
| 991 | 989 | PT-2385 | 0.09087 | 0.34894 | 2.07925 | -2.24699 | -0.24550 | 0.85967 | 1.97555 | -0.50680 | 0.14937 | Hypoxia Inducible Factor 2-alpha (HIF-2alpha) Inhibitor |
| 992 | 990 | PFI 4, PFI-4 | 0.31376 | -0.18044 | -0.90145 | -0.46173 | -0.85682 | -0.14329 | 1.00751 | -2.21991 | -0.21834 | Drugs Targeting specific Bromodomain-Containing Protein |
| 993 | 991 | XD 14, XD-14 | 0.56857 | -1.10599 | 0.62809 | -0.88364 | 0.22174 | 0.20575 | -0.54661 | -0.70674 | -0.10268 | Drugs Targeting Bromodomain and Extra-Terminal (BET) Proteins |
| 994 | 992 | UNC 926 hydrochloride, UNC-926 hydrochloride | 1.37890 | 0.35029 | -0.03807 | 1.45827 | -0.28492 | -0.78053 | 1.08886 | 0.03451 | 0.20344 | Lethal(3)malignant Brain Tumor-like Protein 1 (L3MBTL1) Inhibitor, Lethal(3)malignant Brain Tumor-like Protein 3 (L3MBTL3) Inhibitor |
| 995 | 993 | EGCG, Epigallocatechin gallate, (-)-Epigallocatechin gallate, Sunphenon, Teavigo, Epigallocatechin-3-gallate, (-)-EGCG | 0.40314 | 0.65710 | 0.33522 | -1.42688 | 1.67688 | 1.08602 | 1.03406 | -0.42615 | 0.21181 | beta-Amyloid (Abeta) Aggregation Inhibitor, Proteasome Inhibitor |
| 996 | 994 | Molidustat, BAY-85-3934 | 0.06913 | -1.03910 | 0.37195 | 0.64948 | 0.73625 | -0.30304 | -0.78439 | -1.61374 | -0.12137 | Hypoxia-Inducible Factor (HIF) Prolyl Hydroxylase Inhibitor |
| 997 | 995 | Fisetin | -6.93536 | 0.72724 | -1.70362 | -2.42752 | -1.65558 | 0.23400 | -1.11478 | -4.87322 | -1.12579 | Aurora-B (ARK2) Kinase Inhibitor, NAD-Dependent Protein Deacetylase Sirtuin-1 (SIRT1) Activator, Aldose Reductase Inhibitor, Extracellular-regulated Kinase (ERK) Activator |
| 998 | 996 | PCI 34051, PCI-34051 | -4.49666 | 0.78278 | -1.10702 | -0.34887 | -0.28787 | -0.23498 | 0.89946 | 1.25078 | -0.22469 | Histone Deacetylase 8 (HDAC8) Inhibitor, IL-1beta Production Inhibitor |
| 999 | 997 | GSK LSD 1 dihydrochloride | 2.15989 | 6.82432 | -7.93701 | 2.50832 | 0.08719 | 4.91613 | 1.66459 | 1.39385 | 0.73687 | Lysine-Specific Histone Demethylase 1A (KDM1A; LSD1) Inhibitor |
| 1000 | 998 | JIB 04 | 1.24824 | 0.65300 | -0.34030 | -0.05545 | 1.02211 | 1.52394 | -5.35890 | -8.41090 | -0.61642 | |
| 1001 | 999 | RN 1 dihydrochloride, RN-1 dihydrochloride | 2.80213 | 3.42676 | 4.30464 | 4.85534 | 3.80636 | 2.58910 | 3.08202 | 2.10724 | 1.71090 | Lysine-Specific Histone Demethylase 1A (KDM1A; LSD1) Inhibitor |
| 1002 | 1000 | TC-E 5002 | -2.08984 | 0.65303 | 0.49235 | -0.82436 | 0.36766 | -0.00745 | -1.66399 | 0.10416 | -0.18828 | Lysine-Specific Demethylase 7A (KDM7A; JHDM1D) Inhibitor, Histone Lysine Demethylase PHF8 (KIAA1111; ZNF422) Inhibitor |
| 1003 | 1001 | Furamidine dihydrochloride, DB-075, DB-75 | -0.37888 | 0.34983 | -1.45182 | 0.65130 | 1.31279 | 0.08091 | -0.80369 | -0.73283 | -0.06168 | Protein Arginine N-Methyltransferase 1 (PRMT1) Inhibitor |
| 1004 | 1002 | SGC 707, SGC-707 | 0.01830 | 2.38676 | 0.25563 | -2.36322 | -0.36755 | -0.83322 | 0.35442 | -0.29702 | -0.05365 | Protein Arginine N-Methyltransferase 3 (PRMT3) Inhibitor |
| 1005 | 1003 | TC-E 5003 | -0.10893 | -2.03129 | -0.61116 | 1.62884 | 0.60929 | 0.64871 | 2.69347 | 1.30906 | 0.26247 | Protein Arginine N-Methyltransferase 1 (PRMT1) Inhibitor |
| 1006 | 1004 | FG-4592, ASP-1517, Roxadustat | -0.30444 | -0.72314 | -1.86208 | 0.92340 | -0.60383 | -1.43301 | 0.25488 | 0.49270 | -0.20649 | Hypoxia-Inducible Factor (HIF) Prolyl Hydroxylase Inhibitor |
| 1007 | 1005 | UNC 1999, UNC-1999 | -0.46307 | -0.33861 | 1.34313 | -0.65701 | 1.34480 | 1.45821 | 2.68230 | 2.36743 | 0.49076 | Histone-Lysine N-Methyltransferase EZH1 Inhibitor, Histone-Lysine N-Methyltransferase EZH2 Inhibitor |
| 1008 | 1006 | UNC 2327, UNC-2327 | 0.16469 | -2.09090 | 1.36763 | 0.78702 | 1.10641 | -0.64762 | -0.63740 | -2.64220 | -0.16443 | Protein Arginine N-Methyltransferase 3 (PRMT3) Inhibitor |
| 1009 | 1007 | MM 102, MM-102 | -0.10285 | -0.40104 | -0.56349 | -0.03899 | -0.28595 | 1.19375 | 0.81347 | 0.55822 | 0.07441 | |
| 1010 | 1008 | TC-S 7009 | 0.77407 | -0.28959 | -0.86556 | 0.36259 | -1.25131 | 0.23338 | -0.87508 | -3.11479 | -0.31881 | Hypoxia Inducible Factor 2-alpha (HIF-2alpha) Inhibitor |
| 1011 | 1009 | SirReal 2 | -0.43958 | -0.67568 | 0.95806 | 0.45731 | -1.34178 | 1.27490 | 0.55128 | 0.45020 | 0.07832 | NAD-Dependent Protein Deacetylase Sirtuin-2 (SIRT2) inhibitor |
| 1012 | 1010 | Homoharringtonine, Omacetaxine mepesuccinate, CGX-635, HHT, NSC-141633, ZJ-C, Ceflatonin, Myelostat, Omapro, Synribo, Tekinex | -1.01669 | -6.58248 | -7.31481 | -5.01926 | -11.72927 | -7.48622 | -8.45673 | -7.32723 | -3.48431 | |
| 1013 | 1011 | Simvastatin, MK-733, Synvinolin, Colemin, Denan, Lipex, Lipovas, LodalŠs, Medipo, Pantok, Simvast CR, Sinvacor, Sivastin, Zocor, Zocord | 0.74974 | 0.01616 | -2.03912 | -2.14315 | -1.75223 | -0.19802 | -0.16578 | -1.65623 | -0.45597 | HMG-CoA Reductase Inhibitor |
| 1014 | 1012 | Selinexor, KPT-330 | 0.02126 | -0.66838 | -2.63422 | -3.11418 | -5.57635 | -11.54484 | -11.16077 | -4.99234 | -2.51621 | Exportin-1 (CRM1, XPO1) Antagonist |
| 1015 | 1013 | Flubendazole | 1.33521 | 0.98044 | -1.64329 | 0.04707 | 0.29454 | -0.22604 | 0.03948 | -6.34929 | -0.35025 | Atg4B |
| 1016 | 1014 | Pitavastatin Calcium, NK-104, NKS-104, Itavastatin calcium, Nisvastatin, Alipza, Livalo, Livazo, Redevant | -0.78522 | -0.39890 | 1.32811 | -0.35258 | 1.44673 | -0.03596 | -1.21370 | -6.61351 | -0.42022 | SPP1 (Osteopontin) Expression Inhibitor, HMG-CoA Reductase Inhibitor, APOA1 Expression Enhancer |
| 1017 | 1015 | GSK2879552, GSK-2879552 | 0.63123 | 0.47398 | -0.10508 | 4.63283 | 1.11188 | 1.91615 | 5.48970 | -1.28925 | 0.81579 | Lysine-Specific Histone Demethylase 1A (KDM1A; LSD1) Inhibitor |
| 1018 | 1016 | EPZ011989, EPZ-011989 | -0.61346 | -0.51767 | 1.81939 | 1.87830 | 0.07938 | 1.87396 | 1.39270 | 2.53171 | 0.53561 | Histone-Lysine N-Methyltransferase EZH2 Inhibitor |
| 1019 | 1017 | Ischemin sodium salt | 0.71431 | 0.85720 | -0.55059 | -0.54192 | 0.65803 | 0.62914 | -0.53150 | -0.19732 | 0.06580 | |
| 1020 | 1018 | I-BRD9 | 0.88957 | -0.44898 | -2.82396 | 1.17755 | 0.57696 | 0.51354 | -1.15138 | -6.80347 | -0.51188 | |
| 1021 | 1019 | NI 57 | 1.20754 | -1.12642 | 2.43447 | -0.87497 | -0.38965 | -0.93211 | -1.66731 | -3.93908 | -0.33538 | |
| 1022 | 1020 | MI-2 (2HCl) | -0.05695 | 0.65911 | 2.57100 | 3.38605 | 5.67915 | 2.45491 | 1.73019 | -11.69961 | 0.29963 | Menin/MLL Interaction Inhibitor |
| 1023 | 1021 | CPI-360 | -0.30422 | -0.93266 | 0.41676 | -7.61058 | 1.33088 | -0.25175 | 1.47057 | 0.90056 | -0.31590 | Histone-Lysine N-Methyltransferase EZH2 Inhibitor |
| 1024 | 1022 | MK-4827, Niraparib | 0.31349 | 3.00153 | -0.79327 | 0.73690 | -0.14630 | -0.07271 | -0.91331 | -11.38003 | -0.58695 | Poly(ADP-ribose)polymerase-1 (PARP-1; ARTD1) Inhibitor, Poly(ADP-ribose)polymerase-2 (PARP-2; ARTD2) Inhibitor |
| 1025 | 1023 | CPI-169, CPI-'169 | -9.89711 | 0.02682 | 2.56988 | 0.42816 | -2.10654 | 0.64448 | -0.42207 | 2.66032 | -0.38666 | Histone-Lysine N-Methyltransferase EZH2 Inhibitor |
| 1026 | 1024 | Pontin Reptin inhibitor CPD-113 | 0.31264 | -0.61183 | 1.26487 | -0.34628 | -2.42206 | -0.88446 | -1.33276 | -0.99697 | -0.31821 | Pontin Reptin inhibitor, RuvB-Like 1/RuvB-Like 2 Complex ATPase Inhibitor |
| 1027 | 1025 | Parbendazole | 0.14834 | -0.79483 | -1.23357 | -2.96317 | -2.25038 | 0.76829 | -8.38726 | -11.72100 | -1.67665 | |
| 1028 | 1026 | Montelukast (sodium salt), Montelukast sodium, L-706631, Lukasm, MK-0476, MK-476, Kipres, Singulair | -0.91226 | -0.34612 | -1.05683 | 0.12158 | 0.69083 | -0.38716 | 1.26112 | 0.45079 | -0.01129 | Leukotriene CysLT1 (LTD4) Antagonist, Leukotriene Antagonist |
| 1029 | 1027 | EPZ-5676, Pinometostat, EPZ5676, EPZ-5676d | -0.24375 | -0.89164 | 2.44568 | -2.10734 | 0.39294 | -0.96452 | -0.51587 | 0.58154 | -0.08264 | Histone-Lysine N-Methyltransferase, H3 Lysine-79 Specific (DOT1L) Inhibitor |
| 1030 | 1028 | I-BET726, GSK1324726A, GSK-1324726A, I-BET-726 | 0.47605 | -0.42392 | -3.14539 | -2.60942 | -5.31387 | -5.88271 | -5.96999 | -6.33922 | -1.85266 | Bromodomain-Containing Protein 2 (Brd2) Inhibitor, Bromodomain-Containing Protein 4 (Brd4, HUNK1) Inhibitor |
| 1031 | 1029 | OG-L002 | -0.00333 | -0.56821 | -0.20021 | -1.16476 | 0.55109 | -0.15674 | 1.04470 | 2.16953 | 0.10606 | Lysine-Specific Histone Demethylase 1A (KDM1A; LSD1) Inhibitor |
| 1032 | 1030 | EPZ015666, EPZ-015666, GSK-3235025 | 0.91245 | 0.96354 | -0.62904 | -0.44063 | -1.49413 | -0.41234 | -0.43534 | 0.60356 | -0.05911 | Protein Arginine N-Methyltransferase 5 (PRMT5) Inhibitor |
| 1033 | 1031 | SRT2104 (GSK2245840) | 1.44646 | -0.41506 | -0.94439 | 1.24363 | 1.20159 | 1.75001 | 2.20852 | 4.36987 | 0.68888 | NAD-Dependent Protein Deacetylase Sirtuin-1 (SIRT1) Activator |
| 1034 | 1032 | SP2509, SP-2509 | 0.27827 | -0.27944 | 1.30796 | -2.15287 | 1.15509 | 0.11795 | -1.01390 | -2.25764 | -0.18043 | Lysine-Specific Histone Demethylase 1A (KDM1A; LSD1) Inhibitor |
| 1035 | 1033 | ML324, ML-324 | -0.97754 | -2.43439 | -1.71758 | 1.04864 | 0.70639 | -0.55962 | 1.81103 | 6.06332 | 0.24992 | Lysine-Specific Demethylase 4E (KDM4E; KDM4DL; JMJD2E) Inhibitor |
| 1036 | 1034 | PCI-34051 | -0.91468 | 0.99509 | -0.72713 | -0.43518 | 0.86193 | 1.30973 | 1.56070 | -11.70645 | -0.57441 | Histone Deacetylase 8 (HDAC8) Inhibitor, IL-1beta Production Inhibitor |
| 1037 | 1035 | WIKI4 | 0.20915 | -0.16727 | -0.56368 | 0.08211 | 0.20975 | 0.04079 | 0.27747 | -0.60272 | -0.03263 | Tankyrase (TRF1-Interacting Ankyrin Related ADP-ribose Polymerase) Inhibitor |
| 1038 | 1036 | IWP-L6 | 0.28569 | 0.63470 | 2.97975 | -2.05236 | -1.67848 | 0.18146 | 0.49049 | -0.48144 | 0.02282 | Protein-Serine O-Palmitoleoyltransferase Porcupine (PORCN) Inhibitor |
| 1039 | 1037 | SH-4-54, SH-04-54 | -0.35798 | -0.38396 | 1.31812 | 0.22011 | -0.56316 | -0.65578 | -0.14482 | -1.51286 | -0.13195 | Signal Transducer and Activator of Transcription 3 (STAT3) Inhibitor |
| 1040 | 1038 | PF-5274857, PF-05274857 | -0.44924 | 0.21863 | -0.46713 | 2.21419 | -0.42172 | 0.19537 | 1.77883 | 0.54642 | 0.22932 | Smoothened (Smo) Receptor Antagonist |
| 1041 | 1039 | Smoothened Agonist HCl, SAG hydrochloride | 0.33670 | 1.72192 | 2.00849 | -0.24202 | -0.93529 | -0.31299 | 1.54066 | 0.40214 | 0.28667 | Smoothened (Smo) Receptor Agonist, Neurotrophic Agent |
| 1042 | 1040 | LDE225, NVP-LDE225, Erismodegib, Sonidegib, NVP-LDE-225, Odomzo | 0.39576 | 0.32280 | -2.64396 | -1.20931 | -0.94782 | -0.17932 | -1.07318 | 0.56831 | -0.30235 | Smoothened (Smo) Receptor Antagonist |
| 1043 | 1041 | A-196 | 0.18331 | -0.78087 | 1.02058 | -1.53455 | 0.93248 | 0.30161 | 0.54484 | 0.60971 | 0.08101 | SUV420H1/2 |
| 1044 | 1042 | A-366 | 0.73228 | -0.90033 | 0.04432 | 1.33164 | 0.07719 | -0.62904 | -0.51645 | -10.25212 | -0.64142 | G9a, EHMT1 |
| 1045 | 1043 | A-395 | 0.78117 | -1.84840 | -1.50831 | 1.34457 | -0.72919 | 1.62918 | 1.75791 | 0.74000 | 0.13745 | EED |
| 1046 | 1044 | BAY-598 | 1.01595 | 0.23576 | -1.23504 | -0.24711 | 0.67494 | 1.06318 | 0.67903 | 2.80024 | 0.31632 | SMYD2 |
| 1047 | 1045 | BAZ2-ICR | 0.51919 | 0.81102 | -1.10722 | -2.23893 | 0.04616 | 0.75521 | 0.41605 | 0.72918 | -0.00440 | BAZ2A/B |
| 1048 | 1046 | BI-9564 | 0.40249 | 0.17553 | 2.25991 | 0.21000 | -2.18675 | 0.88036 | -0.09435 | 0.57827 | 0.14116 | BRD9/7 |
| 1049 | 1047 | GSK2801 | -0.74256 | -0.28158 | -1.84161 | -3.04284 | 1.63418 | 0.23571 | 0.43856 | -2.18637 | -0.36703 | BAZ2A/B |
| 1050 | 1048 | GSK343 | -0.10750 | 0.22487 | 1.67909 | -3.01816 | -1.03478 | 0.19742 | -2.81820 | -2.02641 | -0.43789 | EZH2 |
| 1051 | 1049 | GSK484 (hydrochloride) | -0.76349 | 0.62687 | -0.58291 | -0.22901 | 2.10880 | -0.55930 | 2.21301 | 4.28994 | 0.45059 | PAD4 |
| 1052 | 1050 | GSK591 | -0.68947 | 0.59847 | -3.34374 | -0.39544 | -0.33493 | 0.11989 | 0.77337 | 1.17432 | -0.13304 | PRMT5 |
| 1053 | 1051 | GSK864 | -0.29326 | -0.37996 | 2.13531 | 1.55465 | -1.81219 | -0.24749 | 0.23058 | -1.54285 | -0.02253 | IDH1 mutant |
| 1054 | 1052 | GSK-J4 (hydrochloride) | 0.93158 | 0.40819 | -0.53230 | 0.33906 | 0.50400 | -0.20987 | 0.72179 | 1.53550 | 0.23456 | JMJD3, UTX, JARID1B |
| 1055 | 1053 | GSK-LSD1 (hydrochloride) | -0.43346 | 0.41793 | -1.81908 | 1.30658 | -0.21458 | -0.69877 | -0.72869 | 0.26279 | -0.12098 | LSD1 |
| 1056 | 1054 | I-BRD9 | 0.29241 | 0.11371 | -2.28314 | 1.08407 | -6.35220 | -0.35678 | -1.94103 | -10.66648 | -1.27552 | BRD9 |
| 1057 | 1055 | I-CBP112 (hydrochloride) | 2.75083 | 0.28660 | -0.40278 | 3.17828 | 3.96538 | 0.43147 | -1.19370 | 0.67976 | 0.61499 | CBP, EP300 bromo |
| 1058 | 1056 | (+)-JQ1 | -7.99040 | 0.06151 | -0.87899 | -1.79012 | -7.77877 | -7.53898 | -10.58238 | -0.80501 | -2.36609 | BET (BRD2, BRD3, BRD4, BRDT) |
| 1059 | 1057 | MS023 (hydrochloride) | 0.14078 | 0.26483 | -1.35138 | -2.65274 | 0.27706 | 1.65513 | 2.87476 | 7.37664 | 0.54454 | PRMT type I |
| 1060 | 1058 | MS049 (hydrochloride) | 0.21730 | 0.69245 | 0.15584 | 1.22348 | 0.39443 | 0.54714 | -1.29991 | -0.15593 | 0.11257 | PRMT4/6 |
| 1061 | 1059 | NI-57 | -8.91335 | 0.90066 | 2.06264 | 1.71376 | 0.02349 | 1.11820 | 2.71526 | -1.00629 | -0.08789 | BRPF1, BRPF2, BRPF3 |
| 1062 | 1060 | NVS-CECR2-1 | 0.69425 | -1.02936 | 1.13158 | -0.45980 | -2.54166 | 0.64619 | -0.51172 | -6.02994 | -0.51380 | CECR2 |
| 1063 | 1061 | OF-1 | 0.53551 | -1.28928 | 0.98221 | -3.12029 | -3.96505 | -0.44480 | -0.42119 | 1.34412 | -0.40460 | BRPF1, BRPF2, BRPF3 |
| 1064 | 1062 | OICR-9429 | -0.14882 | -0.26926 | -0.50505 | 1.59895 | 1.13748 | 0.49081 | 1.08154 | -0.43405 | 0.18722 | WDR5 |
| 1065 | 1063 | (R)-PFI-2 (hydrochloride) | -0.74760 | -0.99239 | 1.86633 | -0.72634 | 0.52005 | 0.03470 | 0.52375 | 5.09106 | 0.35327 | SETD7 |
| 1066 | 1064 | PFI-3 | 0.31695 | 0.03858 | 1.45673 | 1.53233 | -0.11733 | 0.89593 | 1.15925 | 0.13470 | 0.34360 | Smarca2 Smarca4, PB1 |
| 1067 | 1065 | PFI-4 | 0.23141 | 0.13177 | 1.60323 | -0.81764 | -3.32807 | -4.08608 | -5.07847 | -5.45440 | -1.06549 | BRPF1B |
| 1068 | 1066 | SGC0946 | 0.01893 | -0.01149 | -0.17357 | -1.41579 | -0.80186 | -0.33448 | 0.85334 | 0.39838 | -0.09302 | Dot1L |
| 1069 | 1067 | SGC707 | -0.81720 | -1.33903 | -11.07602 | -0.44073 | 0.97332 | 0.13060 | -0.93192 | 1.44598 | -0.76463 | PRMT3 |
| 1070 | 1068 | SGC-CBP30 | -0.79163 | 0.50126 | -1.06258 | 0.13260 | 0.15493 | -0.55939 | -1.58341 | -1.02020 | -0.26820 | CBP, EP300 bromo |
| 1071 | 1069 | TP-064 | 0.61533 | -0.54591 | -2.03041 | -1.08923 | -1.01197 | 0.39838 | -1.58277 | 0.77810 | -0.28343 | PRMT4 |
| 1072 | 1070 | UNC0642 | 0.18452 | 0.85231 | 0.37507 | -2.07829 | 1.76361 | 1.70334 | -0.35007 | 0.74133 | 0.20245 | G9a, EHMT1 |
| 1073 | 1071 | UNC1215 | 0.00923 | 1.78251 | -1.66625 | -0.32788 | -0.72162 | -1.15118 | -0.53930 | 1.02352 | -0.10091 | L3MBTL3 |
| 1074 | 1072 | UNC1999 | -0.26662 | 0.95499 | -1.20805 | 0.61203 | 0.79312 | 2.12714 | 2.63524 | 3.67496 | 0.59133 | EZH2, EZH1 |
| 1075 | 1073 | Acitretin | -0.69585 | 0.07789 | -1.80517 | -1.65977 | -9.32919 | -13.47065 | -13.50330 | -13.48896 | -3.41722 | Non-selective dopamine receptor antagonist |
| 1076 | 1074 | Amoxapine | -1.18207 | -0.53858 | 0.18577 | -0.41621 | -1.96069 | -0.87960 | 1.00588 | 0.87439 | -0.18465 | Opioid receptor agonist |
| 1077 | 1075 | Aripiprazole | -0.69618 | 0.60966 | 0.94464 | -1.10885 | -0.73163 | 1.28773 | 1.26615 | -8.10170 | -0.41420 | Retinoic acid receptor agonist |
| 1078 | 1076 | Brexpiprazole | -0.11953 | 2.34593 | 80.53561 | 1.42358 | -0.95845 | 0.11738 | -0.95768 | 0.22212 | 5.23978 | Vanilloid receptor agonist |
| 1079 | 1077 | Bromocriptine | 1.05033 | 0.47688 | 0.58828 | 0.56591 | -2.26609 | 0.95193 | 0.42419 | -0.13283 | 0.10520 | D2 dopamine receptor partial agonist |
| 1080 | 1078 | (ñ) Butaclamol | -0.75614 | -1.13218 | -1.67571 | -0.39300 | 0.50630 | -0.69854 | 1.99119 | -1.58051 | -0.23713 | Dopamine receptor antagonist |
| 1081 | 1079 | Capsazepine | 0.45433 | 1.64361 | -2.74977 | 1.89073 | -0.27875 | 0.22301 | -0.70170 | 1.82819 | 0.14650 | D2 dopamine receptor partial agonist |
| 1082 | 1080 | Carbetapentane | -0.70276 | -0.97038 | 0.01664 | 1.13488 | 0.26765 | 2.41276 | 5.40751 | 3.79850 | 0.72085 | Dopamine receptor agonist |
| 1083 | 1081 | cis-(Z)-Flupenthixol | 1.14739 | -1.34453 | 2.33296 | -0.30452 | 2.90361 | 4.03162 | 2.68223 | -1.15916 | 0.65265 | Dopamine receptor agonist |
| 1084 | 1082 | Dihydrocapsaicin | 0.17845 | 1.26639 | -1.41555 | -1.54133 | -1.56947 | 1.74856 | 0.94437 | 0.69920 | 0.01970 | D2/D3 dopamine receptor agonist |
| 1085 | 1083 | Domperidone | -0.06092 | 1.57881 | -9.40569 | 0.82380 | -0.53855 | 1.77369 | 0.41251 | -0.44536 | -0.37180 | Non-selective dopamine receptor agonist |
| 1086 | 1084 | Droperidol | -0.05613 | 0.47054 | -2.86060 | 0.94638 | 1.81405 | -0.17520 | -0.84705 | -2.80140 | -0.22260 | Non-selective dopamine receptor antagonist |
| 1087 | 1085 | Fenretinide | 0.32848 | 0.59999 | -0.95123 | -2.64392 | -1.88880 | -2.59217 | -7.56175 | -4.21544 | -1.20038 | Non-selective dopamine receptor antagonist |
| 1088 | 1086 | Metoclopramide | -0.31645 | 1.54846 | -2.01168 | 0.20301 | 1.42590 | 2.24425 | 0.22653 | 0.23502 | 0.22549 | D2/D3 dopamine receptor antagonist |
| 1089 | 1087 | N-Arachidonoyl dopamine | -0.74024 | 0.97482 | -0.11886 | 2.81277 | 0.03621 | -0.22230 | 1.32362 | 6.81533 | 0.69019 | D2 dopamine receptor antagonist |
| 1090 | 1088 | Olanzapine | 0.72008 | -0.30234 | 2.37054 | -0.78671 | 0.31008 | -0.83747 | -0.21487 | -0.97932 | 0.01776 | D2 dopamine receptor antagonist |
| 1091 | 1089 | Pergolide | 0.11993 | 0.50315 | -0.79610 | 0.18569 | 0.16808 | -0.44006 | -2.48843 | 0.33418 | -0.15309 | Retinoid receptor antagonist |
| 1092 | 1090 | Pramipexole | 0.51434 | -1.77504 | -1.84385 | 1.12143 | 2.66707 | 0.53225 | 0.61079 | 0.78145 | 0.16545 | Vanilloid receptor agonist |
| 1093 | 1091 | Ropinirole | 0.45052 | 1.56151 | -0.79027 | -2.19049 | -1.20426 | 0.05693 | 0.24357 | 1.12878 | -0.04717 | Vanilloid receptor antagonist |
| 1094 | 1092 | BAY-299 | 0.80310 | 0.77029 | 1.50474 | 0.74232 | -0.46283 | 0.98649 | 0.09166 | -1.80489 | 0.16687 | BRD1, TAF1 |
| 1095 | 1093 | GSK6853 | -0.88128 | 1.85730 | -2.21603 | 0.39325 | -1.71628 | -0.40797 | 0.26517 | -0.30758 | -0.19114 | BRPF1 |
| 1096 | 1094 | TP-472 | -4.19531 | 0.43330 | -0.11653 | 0.22073 | -0.70714 | 0.18972 | -0.51841 | 0.18476 | -0.28599 | BRD9, BRD7 |

| | A | B | C | D | E | F | G | H | I | J | K | L |
|---|---|---|---|---|---|---|---|---|---|---|---|---|
| 1097 | 1095 | (+)-?-Tocopherol acetate | -0.15867 | 0.81775 | -0.75751 | 0.48569 | -0.60039 | -2.75351 | -0.58516 | -0.67387 | -0.26803 | Antioxidant - Vitamin E |
| 1098 | 1096 | 2-DG | -0.74975 | -1.19459 | 3.22181 | 1.83241 | 0.66153 | 0.57123 | -1.56012 | -10.11053 | -0.46481 | Glucose analogs |
| 1099 | 1097 | 4μ8C (IRE1 inhibitor III) | -1.18911 | 0.41796 | -0.13547 | 0.51361 | -0.04591 | 0.09521 | 1.12371 | 1.22715 | 0.12731 | IRE1 inhibitor |
| 1100 | 1098 | A-106 (IRE1 inhibitor IV) | -0.94087 | 1.27893 | -2.00950 | -1.64125 | -1.85690 | -0.27704 | -0.93947 | -3.43559 | -0.62298 | IRE1 inhibitor |
| 1101 | 1099 | AH6809 (MW = 298.29) | 0.24553 | 1.42858 | -2.12390 | 2.30838 | 0.57190 | -1.51715 | -1.79712 | -1.67462 | -0.16228 | Prostaglandin receptor inhibitor |
| 1102 | 1100 | Antimycin | 0.02781 | 1.57769 | 0.04970 | 0.13622 | -3.03236 | -2.34405 | -1.98879 | -6.13428 | -0.74263 | ETC Complex III inhibitor |
| 1103 | 1101 | Apocynin (acetovanillone) | 0.76826 | 1.21522 | -2.05094 | -1.54890 | 0.51579 | -0.29268 | -0.29197 | -2.24336 | -0.24919 | NADPH oxidase inhibitor |
| 1104 | 1102 | Atorvastatin | -0.63095 | -0.57579 | -0.36960 | -4.24828 | 1.68052 | 3.58283 | -2.21753 | -7.37345 | -0.64394 | HMG-CoA reductase inhibitor |
| 1105 | 1103 | Bafilomycin | -0.06866 | 1.88686 | -6.86186 | -3.99338 | -7.42765 | -5.95330 | -7.14280 | -1.22295 | -1.95257 | Autophagy inhibiitor |
| 1106 | 1104 | BAM15 | -0.01800 | -0.05965 | -2.01879 | 0.06595 | -0.73800 | -0.54117 | -2.88854 | -7.65405 | -0.87863 | Mitochondrial uncoupler |
| 1107 | 1105 | Bezafibrate | -6.09816 | 1.02264 | 2.81124 | 0.38287 | -0.70888 | -4.50511 | -0.74600 | -1.48187 | -0.59136 | PPARa agonist |
| 1108 | 1106 | BPTES | 0.65863 | 1.52979 | -0.22700 | 3.13250 | 0.43405 | 2.95597 | 2.34855 | 4.44368 | 0.96895 | Glutaminase inhibitor |
| 1109 | 1107 | Bromopyruvic acid | -4.08245 | -1.42881 | 2.14052 | -0.15471 | 1.53636 | 0.59450 | -12.27657 | -1.39503 | -0.95563 | Hexokinase II inhibitor, GAPDH inhibitor |
| 1110 | 1108 | Dinitrophenol | -0.45707 | -1.30124 | 2.66822 | -0.02958 | -0.45566 | 0.83268 | -11.24577 | -8.42381 | -1.16786 | Mitochondrial uncoupler |
| 1111 | 1109 | Ebselen | 0.21058 | 0.36623 | -0.87367 | -0.90416 | -0.46470 | -1.74262 | -1.98780 | -3.10524 | -0.53923 | Antioxidant |
| 1112 | 1110 | Echinomycin | -3.18347 | -8.51523 | -9.24451 | -0.91514 | -7.61343 | -8.24550 | -9.52578 | -4.71648 | -3.29573 | HIF-1 inhibitor |
| 1113 | 1111 | Etomoxir | 0.52648 | 1.18803 | -0.28845 | 1.01247 | 0.90545 | 0.36986 | 1.47160 | -0.06000 | 0.32510 | Carnitine palmitoyltransferase 1 inhibitor |
| 1114 | 1112 | FCCP | -0.25848 | -2.71556 | -1.27620 | -3.04220 | -6.75233 | -9.57155 | -9.12625 | -4.19371 | -2.34282 | Mitochondrial uncoupler |
| 1115 | 1113 | Glycogen Phosphorylase Inhibitor | -0.19288 | -0.62280 | -1.97793 | -2.14246 | -2.46388 | -4.65470 | -0.79435 | -9.61236 | -1.42470 | Glycogen Phosphorylase Inhibitor |
| 1116 | 1114 | GSK621 | -0.50538 | 2.54099 | 0.75699 | 0.38870 | -0.56020 | 0.38308 | -1.55731 | -2.58247 | -0.07203 | AMPK activator |
| 1117 | 1115 | Hydrochloroquine | 0.30740 | -1.17684 | -1.39974 | -0.21669 | 0.14484 | -1.72329 | -3.68207 | -1.40264 | -0.58031 | Autophagy inhibitor, heme aggregator |
| 1118 | 1116 | Kinetin | 0.64622 | 0.01429 | 3.77267 | -1.89867 | 1.48668 | 0.52962 | 0.14355 | -1.80606 | 0.18320 | PINK1 activator |
| 1119 | 1117 | Kinetin Riboside | 1.04661 | 0.79534 | 0.31703 | -3.41022 | -11.47011 | -10.52286 | -8.11199 | -8.14225 | -2.50534 | PINK1 activator |
| 1120 | 1118 | LB100 | -1.02429 | -0.01587 | -0.67503 | 0.69134 | 1.27057 | 0.37189 | -8.00279 | -9.75816 | -1.08732 | PP2A inhibitor |
| 1121 | 1119 | LCS-1 (SOD inhibitor) | 1.10021 | -1.16294 | 0.90706 | -0.35600 | -5.36226 | -8.28350 | -3.13085 | -7.69269 | -1.52108 | SOD1 inhibitor |
| 1122 | 1120 | Linezolid | 0.60403 | 0.36265 | 1.25126 | -1.78970 | -0.36597 | -0.79896 | -0.68643 | -2.20840 | -0.23034 | Mitochondrial translation inhibition |
| 1123 | 1121 | LMK-235 | 1.95002 | 2.31362 | 0.54827 | 2.27982 | -11.15571 | -11.15627 | -9.23617 | -6.76465 | -1.98031 | HDAC4 and HDAC5 inhibitor |
| 1124 | 1122 | Lonidamine | 0.50100 | -0.12424 | 1.23202 | -0.82634 | -1.33377 | -0.28317 | 1.23104 | 3.06142 | 0.21933 | Mitochondrial hexokinase inhibitor |
| 1125 | 1123 | M1-Hydrazone | 0.55574 | -1.22691 | 1.31272 | -2.38248 | -1.29724 | -1.73162 | -4.36700 | -0.61665 | -0.61865 | Activates mitochondrial fusion |
| 1126 | 1124 | Mdivi-1 | -1.34473 | -1.43226 | -1.32368 | -3.58742 | -2.81685 | -6.36060 | -5.34704 | -6.86017 | -1.84405 | DRP-1 inhibitor |
| 1127 | 1125 | Meclizine | -0.81722 | 0.66925 | 7.05253 | 7.08453 | 2.31287 | -10.10246 | -0.32396 | -7.60551 | -0.10973 | Antihistamine H1 antagonist |
| 1128 | 1126 | Meloxicam | -0.24900 | -0.24314 | 2.69018 | 0.11312 | 0.54422 | -0.30185 | 0.36011 | -2.01475 | 0.05702 | NSAID |
| 1129 | 1127 | MHY1485 | 0.77922 | 0.19317 | -0.53674 | 0.29586 | -3.62284 | -2.12226 | -0.36968 | -1.30302 | -0.42410 | mTOR activator |
| 1130 | 1128 | MnTBAP | -0.22360 | 0.68628 | -1.42886 | -0.62797 | 0.93843 | 1.44277 | -0.06069 | -2.63128 | -0.12083 | Superoxide dismutase (SOD) mimetic and peroxynitrite scavenger. |
| 1131 | 1129 | NAMPT inhibitor (FK866 compound) | -1.05476 | -3.94863 | -7.54593 | -7.77202 | -12.96456 | -10.41215 | -9.01618 | -9.85052 | -3.96840 | Nicotinamide phosphoribosyltransferase (NAMPT) inhibitor |
| 1132 | 1130 | Nicotinamide | 0.32439 | -0.11915 | 3.56082 | 0.41979 | -0.84941 | 1.35585 | -0.96860 | -0.26685 | 0.21926 | Vitamin B3 |
| 1133 | 1131 | Niclosamide | -1.17223 | -0.74271 | -1.52012 | -2.84377 | -2.81442 | -3.01153 | -4.68944 | -9.02033 | -1.63738 | Antihelminth |
| 1134 | 1132 | Nicotinic acid | 0.62592 | 0.51633 | -0.00123 | -0.68263 | 1.26219 | 0.52156 | 0.24734 | 1.29994 | 0.24036 | Vitamin B3 |
| 1135 | 1133 | NU7441 | -0.71597 | 0.18376 | -0.66330 | 0.04521 | -0.71301 | -2.35015 | -5.86977 | -1.70844 | -0.74793 | DNA-PK inhibitor |
| 1136 | 1134 | Oligomycin | -10.03410 | -8.31833 | -8.34915 | -6.88912 | -9.49621 | -6.35759 | -3.53066 | -7.00470 | -3.80444 | ATPase inhibitor |
| 1137 | 1135 | PFK15 | -0.64034 | -1.90886 | -1.98280 | -1.93098 | -2.36341 | -10.77324 | -5.12389 | 0.95597 | -1.50755 | 6-phosphofructo-2-kinase (PFKFB3)ÿinhibitor |
| 1138 | 1136 | Phenformin | -0.67230 | -0.37028 | -1.36020 | -0.32394 | -5.00117 | -1.36070 | -7.11026 | -7.76247 | -1.51984 | ETC Complex I inhibitor |
| 1139 | 1137 | Rotenone | -0.35587 | 1.23266 | -0.45886 | -1.88932 | -2.43717 | -1.22062 | -2.24506 | -9.61099 | -1.07735 | ETC Complex I inhibitor |
| 1140 | 1138 | SAR-405 | 0.22245 | -0.47727 | 1.57125 | -0.58463 | -1.00341 | -1.12675 | -9.41155 | -0.91846 | -0.74391 | ÿPI3K class III isoform Vps34 inhibitor |
| 1141 | 1139 | SB203580 | -0.99170 | 1.12829 | -2.11058 | 0.21486 | -3.76953 | -2.84007 | -3.08533 | 1.00023 | -0.66307 | p38 MAPK |
| 1142 | 1140 | SBI-0206965 | -0.55688 | -0.72756 | -1.40453 | -2.02111 | -2.35581 | -4.26075 | -8.51978 | -6.82836 | -1.69195 | ULK1 inhibitor |
| 1143 | 1141 | Spautin-1 | -0.14028 | -1.75238 | -1.75506 | -2.88392 | -5.12906 | -4.87299 | -3.71461 | -3.56823 | -1.51065 | Autophagy inhibitor |
| 1144 | 1142 | Tigecycline | -0.36412 | -0.36256 | -0.45575 | 0.76601 | -0.22224 | 0.08938 | -1.62064 | 1.05210 | -0.07090 | Mitochondrial translation inhibition |
| 1145 | 1143 | Torin-1 | 0.74159 | -0.73636 | -1.24970 | -4.05107 | -2.79268 | -2.04186 | -5.02735 | -0.00378 | -0.96166 | mTOR inhibitor |
| 1146 | 1144 | Tranilast | -0.55899 | 0.31932 | 0.21876 | -0.62508 | 0.35469 | -1.61257 | -2.02423 | -3.50077 | -0.47120 | Aryl hydrocarbon receptor activator, Anti allergic |
| 1147 | 1145 | Trolox | -0.67739 | -1.25603 | -0.86265 | -1.13726 | -0.47851 | -1.50860 | -1.35676 | -2.39069 | -0.61322 | Antioxidant |
| 1148 | 1146 | UK5099 | -0.65340 | -0.32211 | -0.81203 | 0.13701 | -0.99295 | 2.79044 | 4.51476 | 4.79594 | 0.59989 | Mitochondiral pyruvate carrier inhibitor |
| 1149 | 1147 | Urolithin A | -0.84915 | -0.08157 | -1.03614 | -0.55977 | -0.77445 | 2.42602 | -8.15168 | -7.81393 | -1.06818 | Mitophagy inducer |
| 1150 | 1148 | Verapamil | 0.91542 | -0.30482 | 0.03024 | 0.05598 | -0.94537 | 1.23883 | -1.36730 | -5.95527 | -0.40165 | Pgp, Ca Channel inhibitor |
| 1151 | 1149 | 6-Benzylaminopurine | 0.00312 | -0.03764 | -1.45736 | 0.03343 | -1.27740 | -2.97767 | -6.15795 | 0.45612 | -0.72406 | Synthetic cytokinin |
| 1152 | 1150 | Artesunate | 0.55465 | -0.18722 | 2.21643 | 5.62420 | -5.91397 | -10.30201 | -10.35259 | -9.33638 | -1.75678 | Anti-malarial drug |
| 1153 | 1151 | AZD3965 | 1.01553 | 0.38553 | 0.28872 | 0.49392 | 0.15913 | 1.04054 | 0.13879 | -0.57596 | 0.18687 | Monocarboxylate transporter 1 inhibitor |
| 1154 | 1152 | Bisacodyl | -0.23300 | 0.36318 | 1.01224 | 0.01721 | -1.20564 | 0.68386 | -0.24780 | -9.24594 | -0.56172 | Laxative |
| 1155 | 1153 | Brinzolamide | -0.84554 | -0.01105 | -0.64357 | -0.29095 | -0.89163 | -0.45172 | -1.79254 | -2.22760 | -0.45381 | Carbonic anhydrase II inhibitor |
| 1156 | 1154 | BTB06584 | 0.05032 | 0.16206 | -0.38358 | -1.84445 | -0.18154 | 0.86779 | -1.12810 | -4.77972 | -0.45905 | Mitochondiral F1 Fo-ATPase inhibitor |
| 1157 | 1155 | CB-839 | 2.17005 | 0.21489 | 2.46960 | 4.02569 | 2.46389 | 3.85624 | 2.41803 | 3.96910 | 1.36927 | Glutaminase inhibitor |
| 1158 | 1156 | CP-91149 | 0.08534 | -1.01075 | 0.30371 | -0.71038 | 0.51317 | -0.94126 | -2.02567 | -1.99438 | -0.36663 | Glycogen Phosphorylase Inhibitor |
| 1159 | 1157 | CPI-613 | -0.02731 | -0.94693 | -0.08910 | 0.56538 | -0.17659 | -0.76905 | 1.07042 | 1.26377 | 0.05649 | Pyruvate dehydrogenase & alpha-ketoglutarate dehydrogenase inhibitor |
| 1160 | 1158 | Dexrazoxane HCl | -0.62299 | 1.06645 | 2.65696 | 0.96739 | 4.57484 | -3.73146 | -9.28416 | -1.12734 | -0.34888 | Intracellular iron chelator |
| 1161 | 1159 | Disulfiram | 2.48073 | 1.75353 | 1.02681 | 1.45290 | 1.91530 | 3.02648 | -7.79754 | -8.88208 | -0.31866 | ALDH1 inhibitor |
| 1162 | 1160 | Dynasore | -1.18584 | -1.35217 | -2.27723 | -3.35240 | -1.84963 | -7.02313 | -6.88649 | -1.29572 | -1.59984 | Dynamin 1/2 inhibitor |
| 1163 | 1161 | Epacadostat | -0.80188 | -0.93038 | -1.29869 | 0.03670 | 0.42213 | 0.00920 | -0.11641 | -2.22445 | -0.31104 | IDO1 inhibitor |
| 1164 | 1162 | Erastin | -1.12406 | -2.38617 | -2.49301 | -4.24464 | -2.07290 | -2.00785 | -5.89482 | -9.74763 | -1.90103 | Ferroptosis activator |
| 1165 | 1163 | EUK 134 | 1.59690 | 1.02765 | 0.11881 | -0.09063 | 0.45031 | 0.75364 | -7.62713 | -3.51545 | -0.46213 | SOD/catalase mimetic |
| 1166 | 1164 | Fenofibric acid | -1.44745 | 0.33090 | 0.09524 | 0.28214 | -0.41773 | 0.86892 | 1.51534 | -1.08123 | 0.00927 | Fibrate, Targets PPARa |
| 1167 | 1165 | Flufenamic acid | 0.74403 | 1.68673 | -0.75439 | 0.54805 | -0.12214 | -0.40214 | -0.41511 | -4.30738 | -0.19170 | Ion channel modulator |
| 1168 | 1166 | GKT137831 | -0.70861 | -0.93205 | -0.35274 | 0.33343 | -1.13214 | 0.41562 | -2.30443 | -2.92724 | -0.48258 | NADPH oxidase inhibitor |
| 1169 | 1167 | GMX1778 | -1.01584 | -1.31125 | -7.60220 | -9.94666 | -11.96399 | -10.99709 | -11.93738 | -9.22580 | -4.05945 | NAMPT inhibitor |
| 1170 | 1168 | GW9662 | -1.35218 | 0.29853 | -0.62650 | -0.66986 | -0.65256 | 1.77820 | -4.43188 | -8.52334 | -0.89939 | PPARgamma antagonist |
| 1171 | 1169 | JNJ-1661010 | -0.38332 | -0.68549 | 0.62825 | -1.25042 | -1.90713 | -0.71950 | -2.47000 | -1.22243 | -0.50807 | FAAH inhibitor |
| 1172 | 1170 | Leflunomide | -1.65741 | -2.30083 | -3.23488 | -2.20521 | -1.06686 | -3.56483 | -9.56331 | -4.06464 | -1.75431 | Pyrimidine synthesis inhibitor |
| 1173 | 1171 | MK-8245 | -1.09453 | -0.84816 | 0.91711 | -0.73514 | 0.04236 | -0.15653 | -0.45956 | -1.73305 | -0.25800 | Stearoyl-CoA desaturase inhibitor |
| 1174 | 1172 | MRT68921 | -0.31843 | 0.99755 | 0.24318 | 0.05203 | -1.55177 | -10.87434 | -8.97335 | -4.37315 | -1.57292 | ULK1/2 inhibitor |
| 1175 | 1173 | NCT-501 | -1.27881 | 0.44367 | 1.16083 | 0.76680 | 0.55129 | -0.06022 | 1.40156 | 0.23290 | 0.20412 | ALDH1A1 inhibitor |
| 1176 | 1174 | NLG919 | 0.10394 | -0.72616 | 0.61293 | -1.30753 | -2.56241 | -2.83576 | -6.29650 | -6.86059 | -1.26046 | IDO pathway inhibitor |
| 1177 | 1175 | Oltipraz | 0.46125 | -0.90384 | -0.37442 | -4.29469 | -8.34847 | -3.51138 | -3.93397 | -4.51465 | -1.61237 | Nrf2 activator |
| 1178 | 1176 | Omaveloxolone | 0.52331 | 4.29261 | 6.35156 | 9.21360 | 2.53215 | -7.76601 | -4.39328 | -1.21854 | 0.60482 | Nrf2 activator |
| 1179 | 1177 | PX-12 | -1.06810 | -0.53002 | -1.18261 | -0.10507 | -1.45236 | -2.69222 | -11.24232 | -1.96768 | -1.28382 | Thioredoxin-1 inhibitor |
| 1180 | 1178 | Ro 61-8048 | 0.58171 | -0.97686 | 0.26559 | 2.79635 | -0.98326 | -0.58035 | -1.83819 | -5.48965 | -0.39482 | Kynurenine 3-hydroxylase inhibitor |
| 1181 | 1179 | Rosiglitazone maleate | -1.11622 | 0.17975 | -1.59415 | -0.34008 | 0.90002 | -0.77862 | -1.30028 | -6.32912 | -0.65831 | PPAR-gamma agonist |
| 1182 | 1180 | RSL3 | 1.29543 | 0.88722 | 2.18088 | 3.94965 | -11.67960 | -14.18111 | -0.50060 | -4.54640 | -1.43314 | Ferroptosis activator |
| 1183 | 1181 | SirReal2 | -0.52214 | -0.30731 | -1.06399 | 0.45833 | -0.64719 | -1.44952 | -0.77634 | -0.43852 | -0.30108 | SIRT2 inhibitor |
| 1184 | 1182 | Sirtinol | 0.74793 | -1.06878 | 0.35242 | 0.00180 | -0.17799 | -1.81793 | -2.45820 | -1.28100 | -0.36165 | SIRT1 and SIRT2 inhibitor |
| 1185 | 1183 | SRT1720 | 1.64205 | -1.99225 | -1.37637 | -1.01033 | -1.52864 | -6.48777 | -11.03265 | -4.64813 | -1.67668 | SIRT1 activator |
| 1186 | 1184 | STF-083010 | 0.79276 | -0.77957 | -0.61959 | -2.02728 | -1.15172 | -0.50401 | -2.59868 | -9.13583 | -1.01638 | IRE1alpha endonuclease inhibitor |
| 1187 | 1185 | STF-118804 | -0.63479 | -7.84743 | -10.81303 | -8.80685 | -11.43873 | -9.90718 | -8.23174 | -6.25941 | -4.05558 | NAMPT inhibitor |
| 1188 | 1186 | STF-31 | -0.45921 | 0.83499 | -3.39728 | -4.68841 | -11.91346 | -9.54251 | -9.42724 | -10.12505 | -3.09013 | GLUT1 inhibitor |
| 1189 | 1187 | SW033291 | -0.02315 | 0.34718 | -1.82446 | 0.02159 | -1.41447 | -1.46341 | -8.54648 | -11.88199 | -1.57209 | 15-PGDH inhibitor |
| 1190 | 1188 | Thiabendazole | -1.22940 | -1.97944 | -2.40267 | -2.61910 | -3.66826 | -2.24040 | -1.52100 | -2.44354 | -1.14830 | Anti-helminthic agent |
| 1191 | 1189 | Thiamet G | 1.28860 | 4.07742 | 1.70592 | 4.39375 | 2.50767 | 4.32944 | 5.43123 | 2.29316 | 1.65087 | O-GlcNAcase inhibitor |
| 1192 | 1190 | Tioxolone | -0.71169 | -0.56429 | -0.68057 | 3.10447 | 1.55504 | 2.00638 | -2.09666 | -10.85225 | -0.52263 | Carbonic anhydrase I inhibitor |
| 1193 | 1191 | U-104 | 0.34551 | -0.50760 | 1.24439 | -0.83513 | -3.12772 | 0.02731 | 1.85792 | -7.39335 | -0.53208 | Carbonic anhydrase IX and XII inhibitor |
| 1194 | 1192 | Cerulenin | 0.22900 | 0.58416 | -8.81230 | -2.58827 | 1.96829 | -10.25176 | -5.23284 | -7.94252 | -2.03265 | Inhibits fatty acid and steroid synthesis |
| 1195 | 1193 | Indomethacin | 0.37144 | -0.53161 | -0.61526 | -1.65354 | -2.46895 | -4.57927 | -1.48851 | 2.33816 | -0.54723 | NSAID |
| 1196 | 1194 | Miconazole Nitrate | 0.87921 | 3.35330 | 1.14694 | 1.32613 | -3.33304 | -5.50561 | -10.80257 | -4.19060 | -1.08630 | Antifungal agent |
| 1197 | 1195 | Salinomycin | -0.65309 | -1.10071 | -0.56564 | 4.96121 | 8.06393 | 4.18859 | -2.64466 | -5.11875 | 0.45230 | Potassium ionophore |
| 1198 | 1196 | Thapsigargin | -2.36754 | -12.51047 | -12.14335 | -5.23736 | -4.83458 | -6.80256 | -4.68209 | -6.64363 | -3.50263 | Sarco/endoplasmic reticulum Ca2+ ATPase inhibitor |
| 1199 | 1197 | TPEN | -2.08404 | -0.07019 | 2.01748 | 1.11044 | -11.79122 | -7.07246 | -7.78573 | -2.09334 | -1.76136 | Apoptosis inducer and heavy metal chelator |
| 1200 | 1198 | WZB117 | -0.29900 | -2.28742 | 0.78245 | -2.62497 | -1.94957 | -2.58541 | -3.26332 | -3.07810 | -0.97080 | GLUT1 inhibitor |
| 1201 | 1199 | Mycophenolic acid | -0.61164 | -1.59794 | 3.58301 | 2.33409 | -7.20947 | -5.15635 | -9.23561 | -6.88706 | -1.57182 | Inosine monophosphate dehydrogenase inhibitor |
| 1202 | 1200 | Pyrvinium Pamoate | -12.39744 | -14.09193 | -13.60002 | -6.22166 | -4.22498 | -9.90718 | -7.31597 | 80.66731 | 0.81875 | Anti-helminthic agent |
| 1203 | 1201 | Myriocin | -0.77251 | 0.56801 | 4.18429 | 4.92479 | 3.10557 | 5.60510 | 6.88942 | 6.06640 | 1.93908 | Serine palmitoyltransferase |
| 1204 | 1202 | Azithromycin | 0.39227 | 0.82497 | 1.76738 | 0.47501 | 1.38112 | -1.81188 | -1.80785 | -2.81780 | -0.10128 | Mitochondrial translation inhibition |
| 1205 | 1203 | Doxycyline | 0.59965 | 0.64553 | -3.12469 | -0.58386 | -11.51277 | -11.53445 | -7.04382 | -11.79451 | -2.81300 | Mitochondrial translation inhibition |
| 1206 | 1204 | Auranofin | -0.49392 | 0.36407 | -2.56209 | 1.24008 | -4.20593 | -7.66515 | -8.30306 | -8.20674 | -1.89225 | Antirheumatid |
| 1207 | 1205 | TRO-19622, Olesoxime | -2.09608 | -0.37684 | -0.47194 | 0.44389 | -0.41657 | -1.18171 | -12.36552 | -1.81277 | -1.15932 | Prevent mitochondrial pore opening |
| 1208 | 1206 | LRRK2-In-1 | 1.77888 | -3.60804 | -3.31705 | -4.08340 | -4.26749 | -8.98930 | -8.90979 | -4.85875 | -2.29960 | LRR2 inhibitor, LRRK2 binds with parkin |
| 1209 | 1207 | Chloramphenicol | 0.16597 | 0.24896 | 3.63460 | 0.41981 | -0.81272 | 0.31357 | 1.88132 | -1.56515 | 0.27188 | Mitochondrial translation inhibition |
| 1210 | 1208 | 2'-3'-dideoxycytidine | -0.73518 | -1.30906 | 0.19704 | 0.17946 | 0.04113 | -4.54288 | -6.03422 | -5.50632 | -1.12332 | Reverse transcriptase inhibitor |
| 1211 | 1209 | Mitoblock6 | -0.98835 | 0.99393 | 1.29235 | -0.86123 | -5.65815 | -7.88209 | -6.63558 | -8.76048 | -1.80769 | Mitochondrial protein import blocker |
| 1212 | 1210 | Avocatin b | -0.45070 | -0.34088 | -11.20619 | -1.30392 | -8.84799 | -6.23641 | -3.92612 | 0.06020 | -2.04570 | Fatty acid oxiation inhibitor |
| 1213 | 1211 | A-1155463 | -1.10616 | -1.93854 | 0.14735 | -2.68087 | -5.81728 | -7.83361 | -10.07127 | -11.71092 | -2.60130 | Bcl-XL inhibitor |
| 1214 | 1212 | BEC HCL | -0.49996 | 0.24715 | 1.44720 | -1.98467 | -0.88970 | -2.54281 | -1.34811 | -0.15479 | -0.36317 | Arginase inhibitor |
| 1215 | 1213 | DASA-58 | 0.64419 | 1.54871 | 5.51417 | -4.02264 | -12.22908 | -10.65455 | -7.63702 | -8.67505 | -2.25243 | Pyruvate kinase M2 activator |
| 1216 | 1214 | LTX-315 | 1.39803 | -0.84430 | -0.73347 | -1.49627 | 1.85391 | -2.52434 | 2.28785 | -3.50310 | -0.22591 | Bax/Bak-regulated mitochondrial membrane permeabilization |
| 1217 | 1215 | Telotristat Etiprate | -0.75203 | 0.21231 | 2.11254 | -0.43320 | -0.92560 | -0.60256 | 0.07562 | 2.24903 | 0.12280 | Tryptophan hydroxylase inhibitor |
| 1218 | 1216 | Vidofludimus | 0.38534 | -1.82471 | -1.54547 | -2.02650 | -1.12495 | -11.76966 | -7.10539 | -9.29007 | -2.17569 | DHODH inhibitor |
| 1219 | 1217 | Concanamycin | -7.79056 | -7.04824 | -1.64455 | -6.39053 | -6.97584 | -4.79239 | -0.99992 | 0.69430 | -2.21669 | Vacuolar type H+-ATPase inhibitor |
| 1220 | 1218 | Actinonin | -0.58539 | 0.75459 | 80.53561 | -2.06593 | -2.07825 | 6.98719 | -6.02395 | -10.34829 | 4.26086 | Mitochondrial translation inhibition |
| 1221 | 1219 | Tedizolid | -1.34493 | 1.51884 | 2.27375 | 2.51674 | -2.41782 | -0.64185 | -0.23641 | -1.80057 | -0.00839 | Antibiotics |

| drug.ID | Drug name | viable t_score at different concentration (nM) | | | | | | | | | | |
|---|---|---|---|---|---|---|---|---|---|---|---|---|
| | | 1.25 | 3.125 | 5.625 | 10.625 | 20.625 | 40 | 62.5 | 156.25 | 281.25 | 531.25 | 1031.25 |
| 13 | PD-169316 | 0.180439499 | -0.01932952 | -0.462084832 | 0.473182861 | -1.778534012 | -0.864534349 | -0.479134998 | -1.784621028 | -2.720295742 | -1.430192008 | -2.1611018 |
| 14 | PD-180970 | -3.982433908 | 0.036193932 | -0.51488785 | -0.80147897 | -1.506212784 | -2.305551836 | -2.928177174 | -2.568410457 | -3.136048453 | -2.489633146 | -4.5062336 |
| 17 | PIK-75 hydrochloride | -3.305664082 | 0.711044092 | 0.465422197 | 0.182931111 | 0.148200805 | 0.201214086 | 0.809380482 | -0.852833971 | -1.614330388 | 4.905048252 | -0.7918791 |
| 20 | JNJ-10198409, RWJ-540973 | 0.111588787 | -0.146898608 | -0.229537068 | 1.211895252 | -0.111166474 | 0.430939057 | -4.306091708 | -2.88099618 | -5.131013823 | -5.633792133 | -6.6939483 |
| 33 | K-252a, K-2151 | -0.225607846 | 1.064264083 | 0.83279897 | 0.592352951 | 3.284614007 | 1.125142649 | 4.609387355 | -0.317076715 | -2.864099547 | -2.274849154 | -2.7329488 |
| 34 | Lestaurtinib, CEP-701, KT-5555, SPM-924 | 0.289621487 | 1.366592868 | 1.668262661 | 1.964477467 | 5.488847432 | 4.154219206 | 2.686570752 | 4.736295673 | 2.247466748 | -3.050805698 | -2.3958368 |
| 38 | PKC-412, CGP41251, Midostaurin | -1.174452365 | -1.042075872 | 1.180765197 | 0.849174254 | -0.394098724 | -1.280681226 | 0.495271787 | 1.756087237 | 1.497763965 | -4.772658399 | -7.592029 |
| 42 | Staurosporine, AM-2282 | -0.994046437 | 0.900892194 | -0.58465734 | -1.324937639 | -3.431942698 | -6.535240756 | -7.032204421 | -6.631413674 | -4.252902761 | -2.618252105 | 0.13469143 |
| 44 | 6-bromoindirubin-3'-oxime, BIO, 6-BIO, MLS-2407 | 1.60319688 | -0.672633571 | 0.074405579 | -1.405086058 | -2.541991159 | -1.996575936 | -2.825493736 | -4.466788681 | -5.584708672 | -7.445177808 | -6.6854248 |
| 50 | GW441756 hydrochloride, GW-441756 | -0.120006599 | -2.31605377 | -3.641966161 | -3.35925208 | -3.808151057 | -5.201458892 | -2.336554686 | -4.250508518 | -3.348051466 | -3.612269899 | -2.3214808 |
| 61 | SU 6656, SU-6656 | 0.111953195 | -1.07163138 | 0.738236699 | -0.919777706 | -1.166512026 | -1.821300573 | -3.246153829 | -3.210605603 | -3.514858537 | -2.96045683 | -0.7944897 |
| 65 | Compound 52, NG-52, NG 52 | 0.96158541 | -2.399414136 | -0.94902195 | -4.365509111 | -2.665963407 | -1.829028479 | -2.612922242 | -2.609599541 | -2.399288109 | -3.137108726 | -3.1584134 |
| 68 | CGP-57380, CGP 57380 | -1.351387124 | -1.183137049 | -0.032418956 | -0.989860587 | -0.629099927 | -1.22681154 | -2.563163801 | -1.947683964 | -1.094197039 | -2.222640279 | -1.2765187 |
| 70 | GTP 14564, GTP-14564 | -3.049092696 | -0.177404292 | -0.312296536 | -1.078131279 | -0.340240674 | -2.049779907 | -2.367567398 | -3.186954191 | -5.282422903 | -3.980748084 | -3.7723119 |
| 83 | 2-Dimethylamino-4,5,6,7-tetrabromo-1H-benzimidazole, DMAT, Casein Kinase II, K25 | -0.639997906 | 1.0348628 | -0.104397198 | 1.290866364 | -0.754363837 | -1.813584386 | -0.893430336 | -2.771451311 | -2.515995083 | -3.741416241 | -4.0526368 |
| 93 | Kenpaullone, NSC-664704 | 1.631466554 | 0.187251137 | 0.523002912 | -1.60884085 | 0.373507832 | -0.567726929 | 0.402738489 | -3.458275533 | -1.059252839 | -2.776487063 | -4.1438599 |
| 117 | ER 27319 maleate, ER-27319 | -1.889161156 | -0.719820561 | 0.395022657 | -0.206326941 | -2.936217781 | -0.094317269 | -3.785258901 | -2.512894108 | -4.375602519 | -3.553449512 | -8.9382124 |
| 125 | SU 5416, SU-5416, TSU-16, NSC-696819, Semaxanib, Semaxnib, Semoxind | -1.344443516 | -2.032013523 | -4.374031231 | -2.265566364 | -6.574638285 | -5.642142615 | -6.643657037 | -6.558779449 | -6.391380801 | -6.481672501 | -6.4192788 |
| 132 | Necrostatin-1, MTH-DL-Tryptophan, MTH-trp, Nec-1 | -1.425331661 | -0.108236871 | -0.043141419 | -0.222806182 | -0.359207367 | -0.207932263 | -0.419953967 | -1.445796651 | -1.684629318 | -1.886490974 | -2.1403865 |
| 134 | PHA-767491 hydrochloride | 0.672705361 | -0.228200333 | -0.428078561 | -0.627684588 | -0.034451617 | -0.842779894 | -1.047077446 | -0.111837075 | -1.591118344 | -0.900325661 | 3.15537012 |
| 141 | A-443654, A-654 | 0.980937238 | 0.738652875 | 0.801140239 | 1.86553008 | 1.324735859 | 0.162314442 | -0.299618936 | 3.621977556 | 1.678760065 | -3.200996956 | -5.8890252 |
| 142 | CGP 57380, CGP-57380 | -0.589314931 | -1.838320332 | -2.951537725 | -1.988113404 | -2.581956642 | -1.417062471 | -0.533768791 | -1.476396785 | -3.050478591 | -3.970098132 | -0.9128455 |
| 149 | BI 2536, BI-2536 | -0.693489884 | -0.896425683 | 0.604826767 | 0.371080276 | 5.182137585 | -7.196014294 | -5.338565279 | -6.257364509 | -4.869461685 | -5.579931955 | -3.873014 |
| 203 | SB-218078 | 0.128503845 | 3.245983181 | 0.448943949 | 1.979284929 | -1.307617471 | 9.034149368 | 1.451109187 | 3.667212082 | 1.99916183 | -2.097885627 | -1.2863164 |
| 213 | Arcyriaflavin A | 1.433245884 | -0.670403369 | -0.616593501 | -0.355075644 | -0.852060737 | -0.292839573 | 0.123130788 | 1.396525212 | 1.095821266 | 1.076384577 | -6.5119787 |
| 238 | SU 16f | -0.106806018 | -0.665667113 | -0.693695096 | -0.821770559 | -1.356447717 | -3.355330098 | -3.783530688 | -3.675402203 | -1.989980288 | -5.244506464 | -5.1174316 |
| 239 | PIM 1 Inhibitor 2 | 0.913643309 | 0.583652764 | 3.028224949 | -1.875466926 | -0.566570045 | -0.515980224 | -2.66142228 | -4.166982733 | -3.270571888 | -3.980247642 | -3.6731189 |
| 255 | AP-24534, Ponatinib, Iclusig | 0.728970839 | -1.172803938 | -0.021096238 | 0.486939038 | 0.517283999 | -2.869875398 | -1.032188126 | -1.078361111 | -2.147095351 | -3.140749274 | 2.46293522 |
| 260 | PD-0332991, PD-332991, PD-991, Palbociclib, Ibrance | 0.028203934 | -0.288831855 | -0.454599985 | -1.251265271 | -1.522318594 | -4.386183767 | -4.469021238 | -6.19496312 | -5.785105534 | -6.884864144 | -6.3314803 |
| 268 | Alvocidib, HMR-1275, L-868275, MDL-107826A, NSC-649890, L-868276, Flavopiridol | 0.173362494 | 0.483766344 | -1.282435192 | 0.592661856 | -1.304981753 | -0.253887443 | -1.876450351 | 0.526925327 | 2.501128935 | 1.382452494 | 3.34031577 |
| 270 | GSK-461364, GSK461364 | -2.081825375 | 0.061073457 | 0.468886262 | 0.049276643 | 0.339336425 | -4.280241191 | -5.359828795 | -4.368723649 | -4.653594 | -4.621757393 | -4.2623645 |
| 271 | BI-6727, volasertib | 0.165064751 | -0.581613865 | 1.566319437 | -0.164929968 | -0.792452257 | -2.153884305 | -6.427745182 | -4.594166222 | -3.26031572 | -5.408292928 | -4.3242572 |
| 278 | AT9283, AT-9283 | 0.197079602 | 0.048001129 | 0.982661319 | 3.871390698 | 3.600399738 | -3.428503296 | -3.427263911 | -3.048818028 | -1.90737399 | -0.092191485 | 1.3097792 |
| 284 | Bisindoylmaleimide X, HCl salt | 0.212249249 | 1.450095763 | -1.672848517 | 14.58577897 | -0.760594479 | -0.230836012 | -0.411969341 | -0.036882798 | 2.693318002 | -7.411765524 | -1.2043784 |
| 291 | CYC-116 | 0.102051209 | -0.174113812 | 0.343838138 | -2.234338031 | -3.61621189 | -3.207726272 | -2.541441975 | -3.25115506 | -2.412826849 | 1.157648586 | 0.24472593 |
| 299 | PD173955, PD-173955 | 0.141273678 | -0.040524515 | -2.103397986 | -2.193838392 | -2.539876042 | -1.092673022 | -2.142561569 | -3.073906715 | -2.630353544 | -2.244521043 | -3.0662579 |
| 312 | SNS-314 | 0.068291488 | -0.410629937 | -1.001771335 | 2.017818119 | 0.716020904 | -4.854168632 | -4.645578846 | -4.122079838 | -3.268710156 | -5.576228943 | -5.4208001 |
| 327 | (+)-P276, P276-00, Riviciclib hydrochloride | -1.458705029 | -0.431269012 | -0.3903611 | 0.939361326 | -1.040934622 | 0.392201223 | 0.269314713 | -1.367955261 | -0.475134395 | 0.219883281 | 0.73149254 |
| 329 | Dianilinopyrimidine_01 | -0.096362054 | -0.215132874 | -2.082750491 | 1.814137163 | 0.128082918 | -0.247320622 | -0.000179089 | 0.754003959 | -0.010810485 | -2.993271429 | -2.0817744 |
| 339 | AMG-900 | 0.448562448 | -0.550832129 | -0.016660667 | -0.816978507 | 5.056022457 | -2.690263182 | -5.966620665 | 5.06396829 | -1.956495852 | -4.540287793 | -2.7369566 |
| 347 | ASC-069, APY 69 | -0.737900881 | -2.102353005 | -0.329014072 | 4.032320481 | 1.39399377 | -2.440745946 | -0.992877167 | 1.102604167 | 3.066761306 | 1.873695473 | 0.82201992 |
| 348 | ASC-081, APY 81 | 0.236233993 | -0.266265983 | -1.188281827 | -3.693979425 | -0.121032861 | -1.62803704 | -1.165276067 | -3.527905632 | -3.936579067 | -3.031703746 | -2.4330754 |
| 349 | ASC-082, APY 82 | -2.552980673 | -1.547573527 | -1.437646431 | -1.072256531 | 2.495084785 | -0.3643645 | -0.304302965 | -4.580940172 | -6.190101213 | -7.654820002 | 3.52326945 |
| 350 | ASC-086, APY 86 | -0.959642466 | -3.060032523 | -0.927477825 | -1.472862105 | -1.864201058 | -0.929157287 | -2.340358638 | -0.47131965 | 0.186032295 | -1.245347203 | -0.0266779 |
| 362 | Cdk1/2 Inhibitor III | 0.469581942 | -0.01336468 | 0.675882392 | -0.598659892 | -1.13400529 | -2.255991024 | -2.077439236 | 0.577111089 | -6.60403858 | -1.78995908 | 0.03015121 |
| 378 | IRAK4 Inhibitor | -0.964042847 | 1.350408428 | -1.135160273 | -2.363775979 | -3.959163525 | -3.81613042 | -2.197758272 | -1.634205195 | -2.743876158 | -1.748061056 | -1.1504598 |
| 382 | AZD5438, AZD-5438 | 0.347551583 | 2.092792351 | 1.211525694 | -0.136480119 | 0.430932105 | -0.752200244 | -2.267867543 | -3.476660522 | -7.473385294 | -2.74422482 | -0.4808561 |
| 387 | RAF265, RAF-265, CHIR-265 | -0.907805492 | -0.948924108 | -1.075642481 | 0.523989107 | -0.944640253 | -1.194538605 | 1.679995351 | -1.559152907 | -3.66018623 | -2.79218387 | -6.5658055 |
| 424 | Hesperadin | -1.111484679 | -0.096422072 | 2.344603742 | 5.267050376 | -4.370847998 | -3.80702408 | -3.979062989 | -0.019898656 | -2.419782648 | 1.262871875 | 2.35246233 |
| 433 | Indirubin, NSC-105327, Couroupitine B, Indigo red, Indigopurpurin | -1.64373713 | -2.865590113 | -2.111848054 | -2.102464595 | -3.603492457 | -7.211222756 | -5.35739503 | -3.60960314 | -4.64554849 | -4.380839937 | -4.2764438 |
| 449 | NQDI-1 | -0.864659544 | 1.180911953 | 0.536739018 | -1.837153125 | -1.442677122 | -4.139657394 | -3.214925503 | -5.177528913 | -3.168878829 | -4.205991614 | -5.599204 |
| 468 | A 1070722, A-1070722 | 1.346732175 | 0.254234062 | -0.441991035 | -1.392226038 | 1.621099772 | -3.047829374 | -2.806609003 | -0.145081802 | -2.841548587 | -1.103205978 | -2.2746989 |
| 473 | PHA-848125, PHA-848125AC | -0.751963053 | -0.028705228 | 1.10087755 | 0.836374778 | 1.800682872 | -0.733012301 | -0.125633673 | -1.236172521 | -0.295306963 | -2.016936411 | -5.2179437 |
| 480 | Dinaciclib, MK-7965, NSC-727135, SCH-727965 | -0.782113002 | -0.112445368 | 1.657124762 | 12.2161305 | 4.985634 | 0.608033309 | 2.842450283 | 4.868028521 | 1.93844539 | 5.794746776 | 2.94538513 |
| 487 | Trk Inhibitor III, GNF-5837 | -2.579556464 | 0.455967648 | 0.597442819 | 1.268906163 | 0.574864152 | -0.434538252 | -2.400207572 | -2.456838002 | -6.059736265 | -10.70184652 | -11.062474 |
| 505 | LY-2606368 | -0.15326057 | 1.635768236 | 5.961469913 | 12.08408341 | -1.419460228 | -2.908518905 | -1.662633113 | -4.718201132 | -2.973138583 | -4.194651755 | -4.114265 |
| 522 | NMS-1286937, NMS-P937 | 0.972264585 | 0.520056101 | 0.647331099 | 0.793307612 | -0.577817584 | 2.212709704 | -2.907418979 | -4.845122536 | -6.39479147 | -5.692375426 | -6.3189736 |
| 532 | LY2835219, LY-2835219 | 0.224992394 | 0.583719187 | 1.037188865 | -1.375116166 | -1.394516101 | 0.562233024 | -1.123102254 | -2.773596557 | -3.883740684 | -3.360063394 | -2.0734074 |
| 534 | AMG 925 | 0.021247108 | 1.785651031 | -1.562293629 | -0.80558909 | -0.995446086 | -2.368961382 | -2.19741734 | -4.386549545 | -4.623572016 | -5.539380974 | -5.9625482 |
| 549 | AVL-292, CC-292, Spebrutinib | -0.108505271 | -0.359860587 | 1.116519686 | 0.559628177 | -0.918930833 | -0.714730196 | -1.639196135 | -1.765950322 | -2.165705622 | -0.452744858 | -0.2127887 |
| 562 | Bortezomib, Velcade, MG-341, PS-341, DPBA, JNJ-26866138, LDP-341, MLN-341, NSC-681239 | 0.334774336 | 1.305930306 | 0.283395316 | 4.314495594 | 6.765105595 | 0.802047329 | 3.998725493 | -0.042285599 | 2.441282611 | -1.058937571 | 3.63654084 |
| 568 | Ouabain, g-strophanthin, BNC-1 | 1.211151124 | 1.101592083 | 0.534075736 | 0.54870094 | -0.41441206 | -1.845906637 | -4.184974413 | 3.586761396 | 4.579099287 | 6.626564578 | -2.6027528 |
| 589 | Bexarotene, Targretin, Targretyn, Targrexin, LGD-1069, LG 1069, LG100069, LG-100069, SR-11247 | -1.102499947 | 0.825435939 | 0.09447901 | -3.07925591 | -1.740946446 | -5.320858816 | -6.128489837 | -9.863241687 | -10.49209235 | -12.28214078 | -13.73843 |
| 590 | Taxotere, Docetaxol, ANX-514; EmDOC; ML-061; NSC-628503; RP-56976; SD-009; SDP-014; SID-530; SYP-0704A; | 0.79528023 | 2.247053036 | 7.854947425 | 2.332899079 | 6.560278154 | -4.167432234 | -5.371733189 | -9.235609628 | -8.51804262 | -4.950952507 | -4.5385903 |
| 591 | Paclitaxel, Taxol | 0.461301717 | 2.070534279 | 5.519092963 | 15.32572837 | 17.15247226 | 7.766984217 | 4.816855892 | -1.596683708 | -9.76592093 | -9.311666556 | -4.9412343 |
| 593 | ABT-263, Navitoclax, ABT263, Navitodax, RG-7433 | 0.971411661 | 0.096593773 | 1.259351218 | 1.439993916 | 2.388781095 | 2.863615215 | 1.97527514 | 2.958677906 | -2.739491059 | -3.162080915 | -7.8161929 |
| 598 | D07001-F4; Inno-D07001; LY-188011; dFdC, Gemcitabine hydrochloride, Gemzar | -0.263549286 | -4.38179635 | -1.78321659 | -7.374231756 | -6.295542912 | -4.656803409 | -3.499928432 | -7.68859539 | -0.371041505 | -5.119273435 | -6.8061023 |
| 599 | , Alvespimycin, KOS-1022, NSC 07545, 17-(Dimethylaminoethylamino)-17-demethoxygeldanamycin Hydrochloride,BMS | 1.045216177 | -0.224419248 | 0.961135318 | 1.717647484 | 5.113914144 | 10.36032544 | 16.11748855 | 11.42232481 | 7.957888895 | 8.74793159 | 0.02962575 |
| 600 | Ispinesib, SB-715992, CK0238273, CK-0238273, 715992 | 0.724901271 | 0.28859086 | 10.27783736 | -2.682219925 | -3.948683696 | -6.340005634 | -7.350219192 | -7.128501529 | -3.002685536 | -6.567328139 | -7.1687681 |
| 605 | XAV-939, NVP-XAV-939 | -0.223296808 | -0.205012447 | -1.976672834 | -0.172255123 | -2.844837706 | -0.924177562 | -1.657680953 | -1.260558738 | -1.482794649 | -1.518839574 | -1.4557849 |
| 609 | GX15-070 mesylate, GX-015-070 mesylate, Obatoclax Mesylate, CEP-41601 | 0.321767116 | -0.264117299 | -0.491340519 | 1.003552635 | 0.824458011 | 4.854179651 | 3.324239055 | 2.669075287 | -8.33973977 | -7.151619718 | -10.406979 |
| 651 | Pemetrexed disodium heptahydrate, Alimta, LY-231514, MTA, Rolazar, Tifolar | 0.712272049 | 0.171456442 | 1.070710161 | 13.75278568 | -1.800494907 | -1.418867127 | -3.970529186 | -3.258697093 | -1.621783212 | -1.339260142 | -2.7466714 |
| 655 | NVP-AUY922, AUY922, VER-52296 | -0.073518802 | 1.919173261 | 9.925205539 | 2.041008448 | 11.19396846 | 7.968886403 | 8.316072969 | 12.30137072 | 7.757936028 | 1.050245094 | 4.18770266 |
| 657 | LBH-589, Panobinostat, NVP-LBH589, LBH589 | 2.421508738 | 5.844119605 | 6.303185249 | 7.988784127 | 8.520727926 | 1.166745791 | -0.718211113 | -8.01645374 | 4.819191901 | -0.473552589 | 7.32199198 |
| 661 | etinoin, Vitamin A acid, all-trans-Retinoic acid, NSC-122758, Airol, Atralin, Avita, Nexret, Renova, Retacnyl, Retin-A Mic | -1.545497733 | -4.931813482 | -7.619331137 | -10.99581908 | -12.36404773 | -12.47172815 | -13.79970156 | -13.16825473 | -12.21415526 | -13.41506724 | -13.906891 |
| 666 | Daunorubicin hydrochloride, Rubidomycin hydrochloride | 1.104681396 | 1.767375893 | 5.430938299 | 3.3138625 | -1.627657017 | -8.432465749 | -9.105847724 | -7.392753572 | -5.088803361 | 4.709209743 | 1.774623 |
| 667 | Idarubicin hydrochloride, Idamycin, Zavedos, 4-demethoxydaunorubicin | 2.048024338 | 4.750512679 | 1.042777858 | -5.564969215 | -9.833735906 | -8.167424679 | -10.14619781 | -7.332651042 | -3.221954944 | 4.053562336 | 4.97870036 |
| 668 | Epirubicin hydrochloride, Ellence, Pharmorubicin, Epirubicin Ebewe | 1.054799389 | 1.87703129 | 1.291930602 | 4.520165304 | 4.408850174 | 2.186566197 | 2.888723223 | -6.846605338 | -7.900254447 | -9.885277945 | -9.2803852 |
| 672 | Carfilzomib, PR-171, ONO-7057 | 0.209572747 | 3.681577827 | 17.10348572 | 19.45191301 | 3.977169455 | 0.847056392 | 4.853632964 | -2.33134821 | -3.52622917 | 4.008158182 | -3.7460963 |
| 674 | MGCD0103, MGCD-0103, MG-0103, Mocetinostat | 0.664161036 | -1.547526867 | -0.745845304 | -1.502015233 | 1.319877929 | -0.0600797 | -0.949495552 | 5.108284388 | 4.563113043 | 2.404622027 | 7.16746914 |
| 675 | LAQ824, LAQ-824, NVP-LAQ-824, Dacinostat | 1.266856576 | 0.8263016 | 7.337523509 | 7.943512559 | 9.388099885 | -0.947632399 | 4.351035701 | 1.466805823 | 8.343450359 | 5.144392497 | 4.77669319 |
| 685 | JNJ-26481585, Quisinostat | 3.029669495 | 7.776591185 | 9.435690907 | 10.2856811 | -1.906671596 | -3.205367509 | 1.707743842 | 3.390311226 | 1.208191682 | 2.976757612 | 0.98249799 |
| 690 | Pifithrin-alpha | -1.119786306 | 0.271278729 | -1.87787228 | -1.422384066 | -0.93897977 | -1.660003212 | -3.099540791 | -2.912446947 | -1.971952598 | -2.920908541 | -5.9308791 |
| 695 | Cyclic Pifithrin-alpha hydrobromide | 0.240712291 | 0.760537076 | -1.577831389 | -2.402996093 | -0.639904249 | -3.423964924 | -2.123362431 | -2.598827944 | -1.810384452 | -3.112307432 | -3.2294868 |
| 701 | Dexamethasone, ISV-305, OTO-104, Decadron, DexaSite, Maxidex, ProDex | -0.923687199 | -2.024069516 | -3.338270445 | -4.260879521 | -7.899314576 | -8.37487851 | -9.793772701 | -10.071355 | -10.13703273 | -10.53647055 | -10.227328 |
| 703 | SHH Antagonist, JK 184 | -0.357674863 | 0.08895276 | -0.010996866 | -0.050110908 | -0.071738858 | -0.377714173 | 1.726922628 | -3.382853635 | -3.63512718 | -5.889628583 | -5.260845 |
| 714 | SC-1, Pluripotin | 0.15021041 | -0.428383953 | 0.534605058 | -0.259579671 | 0.178401679 | 0.30820019 | 0.128904995 | 1.146008555 | -0.018824537 | 0.824985104 | 0.8002022 |
| 716 | Trapoxin A, RF-1023-A | 3.772627716 | -0.237023579 | -0.709856779 | 3.17496366 | -4.237270935 | 6.082861956 | 1.729828305 | 3.879361387 | 2.35054329 | -0.48949196 | 4.00174998 |
| 717 | Chaetocin, Chetocin | 1.100312793 | -1.231181859 | 0.27454929 | -0.064761478 | -0.228133229 | -2.13006541 | -3.76064745 | 0.738378431 | -3.696613442 | -1.665150005 | 4.822838 |
| 718 | RITA, NSC-652287, NSC-652287D | -0.756062444 | -1.630300532 | 0.254042912 | -1.627750341 | 2.181052949 | -1.727229887 | -2.740919298 | -0.646478064 | -1.311449753 | -0.557881604 | 1.47321099 |
| 720 | OICR0000860A01, SRT2183, SRT-2183 | -1.374526927 | -0.306876841 | -0.448941126 | 0.276269797 | 0.037649095 | 0.657851372 | -1.48312102 | -1.76891619 | -2.402488486 | -3.050682444 | -4.0084669 |
| 727 | bine, Ara-C, Cytosine b-D-arabinofuranoside, AraC, CHX-3311, NSC-63878, U-19920A, Aracytin, Cylocide, Cytosar, Cyt | -2.256770136 | -4.070373363 | -4.467134465 | -7.851779903 | -6.981395385 | -7.303395314 | -4.772100607 | -4.435574213 | -4.760551206 | -7.303839351 | -6.4107319 |
| 729 | crine Hydrochloride, CI-880, NSC-156303, NSC-249992, SN-11841, m-AMSA, NSC-141549 (HCl), Amecrin, Asidine, Ar | -0.109656451 | 1.066173285 | 0.459059111 | 1.378499246 | 3.142095093 | 4.695942827 | 2.065921858 | -0.178525396 | -1.453153945 | -6.828996458 | -8.8373162 |
| 734 | Cladribine; Movectro; NSC-105014; Leustatin; Litak; Leustat; Mylinax; RWJ-26251-000; RWJ-26251; 2-CdA | -0.178987133 | -0.770366275 | -1.193936932 | -2.666499491 | -1.656485878 | -5.589178104 | -6.080625919 | -5.166214148 | -7.339413992 | -3.944262572 | -2.3110929 |
| 735 | Clofarabine; SAR-393590; Clofarex; CAFdA; Evoltra; Clolar; 2-Cl-2'-F-araA | -2.04279018 | -4.202709668 | -5.82660741 | -8.360683678 | -8.042107004 | -7.738535642 | -7.378744177 | -6.855147433 | -4.95131711 | -3.991283374 | -6.1033142 |
| 737 | Fludarabine, F-ara-A, NSC 118218 | 0.438519573 | -0.193267371 | 1.33632633 | -0.22419698 | -1.494394734 | -1.342203111 | -0.12976569 | -1.298273816 | -2.967583937 | -4.751923682 | -6.3529516 |
| 739 | Actinomycin D, Dactinomycin, Cosmegen, NSC-3053 | -1.585386868 | -5.833700841 | -5.483467079 | -3.959900635 | -0.71868526 | 3.09516214 | 4.30635008 | 7.496988355 | 2.282234178 | 3.56013821 | 1.31506413 |
| 745 | Xaluprine; DR-6MP; NCS-755; 6-Mercaptopurine; Purinethol; Mercaptopurine | 0.852764168 | -0.745428711 | -0.305125685 | -0.169875418 | -0.61156825 | -2.508332384 | -0.874026615 | 0.060207101 | -2.019104927 | -1.951385946 | 0.49797213 |
| 747 | Mitomycin C; Mitosol; Mitomycin; MMc; Ametycine; Mito-medac; Mutamycin | -0.553782613 | 0.793606348 | 1.746274721 | 0.972020728 | -0.585325367 | -0.382999683 | 0.359533841 | -2.423493239 | -1.599251607 | -6.484787287 | -2.4022267 |
| 754 | Raltitrexed; ICI-D-1694; NSC-639186; ZN-D1694; ZD-1694; D-1694; Tomudex | -0.991740723 | -0.805550653 | -1.282976372 | -2.268735997 | -5.482235702 | -5.355695483 | -2.030049971 | -4.472089332 | -4.000225523 | -5.317960991 | -3.1889389 |
| 756 | can Hydrochloride Hydrate; Nogitecan hydrochloride; NSC-609699; Hycamtin; NK-211; E-89/001; SK&F-S-104864-A; E | 2.965747411 | 3.413985777 | 4.221593816 | 3.294366389 | 4.143870483 | -3.183900653 | -5.815092669 | -5.483063999 | -2.806496707 | -3.248979121 | -7.0342493 |
| 757 | Thioguanine; Tabloid; NSC-752; Tioguanine; 6-Thioguanine; Lanvis | -1.132913986 | 0.903977718 | 0.35815293 | -1.99533902 | 0.214552792 | 1.568576733 | 0.964845068 | -0.560302163 | 0.58376097 | 2.766715011 | 0.73594502 |
| 771 | Apicidin, OSI-2040 | 0.061210867 | 1.136464656 | -0.673806016 | -0.414176492 | -0.721842216 | 0.356435956 | 1.426136209 | 4.817730974 | 4.243127656 | 2.564960861 | 4.66657688 |
| 782 | n; Cortef; Chronocort; Dermacort; Cortisol; Alphaderm; Aeroseb-HC; Acticort; Plenadren; Proctocort; Anusol-HC; Cetaco | 1.678583453 | -0.240850448 | 0.968995149 | -1.200475475 | -2.462110616 | -3.96714895 | -4.77933766 | -7.769845279 | -7.20134538 | -8.794915721 | -8.9021812 |
| 785 | 7-acetate; Gestoral; Farlutal; MAP; Depo-subQ provera 104; Amen; Medroxyprogesterone acetate; NSC-26386; Depo- | -1.162783598 | -0.069886765 | -3.985359986 | -2.861202527 | -2.035487985 | -3.316089545 | -4.592741982 | -5.960174124 | -6.059788118 | -6.853269621 | -5.3469639 |
| 787 | Prednisolone; Prelone | -0.146650714 | -0.725615236 | -1.091588624 | -3.175150591 | -3.538052287 | -5.974207393 | -6.33848217 | -7.938821016 | -9.178275647 | -9.649844962 | -8.9325472 |
| 796 | Megestrol acetate; DW-ES(B); PAR-100.2; Megace ES; Megestat; Megace | 0.987155606 | 0.283414668 | 0.005458573 | -0.976572164 | -1.774415727 | -0.514650902 | -1.318145308 | -3.14595554 | -4.83554756 | -3.848144909 | -3.5674546 |
| 797 | Teniposide; VM-26; Vumon; Podophyllotoxins | 0.121956615 | 1.300315617 | -6.14300786 | 0.464734005 | 2.643322482 | 2.917912958 | 5.472860005 | 0.017024449 | -0.215409958 | -5.437299316 | -4.8597331 |
| 809 | CX-5461 | 0.566959779 | -0.388686662 | 0.282553941 | 0.777857142 | 2.065179076 | 3.036475694 | 2.041512975 | -4.562506214 | -5.396005201 | -5.911828029 | -7.3634862 |
| 811 | MLN2238, MLN-2238, Ixazomib | -0.382927188 | -0.512710836 | -0.692940849 | -4.257874515 | -0.280190405 | -0.708369364 | -0.209188555 | 5.860728766 | 0.978171284 | -3.598046028 | 3.44404171 |
| 820 | Diphenyleneiodonium chloride, DPI | -0.340207218 | 2.259556274 | -0.73764956 | 0.464776569 | 0.040269463 | 0.570913406 | -0.916043352 | 0.413606308 | 0.234149296 | 1.385222692 | -3.3476528 |
| 821 | Dp44mT, HDp44mT | -0.468131998 | 0.662525544 | -0.730315562 | 0.092302741 | 0.286284964 | 1.669314789 | 0.139935789 | 0.970480498 | 0.692757991 | -2.202056143 | 8.00363177 |
| 837 | rochloride, NSC-123127, Doxorubicin hydrochloride˜(Rec INNM; BAN; USAN; USP), Adriacin; Adriamycin; Adriblastina; F | 1.163598878 | -1.601695232 | 1.381596605 | 0.288017821 | 3.105297456 | -0.174657011 | -2.155792485 | -8.017916637 | -7.050356141 | -6.497384862 | -4.6276879 |
| 860 | CUDC-907 | 7.812440641 | 2.231646322 | 0.358138664 | 0.760928813 | -6.174671012 | 1.280226472 | -1.344763838 | 1.227235675 | 3.49056715 | 2.982576486 | 4.5986861 |
| 879 | ARRY 520 trifluoroacetate, ARRY-520, Filanesib | 0.355040867 | 4.089576104 | 0.115681784 | -4.234787029 | -7.195446264 | -6.061111412 | -5.411929453 | -3.010568563 | -5.896534617 | -4.661114255 | -5.0542028 |
| 881 | Tazarotene, AGN-190168, Avage, Fabior, Suretin, Tazorac, Tazoral, Zorac | 1.245428685 | -0.25234283 | -1.466926891 | -0.654811279 | -0.726994741 | -4.711447157 | -5.388747461 | -8.35166833 | -10.00165332 | -12.15903811 | -11.799675 |
| 882 | AHPN, AGN-192837, CD-437, CD 437 | 0.377060739 | -1.093476893 | -2.198585776 | -0.783838945 | -1.930420096 | -1.08703773 | -0.368933454 | -6.228768403 | -7.486313697 | 0.076683407 | -9.7463485 |
| 883 | Am-580, AM 580, CD-336, Ro-40-6055 | -12.25830746 | -12.13724527 | -11.92855372 | -11.68257336 | -12.56065045 | -12.18454943 | -11.99490476 | -12.31248863 | -12.07575071 | -12.20328577 | -11.72322 |
| 886 | AC-261066 | -0.634920034 | 1.461608035 | 1.660701869 | -0.76542016 | -0.628319275 | -2.644415015 | -2.417980843 | -7.583887091 | -9.794248778 | -11.45522498 | -11.415007 |
| 888 | BMS 753 | 0.383919977 | -0.666930238 | -2.661809911 | -5.801834038 | -4.112848885 | -10.62793056 | -10.77498206 | -12.07384603 | -11.23072147 | -12.07372077 | -12.120292 |
| 892 | CD 2665 | -0.524211712 | 0.494749445 | 1.340029143 | 0.97743584 | -0.047532537 | -0.601593014 | -2.438325938 | -4.364794317 | -5.441338495 | -5.654146124 | -7.5296349 |
| 893 | BMS 195614, BMS-614 | 0.732931413 | 0.430474276 | -0.827266914 | -1.918893811 | -1.894201145 | -1.920243843 | -1.732735736 | -1.635304555 | -2.641382571 | -3.698976443 | -2.1105007 |
| 897 | ER 50891 | -0.881671394 | 1.111774273 | 0.558240281 | 0.485538435 | -1.12129137 | -8.687877678 | -0.024921373 | 0.104912256 | -0.44889054 | -0.002695071 | 0.49406506 |
| 898 | BMS 453, BMS-189453 | -1.804402089 | -0.313628413 | -3.96340211 | -5.929024672 | -8.691613025 | -11.43006987 | -11.10537398 | -11.7471597 | -11.82405409 | -11.81714451 | -12.365375 |
| 906 | Fluorobexarotene | 0.140776717 | 0.70135539 | -0.346371853 | 1.186603924 | -1.842019217 | -3.248735553 | -3.840694791 | -7.091327952 | -8.867490333 | -10.07414655 | -11.84217 |
| 911 | ARRY-520, Filanesib | 1.357076814 | 2.519090716 | -6.453624663 | -6.512709547 | -3.255475999 | -3.816334127 | -8.166370575 | -6.272696667 | -4.930502813 | -5.35366286 | -7.8599602 |
| 918 | OTX015, OTX-015, Y-803 | 0.774731878 | -0.769951821 | -0.644176931 | 0.081549839 | 0.975306427 | 1.700492027 | 0.551953869 | -1.588483137 | -1.162900374 | -0.867824767 | -1.1142789 |
| 926 | HSP990, NVP-HSP990, HSP-990 | 0.788745021 | 0.43788747 | 3.620431243 | 9.96246149 | 1.570535365 | -0.822946716 | 4.253152421 | 1.084000489 | 3.779272647 | 4.612530952 | 3.4294115 |
| 930 | GSK923295, GSK-295, GSK-923295, GSK-923295A | -1.501715144 | 0.655971068 | -1.174037982 | -1.175872832 | -0.137196447 | 1.773508216 | 17.02825237 | -3.981337357 | -4.310438668 | -4.653173647 | -3.1485131 |
| 945 | AMG 232, AMG-232 | -1.203820929 | 2.099074675 | 1.07699501 | -1.74464713 | -0.21562116 | 0.447113354 | 0.351141872 | 2.223500679 | 1.599859601 | -3.891279825 | -4.8765039 |
| 964 | Doxorubicin, NSC-123127, SA-033, Adriacin, Adriamycin, Adriblastina, Resmycin, Rubex | -2.113334378 | 0.979304644 | 0.66494389 | 1.027484294 | 2.391029845 | -1.109774201 | -2.61822045 | -5.756872065 | -7.844865028 | 1.465542822 | -10.057387 |
| 980 | GSK-J4 | 2.613066971 | -0.887854341 | 1.673939963 | -1.01459128 | 1.290845787 | -0.751737305 | 1.378832474 | 1.500372183 | -0.459341329 | -2.129787964 | -3.1644063 |

| | | | | | | | | | | | | |
|---|---|---|---|---|---|---|---|---|---|---|---|---|
| 982 | Bromosporine | 1.601324049 | -0.519384163 | 0.536142965 | 1.173399256 | 0.898572714 | 1.38038999 | 0.463572626 | 0.677311997 | 1.106244778 | -3.887523848 | -1.8543963 |
| 984 | CPI 203, CPI-203, CPI-'203 | 1.596581921 | 1.014130403 | 1.673233849 | 0.007335989 | -0.280288492 | -1.860377982 | 0.50538087 | 1.457495813 | 7.337209051 | 3.529495059 | 6.79239577 |
| 995 | Fisetin | 1.433994416 | 0.960077542 | -0.291827715 | 3.247771641 | -0.11013996 | -1.078417488 | -0.405916488 | 0.654594417 | -0.796443092 | 0.873636088 | -0.2573899 |
| 1010 | rringtonine, Omacetaxine mepesuccinate, CGX-635, HHT, NSC-141633, ZJ-C, Ceflatonin, Myelostat, Omapro, Synribo, | 0.681907495 | -2.284869001 | -4.469300822 | -4.725781838 | -2.613832283 | 3.02705078 | 3.656782563 | -2.47813122 | 0.467548301 | 2.449244349 | 0.05339676 |
| 1012 | Selinexor, KPT-330 | -0.011463414 | 1.102175316 | 0.451907159 | -11.79210093 | -3.014657538 | -1.433057039 | -4.825281644 | -6.971326672 | -4.539786486 | -5.048729545 | -2.2714685 |
| 1025 | Parbendazole | -1.963692626 | -1.557589462 | 1.739043412 | -0.617072186 | 0.66903516 | 0.434970474 | 0.616967632 | 3.018663879 | 3.915995526 | -5.336643438 | -7.1532988 |
| 1028 | I-BET726, GSK1324726A, GSK-1324726A, I-BET-726 | 1.930508858 | -0.045882578 | 0.494562612 | -0.277806462 | -0.282863455 | -0.39931424 | -1.384645496 | -1.869419934 | -0.396001252 | -5.153987252 | 0.72669369 |
| 1041 | A-196 | -0.32550075 | -0.455435707 | -0.352497141 | 6.68453104 | 0.068298514 | 2.846377865 | 1.224538332 | 1.305971851 | 0.019872933 | -0.774924517 | 0.46921953 |
| 1042 | A-366 | 1.356684604 | -1.026916768 | 0.098524818 | -0.503502876 | 1.995538278 | 0.2733227 | 1.745060835 | 0.011007845 | 3.261802187 | 0.385941754 | 0.79870426 |
| 1043 | A-395 | 0.597875468 | -0.569553598 | 0.911760715 | 0.680120715 | 0.789958863 | -0.488919533 | 2.622203258 | 1.509256371 | 1.456619002 | 0.295888293 | 3.63299734 |
| 1044 | BAY-598 | 1.499236875 | -0.1204882 | 0.591346617 | -1.976882464 | 0.134100746 | 0.009691424 | -1.639376635 | 0.229943914 | 1.343133909 | -1.544342974 | 1.6340478 |
| 1045 | BAZ2-ICR | -0.622103494 | -2.236480431 | 0.560998466 | -1.347895732 | 0.197100022 | -0.330249589 | -0.318653967 | 0.756101904 | -0.36592709 | 4.18617199 | 0.26403423 |
| 1046 | BI-9564 | -1.573523411 | -0.51353396 | 0.921257597 | 0.744558702 | 1.7956551 | 1.241853765 | 0.588655602 | 0.684073331 | -0.561554595 | 1.330328567 | 1.13783352 |
| 1047 | GSK2801 | 2.420581189 | -0.430707022 | 1.795101208 | -0.060333169 | -1.284172167 | -0.911732304 | -0.430337299 | 1.243512011 | 0.818278366 | 1.01727699 | 1.16711949 |
| 1048 | GSK343 | 1.48904272 | 1.291035867 | -0.101509727 | 1.005042063 | -0.258251564 | -0.727420781 | 0.895445015 | 1.330485471 | 0.298600328 | -0.46229843 | 2.10653159 |
| 1049 | GSK484 (hydrochloride) | 1.680236381 | 0.405103377 | -0.730291978 | 1.029235422 | -1.434344983 | -1.404773317 | -0.326368317 | 1.710693602 | 1.413380364 | -1.295210909 | 1.55100998 |
| 1050 | GSK591 | -0.002673131 | -0.778924541 | -0.524689599 | -0.433515735 | 0.954067441 | -0.212460902 | -0.16923882 | -0.005709688 | -0.989061617 | -0.302654781 | 1.04429226 |
| 1051 | GSK864 | 0.981318917 | -0.415511337 | 1.86595214 | 0.077294734 | -1.80411903 | -0.467816747 | 1.790497954 | 0.143029731 | 0.991287386 | 0.639217734 | 0.7429562 |
| 1052 | GSK-J4 (hydrochloride) | 0.076380989 | -0.747838409 | 0.918262755 | -0.259801971 | -0.035479823 | 1.09012403 | 0.699285806 | -1.15514405 | -0.435980762 | 1.594987997 | 1.86952399 |
| 1053 | GSK-LSD1 (hydrochloride) | 0.554940638 | 1.75661224 | 1.075347975 | -1.642555369 | -1.778907753 | -1.31092792 | -0.558883062 | -0.529099675 | -0.877972355 | 0.091130224 | -2.339057 |
| 1054 | I-BRD9 | 0.389129509 | 0.563834436 | -0.889690627 | -1.863348725 | 1.181176877 | -0.627218446 | 0.436339344 | -0.806904989 | 0.900536389 | -1.666602958 | -3.9542097 |
| 1055 | I-CBP112 (hydrochloride) | 0.737096221 | 0.560865867 | -0.823575449 | -2.204457616 | -0.722238995 | -0.781885116 | 0.752698829 | -0.608634163 | 0.120059637 | 0.990612679 | -0.009762 |
| 1056 | (+)-JQ1 | 0.104057327 | -0.566759284 | -0.209687695 | 0.226246218 | 1.755383272 | -1.370865137 | -2.03231192 | 2.550735496 | 2.649689729 | 6.993592168 | 7.22756059 |
| 1057 | MS023 (hydrochloride) | -0.553990154 | -0.876334981 | 0.64635865 | -0.502898154 | 0.143882488 | 0.905934953 | 1.356123214 | 2.052389902 | 7.104330509 | 5.083097879 | 8.2031126 |
| 1058 | MS049 (hydrochloride) | 1.024230044 | -0.543006557 | -0.284427483 | -0.332780302 | 1.004784303 | -0.235179437 | 1.422020087 | 1.141384187 | 0.484256535 | 1.775756597 | 1.2217416 |
| 1059 | NI-57 | 0.740689324 | -0.701224417 | 0.611648017 | -1.077284428 | 0.155856111 | -1.056552445 | 0.182669969 | 0.166747331 | 0.133448812 | 1.811541176 | 2.08272125 |
| 1060 | NVS-CECR2-1 | 1.286368052 | 0.445181778 | -0.184401105 | -1.272050957 | -1.408595412 | 0.009473132 | 0.076545596 | 0.295276597 | -0.141335743 | -0.841079516 | -0.1346482 |
| 1061 | OF-1 | 0.306309868 | 0.966504986 | 0.047856296 | 0.463134067 | -1.608304926 | -0.729800436 | -0.560582231 | 1.966337687 | -0.087806699 | -0.661254253 | 1.02935463 |
| 1062 | OICR-9429 | 0.823976943 | 0.045890787 | 0.053088645 | -1.248343549 | 0.244284868 | 0.483649222 | -0.243384761 | 0.512366106 | 1.100612584 | -1.731098918 | -0.1120675 |
| 1063 | (R)-PFI-2 (hydrochloride) | 0.885675476 | -0.446368633 | -1.041424253 | -0.655490307 | 0.06291754 | 0.163390525 | -0.513038821 | -0.906181917 | -0.00862421 | 0.0473948 | -0.6528984 |
| 1064 | PFI-3 | -1.100195838 | 1.008760708 | 1.269545285 | 0.335918292 | 0.68751329 | 0.942915867 | -0.362076363 | 0.442098343 | 1.080741939 | -0.508804275 | 0.72901443 |
| 1065 | PFI-4 | -0.764146133 | -0.847774601 | 0.126061424 | 1.141938972 | 0.811781259 | -0.478124813 | -0.132243372 | -1.112244724 | -1.729819061 | -2.259288124 | -2.8144495 |
| 1066 | SGC0946 | 0.294001356 | -0.17765755 | -1.158306814 | -0.502488811 | 0.441245673 | 0.599959043 | -0.174933708 | -1.21561538 | -0.517172743 | -0.496973724 | 0.598244 |
| 1067 | SGC707 | -0.004368018 | -0.411172092 | -0.549606285 | 1.315186853 | 0.086818455 | -0.193772695 | -0.598314471 | -1.712218341 | 1.340909609 | 2.330601455 | 1.7568702 |
| 1068 | SGC-CBP30 | -1.276141422 | 0.263433347 | -1.165088367 | -1.478315922 | 0.556977406 | 0.706041378 | 0.206115434 | 1.894026879 | -0.341617806 | 0.590921947 | 0.10979397 |
| 1069 | TP-064 | -0.011227359 | -1.338539345 | -0.996223531 | 0.331991865 | -0.347037467 | -0.375533112 | -1.637095479 | 1.35573256 | -2.574418744 | 1.220404745 | 1.89504185 |
| 1070 | UNC0642 | 3.193693098 | 0.432044703 | -0.384420517 | 1.954410577 | -0.156748966 | 0.171335676 | 1.704275604 | -1.346034599 | 1.932212338 | 0.013940355 | 1.60339575 |
| 1071 | UNC1215 | 0.634879803 | 1.137677377 | -1.378731778 | 0.200780594 | -0.182529697 | 0.679639641 | 0.442274405 | 0.156559126 | -0.517606561 | 1.430914465 | 0.33693839 |
| 1072 | UNC1999 | 2.300864858 | -0.162197799 | -0.331460088 | -0.732490242 | 0.647470469 | 1.58237566 | -0.19979847 | 0.560791247 | 4.399791947 | 2.54046928 | 1.87989407 |
| 1073 | Acitretin | -0.71147802 | 0.501043601 | -0.559635639 | -0.785566427 | -3.669143287 | -6.50274495 | -11.82359388 | -13.50588607 | -13.31949811 | -13.11835965 | -13.391991 |
| 1085 | Fenretinide | -0.331953584 | -0.841171063 | -1.1066895 | 0.853694966 | -1.345852506 | 0.879955343 | -0.922639952 | -2.149553385 | -5.945068527 | -7.082052829 | -8.9920203 |
| 1092 | BAY-299 | -0.202186515 | -0.558929612 | 0.142652161 | -0.958500602 | -1.529260537 | 1.127565486 | 0.97493051 | 1.254477166 | 1.231808395 | 3.152372742 | 6.85445143 |
| 1093 | GSK6853 | 0.767128778 | 1.368002864 | 1.25718826 | 0.66491413 | -0.406568579 | 0.970906821 | 2.678352591 | 1.08891196 | 0.561524232 | 2.641595529 | 3.05991679 |
| 1094 | TP-472 | -4.418033317 | -0.503895649 | -0.258658193 | 2.256720047 | -0.05129173 | -0.361813085 | 0.07934635 | 1.080631316 | 5.631253784 | 0.259978336 | 0.18101756 |
| 1103 | Bafilomycin | 1.084440757 | 7.008808354 | 9.735062408 | 4.233580082 | 12.97805979 | 6.420821238 | 5.309980991 | 7.232494962 | 7.80179011 | 3.380502094 | 0.20877153 |
| 1108 | Dinitrophenol | 0.327138957 | 5.051869891 | 0.859622229 | 0.48063904 | 1.456640086 | 1.81338034 | 0.030201684 | 1.65284536 | 2.814815664 | -4.387410622 | -1.4186782 |
| 1110 | Echinomycin | 8.7239832 | 1.161169185 | -3.767137237 | -2.265320122 | 1.26238128 | -0.136022539 | 0.543351287 | 2.935397897 | -2.842764751 | -2.277008005 | 4.22531016 |
| 1112 | FCCP | -1.374906302 | -0.492870791 | -1.303462484 | -2.15322014 | -1.215964393 | -2.441237845 | -2.935759746 | -4.273400834 | -1.354786481 | -3.649698204 | -2.1345024 |
| 1113 | Glycogen Phosphorylase Inhibitor | 0.080851536 | 0.037885096 | -0.460490029 | -0.715888883 | -1.844299355 | -0.244841947 | -0.877781291 | -1.865720745 | -0.794456653 | -2.342366282 | -1.1661052 |
| 1117 | Kinetin Riboside | -1.176088233 | 0.503502655 | 4.904195618 | 10.63067247 | 2.995755604 | 4.08424287 | 1.576473453 | 4.137279174 | -0.296905057 | -1.97124509 | -0.4554608 |
| 1119 | LCS-1 (SOD inhibitor) | -0.081183337 | -1.451003849 | -1.592466137 | 0.065262748 | 2.781281844 | -10.05418157 | -5.080421106 | -3.860967749 | -9.86363429 | 0.880131597 | -9.8195333 |
| 1121 | LMK-235 | 1.836163276 | 6.527109737 | 10.15404377 | 7.90993277 | 3.797135111 | -3.792102879 | 0.951944875 | -2.547971716 | 1.796591371 | -0.740003109 | 1.67114142 |
| 1124 | Mdivi-1 | 0.6231515 | -2.205731038 | -1.944485872 | -2.47184492 | -4.052455616 | -2.420961008 | -2.551858239 | -3.838252422 | -1.369259121 | -0.958706854 | -1.7021141 |
| 1129 | NAMPT inhibitor (FK866 compound) | 0.937218185 | 0.345284262 | -0.825162157 | 0.852088729 | -5.325220372 | -5.781974604 | -4.048318005 | -4.69620804 | -8.201521623 | -4.941475108 | -4.9524185 |
| 1131 | Niclosamide | -0.957299891 | -1.563440665 | 0.700735798 | -0.052903531 | -1.191861466 | 0.217605243 | -1.259776096 | 1.181243906 | -1.932716198 | -2.832483718 | -1.0957732 |
| 1134 | Oligomycin | -4.915537138 | -3.783041737 | -4.425128231 | -3.430636785 | -4.413209595 | -3.223312977 | -5.011235661 | -4.142916459 | -3.449833752 | -3.189154912 | -2.4542259 |
| 1135 | PFK15 | 0.13704847 | 0.238082095 | -0.092038578 | 0.714019193 | 1.61045857 | 0.364533045 | -0.976561832 | 4.501537005 | -6.967196035 | -10.25279676 | -9.6861105 |
| 1136 | Phenformin | -1.114298804 | 1.576022454 | -0.438971094 | 0.149125079 | -1.641830742 | -3.958707167 | 1.269352433 | 0.537306425 | -1.774023389 | 0.50745922 | -1.6961818 |
| 1140 | SBI-0206965 | -2.089796142 | 0.445081168 | 1.625613712 | -0.40647038 | -1.210007325 | 2.068955249 | -3.722799887 | 1.560231842 | 4.526140765 | 2.555335833 | 0.82323725 |
| 1141 | Spautin-1 | -1.233370325 | 0.002490436 | -1.115732966 | -2.289321238 | -2.81556997 | -4.727623308 | -5.043665969 | -6.381924494 | -7.360518495 | -4.50807706 | -3.0171348 |
| 1150 | Artesunate | -0.680398363 | -0.30875905 | 0.412892726 | 0.720207416 | 2.801023127 | 9.543168813 | 9.400473143 | -4.759061566 | -3.537481527 | -8.004092846 | -11.131794 |
| 1160 | Dynasore | -1.855650604 | -0.722819938 | -1.108797128 | 0.186924807 | -0.542484356 | 0.568596396 | 2.302845722 | 8.99679733 | 5.5097421 | 2.024918388 | -0.4293612 |
| 1162 | Erastin | 1.227132579 | 0.65431291 | 1.104121625 | 1.237113543 | -0.074384827 | -0.190883863 | 5.376823009 | 1.566691138 | -0.945969199 | -1.290227739 | -1.624783 |
| 1167 | GMX1778 | 1.228876198 | 1.037070957 | 1.369180135 | -4.413309309 | -5.553255617 | -4.990421272 | -6.991657158 | -6.615580318 | -7.253319165 | -5.318396473 | -2.9693138 |
| 1170 | Leflunomide | -0.634792085 | 0.075925575 | -0.435985724 | -1.400789421 | -2.408174745 | -1.929964985 | -0.547756444 | 0.238121314 | -0.368450653 | -2.311405818 | 1.89704949 |
| 1172 | MRT68921 | 0.96608315 | 0.369665533 | 0.674798339 | -1.256519173 | -1.086640257 | 2.446006277 | 1.059569592 | -2.590442228 | -8.406424666 | -11.77127033 | -1.8223489 |
| 1174 | NLG919 | -0.671796986 | 0.824431277 | 0.09217995 | -0.178533605 | -2.001871106 | -1.889265547 | 0.540145424 | 1.06743655 | 2.086015829 | 4.035535447 | -1.5464005 |
| 1175 | Oltipraz | 1.925696722 | -1.014844549 | -0.139822818 | 1.056370918 | 0.018767826 | 2.286332942 | -0.654222695 | -1.483711157 | -0.65267187 | -2.618325174 | -2.5899683 |
| 1177 | PX-12 | -0.232461975 | 2.085975389 | -0.682539247 | 0.468422459 | -0.685105436 | 0.925596567 | 2.602139902 | 2.859419434 | 4.881709474 | -10.86037103 | -1.590413 |
| 1180 | RSL3 | 0.619124638 | 0.987918217 | -0.000591931 | -1.766798848 | -11.51202488 | -5.747797685 | -12.58692438 | -9.457879189 | -13.29335946 | -11.02714851 | -1.1689732 |
| 1183 | SRT1720 | -0.59767202 | 0.517232737 | 0.482246401 | 0.898711352 | 4.231273562 | -0.603377959 | 3.119418526 | -10.04308967 | -6.83186926 | -5.19466993 | 1.19365266 |
| 1185 | STF-118804 | 0.609047432 | 0.112097905 | -3.413181968 | -5.611141069 | -1.588975754 | -7.418019199 | -4.852305613 | -5.86958543 | -6.094343496 | -5.29531679 | -4.4130295 |
| 1186 | STF-31 | 0.793860281 | -0.792590421 | -0.326026972 | -0.255859391 | -0.384366832 | -5.915295042 | -4.177478305 | -6.443682282 | -6.021878527 | -9.190500815 | -7.0187227 |
| 1187 | SW033291 | 0.211488041 | 1.055992945 | 1.147500586 | 0.136833599 | -0.827538406 | -0.223729148 | -2.860741455 | -9.374885912 | -12.5957414 | -10.08105968 | -12.409831 |
| 1188 | Thiabendazole | -2.232586996 | -0.251421109 | -1.276980408 | -2.029600797 | -1.574926211 | -1.241304048 | -0.416611461 | 1.263899917 | 0.900438991 | 1.677636186 | 4.13378665 |
| 1192 | Cerulenin | 0.325306996 | 1.19426619 | -0.700678681 | -1.512554223 | 0.419947748 | 0.784985389 | 0.94522131 | 3.626624541 | 3.133529 | -8.230969239 | -10.149895 |
| 1196 | Thapsigargin | 0.759383328 | -8.378823137 | -10.93419008 | -11.95018503 | -9.325349286 | -7.153309245 | -3.510331101 | -2.859133286 | -1.629178531 | 1.564416767 | -2.4335481 |
| 1197 | TPEN | -1.89790248 | 0.452610285 | 0.585872713 | -0.908712296 | -0.147323491 | 1.19207244 | -0.417827637 | 1.28813159 | -1.980367422 | -0.94290091 | -3.1674189 |
| 1199 | Mycophenolic acid | 0.063307162 | -1.350382662 | 2.826177585 | 3.711775033 | 7.214910588 | 3.073745819 | 0.155999378 | -1.624531439 | -3.749889064 | -4.035257492 | -3.4300589 |
| 1203 | Doxycyline | 0.329857923 | -0.931410068 | 0.633421532 | -0.131118823 | 2.626882889 | 2.761993151 | 0.147358775 | -9.347633598 | -8.369647175 | -7.814150091 | -5.4841018 |
| 1204 | Auranofin | 0.47086749 | -1.060484978 | -0.506766557 | -0.265398804 | -1.268505635 | 0.015358427 | 1.749106728 | 5.714456509 | -7.310856428 | -10.05272386 | -8.3189119 |
| 1205 | TRO-19622, Olesoxime | 0.615814705 | 0.046109757 | -0.051324375 | 1.224971646 | 0.769487982 | 1.489258029 | -1.66213418 | -0.366044396 | 1.629348883 | 0.019641328 | -0.2543518 |
| 1206 | LRRK2-In-1 | -0.187051803 | -1.121365367 | -1.197453159 | -0.568164509 | -0.562765691 | -3.041108896 | -0.113575058 | 2.005626488 | 4.73079504 | -1.296070276 | -0.5356961 |
| 1208 | 2'-3'-dideoxycytidine | -0.033261774 | 0.566021918 | -0.916949731 | 0.28424175 | 0.015715711 | -0.390845552 | -1.165578701 | -0.847318464 | -4.030101368 | -5.280970631 | -5.5720899 |
| 1209 | Mitoblock6 | -0.741241587 | 1.404624859 | -0.554632909 | 0.378208856 | -2.137259817 | -2.71223766 | -5.190241545 | -7.433183942 | -4.765797816 | -1.377145456 | -4.9842443 |
| 1210 | Avocatin b | -0.743835465 | 0.589169503 | -0.549424324 | -0.880177299 | -10.09007344 | -10.28922661 | -11.00514623 | -8.104039048 | -6.892701996 | -5.753764728 | -4.269859 |
| 1211 | A-1155463 | 1.230038791 | 3.730930172 | 4.056517111 | 2.692101594 | 2.963502862 | -1.234970949 | 2.167371575 | 0.80718118 | -0.665498276 | -0.892164869 | -2.1698635 |
| 1213 | DASA-58 | 1.030522944 | 3.638239387 | 5.202367626 | 13.36255668 | -3.277961134 | -7.447045888 | -2.809258023 | -3.11467216 | -1.711083831 | 3.484349982 | -2.8938251 |
| 1216 | Vidofludimus | -0.181864101 | -0.041350041 | -0.815717839 | -0.253391274 | 0.43196367 | -0.197825689 | 0.220616161 | 3.923666723 | -0.07951724 | -4.614772193 | -7.4911674 |
| 1217 | Concanamycin | -0.181695598 | 3.588644467 | 6.447878004 | 7.182442948 | 3.641810253 | 2.316362356 | 3.987743274 | 6.560228703 | 3.078142294 | 9.155711231 | 6.15870736 |
